# Large differences in carbohydrate degradation and transport potential in the genomes of lichen fungal symbionts

**DOI:** 10.1101/2021.08.01.454614

**Authors:** Philipp Resl, Adina R. Bujold, Gulnara Tagirdzhanova, Peter Meidl, Sandra Freire Rallo, Mieko Kono, Samantha Fernández-Brime, Hörður Guðmundsson, Ólafur Sigmar Andrésson, Lucia Muggia, Helmut Mayrhofer, John P. McCutcheon, Mats Wedin, Silke Werth, Lisa M. Willis, Toby Spribille

## Abstract

Lichen symbioses are generally thought to be stabilized by the transfer of fixed carbon compounds from a photosynthesizing unicellular symbiont to a fungus. In other fungal symbioses, carbohydrate subsidies correlate with genomic reductions in the number of genes for plant cell wall-degrading enzymes (PCWDEs), but whether this is the case with lichen fungal symbionts (LFSs) is unknown. We predicted genes encoding carbohydrate-active enzymes (CAZymes) and sugar transporters in 17 existing and 29 newly sequenced genomes from across the class *Lecanoromycetes*, the largest extant clade of LFSs. Despite possessing lower mean numbers of PCWDE genes compared to non-symbiont *Ascomycota*, all LFS genomes possessed a robust suite of predicted PCWDEs. The largest CAZyme gene numbers, on par with model species such as *Penicillium*, were retained in genomes from the subclass *Ostropomycetidae*, which are found in crust lichens with highly specific ecologies. The lowest numbers were in the subclass *Lecanoromycetidae*, which are symbionts of many generalist macrolichens. Our results suggest that association with phototroph symbionts does not in itself result in functional loss of PCWDEs and that PCWDE losses may have been driven by adaptive processes within the evolution of specific LFS lineages. The inferred capability of some LFSs to access a wide range of carbohydrates suggests that some lichen symbioses may augment fixed CO_2_ with carbon from external sources.

**Significance:** Lichen symbioses are considered self-contained autotrophic systems in which the total carbon economy is the sum of phototroph-fixed CO_2_, supplied to a fungus as sugars. In other fungal-plant symbioses, such as mycorrhizae, plant-derived sugar subsidies are associated with loss of plant cell wall-degrading enzymes (PCWDEs). We compared PCWDE inventories in 46 genomes from the largest group of lichen fungal symbionts (LFSs) with non-symbionts from across *Ascomycota*. We found that despite lower overall gene numbers, all LFSs retain PCWDEs, and some possess gene numbers and functional diversity on par with non-symbionts. Our results suggest that association with a phototroph does not necessarily result in PCWDE loss, and some lichens may obtain carbon from sources other than CO_2_ fixation.

## Introduction

Stable fungal associations with single-celled photosynthetic organisms, usually referred to as lichens, feature prominently in the history of the discovery and study of symbiosis. In describing the pairing of fungi with algae and/or cyanobacteria within lichens for the first time, the Swiss botanist Simon Schwendener proposed that lichen fungal symbionts derive nutrition from “assimilates” of their photosynthesizing symbionts (1). Almost a hundred years later, Smith and colleagues revealed these transferred photosynthates to be polyols and glucose in the case of algae and cyanobacteria, respectively (2). They and others traced the transfer of algal fixed carbon into fungal cells, where they found it to be converted into mannitol and arabitol (3, 4). The fungal-photosymbiont relationship is widely interpreted as conferring net independence from external carbohydrates on the resulting lichen thallus. Accordingly, lichen fungal symbionts have been classified as biotrophs (5), and the symbiotic outcome, the lichen thallus, as a “photosynthetic carbon autotroph” (6) or “composite autotroph” (7).

Fungi are assimilative heterotrophs and thus require a robust machinery of enzymes for scavenging and transporting extracellular nutrients, including carbohydrates. In arbuscular mycorrhizal and ectomycorrhizal fungi, the stable supply of glucose from plants is thought to have led to erosion or loss in many families of plant cell wall-degrading enzymes (PCWDEs) (8, 9), reflecting a common pattern of compensated trait loss in symbioses (10). So what happened to PCWDEs in lichen fungal symbionts? Multiple lines of indirect evidence have emerged over the last 40 years to suggest the retention of PCWDEs in lichen fungal symbionts (LFSs) or their secondary evolutionary derivates. First, molecular phylogenetic studies have shown multiple independent origins of putative saprotroph lineages from lichen fungal symbiont ancestors, both ancient (11) and recent (12,13,14). How these newly evolved lineages acquired the carbohydrate breakdown arsenal they would presumably need for life without an alga has not been explained. Second, some fungi near the symbiont-to-saprotroph transition have been shown to switch between the two lifestyles facultatively, so-called “optional lichens” (15, 16); in these cases, the fungus appears not to be obligately dependent on the alga for nutrition. Third, many lichen symbioses exhibit anomalous “substrate specificity”, i.e., they are restricted to specific organic substrates and unable to colonize others, suggesting lack of nutritional autonomy (17, 18, 19). Fourth, lichen fungi are capable of growing axenically *in vitro* on a variety of sugars other than sugar alcohols, including crystalline cellulose and sucrose (reviewed by 20). Finally, enzymes involved in breakdown of lichen-exogenous carbohydrates have been isolated from lichens in nature, including cellulases and lignolytic peroxidases (reviewed by 21). These phenomena are easy to treat as exceptions, but their distribution across the fungal symbiont tree hints at deeper underlying fungal capabilities, which if combined with phototroph symbiosis could lead to a kind of “hybrid” lifestyle, in which carbohydrates are obtained from multiple sources. The possibility of multiple carbon sources for lichens was even suggested by Schwendener himself in his original 1869 paper, in which he predicted that two tracks of nutrient acquisition would ultimately be proven: one for lichens that have minimal substrate contact, which he predicted to depend mostly on algal assimilates; and one for lichens that closely hug organic substrates such as tree bark or wood (1).

Even with these results and hypotheses, however, the possibility of carbon assimilation from sources other than the phototroph symbiont is still generally discounted in lichen symbiosis research. Lichen ecophysiologists currently calculate total lichen carbon budgets based on the rate and total amount of algal carbon fixation (6, 22). Implicit to this, though to our knowledge never explicitly stated, is that lichen fungal symbionts must lack the ability to break down lichen-exogenous carbohydrates, i.e., carbohydrates found neither in the lichen fungus nor the phototroph symbiont.

Unlike for many fungi, phenotypic carbohydrate use profiles have seldom been developed for LFSs. This is in part due to the recalcitrance of most LFSs to culturing and their extreme slow growth, if culturing is successful. Knowledge gaps around unculturable or slow-growing fungi are common, but have been offset in recent years by genome sequencing. Coupled with the development of widely available databases such as CAZy (23), it has become possible to infer carbohydrate active enzymes (CAZymes) for species for which a genome, but no experimental evidence, is currently available. Comparative genomic overviews of CAZyme repertoires are now available for many symbiotic fungi (9), but no survey exists of comparative CAZyme arsenals in LFSs.

Given the common assumption that lichen symbiont complementarity confers collective autotrophy on the symbiosis, and historical assumptions that they evolved from saprotrophic ancestors, we hypothesized that LFSs would exhibit functional losses in PCWDEs coinciding with the beginning of stable association with phototroph symbionts, similar to what has been found in mycorrhizal fungi (9,24). To test this, however, we would need to map CAZymes across a well-sampled phylogeny and reconstruct the ancestral state of the last common ancestor, which is considerably older than many of the origins of e.g. ectomycorrhizal fungi (25). Here we map the occurrence of genes encoding PCWDEs at two levels: across *Ascomycota* including the origin of the *Lecanoromycetes*, the largest extant lineage of LFSs; and across representatives of major groups within *Lecanoromycetes*, representing different ecological substrate specificities (specialists and generalists) as well as major morphological outcomes of the lichens they occur in (crusts, macrolichens). Our survey of 46 lecanoromycete genomes, 29 of which we sequenced for this project, reveals a complex pattern of retention and loss that lends support to Schwendener’s hypothesis of hidden saprotrophy in some lichens and is not unequivocally consistent with PCWDE erosion upon acquisition of phototroph symbionts.

## Results

### Data set and phylogenomic reconstruction

We assembled a data set of 83 fungal genomes from the phylum *Ascomycota*, including 46 from the class *Lecanoromycetes* (Supplementary Table 1). Because the few published lecanoromycete genomes are not representative of deep evolutionary splits in the group, we generated 29 new genomes for this study, including 18 as metagenome-assembled genomes (MAGs; Supplementary Table 2). Completeness and quality metrics were comparable for genomes derived from culture and MAGs (Supplementary Figures 1 and 2). Phylogenomic analysis based on 1310 inferred universally present single-copy orthologs (Fig. 1; Supplementary Figure 3 and Supplementary Table 7) recovered major clades and sister group relationships found in recent studies, both among class-level clades of *Ascomycota* (26) as well as within the *Lecanoromycetes* (27, 28). For each genome, we performed *ab initio* gene predictions and obtained functional annotations (CAZymes, InterPro IDs, Pfams). To these we assigned activity on the common plant cell wall substrates cellulose, hemicellulose, lignin and pectin following (9).

**Figure 1.**
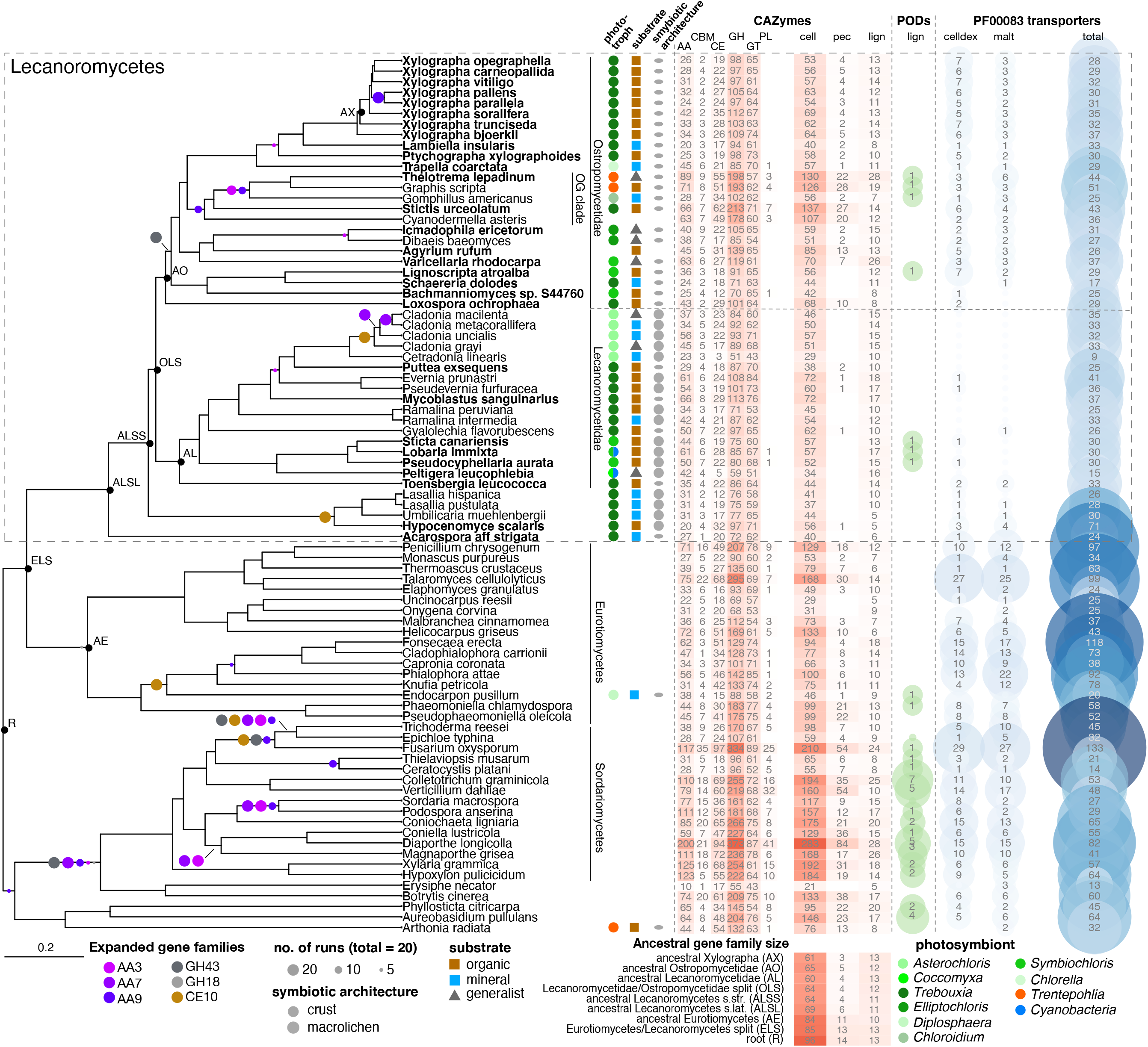
Distribution and ancestral states of CAZymes and selected sugar transporters across the evolution of *Lecanoromycetes* and related classes of *Ascomycota* projected onto a maximum likelihood phylogenomic tree based on 1310 loci. Symbols beside tree tips refer to life history traits and phototrophic partners of LFS under study. Heatmaps with shades of red indicate the number of genes in different CAZyme classes or involved in degrading complex PCW components. Columns from left to right: AA - Auxiliary Activities, CBM - Carbohydrate binding module, CE - Carbohydrate Esterases, GH - Glycoside Hydrolases, GT - Glycosyl Transferases, PL - Polysaccharide Lyases. cell - Number of genes in 35 CAZyme families involved in cellulose and hemicellulose breakdown. pec - Number of genes in 11 CAZyme families involved in pectin breakdown. lign - Number of genes in 3 CAZyme families and class II peroxidases involved in lignin breakdown. Selection of CAZyme sets follows (9, 24). celldex - Number of cellodextrin transporters. malt - number of maltose transporters. total - Total number of sugar transporters. Below the heatmap are ancestral sizes of CAZyme families involved in (hemi-)cellulose, pectin and lignin degradation. Colored circles on tree branches indicate significantly expanded CAZyme families. The size of the circles indicates the number of individual CAFE runs in which a family was found to be significantly expanded.

### CAZymes

Many CAZyme families are shared across *Ascomycota*. Glycosyl transferases (GTs), which are involved in glycosylation and the synthesis of polysaccharides, differ little across all analyzed genomes and do not exhibit any significant reduction in *Lecanoromycetes*, suggesting that a core *Ascomycota* synthetic machinery remains largely unchanged across the evolution of these groups (Fig. 1). While significant within-group variation exists, the mean number of glycoside hydrolase (GH) genes in lecanoromycete genomes is 40.7% lower compared to other *Ascomycota* genomes (p=0; Supplementary Table 9, 10). CAZymes with Auxiliary Activities (AA), Carbohydrate Binding Modules (CBM) and Carbohydrate Esterases (CE) are also reduced significantly in *Lecanoromycetes* (Supplementary Table 10). The bulk of these differences can be attributed to a small number of CAZyme families (Supplementary Figure 11), most of which are plant cell wall degrading enzymes (PCWDEs). Additionally three CAZyme families widespread in *Ascomycota* were not detected in *Lecanoromycetes*. These include PL4, which contains pectin degrading rhamnogalacturonan endolyases; CBM67, which binds to L-rhamnose and frequently occurs in multi-domain protein with enzymes in GH78 and PL1; and AA13, which contains lytic polysaccharide monooxygenases (LPMOs) involved in starch breakdown. In contrast to these reductions, numerous CAZyme families are not reduced at all in *Lecanoromycetes* compared to other *Ascomycota*. Indeed some, including those involved in degradation of endogenous fungal cell wall polysaccharides, such as GH128 and AA5, are even expanded in *Lecanoromycetes* (Supplementary Figure 11).

All lecanoromycete genomes retained genes encoding enzymes predicted to act on plant cell wall carbohydrates, including cellulose and hemicellulose (e.g., GH5 and 43), lignin (AA1, 2 and 5). Depending on the symbiont configuration, these carbohydrates may be produced by the phototroph partner and/or be exogenous to the lichen symbiosis (see Discussion). Phylogenetically corrected principal components analyses (PCA) of CAZyme family numbers in the sampled genomes reveals differences in the amount of variation in CAZyme composition among lecanoromycete and other ascomycete genomes (Fig. 2). For CAZymes predicted to act on cellulose, hemicellulose and pectin, the number of predicted gene families varies less among lecanoromycete genomes than among comparable classes of Ascomycota (Fig. 2A, 2C), reflected in tight clustering in *Lecanoromycetes* versus wide scattering in other ascomycete classes. For lignin-degrading enzymes the variation is similar (Fig. 2B). Within Lecanoromycetes, however, the greatest amount of variation in predicted gene sets is exhibited in the subclass *Ostropomycetidae* in cellulose/hemicellulose (Fig. 2A) and pectin degradation (Fig. 2C), resulting in scattering in the PCA ordination. Genomes from the subclass *Lecanoromycetidae*, by contrast, possess similar predicted CAZyme sets, resulting in tight clusters in the PCA ordination.

**Figure 2.**
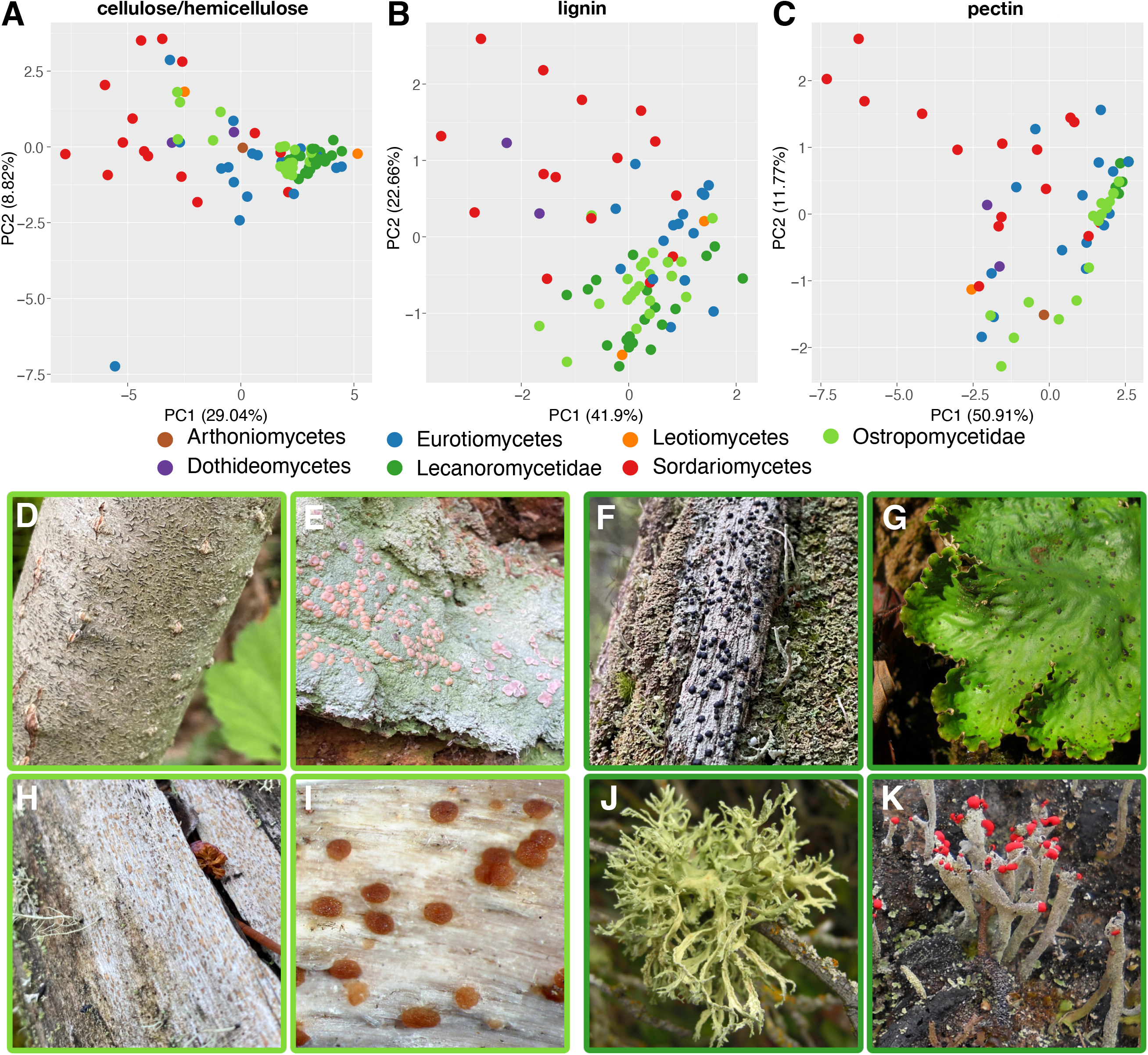
Similarity of CAZyme sets involved in the breakdown of different complex plant-based polysaccharides based on phylogenetically corrected Principal Components Analysis. Different colors indicate taxonomic groups. A - Similarity of CAZyme families involved in cellulose and hemicellulose breakdown. B - Similarity of CAZyme families involved in lignin breakdown, C - Similarity of CAZyme families involved in pectin breakdown. Displayed below are representative members of the two subclasses *Ostropomycetidae* (with light green border) and *Lecanoromycetidae* (dark green border). D - *Graphis scripta*, E - *Icmadophila ericetorum*, F - *Mycoblastus sanguinarius*, G - *Peltigera leucophlebia*, H - *Xylographa carneopallida*, I - *Agyrium rufum*, J - *Evernia prunastri*, K - *Cladonia macilenta*. Image credits: *Agyrium rufum*: Paul Cannon (fungi.myspecies.info); Creative Commons: BY-NC 4.0. *Peltigera leucophlebia*: Jason Hollinger, uploaded by Amada44, CC-BY 2.0, https://commons.wikimedia.org/w/index.php?curid=24213606. *Evernia prunastri*: by Jason Hollinger, CC-BY 2.0, https://commons.wikimedia.org/w/index.php?curid=50595319. *Cladonia macilenta*: Bruce McCune & Sunia Yang - Lichen, CC-BY 4.0-NC, https://lichens.twinferntech.net/pnw/species/Cladonia_macilenta.shtml; other images by the authors.

Genomes from the *Ostropomycetidae* possess higher numbers of predicted CAZyme genes that act on cellulose and hemicellulose than those of *Lecanoromycetidae*, which are reflected in significantly higher numbers of GHs and CEs (p=0 and p=0.0024 respectively; Supplementary Table 10). The differences are driven by one lineage in particular, represented by five genomes from the orders *Ostropales* and *Gyalectales* and referred to here as the OG clade (Fig. 1), and are much less if the two suborders are compared without the OG clade genomes. The OG clade contains numbers of predicted CAZyme genes for cellulose and hemicellulose breakdown that are over fourfold more numerous than the lowest lecanoromycete numbers, which are in *Cetradonia* and *Peltigera*, and equal or exceed the numbers in well-studied eurotiomycete saprotrophs such as *Penicillium*. These disparities are accounted for in large part by gene assignments to two CAZyme families, GH5 and GH43. Because both of these are large, heterogeneous families that include multifunctional CAZymes, we mapped putative GH5 and GH43 orthologs from the analyzed genomes against sequences of experimentally validated enzymes and predicted which are secreted (Fig. 3). Extensive gene duplication in *Ostropomycetidae* is found in gene sequences close to characterized cellulases (GH5 subfamily 5; EC 3.2.1.4) and from both lecanoromycete subclasses in sequences close to characterized 1,3-beta-glucosidases (GH5 subfamily 9; EC 3.2.1.58) and endo-1,4-beta-mannosidases (GH5 subfamily 7; EC 3.2.1.78). Notably, many putative lecanoromycete CAZyme genes from GH5 do not closely cluster with any characterized sequences and form their own clades with sequences from other classes of *Ascomycota* (Fig. 3A).

**Figure 3.**
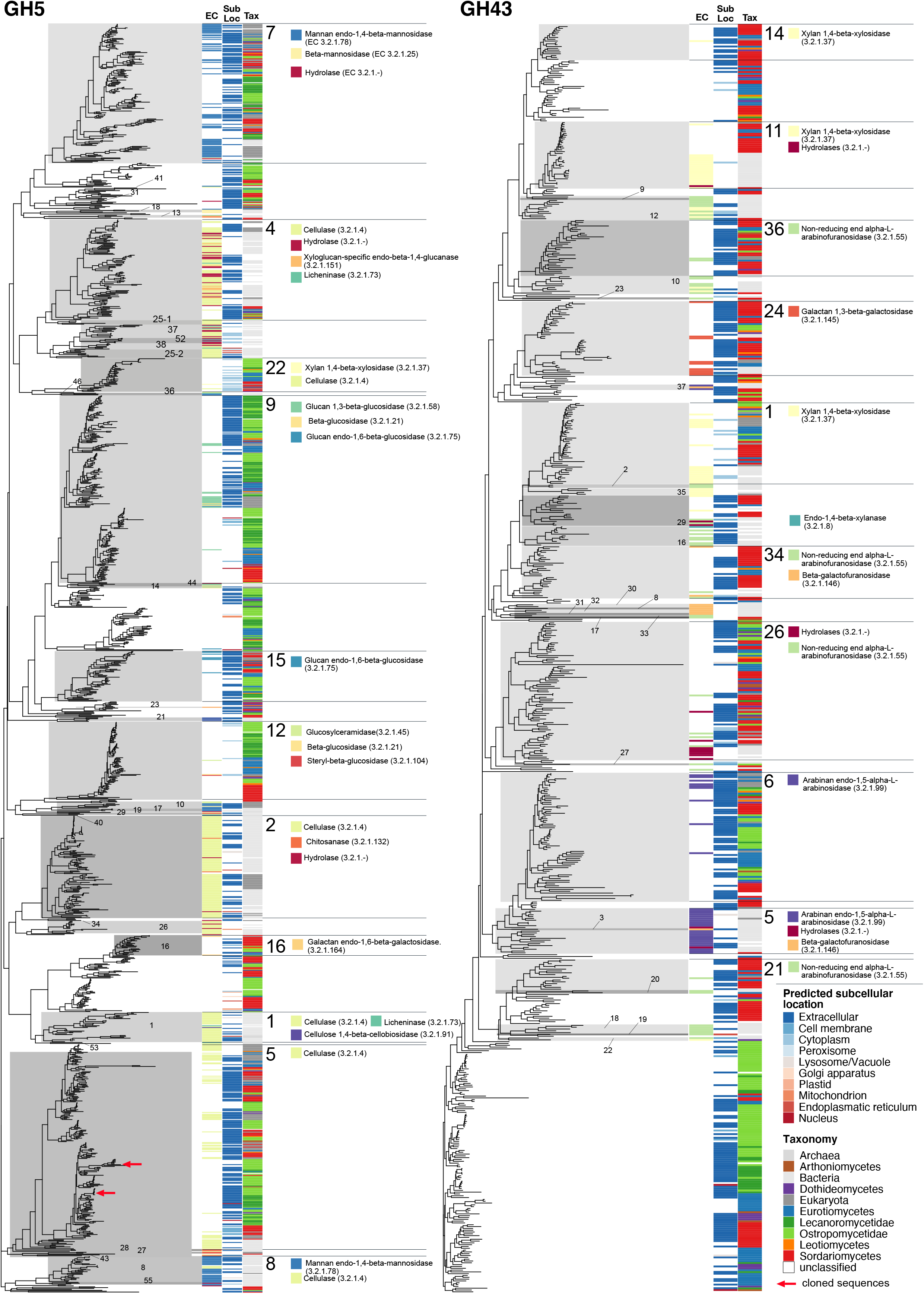
Gene trees of two CAZyme families involved in cellulose (GH5) and hemicellulose (GH43) breakdown. Each tree includes all experimentally characterized sequences combined with all sequences from the 83 genomes studied here. CAZyme Subfamilies are labeled with numbers and grey rectangles. Three columns along tree tips display additional information of corresponding sequences when available. EC - Enzyme Code of experimentally characterized sequences downloaded from cazy.org. Sub Loc - Predicted subcellular location of Enzyme with DeepLoc. Tax - Taxonomic assignment of organisms from which the sequence comes from. For larger subfamilies, functions of characterized sequences based in Enzyme Code numbers are given as colored squares. Sequences used for heterologous expression experiments are marked with red arrows in GH5 subfamily 5.

The subclass *Ostropomycetidae* also possesses a larger proportional representation of genes coding for enzymes predicted to be involved in hemicellulose, specifically xylan, breakdown. Families GH6, GH7, GH11 and GH62, GH67 and GH131 were absent in *Lecanoromycetidae* and are only present in *Ostropomycetidae* in the OG clade and the saprotroph *Agyrium* (Supplementary Figure 11). Hemicellulose breakdown also involves several CEs of which CE2 is present in the majority of *Ostropomycetidae* genomes and absent in *Lecanoromycetidae* (Supplementary Figure 11). CE2 contains acetylxylan esterases (AXEs; (29)); acetylxylan is a major component of hemicellulose. An analysis of predicted GH43 gene sequences and their subcellular location compared to those of characterized enzymes shows that most characterized GH43s are in *Ostropomycetidae* (subfamilies 1, 6, 24, 26 and 36), but as with GH5s, the majority of sequences are not close to any characterized CAZymes (Fig. 3B).

CAZymes involved in lignin degradation are frequently characterized in fungi as encompassing Class II peroxidases (PODs) and selected oxidative CAZymes (17). In *Lecanoromycetes*, CAZymes associated with lignin degradation are predicted in all lecanoromycete genomes, but again the highest numbers are found in *Ostropomycetidae* (Fig. 1). The numbers of predicted genes in *Graphis* and *Varicellaria* are among the highest in the analyzed *Ascomycota* genomes. Most of these genes are members of AA1 and AA3 which contain laccases (EC 1.10.3.2; AA1; Supplementary Figure 9) and glucose-methanol-choline oxidoreductases in three subfamilies involved in lignocellulose degradation (Supplementary Figure 9). Class II PODs, by contrast, are virtually absent across *Lecanoromycetes* (Fig. 1).

The largest differences in predicted PCWDEs among lecanoromycete genomes come from CAZymes acting on pectins. All but three of the *Ostropomycetidae* genomes possess a few predicted CAZymes involved in pectin degradation (Fig. 1). These include GHs (GH28, 49, 53, GH79, GH108), CEs (CE8, CE12, CE15) and the only polysaccharide lyases (PL; from PL1 and PL3, containing pectate lyases, EC 4.2.2.2) predicted in *Lecanoromycetes* (Supplementary Figure 8). *Lecanoromycetidae*, by contrast, almost completely lack gene predictions for pectin degradative enzymes and completely lack PLs. Similar to CAZyme predictions for cellulose, hemicellulose and lignin degradation, the largest numbers of predicted genes involved in pectin degradation come from the OG clade of *Ostropomycetidae*. In these genomes, predicted gene numbers for CAZymes involved in pectin degradation even exceed those of known model saprotrophs in *Eurotiomycetes*, though the difference is not significant.

In addition to the PCWDEs outlined above, all sampled *Lecanoromycetes* possess a gene assigned to GH32, a family which includes invertases involved in the conversion of sucrose to glucose or fructose. Invertases are generally interpreted as an indicator of utilization of apoplastic plant sucrose. However, lecanoromycete GH32s do not cluster closely with any characterized invertases (Supplementary Figure 13).

### Predictions of gene family contraction and expansion

In order to establish whether CAZyme patterns in *Lecanoromycetes* are due to gene gain or loss, we reconstructed ancestral gene numbers for each of the three main groups of PCWDEs (cellulose/hemicellulose, pectin, lignin). In general, the pattern reflects sequential gene loss. All three PCWDE groups are reduced already at the split between *Eurotiomycetes* and *Lecanoromycetes* (ELS node; Fig. 1), especially enzymes involved in (hemi-)cellulose and pectin breakdown. Much of the signal for this pattern comes from GH5 and GH43 which contain many well-characterized cellulases and hemicellulases, respectively (Fig. 2; Supplementary Figure 7). Another apparent shedding of CAZymes, especially of those involved in pectin degradation, occurs concomitantly with the appearance of LFSs at the base of *Lecanoromycetes* (ALSS and ALSL nodes; Fig. 1). Here, too, the signal comes mainly from GHs. In several cases our ancestral state reconstruction indicates complete losses, despite the prediction of several of the relevant families in *Ostropomycetidae* genomes (Supplementary Figure 8).

Gene family expansion analyses of all CAZymes revealed six significantly expanded families along different branches in the phylogeny. The only GHs expanded within Lecanoromycetes are GH43, which are expanded at the last common ancestor of *Ostropomycetidae* (Fig. 1); another, GH18, was expanded in four model runs at the base of *Sordariomycetes*. Other significantly expanded CAZyme families in *Lecanoromycetes* include Auxiliary Activity (AA) families AA3, AA7 and AA9, which are variously expanded in *Xylographa*, the OG clade and *Cladonia*. Auxiliary Activity families typically act in concert with other CAZymes and the three expanded AA families all contain genes involved in cellulose breakdown. The only other expanded CAZyme family was CE10, now considered to be a family of esterases acting on non-carbohydrate substrates, which was expanded at the last common ancestor of *Cladonia* species and *Umbilicariomycetidae*.

### Transporters

As an additional line of evidence for potential use of exogenous carbohydrates, we mapped the numbers of predicted sugar transporters across all genomes (Fig. 1). Most predicted transporters were more or less evenly represented across most genomes, but two groups of transporters exhibited a concerted pattern of absence within *Lecanoromycetes*. Cellodextrin transporters, close orthologs of experimentally demonstrated cellodextrin transporters from *Aspergillus* and *Penicillium* (Supplementary Figure 5), were predicted in all *Ostropomycetidae* genomes except *Schaereria*, and were most numerous in *Lignoscripta*, *Stictis*, *Ptychographa* and *Xylographa*, all of which are LFSs of lichens with high wood or bark specificity (Fig. 1). Cellodextrin transporters are involved in transmembrane import of cellobiose and other cellodextrins, which are short beta-linked glucose fragments of cellulose. Only nine of the remaining 21 lecanoromycete genomes had any predicted cellodextrin transporters, and in these, gene numbers were well below the average observed in *Ostropomycetidae*. We detected a similar pattern for the predicted occurrence of maltose transporters (Fig. 1), involved in transmembrane import of alpha-linked starch breakdown products: 22 of 24 Ostropomycetidae genomes had predicted maltose transporters, but only five lecanoromycete genomes outside of *Ostropomycetidae* had any, mostly in early diverging lineages.

### Evidence of cellulase functionality

To validate the functionality of putative cellulases found in lichens, we selected *X. bjoerkii* genes *Xylbjo000518* and *Xylbjo004565*, hereafter called *cellulase A* and *B*, for further analysis based on their similarity with Cel5A cellulases, for which there has been substantial structural and functional characterization. The cellulase domain of each gene was expressed in *Escherichia coli* as a C-terminal fusion with the maltose binding protein, which enhances protein expression and stability and provides a convenient handle for purification. The partially purified proteins were active on both cellulose (β-1,4-linked glucose) and barley β-glucan (alternating β-1,4/β-1,3-linked glucose) but not xylan (β-1,4-linked xylose containing side branches of α-arabinofuranose and α-glucuronic acids) (Fig. 4A-C). Both exhibited a pH optimum of 5, similar to other Cel5A enzymes but differ slightly in their temperature optima.

**Figure 4.**
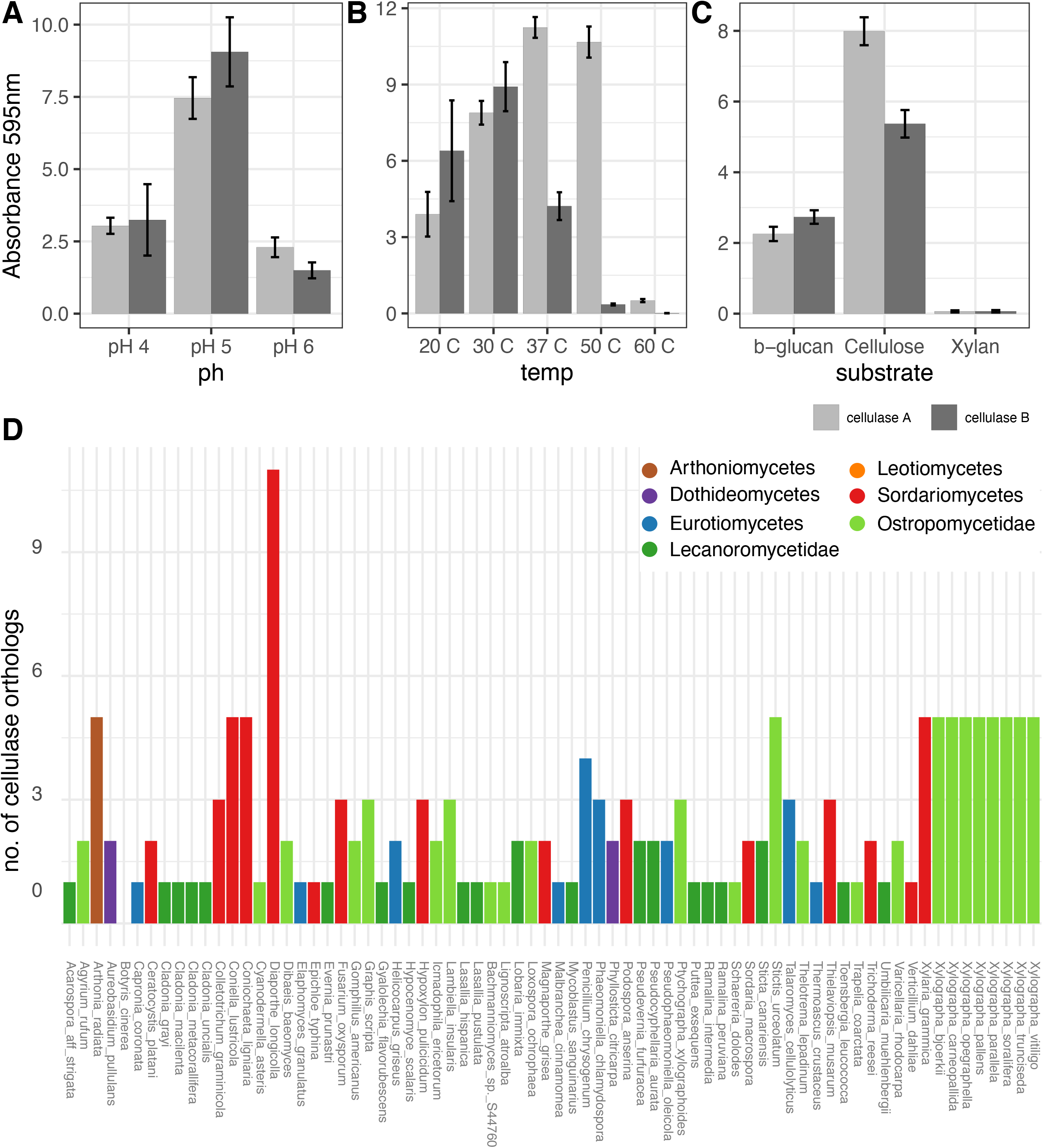
Enzymatic activity of two putative cellulases A and B from *Xylographa bjoerkii* and orthologues of these genes in 83 ascomycete genomes. A - Activity of cellulase A and B at different pH conditions. B - Activity of cellulase A and B at different temperatures. C - Activity of cellulase A and B on different substrates. D - Number of orthologs of cellulase A and B in different genomes.

While both are active at 20 °C, only *cellulase A* is also active at higher temperatures. Closely related orthologs of *cellulase A* and *cellulase B* were recovered in all lecanoromycete genomes. *Lecanoromycetidae* mostly only had a single ortholog with the exception of *Lobaria*, *Pseudocyphellaria*, *Pseudevernia* and *Sticta* which each have two orthologs (Fig. 4D). In contrast, the majority of *Ostropomycetidae* genomes have two or more orthologs, and the highest numbers can be found in *Xylographa* species and *Stictis* which each have five (Fig. 4D).

## Discussion

Our analysis of CAZyme and sugar transporter genes paints a picture of a lecanoromycete PCWDE arsenal that is larger, more diverse and more consistently present than expected. Understanding when and where in the fungal mycelium the PCWDEs are deployed will be critical to determining their biological significance. Our data show a reduction in mean numbers of lecanoromycete degradative CAZyme genes relative to other *Ascomycota*, but also reveal significant differences within *Lecanoromycetes*. Genes for the breakdown of cellulose, hemicellulose and pectin are disproportionately enriched in the subclass *Ostropomycetidae*, and some genomes, notably in the OG clade, retain overall CAZyme numbers and functionality similar to those of well-known model saprotrophs such as *Aspergillus* and *Penicillium*. Furthermore, LFSs associated with the same genus of algal phototroph, *Trebouxia*, can retain multiple genes coding for pectin degradation as well as cellodextrin transporters (in *Lambiella*, *Loxospora*, *Ptychographa* and *Xylographa*), or they can largely lack these genes (in *Ramalina*). This implies that association with *Trebouxia* does not in itself result in gene loss, and the exact nature of the fungal-phototroph interaction, and other aspects of symbiosis biology, may need to be considered.

A central question remains: what are the targets of LFS PCWDEs? The two most obvious candidates are the phototroph itself, on the one hand, and lichen-exogenous plant tissues, such as wood and tree bark, on the other. The first possibility — targeting of the alga — echoes suggestions in studies of ectomycorrhizal fungi, where PCWDEs have been postulated to play a role in “cell softening” of root tissues in their vascular plant symbionts upon contact initiation (30). Cellulases and other CAZymes have been postulated to be involved in haustorial penetration of the algal cell wall (31, 32) and degradation of algal cell walls in fresh lichen growth tips (33). They could also play a role in digestion of dead algal cell walls, especially if algal populations turn over during the “lifespan” of the thallus, as has long been suspected (34). Transcriptomic studies of isolates, co-cultures and natural lichens could provide evidence of this. However, the one study to date to report CAZyme differential expression (35) only detected upregulation of multifunctional GHs that could also be involved in fungal cell wall modification (GH2 and GH12), but none of the core lecanoromycete cellulases or hemicellulases we report here. Most common algal symbionts are thought to contain cellulose in their cell walls but no pectin (36); it is unclear if any lichen algal symbiont possesses pectin in its cell walls.

The second possibility, that LFS PCWDEs target lichen-exogenous plant polysaccharides, is supported by both experimental evidence and inference. Cellulases as well as polygalacturonases, active on pectins, have been detected in incubated whole lichens both in the presence (37, 38) and absence (39) of cellulose-containing algal photosymbionts. Cellulase production furthermore has been reported to vary depending on the species of tree it is growing on (40). Activity has also been demonstrated for laccases and different types of peroxidases (21, 41, 42). Several secondary origins of putative fungal saprotrophs occur within the *Lecanoromycetes*, some of which are closely related to fungi involved in “optional lichens”, in which the fungus can occur either as an LFS or saprotroph (43). Our sampling shows cellulase gene family expansions within the latter group, especially in LFSs of wood-obligate lichens such as *Ptychographa* and *Xylographa*, and our experimental evidence for cellulase activity derives from one of these genomes (*Xylographa bjoerkii*). A further indication of lecanoromycete use of exogenous carbohydrates could be the presence of invertases. The loss of invertases in ectomycorrhizal fungi is thought to limit their ability to access plant sucrose and reinforce their dependence on a plant partner (9). Though the orthologs we found did not exhibit high similarity to characterized sequences, the prediction of a GH32 is consistent with the ability to culture LFSs on sucrose (44) and experimental evidence of invertases in lichen fungi (20, 45).

PCWDE targeting of algal cell walls or lichen-exogenous plant tissues are not mutually exclusive possibilities. However, determining where and when these genes are expressed in nature will require consideration of the life cycle and spatial extent of LFS mycelia, both of which are still poorly understood. The mycelia of sexually reproducing LFSs go through an aposymbiotic stage of unknown length, during which it has been suggested they may be saprotrophic (46). Lichens at high latitudes or under snowpack may go through seasonal fluctuations in fixed carbon input, which could be augmented by other sources (21). Even after symbiosis is established, parts of the mycelium can be free of phototroph cells, and exhibit deviations in carbohydrates that suggest other metabolic processes are in effect than in the phototroph-associated mycelium (47). Many lichens include a phototroph-free “prothallus”, in which fungal hyphae radiate beyond the zone of phototroph cells (48). Others possess a “hypothallus”, a phototroph-free cushion of mycelium in direct contact with the substrate (47). Macrolichens are often anchored onto their substrates by phototroph-free “holdfasts”, “rhizines” or mycelial pegs which have traditionally been interpreted as having an exclusively structural, stabilizing function, but can extend as mycelial networks into xylem (49) or living moss mats (39). Parts of the mycelium with phototroph cells may also differ based on position relative to growth tips, which are thought to include larger proportions of living cells (50), and entire strata can be phototroph-free.

Did stable phototroph association coincide with the loss of PCWDEs in LFSs? Looking only at the reductions in mean CAZyme gene number, the answer would appear to be positive, but the occurrence of LFSs with CAZyme arsenals as large or larger than those of many saprotrophs in the OG clade shows that LFSs do not necessarily lack CAZyme arsenals. Despite their minority representation in our data set, CAZyme-rich LFSs may in fact be numerous: the *Ostropales* and *Gyalectales* which make up the OG clade include over 3200 named species, almost 17% of named LFSs (51). For phototroph acquisition to have no evolutionary consequences for the CAZyme arsenal, symbiont-derived photosynthates would likely have to provide a function other than as a substrate for growth and respiration. In fact this has already long been postulated, in the form of an osmoregulatory role for polyols in managing cell desiccation (52). Some have even suggested this may be the main role for transferred photosynthates (53, 54). A strong inference about whether LFS CAZyme gene reduction coincided with the onset of stable phototroph association requires greater certainty about the ancestral states along the lecanoromycete phylogenetic backbone. Our ancestral state reconstruction suggests a gradual loss of GHs and PLs since the last common ancestor of Lecanoromycetes and Eurotiomycetes, and only pectin degradation genes are inferred to have been abruptly reduced. This reconstruction is sensitive to taxon undersampling and may be a conservative estimate. The large CAZyme arsenal of the OG clade, with dozens of unlinked genes and numerous PLs, is unlikely to have been acquired by lateral gene transfer. If the PL-rich OG clade CAZyme arsenal is in fact ancestral, this would imply that CAZyme loss is not an automatic consequence of stable association with phototroph symbionts, but rather of subsequent events or adaptations. If so, this implies that the OG clade CAZyme arsenal was lost no fewer than seven times in our tree. Support for a scenario of frequent mass gene reduction comes from the fact that one such loss appears to have happened within the OG clade itself: *Gomphillus* exhibits marked gene number reductions compared to its closest sampled relatives. Our data do not currently allow the hypothesis that the OG clade CAZyme arsenal is ancestral to be rejected, and if it is not, it becomes more likely that CAZyme loss in LFSs is driven by additional processes.

What additional adaptational processes could drive CAZyme loss? The most striking apparent functional losses, both in terms of CAZyme genes as well as in cellodextrin and cellobiose transporters, occurred in the *Lecanoromycetes* outside of the *Ostropomycetidae*, especially in the subclass *Lecanoromycetidae*, which largely lacks these two types of transporters. LFSs in *Lecanoromycetidae*, like those in *Ostropomycetidae*, are considered obligate symbionts of algae or cyanobacteria. They share many of the same algal symbionts, especially *Trebouxia*, and have no known lifecycle differences. However, they differ in general thallus architecture. *Ostropomycetidae* almost exclusively form crustose thalli in which thallus-to-substrate surface area contact is maximized. *Lecanoromycetidae*, and to some extent *Acarosporomycetidae* and *Umbilicariomycetidae*, include LFSs involved both in crustose thalli as well as so-called macrolichens, in which the thallus often becomes greatly expanded into leaf-, hair- or shrub-like thalli, in which surface contact is minimized. *Lecanoromycetidae* macrolichens include some of the largest lichens by biomass, and in theory these would require more carbon, not less. Lichens involving *Lecanoromycetidae* differ from those involving *Ostropomycetidae* in at least two traits that could be implicated in sugar uptake. In the first, the type of fungal-algal contact differs, from intracellular haustoria in *Ostropomycetidae* to so-called intraparietal haustoria, which do not breach the cell wall, in *Lecanoromycetidae* (55). The exact consequence of this is unknown, but intracellular haustoria are characteristic of pathogenic fungi and may require a greater variety of CAZymes to penetrate the algal cell wall. In the second, macrolichens involving *Lecanoromycetidae* owe their architecture to a well-developed exopolysaccharide gel, termed cortex, which forms a rigid structural exoskeleton considered a prerequisite to macrolichen formation (56). The polysaccharide composition of this layer varies widely across lichen symbioses (57) and is poorly characterized and mapped across the phylogenetic tree. Additional evolution in cortex layering happened in many lichens involving *Lecanoromycetidae* (58), though in how many remains unclear. The cortex, which is hydrophilic, also mediates water retention (57) and has been shown to operate like a sponge for passive uptake of dissolved nutrients (59) and glucose (60). One of the largest epiphytic macrolichens has been experimentally demonstrated to take up tree-derived glucose (61). Whether this is specifically facilitated by the cortex remains to be tested, but as all environmental molecules enter macrolichens through the cortex, it seems likely. If capture of simple sugars is found to be a general function of the cortex in different environments, it could be expected to have significant evolutionary consequences for maintenance of CAZyme genes used for more costly carbohydrates.

The existence of a robust carbohydrate breakdown machinery across a large swathe of LFS evolution calls into question the assumption, underlying decades of ecophysiological work on lichens, that the lichen carbon economy is exclusively the sum of algal CO_2_ fixation. The finding that some LFSs have CAZyme arsenals on par with saprotrophs lends support to Schwendener’s hypothesis that different types of lichens exist: those that depend mostly on their phototroph, and those that augment their carbon assimilation from multiple sources (1). The longstanding assumption that LFSs solely utilize fixed carbon must now be weighed against competing hypotheses, including A) that some or many LFSs build their mycelia from non-algal carbon sources, including absorbed monosaccharides and complex polysaccharides, before or during symbiosis; and B) that many different models of carbon acquisition — fixed, seasonal, facultative, scavenged and/or absorbed — may exist under the umbrella of what we currently call lichen symbiosis.

## Materials and Methods

Comprehensive Materials and Methods as well as supplementary results are available as supplementary material.

### Used genomes

We built a data set consisting of 83 fungal genomes from the phylum Ascomycota, including 46 genomes from the Lecanoromycetes, twenty-nine of which were newly generated for this study, and 37 genomes from the related classes *Eurotiomycetes*, *Dothideomycetes*, *Arthoniomycetes* and *Sordariomycetes*. Within *Lecanoromycetes*, the acquisition of new genomes was targeted to include a representation of lineages including 1) LFSs of macrolichens (*Cladonia*, *Evernia*, *Lobaria*, *Peltigera*, *Pseudevernia*, *Pseudocyphellaria*, *Ramalina*, *Sticta*, *Umbilicaria*); 2) LFS lineages involved in crust-forming lichens for which carbon acquisition from the substrate has been postulated, including wood specialists (*Lecidea scabridula*, *Lignoscripta*, *Ptychographa*, *Puttea*, *Xylographa* (18)), bark specialists (*Graphis*, *Loxospora*, *Schaereria*, *Stictis*, *Thelotrema*, *Varicellaria*), specialists of decaying plant matter (*Gomphillus*, *Icmadophila*), mineral soil (*Dibaeis*), rock (*Acarospora*, *Trapelia*) and a lichen-on-lichen “parasite” (*Lambiella*); 3) LFS lineages from crust-forming lichens that behave as ecological generalists (*Gyalolechia*, *Mycoblastus*, *Toensbergia*); and 4) a lecanoromycete saprotroph (*Agyrium*) and endophyte (*Cyanodermella* (62)). Our sampling simultaneously represents a cross-section of lecanoromycete evolution, with 24 genomes from the subclass *Ostropomycetidae*, 17 from the subclass *Lecanoromycetidae*, and four and one each from the species-poor subclasses *Acarosporomycetidae* and *Umbilicariomycetidae*, respectively. Ten genomes were derived from cultured samples and 18 are metagenome-assembled genomes (MAGs) assembled and binned according to (63). Culture-derived genomes differed little from MAGs in estimated completeness and gene numbers (Supplementary Figures 1 and 2).

### Genome assembly and filtering

Raw sequence data was inspected with FastQC 0.11.7 and trimmed with trimmomatic 0.29 to remove adapter remnants and low-quality reads. Genomes were assembled using SPAdes 3.12.0 or Abyss 2.0.2. To extract LFS contigs, draft assemblies were filtered using blobtools 1.1.1 or CONCOCT 1.2 and assembly completeness assessed with Quast 4.6.3 and BUSCO 3.0.2 We additionally extracted mitochondrial contigs by blasting mitochondrial genes downloaded from NCBI against each *de novo* assembly. We then discarded contigs with blast hits of e < 1e-03 and alignment length > 500 bp.

### Gene calling and functional annotation

We used funannotate 1.8.7 to perform gene-calling and functional annotation for all used genomes in the same way. This reduces potential biases introduced by different gene-calling and annotation methods. *De novo* sequenced genomes were repeat-masked using RepeatModeller and RepeatMasker. We then used Augustus 3.3, snap 2013_11_29, GeneMark-ES 4.62 and GlimmerHMM 3.0.4 and tRNA-Scan 2.0.5. Functional annotations for all predicted protein sequences were inferred using Interproscan-5.48-83.0, HMMer3 searches against dbCAN (v9; CAZymes) and Pfam (33.1) as well as eggnog-scanner searches against EggNOG (4.5.1; various annotations) databases.

### Phylogenomics

Phylogenomic trees were calculated using the phylociraptor pipeline (https://github.com/reslp/phylociraptor). We ran BUSCO 3.0.2 for each genome to identify single copy-orthologous, combined amino-acid sequences of each BUSCO gene from all genomes. Only genes which were present in >50% of genomes were aligned using mafft 7.464 and trimmed using trimal 1.4.1. We calculated single-gene trees using iqtree 2.0.7 and a species tree using ASTRAL 5.7.1. We created a concatenated alignment from all alignments, estimated the best substitution model for each and calculated a tree based on a partitioned analysis of the concatenated alignment using iqtree 2.0.7. We used the concatenated phylogeny to generate an ultrametric tree using r8s 1.81. This tree was used for subsequent analyses. We used custom python and R scripts to plot phylogenomic trees.

### Selection of CAZyme groups

We selected sets of CAZyme families involved in (hemi-)cellulose, pectin and lignin degradation based on previous studies (9, 24). For the lignin set we additionally identified class II peroxidases based on similarity of all Ascomycota class II peroxidases downloaded from RedOxiBase (accessed Jul. 14, 2021; http://peroxibase.toulouse.inra.fr/) using Orthofinder 2.5.2.

### Ancestral state reconstruction of CAZyme families and similarity of CAZyme sets

We reconstructed the ancestral size of each CAZyme family using CAZyme counts from our genome annotations and our ultrametric phylogenomic tree in R. We used anc.ML from phytools under an Ornstein-Uhlenbeck model of trait evolution. We then used custom R scripts to visualize ancestral states.

We calculated phylogenetically corrected PCAs in R as implemented in the phyltools function *phyl.pca* using maximum-likelihood optimization. We used log-transformed gene-counts of CAZyme families and our ultrametric phylogeny as input for phyl.pca. The PCAs were visualized using custom R scripts.

### Gene family expansion analysis

We analyzed gene family expansions of CAZyme families, based on CAZyme family count data from funannotate and our ultrametric phylogenomic tree using CAFE 5 (git commit 08d27a1). We ran CAFE twenty times while accounting for different gene-birth rates and error-models and summarized the results using custom R and python scripts.

### Additional characterization of CAZymes

CAZyme families known to contain important degradative enzymes (e.g., cellulases, hemicellulases) were further characterized using Saccharis v1 (git commit 9a748be). For each CAZyme family, we downloaded all characterized genes from cazy.org including additional information such as taxonomy, accession numbers and CAZyme (sub-)family assignments. Using Saccharis, we created MUSCLE 3.8.31 alignments for each gene family of all (from genomes used in this study plus characterized genes from cazy.org) genes and created maximum-likelihood trees using Fasttree 2.1.10. For all genes included in our phylogenetic reconstructions, we also predicted subcellular locations using DeepLoc 1.0. We only considered predictions for subcellular locations if the predicted probability was >70%. We used custom R and python scripts to visualize the trees.

### Heterologous expression of putative cellulases

Sequences assigned to GH5 subfamily 5, with confirmed cellulolytic activity, from obligately wood inhabiting LFS *Xylographa* species were aligned to the characterized and crystalized cellulase domain of *Trichoderma reesei* (PDB: 3QR3; https://www.rcsb.org/structure/3QR3). Based on sequence similarity, we selected two candidate cellulase sequences from *Xylographa bjoerkii* for testing cellulolytic activity.

Enzyme activity of the two candidate cellulases A and B was tested by combining in a 1.5 mL microcentrifuge tube 50 μL enzyme; 100 μL buffer A at either pH 4, 5, or 6; and 50 μL either AZCL-HE-cellulose, AZCL-β-glucan, or AZCL-xylan. Tubes were incubated at 4, 20, 37, 50, or 60°C for 48 h. To measure activity, samples were centrifuged at 13,000 rpm for 5 min to settle any debris, then 100 μL supernatant was removed to a 96-well flat-bottom plate. Absorbance at 595 nm was measured by plate reader and blanked with a sample containing water instead of enzyme.

### Identification of sugar- and sugar-alcohol transporters

To identify putative sugar- and sugar-alcohol transporters we used Orthofinder 2.5.2 on all sequences from all genomes with Pfam annotation PF00083 (Sugar_tr; http://pfam.xfam.org/family/sugar_tr) combined with characterized sugar transporter sequences from the PF00083 seed set. Additionally we added characterized cellodextrin (MH648002.1 (NCBI; from *Aspergillus niger*), S8AIR7 (UniProtKB; from *Penicillium oxalicum*)) and sugar alcohol transporters (AAX98668.1; from *Ambrosiozyma monospora*, CAR65543.1, CAG86001.1; from *Debaryomyces hansenii*, NP_010036.1; from *Saccharomyces cerevisiae*). We used the presence of characterized sequences in the inferred orthogroups to identify orthologs in each genome included in this study. The number of orthologs of different transporters per genome were visualized using custom R scripts.

### Identification of class II peroxidases

First we used diamond 0.9.22 to search all Ascomycota class II peroxidases downloaded from RedOxiBase (accessed Jul. 14, 2021) against the predicted proteins in all 83 genomes studied here. We then extracted all genomes which had a diamond hit to any of the downloaded sequences. Similar to the identification of sugar transporters we now used Orthofinder 2.5.2 to classify putative class II peroxidases persent in our genomes based on the presence of downloaded genes in individual orthogroups. The number of orthologs per genome for different peroxidases were visualized using custom R scripts.

## Data Availability

The complete analysis workflow used to acquire the results in this paper, including all custom python and R scripts, are available on Github (https://github.com/reslp/LFS-cazy-comparative). The workflow used to calculate phylogenomic trees is available on Github (https://github.com/reslp/phylociraptor). Genome assemblies and corresponding functional annotations will be deposited at the European Nucleotide Archive (ENA). Accession numbers are provided in supplementary material.

## Acknowledgments

The early phases of this project were funded by the Austrian Science Fund (FWF grant P25237, “Evolution of Substrate Specificity in Lichens”) and carried out at the Institute of Plant Sciences (Uni Graz). PR would like to acknowledge the Theodor Körner Funds (Vienna, Austria) and Network of Biological Systematics Austria (NOBIS; Vienna, Austria) for funding that enabled genome sequencing of several species. SW received funding from the Icelandic Research Fund IRF (174307-051), from Deutsche Forschungsgemeinschaft DFG (WE 6443/1-1) and from LMU Munich (startup funds). MW received funding from the Swedish Research Council, grant VR 2016-03589. MW and MK would like to acknowledge support from the NRM Deptartment of Bioinformatics and Genetics, the National Genomics Infrastructure in Stockholm, the SNIC/Uppsala Multidisciplinary Center for Advanced Computational Science and the UPPMAX computational infrastructure, and Linda Phillips for assistance with material. TS would like to acknowledge an NSERC Discovery Grant, a Canada Research Chair in Symbiosis, and the generosity of Susan Dalby and Eskild Petersen, who provided a place to work on this manuscript. Special thanks go to Sophie Dang at the Molecular Biology Service Unit, University of Alberta Department of Biological Sciences, for help in data acquisition, and members of the U of A Lichen Evolution Lab for reading and commenting on the manuscript. We are also grateful to Tanja Ernst and Andrea Brandl for laboratory assistance.

## Notes

### Competing Interest Statement

The authors have declared no competing interest.

