## Supplementary Material for "Large differences in carbohydrate degradation and transport potential in the genomes of lichen fungal symbionts"

**References** **111**

### Extended Materials and Methods

#### Overview of analysis workflow

The complete analysis workflow used to acquire the results in this paper, including all custom python and R scripts is available on Github (https://github.com/reslp/LFS-cazy-comparative). The workflow used to calculate phylogenomic trees is also available on Github (https://github.com/reslp/phylociraptor). An overview of the used software including version numbers is given in Table 3.

#### Dataset construction

This study is based on the analysis of 83 fungal genomes (Table 1). The selected genomes span six classes of Ascomycetes with focus on the largest radiation of lichen-fungal symbionts (LFS) Lecanoromycetes and their sister group Eurotiomycetes. We provide 29 *de-novo* sequenced draft genomes of LFSs and *Agyrium rufum*, which is a saprotrophic member of the Lecanoromycetes. Additional genomes include members of Dothideomycetes, Leotiomycetes, Sordariomycetes, Coniocybomycetes and Arthoniomycetes. Our study also includes published genomes of LFS genomes from the Lecanoromycetes, Arthoniomycetes and Eurotiomycetes (Table 1). For Eurotiomycetes, which are the sister-group to Lecanoromycetes, we included two members of each order (Table 1).

#### *De-novo* sequencing of LFS genomes

We report the genome sequences of 29 *de-novo* sequenced lichen fungal symbiont genomes. The genomes were sequenced between 2014 and 2019 with different sequencing technologies and from different starting material: Some are derived from axenic cultures of the fungal component of lichens while others have been extracted from sequenced whole-lichen metagenomes. Voucher information, and DNA extraction method is summarized in Table 2.

##### *Agyrium rufum*, *Lambiella insularis*, *Trapelia coarctata*, *Xylographa parallela*,*Xylographa pallens*

Axenic mycobiont cultures were obtained from hymenial fragments of freshly lichen material on Malt-Yeast extract agar following the protocol of (Molina and Crespo 2000) Tissue samples from axenic cultures of these four species were snap frozen at -80C and ground in a Retsch TissueLyser (Qiagen) immediately preceding nucleic acid extractions. We extracted DNA with the PowerBiofilm DNA Isolation Kit (MO BIO) and purified with Agencourt AMPure magnetic beads (Beckman Coulter) following manufacturers instructions. A minimum of 1$\mu$g of genomic DNA was used for library preparation. PCR-free libraries were prepared at the High Throughput Genomics Lab of the Huntsman Cancer Institute, Salt Lake City, UT with a mean insert size of 350 bp. These libraries were sequenced at the High Throughput Genomics Lab of the Huntsman Cancer Institute (Salt Lake City, UT, USA) on an Illumina HighSeq 2500 machine to 100 bp paired-end reads.

##### ***Lobaria immixta*, *Pseudocyphellaria aurata*, *Xylographa opegraphella*, *Xylographa soralifera*, *Xylographa bjoerkii* and *Xylographa trunciseda***

We extracted DNA from lichen fragments of *Lobaria immixta* and *Pseudocyphellaria aurata* and from axenic culture material of *Xylographa bjoerkii*, *X. opegraphella*, *X. pallens*, *X. soralifera* and *X. trunciseda* using a modified chloroform-phenol extraction method developed by the lab of Paul Dyer, University of Nottingham. We used about 2 $cm^2$ cleaned terminal pieces of the lichens or with pea-sized pieces of axenic culture as staring material, respectively.

1) 5PRIME Phase Lock Gel Light (hereafter: PLG; QuantaBio) tubes were centrifuged for 30s at 12,000 g immediately prior to use.
2) Then, 0.5 ml of DNA extraction buffer (room temperature; 250 mM Tris-HCl pH 8.5, 250 mM NaCl, 25 mM NaEDTA, 0.5% SDS) was added to the PLG tubes.
3) Freeze-dried material was ground thoroughly under liquid nitrogen in a mortar with a pistil and added to the DNA extraction buffer in the PLG tube..
4) One equal volume (0.5 ml) of ice-cold phenol:chloroform:isoamyl alcohol (25:24:1) was added to the ground mycelium in DNA extraction buffer, mixing thoroughly by inversion for 2 minutes, followed by centrifugation at 12,000 g at 4°C for 10 min.
5) The upper, aqueous phase was removed to a fresh tube, taking care not to disturb the interphase. 5 $\mu$L of RNAse (20 mg $ml^{-1}$) were added, incubating for 30 min at 37°C. Then, 1 $\mu$L of proteinase K was added to a final concentration of 10 $\mu$g/ml, incubating 40 min at 37°C. Finally, 1 volume (0.5 ml) of ice cold chloroform:isoamyl alcohol (24:1) was added, the solution was mixed thoroughly and centrifuged at 4°C and 12,000 g for 10 min. The upper (aqueous) phase was transferred to a new tube.
6) 0.54 volumes of isopropanol (-20°C ) were added and mixed by inversion. The solution was left for 20 minutes on ice to precipitate the DNA.
7) The solution was centrifuged for 20 min at 4°C and 12,000g to pellet the DNA and the supernatant was discarded after centrifugation.
8) 500 $\mu$L 70% ethanol (-20°C) were added to the pellet and the pellet was washed by gentle vortexing. Then, the solution was centrifuged at 4°C and 12,000 g for 5 minutes before removing the supernatant. Another wash step was performed.
9) The pellet was air dried at room temperature for 15-20 min, then the pellet was resuspended in 100 $\mu$L TE buffer (10 mM Tris-HCl, 1 mM EDTA, pH 8.0) overnight at 4°C. The extracted DNA was subsequently cleaned using Agencourt AMPure magnetic beads (Beckman Coulter) following manufacturer's instructions. Library preparation was performed by the Research Center for Molecular Medicine of the Austrian Academy of Sciences (CEMM) using TrueSeq Nano (Illumina) Library Preparation kits. The seven prepared libraries were sequenced on a single lane of an Illumina HighSeq 3000 machine to 150 bp paired-end reads.

##### ***Loxospora ochrophaea*, *Schaereria dolodes***

We extracted DNA using the DNeasy Plant Mini Kit (Qiagen, GmbH, Hilden, Germany) and prepared metagenomic libraries using TruSeq DNA PCR-Free Low Throughput Library Prep Kit (Illumina, San Diego, CA, USA). The libraries were sequenced by the Huntsman Cancer Center (Salt Lake City, UT, USA) on an Illumina HiSeq 2500 to 125 bp paired-end reads.

##### ***Ptychographa xylographoides*, *Xylographa vitiligo***

We extracted DNA from thallus parts indicated in Table 2 using the DNeasy Plant Mini Kit (Qiagen, GmbH, Hilden, Germany). Libraries were prepared using NEBNext Ultra II DNA Library Prep Kit (New England BioLabs, Ipswich, MA, USA) and sequenced by the Genome Sciences Centre (BC Cancer, Vancouver, BC, Canada) on an Illumina HiSeq X to 150 bp paired-end reads.

***Acarospora aff strigata*, *Bachmanniomyces* sp. TS44760, *Hypocenomyce scalaris*, *Lignoscripta atroalba*, *Mycoblastus sanguinarius*, *Puttea exsequens*, *Thelotrema lepadinum*, *Toensbergia leucococca*, *Varicellaria rhodocarpa*, *Xylographa carneopallida***

We extracted DNA from thallus parts indicated in Table 2 using the QIAamp Investigator kit (Qiagen, GmbH, Hilden, Germany) following the blood and tissue protocol. Libraries were prepared using NEBNext Ultra II DNA Library Prep Kit (New England BioLabs, Ipswich, MA, USA) and sequenced by the Genome Sciences Centre (BC Cancer, Vancouver, BC, Canada) on an Illumina HiSeq X to 150 bp paired-end reads.

***Stictis urceolatum***

We extracted DNA from apothecia collected from field-collected thalli of *S. urceolatum* using a CTAB method described in (Huang et al. 2018) with slight modifications by Mieko Kono. Library preparation and sequencing were done by SciLifeLab (Solna, Stockholm, Sweden) using SMARTer ThruPLEX DNA-seq kit (Takara Bio, Shiga, Japan) on an Illumina Miseq v3 to 300 bp paired-end reads.

***Peltigera leucophlebia* and *Sticta canariensis***

DNA was extracted from *Peltigera leucophlebia* apothecia cleaned of algae on the lower surface and for a chloromorph thallus of *Sticta canariensis* as follows: Samples were ground in a mortar with a pistil after immersion in liquid nitrogen for 2.5 minutes. The resulting fine powder was placed in a 2 ml tube and 450 $\mu$L lysis buffer (250 mM Tris-HCl pH 8.5, 250 mM NaCl, 25 mM NaEDTA, 0.5% SDS) and 4 $\mu$L RNAse A (20 mg/ml) were added. The sample was mixed by inversion and placed at -50°C for 20 min. Then, the tubes were spun at maximum speed (12000g) for 25 minutes at 20°C. The resulting supernatant (375 $\mu$L) was pipetted into 2 ml collection tubes. Next, 55 $\mu$L binding buffer (2M guanidine hydrochloride in 95% EtOH) were added and the sample was mixed well by pipetting and releasing the entire volume. The sample was added to a 3$\mu$m glass fiber plate (Chromafil Multi 96 GF 3 $\mu$m, Cat. Nr. 738658.M, Macherey-Nagel), spun for 5 min at 3800 g at 20°C, discarding the flow-through. Then, two washing steps followed where 500 $\mu$L of 70% EtOH were pipetted into the glass fiber plate wells and the plate was spun for 3 min at 3800 g at 20°C. After this, the plate was centrifuged for 15 min at 3800g and 20°C to remove residual ethanol. Now, the glass fiber plate was placed on a PCR plate and 55 $\mu$L TE elution buffer were added (5 mM Tris pH 8.0, 0.05mM EDTA) and the plate was spun for 1 min at 3800 g and at 20°C. The elution was repeated as above to obtain a total of 110 $\mu$L DNA. The sequencing libraries were prepared using the Nextera XT DNA Library Prep Kit according to manufacturers instructions and sequenced using the Illumina Miseq v3 platform generating 300 bp paired-end reads.

#### Raw sequencing data cleaning and filtering

Raw paired-end read data of the *de-novo* sequenced genomes were cleaned using trimmomatic after inspection with FastQC. Trimming parameters and the used trimmomatic versions are summarized in Table 4. Trimmed read files were inspected again with FastQC. Only trimmed reads were used for subsequent genome assembly.

#### *De-novo* assembly, quality assessment and filtering

We used different *de-novo* genome assemblers to assemble the trimmed reads of different species. For a small subset of species consisting of *Xylographa parallela*, *X. pallens*, *X. bjoerkii*, *X. soralifera*, *X. trunciseda* and *X. opegraphella* we created a number of test assemblies to investigate the impact of different assemblers on common genome and BUSCO completeness etc. For these species we assembled trimmed reads using SPAdes, Platanus, Minia, Abyss and Velvet. The quality of the generated test assemblies was assessed using QUAST and by comparing completeness of fungal (fungi_odb9) and ascomycete (ascomycota_odb9) BUSCOs using BUSCO 3. We used the best assemblies for the species included in our initial tests. We realized that SPAdes and Abyss produced consistently good assemblies in terms of BUSCO completeness and N50 while also being fast. Consequently we used either SPAdes or Abyss to assembly all *de-novo* sequenced genomes and the genome of *Graphis scripta*, for which only Illumina reads are available at the Sequence Read Archive (SRA) but no assembly. For each *de-novo* assembled genome the assembler and used parameters are summarized in Table 5.

*De-novo* assembled genomes were filtered using blobtools as implemented in the binner pipeline after assembly to filter out contaminant non-fungal contigs. After this first filtering step, genome completeness was assessed using QUAST and BUSCO 3. Blobtools filtering worked well for most species, however for *Bachmanniomyces sp. S44760*, *Puttea exsequens*, *Toensbergia leucococca* and *Mycoblastus sanguinarius* we had to use additional binning methods. *Mycoblastus sanguinarius* was binning using CONCOCT as implemented in the binner pipeline. The remaining three (*Bachmanniomyces sp. S44760*, *Puttea exsequens*, *Toensbergia leucococca*) species were binned using CONCOCT within the metaWRAP pipeline; the lecanoromycete genomes were detected and assessed by screening the bins with BUSCO 4 in the automated lineage selection mode.

Published assemblies downloaded from NCBI contain only the nuclear genome. Consequently we also limited our downstream analyses to the nuclear genome only. After the initial binning steps (see above) we filtered out candidate mitochondrial contigs based on BLAST similarity to published mitochondrial genes and complete mitochondrial genomes of *Graphis lineola* (Accession KY460674) and *Cladonia rangiferina* (Accession KY315996) (Pogoda et al. 2018). We selected these two species because they are members of the two large sub-groups of lichen-forming fungi Ostropomycetidae (*Graphis lineola*) and Lecanoromycetidae (*Cladonia rangiferina*). Using ncbi-blast+, we created BLAST databases for each binner filtered *de-novo* assembly and blasted all mitochondrial genes and genomes of *G. scripta* and *C. rangiferina* against the databases. Next we filtered the best BLAST hits for each query with an e-value cutoff of 1e-03 and a minimum alignment length of 500 base pairs to remove spurious hits. We extracted the target sequence names of these hits and filtered the input assemblies to remove all contigs containing hits from the query sequences. The so filtered genome assemblies were then used in all downstream analyses. An overview of common genome metrics is given in Figure 1.

#### Gene-calling and functional annotation

The quality and completeness of gene-models and functional annotations can vary greatly. Well-studied species often have reference genomes with high-quality, manually curated gene models and comprehensive functional annotations. For the majority of organisms however, high quality assemblies, gene-models and annotations are still lacking. In our study we combined heterogenic genomic data from a range of different sources, generated with different sequencing technologies at different times. This could introduce bias, potentially influencing the number of predicted genes, quality of functional annotation and thus affecting results of our study. To reduce potential bias of inconsistent gene calling and functional annotation to a minimum, we used the same gene calling and functional annotation regime for all included genomes, regardless of them being publicly available (and thus already having functional annotations) or *de-novo* sequenced. This ensures both comparability and reproducability of the results presented in this study. For all genomes we used the funannotate pipeline to predict genes and provide functional annotations. Funannotate provides wrapper scripts around several important steps of genome annotation and produces output which can easily be incorporated into downstream analyses. First we cleaned raw assemblies with *funannotate clean* to remove duplicated (percent identity > 95%) or too short (length < 500bp) contigs from subsequent annotation steps. Next we sorted the contigs in each assembly by length using *funannotate sort*. After this, we masked repetitive regions in the assemblies with *funannotate mask* using custom generated repeat libraries with RepeatModeler and RepeatMasker. Repeatmasked assembles were then subjected to gene-calling using the gene-callers Augustus, GeneMark-ES, snap and GlimmerHMM. tRNAs were predicted using tRNAScan. The so generated gene-models were subjected to searches against various databases to subsequently add functional annotations. Specifically we searched the predicted gene models against the InterPro databses using InterProScan and the PFAM database (version 33.1; Apr. 2020) using HMMer. Carbohydrate active enzymes (CAZymes) were predicted by HMMer against the dbCAN database version 9 (08 Apr. 2020; http://bcb.unl.edu/dbCAN2/) as part of the *funannotate annotate* step. Additional functional annotations were recovered from the EggNOG (version 4.5.1; 24 Mar. 2020; http://eggnog5.embl.de/) database. An overview of the number of called genes in each genome and annotated CAZymes is given in Figure 1.

#### Phylogenomic analyses and ultrametric tree reconstruction

We used phylociraptor for all steps of phylogenomic tree inference: For the initial phylogenomic estimation we used all single-copy BUSCO genes of the ascomycota_odb9 set found in at least 80% of genomes. An overview of BUSCO results are given in Figure 2. For each gene we created MAFFT alignments of the amino-acid sequence using the -auto flag. We trimmed alignments using trimAL with the -gappyout flag. This resulted in 1310 alignments (Table **??**), which were used for subsequent phylogenomic reconstruction. For each single-gene alignment we calculated the best substitution model using IQ-Tree's modeltest functionality and a maximum-likelihood tree. Using phylociraptor we created a concatenated alignment from all alignments and reconstructed a tree again with IQ-Tree. We performed a partitioned analysis using the best substitution model estimated for each gene.

As one measure of node support we calculated 1000 replicates of ultra fast bootstrap approximations (Hoang, biology, and 2017, n.d.). Since bootstrap has been identified to overestimate node support in phylogenomic datasets (Salichos and Rokas 2013) we also provide gene concordance and site concordance as two additional measures of node-support (Minh, Hahn, and Lanfear 2020). Gene concordance factors describe the proportion of genes supporting a node based on individual gene-trees, while site concordance factors describe the proportion of informative sites in the alignments supporting a node (Minh, Hahn, and Lanfear 2020). We calculated gene concordance and site concordance factors with IQ-Tree Additionally, we created a species tree using ASTRAL from the single-locus maximum-likelihood trees. We visualized all trees with custom R scripts (Figure 3 and 4).

We used r8s to transform the branch lengths of the concatenated tree topology to produce an ultrametric tree. We fixed the age of the tree root to an arbitrary value of 1000 (fixage taxon=root age=1000) and estimated the divergence times of descending nodes with a combination of penalized likelihood (Sanderson 2003) and truncated network method (divtime method=PL algorithm=TN). The smoothing parameter for the penalized likelihood approach was set to 500 (set smoothing=500). This ultrametric tree was used for downstream analyses.

#### Selection of plant cell wall degrading enzymes

The degradative potential of plant cell wall degrading enzymes (PCWDEs) of fungi is manifold. Previous genomic studies suggest that breakdown of common plant cell-wall components can involve many gene families acting in concertation (Miyauchi et al. 2020) (Floudas et al. 2012). We followed these studies on PCWD capabilities of fungi to select gene families potentially involved in degrading the three most common plant cell wall components (hemi)cellulose, pectin and lignin (Table 6).

#### Ancestral size and evolutionary dynamics of CAZyme families

To study the evolution of CAZyme families involved in PCWD we reconstructed ancestral family sizes along our phylogenomic tree. We used gene family count information of each CAZyme family size involved in (hemi-)cellulose, pectin and lignin degradation as inferred using funannotate for each species as input. We were specifically interested in nodes close to the evolution of major lichen-forming fungal groups. We thus reconstructed the ancestral size of CAZyme families for the root of the tree (R), the node of the Eurotiomycete and Lecanoromycete split (ELS), the ancestral node of Eurotiomycetes (AE), the node of the most recent common ancestor of Lecanoromycetes *sensu lato* (including *Acarospora* and *Umbilicaria*; ALSL), the node of the most recent common ancestor of Lecanoromycetes *sensu stricto* (ALSS), the node at the split between the two large subclasses Ostropomycetidae and Lecanoromycetidae (OLS), the MRCA nodes of Lecanoromycetidae (AL) and Ostropomycetidae (AO) as well as the MRCA of the genus *Xylographa* (AX). For ancestral state reconstruction we used the anc.ML method from phytools (Revell 2013) under an Ornstein-Uhlenbeck model of trait evolution (Butler and King 2004).

As a second approach and without restricting ourselves to *a-priori* defined sets of genes we analyzed all CAZyme families with CAFE. This allowed us to identify gene families with significantly elevated birth-rates, regardless of their affinity to different PCWDE groups. We used CAZyme family counts from funannotate and our ultrametric phylogenomic tree as input for CAFE. We imposed two models, one with a single gene-birth rate parameter for the whole tree and one with two independent birth rates; one for Lecanoromycetes and one the rest of the tree. The second model was introduced to test the assumption that some gene families, playing an important role in Lecanoromycetes, could also have significantly different birth-rates and thus a model assuming a single birth rate could be too simplistic. For each birth-rate scenario we also estimated a specific error model with CAFE. We ran the CAFE analysis five times for each birth-rate parameter model, with and without the specific error model. This lead us to 20 total CAFE runs (Table 8). We used custom R and python scripts to visualize expanded gene families and in how many CAFE runs the family was identified as significantly expanded ($p<0.05$) (Table 8, Figure 12).

#### Overall similarity of CAZyme sets

After investigating CAZyme families in an evolutionary context by reconstruction ancestral states, we were further interested in the overall similarity of extant CAZyme sets. Gene numbers for each species and each CAZyme family included in the pectin and (hemi-)cellulose sets of CAZymes as well as CAZymes of the lignin set combined with the estimated numbers of class II peroxidases (see below) were log transformed and subjected to phylogenetically informed PCA (Revell 2009), as implemented in the phytools function *phyl.pca*. We used a Maximum-Likelihood optimization method (opt="ML") and the mod="corr" parameter. To be able to compare overall similarity of extant species and the CAZyme compositions based on ancestral state reconstruction for the nine nodes along the tree backbone (see above) we additionally performed a classical PCA. Again we used log-transformed gene-count information for each CAZyme family of each species and reconstructed ancestral nodes and calculated a PCA using the R function *prcomp* with the parameter center=T. We visualized the first two principal components for each PCA in R using ggplot2 (Wickham 2016) (Figure 10). We also compared the mean numbers of CAZymes in the five CAZyme categories (AA, CBM, GH, GT and PL) between Lecanoromycetes groups and other fungi. First we calculated mean values of CAZyme counts for each CAZyme groups for all Lecanoromycetes, Lecanoromycetidae, the five members of the OG clade (see Fig.1 in the main text) and Ostropomycetidae (without OG clade members) and all other fungi. Next we calculated the difference of these values between LFS groups and all other fungi as follows:

$$\frac{|LFSgroup - OtherFungi|}{\frac{LFSgroup+OtherFungi}{2}} * 100 = \%Difference$$

Differences in % are summarized in (Table 9).

Next we compared the distributions of mean values for different LFS groups from above with a Wilcoxon rank-sum test in R to other fungi (Table 10).

Additionally we calculated the percent increase (positive numbers) or decrease (negative numbers) of CAZyme numbers between LFS groups and all other fungi (Table 11):

$$\frac{LFSgroup - OtherFungi}{OtherFungi} * 100 = \%Increase$$

#### Additional characterization of carbohydrate active enzymes

To characterize carbohydrate active enzymes beyond presence and absence in different genomes, we used a phylogenetic approach in combination with subcellular localization prediction and information of experimentally characterized enzymes listed in the cazy.org database. We started with downloaded information about experimentally characterized CAZymes by parsing the cazy.org website using a custom python script. This information contains broad taxonomic assignments (on the level of domains) as well as GenBank accession numbers, as well as descriptions of the activity of enzymes including Enzyme Code (EC) numbers. Cazy.org contains a large number of characterized CAZymes and CAZymes can act on many substrates apart from plant cell walls. This is especially important for some of the key families involved in degrading cellulose and hemicellulose, families GH5 and GH43. GH5 was originally called the "cellulase family" (Henrissat et al. 1989) however subsequent analyses revealed a large number of different enzymatic activities besides celluse activity (Aspeborg et al. 2012). Similarly GH43, a family involved in hemicellulose and pectin breakdown which is often expanded in PCWD organisms (Kohler et al. 2015) has been divided in many subfamilies to increase accuracy of predicted functions (Mewis et al. 2016). It thus can be expected that future studies will reveal a similar pattern in several other CAZyme familes.

To characterize functions of CAZymes in greater detail while taking into account potential multifunctionality within CAZyme families we blended information on characterized genes from each gene family listed in Table 6 with results from Saccharis. Saccharis is a phylogenetic pipeline specifically developed for CAZyme characterization. It takes CAZyme families as input and downloads all characterized sequences listed in cazy.org from GenBank. Then it searches the predicted protein sequences from our included genomes against HMM profiles (from dbcan) of the specified family to identify additional sequences with affinity to the CAZyme family under study. The so identified sequences are aligned together with the downloaded characterized sequences using MUSCLE and a phylogenetic tree is produced based on this alignment using FastTree.

Since plant-cell wall degrading enzymes have to act outside the fungal cell to have access to the long-chained carbohydrate substrates we predicted the subcellular location of CAZyme sequences using DeepLoc. Using protein sequences as input DeepLoc predicts the probability of an enzyme to be located in each of ten different subcellular locations. It is pre-trained on a large data set of UNIPROT sequences for which subcellular locations are known. We ran DeepLoc on all sequences assigned to a single CAZyme family by Saccharis including all sequences from characterized genes. We filtered DeepLoc results and kept location predictions only when probability for one location was above 70%. Finally we summarized information retrieved from cazy.org with the trees generated with Saccharis and subcellular location predictions as well as taxonomic assignments. We created a custom R script to visualize phylogenetic tree plots for each CAZyme family (Figure 3 in the main text; Figure 13.

#### Selection and heterologous expression of putative LFS cellulases

Cellulase candidate selection: We found that all characterized sequences in GH5 subfamily 5 have Cellulase activity (EC 3.2.1.4) and the vast majority come from eukaryotes (see Figure 3 in the main text). We aligned the cellulase domain of multiple GH5 subfamily 5 sequences from different *Xylographa* species to a previously epxressed and crystalized cellulase domain (PDB: 3QR3, https://www.rcsb.org/structure/3QR3) from *Trichoderma reesei*. We selected *Xylographa* for this experiment due to the obligate lichen-forming lifestyle and the obligate and close association with decaying wood of all members of this genus. Based on initial alignments using Clustal Omega we selected two candidate genes from *Xylographa bjoerkii* whith high similarity to 3QR3 for heterologous expression experiments. These genes are subsequently referred to cellulase A and B.

Cloning and expression: Synthetic genes of cellulase A and B were sub-cloned from pET28a into pMBPT via PCR amplification, SalI/NotI double-digestion, ligation, and transformation into *E. coli* BL21. Transformants were confirmed by sequencing. Cultures were routinely grown on LB agar or in LB broth (Fisher Scientific, Mississauga, ON) containing 100 $\mu$g/mL ampicillin. For expression, overnight cultures seeded from a single colony isolate were used to inoculate 600 mL LB broth with ampicillin, which was then grown at 37ºC with 180 rpm shaking until an OD600nm of ~0.4-0.6 was reached. Cultures were then induced with 0.5 mM IPTG and incubated for 24 h at 20ºC with 180 rpm shaking. Cells were harvested at 10,000 $\times$g for 15 min at 4ºC and stored at -80ºC until protein purification.

Cell lysis: Frozen cell pellets were mixed with an equal amount of diatomaceous earth, 1 uL Benzonase, and a protease inhibitor tablet (Roche), then ground manually by mortar and pestle for several minutes until homogeneous. Buffer A (100 mM Tris, 200 mM NaCl, pH 5.5) was added one mL at a time until a volume of 10 mL/g pellet was achieved. The mixture was transferred to a 50 mL conical tube and centrifuged at 10,000 $\times$g for 10 min at 4ºC to pellet debris. The supernatant was filtered through a 0.45 $\mu$m syringe-top filter to remove residual particulate matter prior to purification.

Protein purification: Filtered cell lysates were loaded onto a 5 mL MBPT column at approximately 1.5 mL/min by syringe, then the column was washed with 3 column volumes (CV) of buffer A. The protein was eluted from the column in 1.5 mL fractions with 1 CV of a 50% mixture of buffer A and buffer B (100 mM Tris, 200 mM NaCl, 10 mM maltose, pH 5.5), followed by 3 CV of 100% buffer B. The fraction containing the highest concentration of the target protein was determined visually by SDS-PAGE and confirmed by Western blot with anti-His-HRP.

Enzyme assay optimization: Enzyme activity was tested at pH 3, 4, 5, 6, or 7; and incubated at 4, 20, 37, 50, or 60ºC for 24, 48, or 72 h in a flat-bottom plate. Absorbance at 595 nm was measured by plate reader and blanked with a sample containing water instead of enzyme. Only optimal pH range and incubation time were presented.

#### Orthologue identification of cloned cellulases

To identify orthologues of the cloned cellulases we inferred orthogroups for the protein files produced with funannotate with Orthofinder. Orthofinder uses a hybrid approach based on sequence similarity estimated by a blast all-vs-all search with diamond and subsequent reconstruction of gene trees and a species tree to identify (single copy) orthologs and paralogs (Emms and Kelly 2019). The trees utilized by Orthofinder were reconstructed using Fasttree. The two cloned sequences were recovered in a single orthogroup. We used custom python and R scripts to count and visualize the number of orthologues in this orthogroup for each species.

#### Identification of sugar and sugar-alcohol transporters

Apart from the ability to break down PCW material, fungi also need the ability to transport breakdown products across the cell membrane to be able to metabolize them. To identify sugar transporter orthologues in LFSs we took all genes with PFAM annotations for sugar transporters (PF00083) and combined them with the seed set of characterized sugar transporters from the PFAM database. The PFAM seed set consists of 33 experimentally characterized sequences from a wide range of different organisms and covers transporters for different sugars. Additional information can be found here: http://pfam.xfam.org/family/sugar_tr.

To this initial set of sequences we added experimentally characterized fungal cellodextrin (*Aspergillus niger*: MH648002.1 (NCBI); *Penicillium oxalicum*: S8AIR7 (UniProtKB)) and sugar-alcohol (*Ambrosiozyma monospora*: AAX98668.1; Debaryomyces hansenii: CAR65543.1, CAG86001.1; *Saccharomyces cerevisiae*: NP_010036.1) transporter sequences. Again we used Orthofinder to assign all sugar transporters to orthogroups and used the presence of characterized sequences in each orthogroup to assign putative functions to sequences from the 83 genomes under study. We used custom python and R scripts to count and visualize the number of orthologues in each sugar-transporter orthogroup (Figure 5).

#### Identification of fungal peroxidases

To our knowledge class II peroxidases (PODs) have not been surveyed for LFSs so far despite their importance in lignin breakdown in fungi (Miyauchi et al. 2020; Kohler et al. 2015; Floudas et al. 2012) and their recently discovered unexpectedly high phylogenetic diversity (Mathé et al. 2019). To acquire a complete picture of the degradation potential of LFSs fungi we also identified a range of different peroxidases including class II peroxidases involved in lignin breakdown in the 83 genomes studied here. First we downloaded all 1290 Ascomycota peroxidase sequences from RedOxiBase (accessed Jul. 14 2021). Next, we used diamond 0.9.22 to search all characterized sequences against all predicted proteins from our genomes. We used the diamond results to subsample predicted proteins to only those sequences which had diamond hits to a characterized sequence. Next we used Orthofinder on the reduced protein sequence sets and all characterized peroxidase sequences to cluster them into orthogroups. We used the presence of characterized sequences in individual orthogroups to assigned names to orthogroups. Finally we used custom python and R scripts to count and visualize the number of orthologues in each peroxidase orthogroup (Figure 6).

### Supplementary Results

#### Table 1: Genomes included in this study

Table 1: List of fungal genomes analyzed in this study.

| isolate | de.novo.sequenced | origin | Genbank..SRA.accession | class.or.subclass |
|---|---|---|---|---|
| Acarospora aff strigata | yes | whole lichen | ?? | Lecanoromycetidae |
| Agyrium rufum | yes | axenic culture | ?? | Ostropomycetidae |
| Arthonia radiata | no | NCBI Genbank | GCA_002989075.1 | Arthoniomycetes |
| Aureobasidium pullulans | no | NCBI Genbank | GCA_003336255.1 | Dothideomycetes |
| Botrytis cinerea | no | NCBI Genbank | GCA_000143535.4 | Leotiomycetes |
| Capronia coronata | no | NCBI Genbank | GCA_000585585.1 | Eurotiomycetes |
| Ceratocystis platani | no | NCBI Genbank | GCA_000978885.1 | Sordariomycetes |
| Cetradonia linearis | no | NCBI Genbank | GCA_003521265.1 | Lecanoromycetidae |
| Cladonia grayi | no | NCBI Genbank | DOE JGI Portal - version 3 | Lecanoromycetidae |
| Cladonia macilenta | no | NCBI Genbank | GCA_000444155.1 | Lecanoromycetidae |
| Cladonia metacorallifera | no | NCBI Genbank | GCA_000482085.2 | Lecanoromycetidae |
| Cladonia uncialis | no | NCBI Genbank | GCA_002927785.1 | Lecanoromycetidae |
| Cladophialophora carrionii | no | NCBI Genbank | GCA_000365165.2 | Eurotiomycetes |
| Colletotrichum graminicola | no | NCBI Genbank | GCA_000149035.1 | Sordariomycetes |
| Coniella lustricola | no | NCBI Genbank | GCA_003019895.1 | Sordariomycetes |
| Coniochaeta ligniaria | no | NCBI Genbank | GCA_001879275.1 | Sordariomycetes |
| Cyanodermella asteris | no | NCBI Genbank | GCA_900618795.1 | Ostropomycetidae |
| Diaporthe longicolla | no | NCBI Genbank | GCA_000800745.1 | Sordariomycetes |
| Dibaeis baeomyces | no | Sequence Read Archive | SRX665192 | Ostropomycetidae |
| Elaphomyces granulatus | no | NCBI Genbank | GCA_002240705.1 | Eurotiomycetes |
| Endocarpon pusillum | no | NCBI Genbank | GCA_000464535.1 | Eurotiomycetes |
| Epichloe typhina | no | NCBI Genbank | GCA_000308955.1 | Sordariomycetes |
| Erysiphe necator | no | NCBI Genbank | GCA_000798715.1 | Leotiomycetes |
| Evernia prunastri | no | NCBI Genbank | GCA_003184365.1 | Lecanoromycetidae |
| Fonsecaea erecta | no | NCBI Genbank | GCA_001651985.1 | Eurotiomycetes |
| Fusarium oxysporum | no | NCBI Genbank | GCA_000149955.2 | Sordariomycetes |
| Graphis scripta | no | Sequence Read Archive | SRX665192 | Ostropomycetidae |
| Gomphillus americanus | no | NCBI Genbank | GCA_905337335.1 | Ostropomycetidae |
| Gyalolechia flavorubescens | no | NCBI Genbank | GCA_000442125.1 | Lecanoromycetidae |
| Helicocarpus griseus | no | NCBI Genbank | GCA_002573585.1 | Eurotiomycetes |
| Hypocenomyce scalaris | yes | whole lichen | ?? | Lecanoromycetidae |
| Hypoxylon pulicicidum | no | NCBI Genbank | GCA_002775035.1 | Sordariomycetes |
| Icmadophila ericetorum | yes | whole lichen | ?? | Ostropomycetidae |
| Knufia petricola | no | NCBI Genbank | GCA_002319055.1 | Eurotiomycetes |
| Lambiella insularis | yes | axenic culture | ?? | Ostropomycetidae |
| Lasallia hispanica | no | NCBI Genbank | GCA_003254425.1 | Lecanoromycetidae |
| Lasallia pustulata | no | NCBI Genbank | GCA_900169345.1 | Lecanoromycetidae |
| Bachmanniomyces sp. S44760 | yes | whole lichen | ?? | Ostropomycetidae |
| Lignoscripta atroalba | yes | whole lichen | ?? | Lecanoromycetidae |
| Lobaria immixta | yes | whole lichen | ?? | Ostropomycetidae |
| Loxospora ochrophaea | yes | axenic culture | ?? | Lecanoromycetidae |
| Magnaporthe grisea | no | NCBI Genbank | GCA_002871045.1 | Sordariomycetes |
| Malbranchea cinnamomea | no | NCBI Genbank | GCA_900128795.2 | Leotiomycetes |
| Monascus purpureus | no | NCBI Genbank | GCA_003184285.1 | Eurotiomycetes |
| Mycoblastus sanguinarius | yes | whole lichen | ?? | Ostropomycetidae |
| Onygena corvina | no | NCBI Genbank | GCA_000812245.1 | Eurotiomycetes |
| Peltigera leucophlebia | yes | whole lichen | ?? | Lecanoromycetidae |
| Penicillium chrysogenum | no | NCBI Genbank | GCA_000710275.1 | Eurotiomycetes |
| Phaeomoniella chlamydospora | no | NCBI Genbank | GCA_001006345.1 | Eurotiomycetes |
| Phialophora attae | no | NCBI Genbank | GCA_001299255.1 | Eurotiomycetes |
| Phyllosticta citricarpa | no | NCBI Genbank | GCA_001604955.1 | Dothideomycetes |
| Podospora anserina | no | NCBI Genbank | GCA_000226545.1 | Sordariomycetes |
| Pseudevernia furfuracea | no | NCBI Genbank | GCA_003184345.1 | Lecanoromycetidae |
| Pseudocyphellaria aurata | yes | whole lichen | ?? | Lecanoromycetidae |
| Pseudophaeomoniella oleicola | no | NCBI Genbank | GCA_003868215.1 | Eurotiomycetes |
| Ptychographa xylographoides | yes | whole lichen | ?? | Ostropomycetidae |
| Puttea exsequens | yes | whole lichen | ?? | Lecanoromycetidae |
| Ramalina intermedia | no | NCBI Genbank | GCA_003073195.1 | Lecanoromycetidae |
| Ramalina peruviana | no | NCBI Genbank | GCA_001956345.1 | Lecanoromycetidae |
| Schaereria dolodes | yes | whole lichen | ?? | Ostropomycetidae |
| Sordaria macrospora | no | NCBI Genbank | GCA_000182805.2 | Sordariomycetes |

Table 1: List of fungal genomes analyzed in this study. *(continued)*

| isolate | de.novo.sequenced | origin | Genbank..SRA.accession | class.or.subclass |
|---|---|---|---|---|
| Sticta canariensis | yes | whole lichen | ?? | Lecanoromycetidae |
| Stictis urceolatum | yes | whole lichen | ?? | Ostropomycetidae |
| Talaromyces cellulolyticus | no | NCBI Genbank | GCA_000829775.1 | Eurotiomycetes |
| Thelotrema lepadinum | yes | whole lichen | ?? | Ostropomycetidae |
| Thermoascus crustaceus | no | NCBI Genbank | GCA_001599835.1 | Eurotiomycetes |
| Thielaviopsis musarum | no | NCBI Genbank | GCA_001513885.1 | Sordariomycetes |
| Toensbergia leucococca | yes | whole lichen | ?? | Lecanoromycetidae |
| Trapelia coarctata | yes | axenic culture | ?? | Ostropomycetidae |
| Trichoderma reesei | no | NCBI Genbank | GCA_000167675.2 | Sordariomycetes |
| Umbilicaria muehlenbergii | no | NCBI Genbank | GCA_000611775.1 | Lecanoromycetidae |
| Uncinocarpus reesii | no | NCBI Genbank | GCA_000003515.2 | Eurotiomycetes |
| Varicellaria rhodocarpa | yes | whole lichen | ?? | Ostropomycetidae |
| Verticillium dahliae | no | NCBI Genbank | GCA_000150675.2 | Sordariomycetes |
| Xylaria grammica | no | NCBI Genbank | GCA_004353285.1 | Sordariomycetes |
| Xylographa bjoerkii | yes | axenic culture | ?? | Ostropomycetidae |
| Xylographa carneopallida | yes | whole lichen | ?? | Ostropomycetidae |
| Xylographa opegraphella | yes | axenic culture | ?? | Ostropomycetidae |
| Xylographa pallens | yes | axenic culture | ?? | Ostropomycetidae |
| Xylographa parallela | yes | axenic culture | ?? | Ostropomycetidae |
| Xylographa soralifera | yes | axenic culture | ?? | Ostropomycetidae |
| Xylographa trunciseda | yes | axenic culture | ?? | Ostropomycetidae |
| Xylographa vitiligo | yes | whole lichen | ?? | Ostropomycetidae |

#### Table 2: Sample information of *de-novo* sequenced genomes

Table 2: List of de-novo sequence

| isolate | extraction.method | class.or.subclass | voucher | material | date | comments |
|---|---|---|---|---|---|---|
| Acarospora aff strigata | QIAamp DNA Investigator Kit | Lecanoromycetidae | Spribille 44754, Isolate T1882, Canada:Alberta,Donalda | 10-15 thalli with apothecia | Mar. 31 2019 | Lichen mon |
| Agyrium rufum | PowerBiofilm DNA Isolation Kit | Ostropomycetidae | Spribille 39798, Canada: British Columbia, Gnat Pass | small pieces of mycelium | Aug. 13 2012 | |
| Hypocenomyce scalaris | QIAamp DNA Investigator Kit | Lecanoromycetidae | Spribille 44763, Isolate T1881, Canada:Alberta, E of Hinton along Hwy 16 | many individual squamules | Apr. 7 2019 | |
| Icmadophila ericetorum | QIAamp DNA Investigator Kit | Ostropomycetidae | Spribille 43583, isolate T1913, Canada: Manitoba,HIghway 6 N of Grand Rapids | sterile thallus only | Jul. 4 2018 | |
| Lambiella insularis | PowerBiofilm DNA Isolation Kit | Ostropomycetidae | Spribille 39820, USA: Montana near N end of Salmon Lake, Lincoln. Co. | cultured mycelium | Sept. 7 2012 | |
| Bachmanniomyces sp. S44760 | QIAamp DNA Investigator Kit | Ostropomycetidae | Spribille 44760, isolate T1894, Canada: Alberta, E of Hinton along Hwy 16 | seven or eight apothecia | Apr. 7 2019 | This is Lec |
| Lignoscripta atroalba | QIAamp DNA Investigator Kit | Lecanoromycetidae | Spribille 44772, isolate T1887, Canada: Alberta, Athabasca R near Brule | many apothecia | Apr. 7 2019 | |

Table 2: List of de-novo sequenced LFS

| isolate | extraction.method | class.or.subclass | voucher | material | date | comments |
|---|---|---|---|---|---|---|
| Lobaria immixta | Phenol-Chloroform extraction | Ostropomycetidae | - | - | - | |
| Loxospora ochrophaea | DNeasy Plant Mini | Lecanoromycetidae | Spribille 41460, isolates: T1860-1863 pooled, USA:New Hampshire, Kancamagus Highway | cultured mycelium | June 26, 2016 | extraction |
| Mycoblastus sanguinarius | QIAamp DNA Investigator Kit | Ostropomycetidae | Spribille 43910, isolate T1914, Canada: Saskatchewan, Wapawekka Hills | sterile thallus only | Jul. 07 2018 | |
| Peltigera leucophlebia | DNeasy Plant Mini?? | Lecanoromycetidae | isolate: Pleu1, Iceland: Heidmoerk (Lat: 64.06833844, Lng: -21.72876066) | mycelium from the backside of apothecia | - | |
| Pseudocyphellaria aurata | Phenol-Chloroform extraction | Lecanoromycetidae | - | - | - | |
| Ptychographa xylographoides | DNeasy Plant Mini | Ostropomycetidae | Spribille 42058, isolate T1868, Canada: Alberta, Rock Lake | | Sep. 25 2017 | |
| Puttea exsequens | QIAamp DNA Investigator Kit | Lecanoromycetidae | Spribille 42807, isolate T1888, Canada: Alberta, W of Clear Prairie | many apothecia | Sep 30 2017 | |
| Schaereria dolodes | DNeasy Plant Mini | Ostropomycetidae | Spribille 41586, Spribille 41654, isolates T1822 and T1849 pooled, USA: Lake Co, Tim Wheeler residence, Jocko River swimming hole | not recorded | Jan.-Mar. 2017 | |
| Sticta canariensis | Phenol-Chloroform extraction | Lecanoromycetidae | - | - | - | |
| Stictis urceolatum | CTAB Method | Ostropomycetidae | USA: Virginia, Smyth County, Grindstone Campground, on hardwood, leg. Linda Phillips (S) | apothecia | May 1 2017 | |
| Thelotrema lepadinum | QIAamp DNA Investigator Kit | Ostropomycetidae | Spribille 44606, isolate T1916 , Canada: British Columbia, Frisby Creek near Revelstoke | apotheica | Oct. 20 2018 | |
| Toensbergia leucococca | QIAamp DNA Investigator Kit | Lecanoromycetidae | Spribille 42847, isolate T1904, Canada: Saskatchewan, Wapawekka Hills | many squamules | Apr. 28 2018 | |

Table 2: List of de-novo sequenced LFS

| isolate | extraction.method | class.or.subclass | voucher | material | date | comments |
|---|---|---|---|---|---|---|
| Trapelia coarctata | PowerBiofilm DNA Isolation Kit | Ostropomycetidae | Resl 1158, isolate: P141, Austria: Styria, Schoeckl | small pieces of mycelium | Apr. 4 2012 | |
| Varicellaria rhodocarpa | QIAamp DNA Investigator Kit | Ostropomycetidae | Spribille 41938, isolate T1912, Canada: Northwest Territories, dolomite cliffs ca 8.5 km SW of Edzo | sterile thallus | Aug.17 | |
| Xylographa bjoerkii | Phenol-Chloroform extraction | Ostropomycetidae | Spribille 41600, USA: Oregon, beach across from Neah-Kah-Nie School | cultured mycelium | Jan. 29 2017 | |
| Xylographa carneopallida | QIAamp DNA Investigator Kit | Ostropomycetidae | Spribille 44761, isolate T1889, Canada: Alberta, E of Hinton along Hwy 16 | 10 - 15 apothecia and thallus | Apr. 07 2019 | |
| Xylographa opegraphella | Phenol-Chloroform extraction | Ostropomycetidae | Spribille 41601, USA: Oregon, beach across from Neah-Kah-Nie School | cultured mycelium | Jan 29, 2017 | |
| Xylographa pallens | Phenol-Chloroform extraction | Ostropomycetidae | Resl 1159 | small pieces of mycelium | | |
| Xylographa parallela | PowerBiofilm DNA Isolation Kit | Ostropomycetidae | Resl 1145, isolate T1151, Austria: Carinthia, Hochrindl | small pieces of mycelium | Jun. 5 2013 | |
| Xylographa soralifera | Phenol-Chloroform extraction | Ostropomycetidae | Spribille 41478, USA: Montana Silver Butte Pass | cultured mycelium | Aug. 15 2016 | |
| Xylographa trunciseda | Phenol-Chloroform extraction | Ostropomycetidae | Spribille 41477, USA: Montana Silver Butte Pass | cultured mycelium | Aug. 15 2016 | |
| Xylographa vitiligo | DNeasy Plant Mini | Ostropomycetidae | Spribille 42472, isolate T1866, Canada: Alberta, Swan Hills | soredia including wood grains | Oct. 28 2017 | |

#### Table 3: Used software

Table 3: Software used in this study.

| software | version | step | citation |
|---|---|---|---|
| trimmomatic | 0.38 | data cleaning | Bolger (2014) |
| FastQC | 0.11.7 | data cleaning | Andrews (2010) |
| binner | 0.1 | data cleaning | Resl (2020) |
| SPAdes | 3.12.0 | assembly | Bankevich et al. (2012) |
| platanus | 1.2.4 | assembly | Kajitani et al. (2014) |
| minia3 | git commit 1d5b8f4 | assembly | Chiki and Rizk (2012) |
| abyss | 2.0.1 | assembly | Jackman et al. (2017) |
| velvet | 1.2.0 | assembly | Zerbino and Birney (2008) |
| QUAST | 4.6.3 | assembly | Gurevich et al. (2013) |
| BUSCO | 3.0.2, 4.0.2 | assembly/phylogenomics | Waterhouse et al. (2017) |
| blobtools | 1.1.1 | assembly filtering | Laetsch and Blaxter (2017) |
| MetaWrap | ??? | assembly filtering | Uritskiy et al. (2018) |
| CONCOCT | 1.2 | assembly filtering (within MetaWrap) | Alneberg et al. (2014) |
| ncbi-blast+ | 2.9.0 | assembly filtering | Camacho et al. (2009) |
| funannotate | 1.8.7 | genome annotation | Palmer and Stajich (2021) |
| RepeatModeller | 1.0.11 | genome annotation | Smit el al. (2008) |
| RepeatMasker | 4.0.7 | genome annotation | Smit et al. (2013) |
| Augustus | 3.3.2 | genome annotation (within funannotate) | Stanke et al (2006) |
| GlimmerHMM | 3.0.4 | genome annotation (within funannotate) | Majoros et al. (2004) |
| snap | 2006-07-28 | genome annotation (within funannotate) | Korf (2004) |
| GeneMark-ES | 4.62 (Jan. 2020) | genome annotation (within funannotate) | Ter-Hovhannisyan et al. (2008) |
| tRNA-Scan | 2.0.5 | genome annotation (within funannotate) | Lowe and Eddy (1997) |
| InterproScan | 5.48-83.0 | genome annotation | Jones et al (2014) |
| eggnog-mapper | 1.0.3?? | genome annotation | Huerta-Cepas et al. (2017) |
| phylociraptor | git commit a93b4c8 | phylogenomics | Resl and Hahn (2021) |
| mafft | 7.464 | phylogenomics (within phylociraptor) | Katoh and Standley (2013) |
| trimAL | 1.4.1 | phylogenomics (within phylociraptor) | Capella-Gutierrez et al. (2009) |
| IQ-Tree | 2.0.7 | phylogenomics (within phylociraptor) | Minh et al. (2020) |
| ASTRAL | 5.7.1 | phylogenomics (within phylociraptor) | Mirarab et al. (2014) |
| r8s | 1.81 | phylogenomics | Sanderson (2003) |
| CAFE | 5.0.0b2 | gene family evolution analysis | Mendes et al. (2020) |
| Saccharis | git commit 9a748be | cazyme characterization | Jones et al. (2018) |
| MUSCLE | 3.8.31 | cazy characterization (within Saccharis) | Edgar (2004) |
| hmmer | 3.1b2 | cazy characterization (within Saccharis) | Mistry et al. (2013) |
| raxml | 8.2.12 | cazy characterization (within Saccharis) | Stamatakis (2014) |
| DeepLoc | 1.0 | cazyme characterization | Almagro Armenteros et al. (2017) |
| Clustal Omega | 1.2.4 | cazy characterization | Sievers et al. (2011) |
| Orthofinder | 2.5.2 | orthology detection | Emms and Kelly (2019) |
| diamond | 0.9.24, 0.9.22 | orthology detecion (within Orthofinder) | Buchfink et al. (2021) |
| Fasttree | 2.1.10 | orthology detection (within Orthofinder) | Price et al. (2010) |

#### Table 4: Read trimmers and parameters

Table 4: Software and parameters used in initial trimming of raw data of different species.

| | **trimmer** | **trimming__parameters** |
|---|---|---|
| Acarospora__aff__strigata | Trimmomatic 0.38 | ILLUMINACLIP:all__PE.fa:2:30:10 LEADING:30 TRAILING:30 SLIDINGWINDOW:4:15 MINLEN 80 |
| Agyrium__rufum | Trimmomatic 0.35 | LEADING:5 TRAILING:5 SLIDINGWINDOW:4:15 MINLEN:50 |
| Dibaeis__baeomyces | Trimmomatic 0.35 | ILLUMINACLIP:all__PE.fa:2:20:7:1:false LEADING:28 TRAILING:28 SLIDINGWINDOW:4:15 MINLEN:50 |
| Graphis__scripta | Trimmomatic 0.35 | ILLUMINACLIP:TruSeq2-PE.fa:2:30:10 LEADING:30 TRAILING:30 SLIDINGWINDOW:4:15 MINLEN:36 |
| Hypocenomyce__scalaris | Trimmomatic 0.38 | ILLUMINACLIP:all__PE.fa:2:30:10 LEADING:30 TRAILING:30 SLIDINGWINDOW:4:15 MINLEN 80 |
| Icmadophila__ericetorum | Trimmomatic 0.38 | ILLUMINACLIP:all__PE.fa:2:30:10 LEADING:30 TRAILING:30 SLIDINGWINDOW:4:15 MINLEN 80 |
| Lambiella__insularis | Trimmomatic 0.38 | ILLUMINACLIP:all__PE.fa:2:30:10 LEADING:30 TRAILING:30 SLIDINGWINDOW:4:15 MINLEN 80 |
| Bachmanniomyces__sp.__S44760 | Trimmomatic 0.38 | ILLUMINACLIP:all__PE.fa:2:30:10 LEADING:30 TRAILING:30 SLIDINGWINDOW:4:15 MINLEN 80 |
| Lignoscripta__atroalba | Trimmomatic 0.38 | ILLUMINACLIP:all__PE.fa:2:30:10 LEADING:30 TRAILING:30 SLIDINGWINDOW:4:15 MINLEN 80 |
| Lobaria__immixta | Trimmomatic 0.38 | ILLUMINACLIP:all__PE.fa:2:30:10 LEADING:33 TRAILING:33 SLIDINGWINDOW:15:20 MINLEN:36 |
| Loxospora__ochrophaea | Trimmomatic 0.38 | ILLUMINACLIP:all__PE.fa:2:30:10 LEADING:30 TRAILING:30 SLIDINGWINDOW:4:15 MINLEN 80 |
| Mycoblastus__sanguinarius | Trimmomatic 0.38 | ILLUMINACLIP:all__PE.fa:2:30:10 LEADING:30 TRAILING:30 SLIDINGWINDOW:4:15 MINLEN 80 |
| Peltigera__leucophlebia | Trimmomatic 0.36 | ILLUMINACLIP:NexteraPE-PE.fa:2:30:10 LEADING:30 TRAILING:30 SLIDINGWINDOW:4:15 MINLEN 50 |
| Pseudocyphellaria__aurata | Trimmomatic 0.38 | ILLUMINACLIP:all__PE.fa:2:30:10 LEADING:33 TRAILING:33 SLIDINGWINDOW:15:20 MINLEN:36 |
| Ptychographa__xylographoides | Trimmomatic 0.38 | ILLUMINACLIP:all__PE.fa:2:30:10 LEADING:30 TRAILING:30 SLIDINGWINDOW:4:15 MINLEN 80 |
| Puttea__exsequens | Trimmomatic 0.38 | ILLUMINACLIP:all__PE.fa:2:30:10 LEADING:30 TRAILING:30 SLIDINGWINDOW:4:15 MINLEN 80 |
| Schaereria__dolodes | Trimmomatic 0.38 | ILLUMINACLIP:all__PE.fa:2:30:10 LEADING:30 TRAILING:30 SLIDINGWINDOW:4:15 MINLEN 80 |
| Sticta__canariensis | Trimmomatic 0.38 | ILLUMINACLIP:all__PE.fa:2:30:10 LEADING:33 TRAILING:33 SLIDINGWINDOW:15:20 MINLEN:36 |
| Stictis__urceolatum | Trimmomatic 0.38 | ILLUMINACLIP:all__PE.fa:2:30:10 LEADING:30 TRAILING:30 SLIDINGWINDOW:4:15 MINLEN:100 |
| Thelotrema__lepadinum | | |
| Toensbergia__leucococca | Trimmomatic 0.38 | ILLUMINACLIP:all__PE.fa:2:30:10 LEADING:30 TRAILING:30 SLIDINGWINDOW:4:15 MINLEN 80 |
| Trapelia__coarctata | Trimmomatic 0.35 | LEADING:5 TRAILING:5 SLIDINGWINDOW:4:15 MINLEN:50 |
| Varicellaria__rhodocarpa | Trimmomatic 0.38 | ILLUMINACLIP:all__PE.fa:2:30:10 LEADING:30 TRAILING:30 SLIDINGWINDOW:4:15 MINLEN 80 |
| Xylographa__bjoerkii | Trimmomatic 0.38 | ILLUMINACLIP:all__PE.fa:2:30:10 LEADING:30 TRAILING:30 SLIDINGWINDOW:4:15 MINLEN 100 |
| Xylographa__carneopallida | Trimmomatic 0.38 | ILLUMINACLIP:all__PE.fa:2:30:10 LEADING:30 TRAILING:30 SLIDINGWINDOW:4:15 MINLEN 80 |
| Xylographa__opegraphella | Trimmomatic 0.38 | ILLUMINACLIP:all__PE.fa:2:30:10 LEADING:30 TRAILING:30 SLIDINGWINDOW:4:15 MINLEN 80 |
| Xylographa__pallens | Trimmomatic 0.38 | ILLUMINACLIP:all__PE.fa:2:30:10 LEADING:30 TRAILING:30 SLIDINGWINDOW:4:15 MINLEN 60 |
| Xylographa__parallela | Trimmomatic 0.38 | ILLUMINACLIP:all__PE.fa:2:30:10 LEADING:30 TRAILING:30 SLIDINGWINDOW:4:15 MINLEN 80 |
| Xylographa__soralifera | Trimmomatic 0.38 | ILLUMINACLIP:all__PE.fa:2:30:10 LEADING:30 TRAILING:30 SLIDINGWINDOW:4:15 MINLEN 100 |
| Xylographa__trunciseda | Trimmomatic 0.38 | ILLUMINACLIP:all__PE.fa:2:30:10 LEADING:30 TRAILING:30 SLIDINGWINDOW:4:15 MINLEN 60 |
| Xylographa__vitiligo | Trimmomatic 0.38 | ILLUMINACLIP:all__PE.fa:2:30:10 LEADING:30 TRAILING:30 SLIDINGWINDOW:4:15 MINLEN 80 |

#### Table 5: Genome assemblers and parameters

Table 5: Software and parameters used to assemble the genomes of different species.

| | assembler | parameters |
|---|---|---|
| Acarospora__aff__strigata | SPAdes v.3.12.0 | -k 21,31,51,71,81,101,127 –careful |
| Agyrium__rufum | SPAdes v.3.6.2 | -k 21,33,55,77,89 –careful |
| Dibaeis__baeomyces | SPAdes v.3.6.2 | -k 21,33,41,49 –careful |
| Graphis__scripta | SPAdes v.3.6.2 | -k 21,33,55,77,89 –careful |
| Hypocenomyce__scalaris | SPAdes v.3.12.0 | -k 21,31,51,71,81,101,127 –careful |
| Icmadophila__ericetorum | SPAdes v.3.12.0 | -k 21,31,51,71,81,101,127 –careful |
| Lambiella__insularis | SPAdes v.3.12.0 | -k 21,31,51,71,81,101,127 –careful |
| Bachmanniomyces__sp.__S44760 | SPAdes v.3.12.0 | -k 21,31,51,71,81,101,127 –careful |
| Lignoscripta__atroalba | SPAdes v.3.12.0 | -k 21,31,51,71,81,101,127 –careful |
| Lobaria__immixta | SPAdes v.3.12.0 | -k 31,43,55,67,79,91,103,115,127 –careful |
| Loxospora__ochrophaea | SPAdes v.3.12.0 | -k 21,31,51,71,81,101,127 –careful |
| Mycoblastus__sanguinarius | SPAdes v.3.12.0 | -k 21,31,51,71,81,101,127 –careful |
| Peltigera__leucophlebia | SPAdes v.3.12.0 | -k 21,31,51,71,89 –careful |
| Pseudocyphellaria__aurata | SPAdes v.3.12.0 | -k 31,43,55,67,79,91,103,115,127 –careful |
| Ptychographa__xylographoides | SPAdes v.3.12.0 | -k 21,31,51,71,81,101,127 –careful |
| Puttea__exsequens | SPAdes v.3.12.0 | -k 21,31,51,71,81,101,127 –careful |
| Schaereria__dolodes | SPAdes v.3.12.0 | -k 21,31,51,71,81,101,127 –careful |
| Sticta__canariensis | SPAdes v.3.12.0 | -k 31,43,55,67,79,91,103,115,127 –careful |
| Stictis__urceolatum | SPAdes v.3.12.0 | -k 21,31,51,71,81,101,127 –careful |
| Thelotrema__lepadinum | | |
| Toensbergia__leucococca | SPAdes v.3.12.0 | -k 21,31,51,71,81,101,127 –careful |
| Trapelia__coarctata | SPAdes v.3.6.2 | -k 21,33,55,77,89,101 –careful |
| Varicellaria__rhodocarpa | SPAdes v.3.12.0 | -k 21,31,51,71,81,101,127 –careful |
| Xylographa__bjoerkii | Abyss 2.0.1 | K=55 |
| Xylographa__carneopallida | SPAdes v.3.12.0 | -k 21,31,51,71,81,101,127 –careful |
| Xylographa__opegraphella | Abyss 2.0.1 | K=65 |
| Xylographa__pallens | Abyss 2.0.2 | K=90 |
| Xylographa__parallela | SPAdes v.3.12.0 | -k 61 –careful |
| Xylographa__soralifera | Abyss 2.0.1 | K=90 |
| Xylographa__trunciseda | SPAdes v.3.12.0 | -k 31 –careful |
| Xylographa__vitiligo | SPAdes v.3.12.0 | -k 21,31,51,71,81,101,127 –careful |

#### Table 6: Studied CAZyme families

Table 6: CAZyme families involved in degrading different PCW components studied here.

| | Set of gene families |
|---|---|
| (hemi)cellulose | AA3, AA9, CBM1, CBM13, CBM35, CBM6, CBM66, GH10, GH1, GH11, GH12, GH141, GH16, GH2, GH26, GH27, GH29, GH30, GH3, GH31, GH35, GH36, GH39, GH43, GH45, GH5, GH51, GH55, GH6, GH61, GH67, GH7, GH72, GH74, GH95 |
| pectin | CE8, GH105, GH28, GH49, GH53, GH79, GH88, PL1, PL3, PL4, PL9 |
| lignin | AA1, AA2, AA5 |

#### Table 7: Overview of single-copy genes and alignments used for phylogenomic reconstruction.

Table 7: Overview of alignments of genes used for phylogenomic reconstruction. Table generated with phylociraptor.

| gene | length | no. of sequences | no. of parsimony informative sites | no. of variable sites | no. of fixed sites | best model |
|---|---|---|---|---|---|---|
| EOG092D4D0M | 150 | 78 | 54 | 89 | 61 | DCMut+I+G4 |
| EOG092D0GSP | 963 | 81 | 442 | 889 | 74 | LG+F+I+G4 |
| EOG092D0VUY | 671 | 69 | 364 | 621 | 50 | LG+I+G4 |
| EOG092D1179 | 819 | 82 | 434 | 648 | 171 | JTT+F+I+G4 |
| EOG092D11OI | 1068 | 81 | 542 | 842 | 226 | LG+I+G4 |
| EOG092D1Q73 | 460 | 82 | 192 | 429 | 31 | LG+I+G4 |
| EOG092D09NC | 1338 | 82 | 435 | 1088 | 250 | JTT+F+I+G4 |
| EOG092D1P7M | 850 | 80 | 320 | 828 | 22 | JTT+I+G4 |
| EOG092D3CGZ | 248 | 74 | 102 | 237 | 11 | LG+F+I+G4 |
| EOG092D0S24 | 1206 | 78 | 309 | 1140 | 66 | JTT+F+I+G4 |
| EOG092D3M5I | 283 | 80 | 179 | 248 | 35 | LG+I+G4 |
| EOG092D2MTA | 1418 | 78 | 466 | 1398 | 20 | JTT+F+I+G4 |
| EOG092D2A39 | 972 | 81 | 342 | 922 | 50 | JTT+F+I+G4 |
| EOG092D1SW4 | 422 | 82 | 235 | 337 | 85 | LG+I+G4 |
| EOG092D4CWZ | 179 | 76 | 128 | 163 | 16 | LG+F+I+G4 |
| EOG092D01J4 | 2451 | 82 | 963 | 2413 | 38 | JTT+F+I+G4 |
| EOG092D3J3N | 265 | 79 | 165 | 229 | 36 | JTTDCMut+I+G4 |
| EOG092D1ODS | 436 | 83 | 253 | 404 | 32 | LG+F+I+G4 |
| EOG092D42U2 | 225 | 76 | 140 | 201 | 24 | LG+I+G4 |
| EOG092D2V2F | 317 | 82 | 32 | 262 | 55 | JTT+F+I+G4 |
| EOG092D4NC2 | 121 | 81 | 86 | 98 | 23 | LG+I+G4 |
| EOG092D3TIT | 627 | 80 | 163 | 585 | 42 | LG+F+I+G4 |
| EOG092D13U3 | 725 | 73 | 338 | 688 | 37 | JTT+I+G4 |
| EOG092D2GBR | 378 | 67 | 231 | 347 | 31 | LG+I+G4 |
| EOG092D26GB | 818 | 81 | 196 | 794 | 24 | VT+F+I+G4 |
| EOG092D4CNZ | 273 | 64 | 146 | 261 | 12 | JTT+F+I+G4 |
| EOG092D3SSN | 179 | 80 | 93 | 128 | 51 | LG+I+G4 |
| EOG092D1KAG | 747 | 80 | 485 | 697 | 50 | JTTDCMut+I+G4 |
| EOG092D0T1H | 820 | 76 | 506 | 720 | 100 | LG+I+G4 |
| EOG092D28JA | 383 | 77 | 240 | 322 | 61 | LG+F+I+G4 |
| EOG092D2ZQ2 | 433 | 69 | 195 | 365 | 68 | LG+F+I+G4 |
| EOG092D1FF7 | 548 | 83 | 206 | 265 | 283 | LG+I+G4 |
| EOG092D0VMZ | 703 | 83 | 394 | 660 | 43 | LG+F+I+G4 |
| EOG092D4HEW | 193 | 83 | 102 | 166 | 27 | LG+I+G4 |
| EOG092D3ZQC | 738 | 81 | 140 | 718 | 20 | JTT+F+I+G4 |
| EOG092D2MZB | 393 | 81 | 211 | 354 | 39 | LG+I+G4 |
| EOG092D1S42 | 413 | 80 | 181 | 375 | 38 | LG+I+G4 |
| EOG092D4KES | 139 | 70 | 21 | 64 | 75 | LG+G4 |
| EOG092D0GFU | 958 | 72 | 653 | 911 | 47 | LG+F+I+G4 |
| EOG092D1KPD | 549 | 81 | 246 | 341 | 208 | LG+I+G4 |
| EOG092D47B5 | 978 | 83 | 449 | 822 | 156 | JTT+I+G4 |
| EOG092D1SLW | 428 | 82 | 196 | 346 | 82 | LG+I+G4 |
| EOG092D4D2W | 178 | 77 | 126 | 171 | 7 | LG+I+G4 |
| EOG092D0DUF | 1011 | 79 | 561 | 800 | 211 | LG+I+G4 |
| EOG092D4O4M | 190 | 83 | 75 | 167 | 23 | JTT+I+G4 |
| EOG092D069S | 1210 | 75 | 499 | 874 | 336 | LG+F+I+G4 |
| EOG092D2RN7 | 383 | 80 | 215 | 333 | 50 | LG+I+G4 |
| EOG092D172M | 543 | 83 | 260 | 464 | 79 | LG+I+G4 |
| EOG092D24ND | 672 | 74 | 299 | 646 | 26 | WAG+G4 |
| EOG092D2DN8 | 379 | 81 | 126 | 224 | 155 | LG+I+G4 |
| EOG092D2GPJ | 470 | 76 | 210 | 412 | 58 | LG+I+G4 |
| EOG092D076U | 1032 | 79 | 568 | 754 | 278 | LG+I+G4 |
| EOG092D0ELD | 1809 | 81 | 416 | 1764 | 45 | JTT+F+I+G4 |
| EOG092D3B56 | 715 | 81 | 152 | 588 | 127 | JTT+F+I+G4 |

Table 7: Overview of alignments of genes used for phylogenomic reconstruction. Table generated with phylociraptor. *(continued)*

| gene | length | no. of sequences | no. of parsimony informative sites | no. of variable sites | no. of fixed sites | best model |
|---|---|---|---|---|---|---|
| EOG092D0M7O | 735 | 83 | 349 | 689 | 46 | LG+F+I+G4 |
| EOG092D3FK5 | 179 | 70 | 102 | 177 | 2 | LG+G4 |
| EOG092D0J6Y | 766 | 75 | 367 | 570 | 196 | JTT+I+G4 |
| EOG092D4A2D | 273 | 81 | 176 | 235 | 38 | LG+I+G4 |
| EOG092D0CVR | 947 | 71 | 557 | 926 | 21 | LG+F+I+G4 |
| EOG092D0Q9D | 640 | 81 | 306 | 432 | 208 | LG+I+G4 |
| EOG092D17F4 | 536 | 78 | 323 | 445 | 91 | JTTDCMut+I+G4 |
| EOG092D3PL7 | 400 | 83 | 193 | 334 | 66 | LG+F+I+G4 |
| EOG092D3D0W | 503 | 78 | 96 | 455 | 48 | JTT+F+I+G4 |
| EOG092D3BKV | 405 | 83 | 201 | 307 | 98 | JTT+F+I+G4 |
| EOG092D1GOQ | 426 | 77 | 205 | 323 | 103 | LG+I+G4 |
| EOG092D07WZ | 1021 | 77 | 749 | 921 | 100 | LG+F+I+G4 |
| EOG092D430X | 488 | 81 | 256 | 444 | 44 | JTT+F+I+G4 |
| EOG092D4CHO | 191 | 81 | 69 | 152 | 39 | LG+I+G4 |
| EOG092D4DWT | 170 | 80 | 101 | 131 | 39 | LG+G4 |
| EOG092D1W8I | 454 | 75 | 323 | 401 | 53 | LG+I+G4 |
| EOG092D3IB3 | 419 | 82 | 164 | 368 | 51 | JTT+I+G4 |
| EOG092D3ALG | 297 | 82 | 170 | 251 | 46 | LG+I+G4 |
| EOG092D3JAI | 492 | 82 | 162 | 470 | 22 | JTT+I+G4 |
| EOG092D23DO | 653 | 82 | 118 | 612 | 41 | JTT+F+I+G4 |
| EOG092D3B18 | 253 | 74 | 145 | 238 | 15 | LG+I+G4 |
| EOG092D1U19 | 623 | 83 | 297 | 536 | 87 | WAG+I+G4 |
| EOG092D0QVE | 906 | 82 | 341 | 714 | 192 | JTTDCMut+F+I+G4 |
| EOG092D4TKE | 170 | 67 | 70 | 160 | 10 | VT+I+G4 |
| EOG092D09F6 | 970 | 74 | 565 | 692 | 278 | LG+I+G4 |
| EOG092D1YB4 | 569 | 77 | 333 | 517 | 52 | JTT+F+I+G4 |
| EOG092D2SHG | 336 | 82 | 205 | 258 | 78 | LG+I+G4 |
| EOG092D2UBG | 501 | 72 | 312 | 464 | 37 | LG+F+I+G4 |
| EOG092D1UUH | 505 | 82 | 214 | 400 | 105 | LG+I+G4 |
| EOG092D2AVN | 286 | 76 | 215 | 274 | 12 | LG+I+G4 |
| EOG092D1BFC | 538 | 74 | 209 | 288 | 250 | LG+I+G4 |
| EOG092D0SFV | 663 | 82 | 516 | 634 | 29 | JTT+I+G4 |
| EOG092D0UBQ | 610 | 82 | 342 | 515 | 95 | JTTDCMut+I+G4 |
| EOG092D431N | 180 | 67 | 112 | 155 | 25 | VT+G4 |
| EOG092D4380 | 176 | 81 | 113 | 139 | 37 | LG+I+G4 |
| EOG092D2N4T | 500 | 82 | 326 | 443 | 57 | LG+I+G4 |
| EOG092D47GI | 183 | 73 | 99 | 169 | 14 | LG+G4 |
| EOG092D0L00 | 836 | 82 | 439 | 814 | 22 | JTT+F+I+G4 |
| EOG092D0P1F | 775 | 83 | 297 | 729 | 46 | JTT+I+G4 |
| EOG092D2DNW | 607 | 80 | 254 | 569 | 38 | LG+F+I+G4 |
| EOG092D3VG3 | 300 | 79 | 186 | 262 | 38 | LG+I+G4 |
| EOG092D0NEP | 969 | 72 | 392 | 948 | 21 | LG+F+I+G4 |
| EOG092D1LJK | 704 | 81 | 248 | 601 | 103 | JTT+F+I+G4 |
| EOG092D1RDH | 447 | 77 | 250 | 413 | 34 | LG+I+G4 |
| EOG092D2BJL | 151 | 73 | 64 | 126 | 25 | LG+I+G4 |
| EOG092D3CQ9 | 348 | 78 | 150 | 326 | 22 | JTTDCMut+I+G4 |
| EOG092D4H8N | 167 | 80 | 63 | 157 | 10 | WAG+F+I+G4 |
| EOG092D1KTN | 429 | 70 | 243 | 374 | 55 | LG+I+G4 |
| EOG092D2W7U | 332 | 82 | 89 | 269 | 63 | JTT+G4 |
| EOG092D1ULX | 404 | 78 | 283 | 346 | 58 | LG+I+G4 |
| EOG092D4DTK | 137 | 78 | 61 | 127 | 10 | VT+I+G4 |
| EOG092D3ECY | 343 | 79 | 214 | 316 | 27 | LG+F+I+G4 |
| EOG092D011S | 2334 | 82 | 644 | 2197 | 137 | JTT+F+I+G4 |
| EOG092D47YB | 234 | 81 | 148 | 201 | 33 | LG+I+G4 |
| EOG092D01WH | 1137 | 77 | 362 | 1128 | 9 | JTT+F+I+G4 |
| EOG092D357K | 408 | 82 | 245 | 353 | 55 | LG+I+G4 |
| EOG092D27ZM | 390 | 81 | 77 | 191 | 199 | LG+I+G4 |

Table 7: Overview of alignments of genes used for phylogenomic reconstruction. Table generated with phylociraptor. *(continued)*

| gene | length | no. of sequences | no. of parsimony informative sites | no. of variable sites | no. of fixed sites | best model |
|---|---|---|---|---|---|---|
| EOG092D4C1H | 145 | 54 | 62 | 129 | 16 | LG+G4 |
| EOG092D1NDG | 398 | 80 | 196 | 295 | 103 | LG+I+G4 |
| EOG092D3VIZ | 807 | 82 | 410 | 721 | 86 | JTT+F+I+G4 |
| EOG092D1T7G | 748 | 82 | 418 | 643 | 105 | JTT+I+G4 |
| EOG092D2ZWY | 322 | 74 | 177 | 275 | 47 | LG+I+G4 |
| EOG092D0JA2 | 750 | 77 | 522 | 651 | 99 | LG+F+I+G4 |
| EOG092D1L0N | 506 | 79 | 294 | 415 | 91 | LG+F+I+G4 |
| EOG092D2MAK | 693 | 79 | 447 | 675 | 18 | LG+I+G4 |
| EOG092D4H14 | 319 | 79 | 129 | 301 | 18 | LG+I+G4 |
| EOG092D3BBO | 653 | 77 | 301 | 595 | 58 | JTT+I+G4 |
| EOG092D3OFG | 401 | 79 | 162 | 340 | 61 | JTTDCMut+F+I+G4 |
| EOG092D2RP6 | 354 | 80 | 212 | 298 | 56 | LG+I+G4 |
| EOG092D04ZP | 1166 | 81 | 570 | 829 | 337 | LG+I+G4 |
| EOG092D3RZ8 | 274 | 76 | 169 | 243 | 31 | LG+F+I+G4 |
| EOG092D4G83 | 223 | 79 | 100 | 207 | 16 | JTT+I+G4 |
| EOG092D3LRW | 326 | 78 | 151 | 304 | 22 | JTTDCMut+F+I+G4 |
| EOG092D0NV0 | 663 | 76 | 361 | 631 | 32 | JTT+I+G4 |
| EOG092D13KG | 862 | 81 | 319 | 782 | 80 | LG+F+I+G4 |
| EOG092D1FGG | 511 | 82 | 180 | 259 | 252 | LG+I+G4 |
| EOG092D10DF | 848 | 82 | 497 | 779 | 69 | JTTDCMut+I+G4 |
| EOG092D4JU7 | 554 | 76 | 143 | 546 | 8 | VT+F+I+G4 |
| EOG092D3LA8 | 330 | 80 | 96 | 288 | 42 | LG+I+G4 |
| EOG092D444G | 150 | 73 | 93 | 125 | 25 | WAG+I+G4 |
| EOG092D3V1W | 168 | 71 | 84 | 158 | 10 | LG+I+G4 |
| EOG092D07BL | 1339 | 80 | 731 | 1136 | 203 | LG+I+G4 |
| EOG092D16TH | 899 | 82 | 324 | 746 | 153 | JTT+I+G4 |
| EOG092D4AY8 | 160 | 77 | 123 | 140 | 20 | LG+I+G4 |
| EOG092D3XVK | 216 | 78 | 67 | 128 | 88 | LG+I+G4 |
| EOG092D34ZI | 401 | 79 | 180 | 344 | 57 | JTT+F+I+G4 |
| EOG092D2DQ7 | 428 | 80 | 242 | 366 | 62 | LG+I+G4 |
| EOG092D4XRX | 903 | 83 | 411 | 897 | 6 | JTT+F+I+G4 |
| EOG092D3DNT | 291 | 76 | 217 | 281 | 10 | LG+G4 |
| EOG092D4KVP | 245 | 78 | 171 | 232 | 13 | LG+I+G4 |
| EOG092D070E | 1507 | 83 | 739 | 1280 | 227 | JTT+I+G4 |
| EOG092D248P | 555 | 83 | 291 | 463 | 92 | LG+F+I+G4 |
| EOG092D4RYQ | 118 | 82 | 28 | 48 | 70 | JTT+I+G4 |
| EOG092D44BU | 202 | 80 | 126 | 174 | 28 | VT+I+G4 |
| EOG092D4PUW | 184 | 77 | 87 | 156 | 28 | Dayhoff+F+I+G4 |
| EOG092D33KV | 437 | 76 | 203 | 378 | 59 | LG+I+G4 |
| EOG092D3IVF | 336 | 81 | 158 | 297 | 39 | JTT+F+I+G4 |
| EOG092D2LES | 430 | 82 | 285 | 387 | 43 | LG+G4 |
| EOG092D48ZU | 185 | 71 | 79 | 142 | 43 | VT+G4 |
| EOG092D2VOM | 299 | 82 | 124 | 204 | 95 | LG+F+I+G4 |
| EOG092D4H30 | 92 | 80 | 7 | 68 | 24 | VT+G4 |
| EOG092D0HE1 | 947 | 80 | 564 | 823 | 124 | LG+F+I+G4 |
| EOG092D2C76 | 799 | 78 | 392 | 756 | 43 | JTT+I+G4 |
| EOG092D434J | 245 | 80 | 160 | 225 | 20 | LG+I+G4 |
| EOG092D4D4F | 134 | 74 | 110 | 129 | 5 | LG+I+G4 |
| EOG092D1Q04 | 579 | 75 | 287 | 567 | 12 | JTT+I+G4 |
| EOG092D1V8V | 432 | 83 | 114 | 383 | 49 | JTTDCMut+I+G4 |
| EOG092D0ITM | 825 | 81 | 562 | 742 | 83 | JTT+F+I+G4 |
| EOG092D4MNU | 177 | 76 | 74 | 134 | 43 | LG+I+G4 |
| EOG092D1DIJ | 502 | 83 | 317 | 418 | 84 | JTT+I+G4 |
| EOG092D3PNL | 402 | 72 | 237 | 381 | 21 | JTT+I+G4 |
| EOG092D23WM | 556 | 83 | 132 | 520 | 36 | JTT+I+G4 |
| EOG092D4N86 | 198 | 81 | 63 | 162 | 36 | LG+I+G4 |
| EOG092D12CU | 672 | 81 | 218 | 339 | 333 | LG+I+G4 |

Table 7: Overview of alignments of genes used for phylogenomic reconstruction. Table generated with phylociraptor. *(continued)*

| gene | length | no. of sequences | no. of parsimony informative sites | no. of variable sites | no. of fixed sites | best model |
|---|---|---|---|---|---|---|
| EOG092D35K7 | 284 | 82 | 99 | 278 | 6 | JTT+I+G4 |
| EOG092D3UVA | 468 | 83 | 339 | 458 | 10 | LG+I+G4 |
| EOG092D2DE1 | 350 | 78 | 213 | 274 | 76 | LG+F+I+G4 |
| EOG092D4KSS | 113 | 52 | 61 | 85 | 28 | LG+G4 |
| EOG092D38OJ | 268 | 79 | 182 | 256 | 12 | JTTDCMut+I+G4 |
| EOG092D2NK1 | 471 | 82 | 276 | 419 | 52 | LG+F+I+G4 |
| EOG092D1718 | 1493 | 81 | 264 | 1470 | 23 | JTT+F+I+G4 |
| EOG092D2A7Q | 409 | 79 | 274 | 370 | 39 | LG+I+G4 |
| EOG092D2XJY | 313 | 61 | 192 | 305 | 8 | LG+I+G4 |
| EOG092D4C59 | 157 | 81 | 100 | 122 | 35 | LG+I+G4 |
| EOG092D3KF0 | 428 | 82 | 196 | 389 | 39 | LG+F+I+G4 |
| EOG092D0590 | 1318 | 82 | 697 | 1201 | 117 | JTT+F+I+G4 |
| EOG092D0YSG | 924 | 80 | 349 | 825 | 99 | JTT+I+G4 |
| EOG092D27VI | 415 | 81 | 193 | 387 | 28 | JTT+I+G4 |
| EOG092D2U5N | 545 | 80 | 142 | 533 | 12 | JTT+I+G4 |
| EOG092D4VBM | 139 | 12 | 57 | 111 | 28 | LG+F+G4 |
| EOG092D2PEN | 416 | 78 | 239 | 331 | 85 | LG+I+G4 |
| EOG092D2VEC | 395 | 78 | 205 | 287 | 108 | LG+I+G4 |
| EOG092D39EO | 298 | 76 | 204 | 283 | 15 | LG+I+G4 |
| EOG092D28EY | 1131 | 82 | 450 | 1070 | 61 | JTT+F+I+G4 |
| EOG092D3FDW | 298 | 79 | 205 | 284 | 14 | LG+I+G4 |
| EOG092D3GMS | 111 | 76 | 48 | 67 | 44 | WAG+G4 |
| EOG092D1LTM | 530 | 81 | 385 | 483 | 47 | LG+F+I+G4 |
| EOG092D0YW3 | 665 | 79 | 231 | 560 | 105 | JTT+I+G4 |
| EOG092D43MS | 186 | 79 | 75 | 139 | 47 | LG+I+G4 |
| EOG092D35VJ | 480 | 82 | 207 | 371 | 109 | JTTDCMut+F+I+G4 |
| EOG092D2ICY | 354 | 78 | 125 | 220 | 134 | WAG+I+G4 |
| EOG092D1ZZP | 504 | 77 | 345 | 435 | 69 | LG+I+G4 |
| EOG092D2UWC | 308 | 79 | 172 | 250 | 58 | LG+I+G4 |
| EOG092D4OXW | 113 | 82 | 48 | 83 | 30 | LG+I+G4 |
| EOG092D2XOE | 611 | 81 | 184 | 569 | 42 | LG+I+G4 |
| EOG092D3J4Y | 316 | 82 | 153 | 294 | 22 | WAG+I+G4 |
| EOG092D28BN | 605 | 69 | 136 | 570 | 35 | LG+I+G4 |
| EOG092D2W27 | 301 | 77 | 203 | 263 | 38 | JTT+F+I+G4 |
| EOG092D0KUS | 778 | 78 | 483 | 673 | 105 | JTT+I+G4 |
| EOG092D4FS0 | 281 | 80 | 139 | 226 | 55 | LG+I+G4 |
| EOG092D3TIC | 433 | 73 | 103 | 410 | 23 | WAG+I+G4 |
| EOG092D1H5T | 557 | 72 | 210 | 408 | 149 | LG+I+G4 |
| EOG092D16KC | 848 | 81 | 375 | 833 | 15 | JTT+I+G4 |
| EOG092D2N6R | 191 | 78 | 101 | 170 | 21 | WAG+I+G4 |
| EOG092D1RUY | 584 | 81 | 360 | 540 | 44 | LG+F+I+G4 |
| EOG092D3NXJ | 852 | 79 | 219 | 749 | 103 | JTT+F+I+G4 |
| EOG092D1QLK | 828 | 82 | 414 | 608 | 220 | LG+I+G4 |
| EOG092D4MRW | 119 | 78 | 38 | 81 | 38 | JTT+G4 |
| EOG092D05RI | 1736 | 81 | 447 | 1679 | 57 | JTT+F+I+G4 |
| EOG092D0KJL | 1155 | 82 | 532 | 1084 | 71 | JTT+F+I+G4 |
| EOG092D04CD | 1240 | 80 | 240 | 1217 | 23 | JTT+F+I+G4 |
| EOG092D0OMH | 724 | 79 | 209 | 669 | 55 | LG+F+I+G4 |
| EOG092D3HJ3 | 276 | 82 | 114 | 148 | 128 | LG+I+G4 |
| EOG092D2FMD | 401 | 73 | 300 | 364 | 37 | LG+I+G4 |
| EOG092D29G7 | 373 | 81 | 181 | 291 | 82 | LG+F+I+G4 |
| EOG092D2MH0 | 474 | 79 | 226 | 421 | 53 | LG+I+G4 |
| EOG092D38AL | 601 | 79 | 233 | 493 | 108 | JTT+F+I+G4 |
| EOG092D2ZRK | 330 | 68 | 233 | 321 | 9 | LG+F+G4 |
| EOG092D1L48 | 905 | 83 | 211 | 843 | 62 | JTT+F+I+G4 |
| EOG092D1OKZ | 530 | 70 | 199 | 393 | 137 | LG+I+G4 |
| EOG092D4RG5 | 89 | 77 | 64 | 71 | 18 | LG+I+G4 |

Table 7: Overview of alignments of genes used for phylogenomic reconstruction. Table generated with phylociraptor. *(continued)*

| gene | length | no. of sequences | no. of parsimony informative sites | no. of variable sites | no. of fixed sites | best model |
|---|---|---|---|---|---|---|
| EOG092D2VEA | 387 | 76 | 207 | 378 | 9 | JTT+I+G4 |
| EOG092D3ASB | 314 | 80 | 189 | 301 | 13 | LG+I+G4 |
| EOG092D3M1U | 364 | 80 | 201 | 284 | 80 | JTT+I+G4 |
| EOG092D0XCU | 1007 | 74 | 316 | 992 | 15 | JTT+F+I+G4 |
| EOG092D0RST | 988 | 82 | 614 | 940 | 48 | LG+F+I+G4 |
| EOG092D092D | 748 | 63 | 436 | 729 | 19 | LG+I+G4 |
| EOG092D3SJS | 821 | 75 | 131 | 807 | 14 | JTT+I+G4 |
| EOG092D2UCZ | 226 | 80 | 120 | 191 | 35 | LG+I+G4 |
| EOG092D0F10 | 1056 | 49 | 653 | 983 | 73 | LG+I+G4 |
| EOG092D4CBH | 184 | 70 | 112 | 171 | 13 | LG+I+G4 |
| EOG092D2RAL | 313 | 78 | 153 | 268 | 45 | LG+G4 |
| EOG092D10ZS | 703 | 81 | 347 | 605 | 98 | JTT+I+G4 |
| EOG092D1G5D | 589 | 75 | 308 | 497 | 92 | LG+F+I+G4 |
| EOG092D10UA | 698 | 82 | 379 | 613 | 85 | LG+I+G4 |
| EOG092D1QZM | 511 | 77 | 262 | 444 | 67 | LG+I+G4 |
| EOG092D1VUE | 671 | 81 | 236 | 562 | 109 | LG+F+I+G4 |
| EOG092D3V2Z | 331 | 78 | 164 | 318 | 13 | LG+F+I+G4 |
| EOG092D1H9Y | 440 | 82 | 247 | 393 | 47 | LG+I+G4 |
| EOG092D128P | 993 | 78 | 370 | 986 | 7 | LG+I+G4 |
| EOG092D3BUG | 360 | 80 | 170 | 326 | 34 | JTT+I+G4 |
| EOG092D4124 | 288 | 78 | 162 | 271 | 17 | LG+I+G4 |
| EOG092D3M40 | 574 | 79 | 281 | 530 | 44 | JTT+F+I+G4 |
| EOG092D2D20 | 687 | 83 | 94 | 640 | 47 | JTT+F+I+G4 |
| EOG092D20CT | 791 | 74 | 452 | 765 | 26 | JTT+I+G4 |
| EOG092D3MB4 | 244 | 77 | 92 | 231 | 13 | LG+F+I+G4 |
| EOG092D2A6H | 439 | 75 | 294 | 410 | 29 | LG+I+G4 |
| EOG092D19A4 | 696 | 82 | 212 | 308 | 388 | LG+I+G4 |
| EOG092D3S7I | 148 | 76 | 51 | 145 | 3 | JTT+I+G4 |
| EOG092D3T80 | 200 | 83 | 116 | 152 | 48 | LG+I+G4 |
| EOG092D1RSN | 675 | 80 | 407 | 609 | 66 | JTT+I+G4 |
| EOG092D3OBO | 336 | 68 | 154 | 314 | 22 | LG+I+G4 |
| EOG092D1A3F | 586 | 83 | 330 | 520 | 66 | JTT+I+G4 |
| EOG092D4CO8 | 228 | 80 | 120 | 223 | 5 | JTTDCMut+I+G4 |
| EOG092D48Q8 | 180 | 76 | 109 | 153 | 27 | LG+I+G4 |
| EOG092D2AGS | 504 | 81 | 211 | 441 | 63 | LG+I+G4 |
| EOG092D4BKP | 178 | 83 | 97 | 132 | 46 | JTT+I+G4 |
| EOG092D0RR7 | 1027 | 68 | 745 | 1009 | 18 | JTT+F+I+G4 |
| EOG092D4JAO | 234 | 80 | 98 | 220 | 14 | LG+F+I+G4 |
| EOG092D45TL | 352 | 69 | 158 | 348 | 4 | JTTDCMut+F+I+G4 |
| EOG092D3XGU | 220 | 73 | 139 | 216 | 4 | LG+I+G4 |
| EOG092D0PYM | 751 | 83 | 267 | 479 | 272 | LG+I+G4 |
| EOG092D1UKL | 637 | 79 | 265 | 522 | 115 | JTT+I+G4 |
| EOG092D07ZA | 1058 | 76 | 581 | 907 | 151 | LG+I+G4 |
| EOG092D4F81 | 244 | 74 | 128 | 227 | 17 | LG+F+I+G4 |
| EOG092D3XC6 | 247 | 82 | 130 | 200 | 47 | Dayhoff+I+G4 |
| EOG092D2N6E | 340 | 80 | 122 | 201 | 139 | LG+I+G4 |
| EOG092D1RV3 | 708 | 83 | 256 | 673 | 35 | JTTDCMut+F+I+G4 |
| EOG092D0HKD | 1117 | 83 | 245 | 932 | 185 | JTT+I+G4 |
| EOG092D3T99 | 307 | 79 | 171 | 290 | 17 | JTTDCMut+I+G4 |
| EOG092D4ENZ | 282 | 81 | 185 | 271 | 11 | LG+I+G4 |
| EOG092D4CP0 | 265 | 82 | 179 | 242 | 23 | JTTDCMut+I+G4 |
| EOG092D0UX9 | 718 | 80 | 363 | 482 | 236 | LG+I+G4 |
| EOG092D14CG | 695 | 79 | 237 | 685 | 10 | LG+I+G4 |
| EOG092D1JIT | 495 | 81 | 270 | 415 | 80 | LG+I+G4 |
| EOG092D1QPX | 672 | 77 | 365 | 645 | 27 | LG+F+I+G4 |
| EOG092D3223 | 520 | 80 | 212 | 444 | 76 | JTT+I+G4 |
| EOG092D2P76 | 347 | 81 | 219 | 278 | 69 | LG+I+G4 |

Table 7: Overview of alignments of genes used for phylogenomic reconstruction. Table generated with phylociraptor. *(continued)*

| gene | length | no. of sequences | no. of parsimony informative sites | no. of variable sites | no. of fixed sites | best model |
|---|---|---|---|---|---|---|
| EOG092D0XB4 | 868 | 83 | 466 | 772 | 96 | LG+F+I+G4 |
| EOG092D24SJ | 451 | 78 | 274 | 367 | 84 | LG+I+G4 |
| EOG092D4SCY | 198 | 78 | 106 | 173 | 25 | LG+I+G4 |
| EOG092D28M7 | 402 | 83 | 215 | 350 | 52 | LG+F+I+G4 |
| EOG092D0WLP | 1160 | 81 | 430 | 935 | 225 | JTT+F+I+G4 |
| EOG092D30H2 | 481 | 77 | 258 | 358 | 123 | LG+I+G4 |
| EOG092D19HX | 703 | 82 | 355 | 638 | 65 | LG+F+I+G4 |
| EOG092D2HRE | 443 | 82 | 121 | 389 | 54 | LG+F+I+G4 |
| EOG092D0E6G | 766 | 81 | 281 | 488 | 278 | LG+I+G4 |
| EOG092D1XFN | 419 | 79 | 141 | 385 | 34 | WAG+F+I+G4 |
| EOG092D0IU7 | 728 | 81 | 362 | 463 | 265 | LG+I+G4 |
| EOG092D2TPW | 369 | 82 | 170 | 301 | 68 | LG+I+G4 |
| EOG092D27RI | 440 | 81 | 172 | 306 | 134 | JTT+I+G4 |
| EOG092D4E63 | 150 | 76 | 23 | 46 | 104 | LG+I+G4 |
| EOG092D4A46 | 547 | 81 | 56 | 517 | 30 | JTT+F+I+G4 |
| EOG092D3OHP | 296 | 81 | 188 | 281 | 15 | LG+I+G4 |
| EOG092D0OAE | 1131 | 81 | 558 | 1058 | 73 | JTT+I+G4 |
| EOG092D13HB | 614 | 83 | 274 | 400 | 214 | LG+I+G4 |
| EOG092D1YWG | 464 | 82 | 77 | 274 | 190 | LG+I+G4 |
| EOG092D3KSF | 265 | 78 | 163 | 203 | 62 | LG+I+G4 |
| EOG092D0OOP | 692 | 79 | 235 | 314 | 378 | LG+I+G4 |
| EOG092D4F44 | 197 | 60 | 141 | 184 | 13 | LG+G4 |
| EOG092D0CFT | 1030 | 83 | 534 | 729 | 301 | LG+I+G4 |
| EOG092D3014 | 508 | 82 | 224 | 395 | 113 | LG+F+I+G4 |
| EOG092D2S6W | 313 | 65 | 222 | 305 | 8 | LG+I+G4 |
| EOG092D3SP3 | 310 | 80 | 177 | 256 | 54 | LG+I+G4 |
| EOG092D0T75 | 657 | 77 | 375 | 515 | 142 | LG+F+I+G4 |
| EOG092D0CWO | 1013 | 82 | 596 | 728 | 285 | LG+F+I+G4 |
| EOG092D1J3R | 476 | 82 | 210 | 326 | 150 | LG+I+G4 |
| EOG092D4OVK | 170 | 79 | 85 | 151 | 19 | JTT+I+G4 |
| EOG092D25EQ | 367 | 82 | 180 | 312 | 55 | LG+I+G4 |
| EOG092D2I7R | 362 | 81 | 205 | 293 | 69 | LG+I+G4 |
| EOG092D0YQS | 722 | 82 | 452 | 607 | 115 | LG+I+G4 |
| EOG092D0431 | 1687 | 79 | 654 | 1657 | 30 | JTT+F+I+G4 |
| EOG092D2Q2D | 626 | 83 | 227 | 585 | 41 | JTT+F+I+G4 |
| EOG092D28LI | 379 | 82 | 157 | 235 | 144 | LG+I+G4 |
| EOG092D0XW1 | 1013 | 81 | 496 | 917 | 96 | JTT+F+I+G4 |
| EOG092D2FXY | 287 | 78 | 176 | 276 | 11 | LG+I+G4 |
| EOG092D4GR9 | 178 | 82 | 97 | 153 | 25 | LG+I+G4 |
| EOG092D37TR | 311 | 79 | 133 | 245 | 66 | JTTDCMut+I+G4 |
| EOG092D4383 | 188 | 80 | 63 | 111 | 77 | LG+I+G4 |
| EOG092D339Q | 428 | 80 | 184 | 362 | 66 | WAG+I+G4 |
| EOG092D2HX1 | 362 | 79 | 187 | 267 | 95 | LG+F+I+G4 |
| EOG092D0E4N | 896 | 80 | 195 | 891 | 5 | JTTDCMut+I+G4 |
| EOG092D1LC1 | 786 | 77 | 276 | 759 | 27 | JTT+F+I+G4 |
| EOG092D37V8 | 262 | 79 | 189 | 232 | 30 | LG+I+G4 |
| EOG092D07LW | 1196 | 81 | 525 | 1167 | 29 | JTT+F+I+G4 |
| EOG092D3HQ5 | 256 | 80 | 164 | 209 | 47 | LG+I+G4 |
| EOG092D4Q5Q | 136 | 76 | 43 | 109 | 27 | LG+I+G4 |
| EOG092D1Q7K | 497 | 80 | 240 | 406 | 91 | JTT+I+G4 |
| EOG092D29P8 | 357 | 78 | 164 | 320 | 37 | LG+I+G4 |
| EOG092D2N6K | 303 | 79 | 198 | 296 | 7 | LG+G4 |
| EOG092D3JQ0 | 200 | 79 | 109 | 181 | 19 | LG+I+G4 |
| EOG092D4O5W | 103 | 78 | 55 | 80 | 23 | LG+G4 |
| EOG092D1AZP | 1043 | 82 | 362 | 918 | 125 | JTT+F+I+G4 |
| EOG092D15YH | 678 | 79 | 394 | 630 | 48 | JTTDCMut+I+G4 |
| EOG092D1VPO | 467 | 82 | 217 | 327 | 140 | LG+I+G4 |

Table 7: Overview of alignments of genes used for phylogenomic reconstruction. Table generated with phylociraptor. *(continued)*

| gene | length | no. of sequences | no. of parsimony informative sites | no. of variable sites | no. of fixed sites | best model |
|---|---|---|---|---|---|---|
| EOG092D3Q3O | 240 | 66 | 102 | 221 | 19 | LG+I+G4 |
| EOG092D45MK | 240 | 78 | 121 | 224 | 16 | JTT+F+I+G4 |
| EOG092D3BOU | 477 | 81 | 121 | 458 | 19 | LG+F+I+G4 |
| EOG092D3GX0 | 305 | 57 | 139 | 292 | 13 | LG+F+I+G4 |
| EOG092D0O5Q | 757 | 83 | 367 | 575 | 182 | LG+I+G4 |
| EOG092D23P5 | 364 | 83 | 174 | 290 | 74 | JTTDCMut+I+G4 |
| EOG092D2TS7 | 501 | 81 | 243 | 449 | 52 | JTT+I+G4 |
| EOG092D2MZ4 | 323 | 83 | 125 | 303 | 20 | LG+I+G4 |
| EOG092D01YA | 2272 | 83 | 900 | 1888 | 384 | LG+I+G4 |
| EOG092D0GJX | 716 | 76 | 401 | 655 | 61 | JTT+I+G4 |
| EOG092D0B03 | 1230 | 83 | 435 | 1093 | 137 | JTT+I+G4 |
| EOG092D20QX | 410 | 75 | 198 | 408 | 2 | JTT+F+I+G4 |
| EOG092D3NFG | 288 | 81 | 165 | 281 | 7 | JTT+G4 |
| EOG092D09R6 | 952 | 74 | 593 | 769 | 183 | LG+I+G4 |
| EOG092D4I2T | 152 | 71 | 81 | 140 | 12 | JTT+I+G4 |
| EOG092D10GL | 561 | 82 | 324 | 456 | 105 | JTT+I+G4 |
| EOG092D4GCP | 325 | 81 | 135 | 304 | 21 | LG+I+G4 |
| EOG092D43HN | 188 | 75 | 107 | 148 | 40 | LG+G4 |
| EOG092D0FWO | 1048 | 78 | 889 | 1011 | 37 | LG+F+I+G4 |
| EOG092D22W1 | 648 | 81 | 221 | 512 | 136 | LG+F+I+G4 |
| EOG092D2GZ4 | 314 | 73 | 232 | 310 | 4 | JTTDCMut+G4 |
| EOG092D3TKU | 184 | 70 | 127 | 152 | 32 | LG+I+G4 |
| EOG092D49R5 | 189 | 62 | 132 | 186 | 3 | LG+F+I+G4 |
| EOG092D2J6Q | 591 | 76 | 255 | 556 | 35 | JTTDCMut+I+G4 |
| EOG092D4DNZ | 166 | 78 | 87 | 144 | 22 | LG+I+G4 |
| EOG092D37F4 | 403 | 77 | 220 | 369 | 34 | LG+F+I+G4 |
| EOG092D00VT | 2206 | 82 | 1048 | 1685 | 521 | JTT+I+G4 |
| EOG092D0OVH | 1101 | 81 | 246 | 914 | 187 | JTT+F+I+G4 |
| EOG092D37ON | 315 | 82 | 109 | 199 | 116 | JTT+I+G4 |
| EOG092D054H | 1347 | 82 | 855 | 1260 | 87 | JTT+F+I+G4 |
| EOG092D4G9R | 173 | 76 | 89 | 148 | 25 | WAG+I+G4 |
| EOG092D2RJ9 | 412 | 79 | 250 | 370 | 42 | LG+I+G4 |
| EOG092D3VHD | 232 | 81 | 137 | 184 | 48 | LG+I+G4 |
| EOG092D0HWZ | 831 | 80 | 408 | 662 | 169 | LG+F+I+G4 |
| EOG092D2NO9 | 314 | 81 | 83 | 154 | 160 | JTT+I+G4 |
| EOG092D0KGE | 1199 | 82 | 254 | 970 | 229 | JTT+F+I+G4 |
| EOG092D16B7 | 585 | 82 | 294 | 525 | 60 | LG+I+G4 |
| EOG092D0SNY | 572 | 76 | 323 | 466 | 106 | LG+I+G4 |
| EOG092D46DE | 188 | 79 | 125 | 166 | 22 | LG+I+G4 |
| EOG092D4J6A | 230 | 77 | 104 | 222 | 8 | WAG+I+G4 |
| EOG092D1HUK | 529 | 81 | 195 | 272 | 257 | LG+I+G4 |
| EOG092D4IJQ | 122 | 76 | 91 | 109 | 13 | LG+I+G4 |
| EOG092D49MT | 211 | 81 | 134 | 183 | 28 | LG+G4 |
| EOG092D1UCZ | 471 | 83 | 330 | 423 | 48 | LG+F+I+G4 |
| EOG092D4MX9 | 142 | 78 | 105 | 132 | 10 | LG+G4 |
| EOG092D2PES | 330 | 83 | 177 | 307 | 23 | LG+I+G4 |
| EOG092D3I0C | 239 | 81 | 136 | 166 | 73 | LG+I+G4 |
| EOG092D3QR7 | 299 | 79 | 122 | 215 | 84 | LG+I+G4 |
| EOG092D2YGY | 402 | 82 | 231 | 324 | 78 | LG+I+G4 |
| EOG092D1GUU | 825 | 81 | 396 | 675 | 150 | JTT+I+G4 |
| EOG092D3E5G | 256 | 72 | 165 | 212 | 44 | JTTDCMut+I+G4 |
| EOG092D31U4 | 389 | 77 | 93 | 371 | 18 | LG+F+I+G4 |
| EOG092D2BYU | 426 | 81 | 187 | 274 | 152 | LG+I+G4 |
| EOG092D2QY5 | 847 | 82 | 322 | 749 | 98 | JTT+F+I+G4 |
| EOG092D41CK | 303 | 77 | 166 | 258 | 45 | LG+F+I+G4 |
| EOG092D23JK | 514 | 83 | 290 | 452 | 62 | LG+I+G4 |
| EOG092D44MC | 190 | 73 | 141 | 178 | 12 | LG+G4 |

Table 7: Overview of alignments of genes used for phylogenomic reconstruction. Table generated with phylociraptor. *(continued)*

| gene | length | no. of sequences | no. of parsimony informative sites | no. of variable sites | no. of fixed sites | best model |
|---|---|---|---|---|---|---|
| EOG092D2HF9 | 417 | 82 | 197 | 280 | 137 | LG+F+I+G4 |
| EOG092D0A17 | 1127 | 82 | 291 | 1022 | 105 | JTT+I+G4 |
| EOG092D3XG3 | 221 | 79 | 99 | 158 | 63 | Dayhoff+I+G4 |
| EOG092D00LL | 2345 | 82 | 630 | 1045 | 1300 | LG+I+G4 |
| EOG092D0H4I | 1026 | 82 | 341 | 984 | 42 | JTT+F+I+G4 |
| EOG092D49G9 | 211 | 82 | 110 | 208 | 3 | JTTDCMut+I+G4 |
| EOG092D3OPM | 248 | 77 | 180 | 212 | 36 | LG+F+I+G4 |
| EOG092D37JC | 292 | 81 | 194 | 275 | 17 | LG+I+G4 |
| EOG092D3ZUC | 245 | 79 | 151 | 216 | 29 | LG+I+G4 |
| EOG092D1R2W | 794 | 83 | 395 | 742 | 52 | JTTDCMut+I+G4 |
| EOG092D0Y8Q | 881 | 80 | 474 | 764 | 117 | JTTDCMut+I+G4 |
| EOG092D041E | 1269 | 71 | 581 | 1228 | 41 | LG+F+I+G4 |
| EOG092D11EA | 674 | 83 | 333 | 541 | 133 | LG+I+G4 |
| EOG092D4FU5 | 189 | 77 | 103 | 167 | 22 | WAG+I+G4 |
| EOG092D41S5 | 485 | 82 | 69 | 388 | 97 | LG+I+G4 |
| EOG092D3GEY | 354 | 80 | 211 | 308 | 46 | LG+I+G4 |
| EOG092D26ZW | 414 | 82 | 178 | 301 | 113 | JTTDCMut+I+G4 |
| EOG092D2U1L | 335 | 81 | 206 | 294 | 41 | LG+I+G4 |
| EOG092D1UL3 | 669 | 83 | 287 | 591 | 78 | JTT+F+I+G4 |
| EOG092D0643 | 1358 | 82 | 730 | 1320 | 38 | LG+F+I+G4 |
| EOG092D1TC1 | 589 | 81 | 169 | 551 | 38 | WAG+I+G4 |
| EOG092D3S4W | 395 | 83 | 141 | 373 | 22 | WAG+I+G4 |
| EOG092D06GR | 1029 | 81 | 575 | 917 | 112 | LG+I+G4 |
| EOG092D2ZDK | 324 | 50 | 110 | 226 | 98 | LG+G4 |
| EOG092D225O | 493 | 82 | 308 | 407 | 86 | JTTDCMut+I+G4 |
| EOG092D09PM | 1142 | 83 | 390 | 1069 | 73 | JTT+F+I+G4 |
| EOG092D467U | 347 | 65 | 206 | 345 | 2 | LG+I+G4 |
| EOG092D4EPH | 190 | 81 | 97 | 157 | 33 | LG+I+G4 |
| EOG092D4O6D | 119 | 80 | 92 | 110 | 9 | JTT+G4 |
| EOG092D05LP | 1249 | 81 | 896 | 1072 | 177 | JTTDCMut+F+I+G4 |
| EOG092D3G4V | 193 | 80 | 85 | 182 | 11 | LG+I+G4 |
| EOG092D0LL2 | 813 | 81 | 410 | 684 | 129 | LG+F+I+G4 |
| EOG092D19T7 | 759 | 78 | 437 | 636 | 123 | LG+I+G4 |
| EOG092D3K9O | 269 | 80 | 110 | 193 | 76 | LG+I+G4 |
| EOG092D27I5 | 386 | 81 | 182 | 314 | 72 | LG+I+G4 |
| EOG092D1Q94 | 669 | 80 | 430 | 612 | 57 | JTT+I+G4 |
| EOG092D2OLS | 728 | 82 | 126 | 696 | 32 | JTT+I+G4 |
| EOG092D4BSZ | 175 | 78 | 78 | 116 | 59 | LG+I+G4 |
| EOG092D38RE | 403 | 82 | 233 | 374 | 29 | LG+I+G4 |
| EOG092D329B | 532 | 80 | 355 | 501 | 31 | LG+F+I+G4 |
| EOG092D25JG | 445 | 79 | 256 | 383 | 62 | LG+I+G4 |
| EOG092D116A | 789 | 77 | 485 | 772 | 17 | JTT+I+G4 |
| EOG092D4HT4 | 209 | 77 | 104 | 199 | 10 | LG+F+I+G4 |
| EOG092D36PK | 333 | 70 | 143 | 277 | 56 | LG+F+I+G4 |
| EOG092D47GJ | 194 | 81 | 113 | 162 | 32 | LG+I+G4 |
| EOG092D20Y4 | 523 | 81 | 304 | 433 | 90 | LG+I+G4 |
| EOG092D2457 | 1095 | 81 | 191 | 1067 | 28 | JTT+F+I+G4 |
| EOG092D034L | 2042 | 82 | 456 | 2015 | 27 | JTT+F+I+G4 |
| EOG092D12AA | 570 | 82 | 374 | 479 | 91 | LG+I+G4 |
| EOG092D1OL9 | 808 | 79 | 363 | 598 | 210 | JTTDCMut+I+G4 |
| EOG092D08FO | 1178 | 80 | 652 | 890 | 288 | JTT+F+I+G4 |
| EOG092D1ZGE | 763 | 83 | 295 | 716 | 47 | JTT+F+I+G4 |
| EOG092D3UPV | 210 | 72 | 49 | 75 | 135 | JTT+I+G4 |
| EOG092D4J2E | 229 | 76 | 87 | 211 | 18 | JTT+F+I+G4 |
| EOG092D2RJR | 759 | 82 | 288 | 616 | 143 | JTT+F+I+G4 |
| EOG092D09TB | 974 | 81 | 512 | 772 | 202 | LG+I+G4 |
| EOG092D47XX | 181 | 83 | 101 | 160 | 21 | LG+I+G4 |

Table 7: Overview of alignments of genes used for phylogenomic reconstruction. Table generated with phylociraptor. *(continued)*

| gene | length | no. of sequences | no. of parsimony informative sites | no. of variable sites | no. of fixed sites | best model |
|---|---|---|---|---|---|---|
| EOG092D3VPV | 552 | 81 | 211 | 501 | 51 | JTT+F+I+G4 |
| EOG092D0ZNF | 929 | 81 | 413 | 790 | 139 | JTT+I+G4 |
| EOG092D4X9E | 96 | 80 | 38 | 72 | 24 | LG+G4 |
| EOG092D2F5H | 423 | 78 | 218 | 373 | 50 | LG+I+G4 |
| EOG092D2HI6 | 562 | 80 | 191 | 510 | 52 | LG+I+G4 |
| EOG092D48MS | 302 | 79 | 194 | 289 | 13 | LG+I+G4 |
| EOG092D1KWA | 583 | 83 | 360 | 491 | 92 | LG+F+I+G4 |
| EOG092D2YOQ | 450 | 78 | 262 | 408 | 42 | LG+I+G4 |
| EOG092D4CCD | 147 | 78 | 47 | 101 | 46 | LG+G4 |
| EOG092D3YZ8 | 150 | 39 | 104 | 133 | 17 | LG+I+G4 |
| EOG092D42VQ | 229 | 54 | 136 | 192 | 37 | LG+I+G4 |
| EOG092D14XV | 895 | 80 | 385 | 873 | 22 | JTT+F+I+G4 |
| EOG092D247K | 398 | 76 | 231 | 332 | 66 | LG+I+G4 |
| EOG092D23EJ | 473 | 82 | 242 | 391 | 82 | LG+F+I+G4 |
| EOG092D1Q2Q | 289 | 82 | 216 | 276 | 13 | LG+I+G4 |
| EOG092D0NUZ | 683 | 82 | 305 | 495 | 188 | LG+I+G4 |
| EOG092D13QH | 639 | 79 | 363 | 468 | 171 | LG+I+G4 |
| EOG092D4557 | 175 | 80 | 48 | 76 | 99 | LG+I+G4 |
| EOG092D0DYZ | 831 | 73 | 479 | 754 | 77 | JTT+I+G4 |
| EOG092D1GCX | 681 | 80 | 345 | 555 | 126 | JTTDCMut+I+G4 |
| EOG092D0LJL | 951 | 82 | 549 | 898 | 53 | LG+F+I+G4 |
| EOG092D39BT | 192 | 79 | 114 | 161 | 31 | JTT+I+G4 |
| EOG092D24WU | 724 | 72 | 406 | 653 | 71 | LG+I+G4 |
| EOG092D0JFU | 901 | 67 | 123 | 899 | 2 | JTT+F+I+G4 |
| EOG092D2B2Y | 418 | 79 | 267 | 340 | 78 | LG+I+G4 |
| EOG092D3FNY | 551 | 82 | 217 | 519 | 32 | JTT+F+I+G4 |
| EOG092D261T | 902 | 82 | 572 | 768 | 134 | LG+I+G4 |
| EOG092D3OQG | 309 | 75 | 166 | 226 | 83 | JTT+I+G4 |
| EOG092D13AE | 560 | 81 | 405 | 504 | 56 | LG+I+G4 |
| EOG092D0H69 | 938 | 75 | 305 | 927 | 11 | LG+F+I+G4 |
| EOG092D1EUT | 200 | 81 | 94 | 136 | 64 | LG+I+G4 |
| EOG092D4MKM | 182 | 80 | 79 | 138 | 44 | LG+G4 |
| EOG092D03HZ | 1765 | 77 | 1290 | 1677 | 88 | LG+F+I+G4 |
| EOG092D4EV2 | 292 | 81 | 122 | 286 | 6 | JTTDCMut+F+I+G4 |
| EOG092D1QLG | 748 | 82 | 257 | 637 | 111 | JTT+F+I+G4 |
| EOG092D0EGX | 945 | 82 | 360 | 663 | 282 | JTT+F+I+G4 |
| EOG092D4CHM | 673 | 82 | 257 | 663 | 10 | JTTDCMut+I+G4 |
| EOG092D2TLM | 317 | 68 | 225 | 277 | 40 | LG+I+G4 |
| EOG092D2M1W | 386 | 78 | 202 | 332 | 54 | JTT+I+G4 |
| EOG092D3TL4 | 205 | 81 | 63 | 116 | 89 | LG+I+G4 |
| EOG092D03AT | 1681 | 82 | 235 | 1639 | 42 | JTT+F+I+G4 |
| EOG092D09SW | 981 | 80 | 618 | 930 | 51 | JTT+F+I+G4 |
| EOG092D1ZEW | 545 | 79 | 316 | 487 | 58 | LG+F+I+G4 |
| EOG092D0JGD | 956 | 83 | 498 | 895 | 61 | JTT+F+I+G4 |
| EOG092D4APA | 116 | 33 | 91 | 114 | 2 | LG+G4 |
| EOG092D1NW2 | 516 | 66 | 282 | 452 | 64 | LG+F+I+G4 |
| EOG092D4ISY | 124 | 51 | 19 | 71 | 53 | Dayhoff+G4 |
| EOG092D2XUX | 272 | 79 | 180 | 243 | 29 | LG+F+I+G4 |
| EOG092D2806 | 498 | 81 | 208 | 449 | 49 | JTT+F+I+G4 |
| EOG092D4V60 | 68 | 78 | 44 | 50 | 18 | LG+I+G4 |
| EOG092D105K | 655 | 83 | 292 | 554 | 101 | JTTDCMut+I+G4 |
| EOG092D3Y1Q | 235 | 77 | 165 | 214 | 21 | LG+I+G4 |
| EOG092D3UQ8 | 257 | 78 | 166 | 199 | 58 | LG+I+G4 |
| EOG092D0CUY | 1434 | 64 | 314 | 1429 | 5 | LG+F+I+G4 |
| EOG092D1UWS | 710 | 83 | 273 | 635 | 75 | JTT+F+I+G4 |
| EOG092D0UKN | 950 | 81 | 577 | 850 | 100 | LG+I+G4 |
| EOG092D0S50 | 1111 | 82 | 374 | 1046 | 65 | JTT+F+I+G4 |

Table 7: Overview of alignments of genes used for phylogenomic reconstruction. Table generated with phylociraptor. *(continued)*

| gene | length | no. of sequences | no. of parsimony informative sites | no. of variable sites | no. of fixed sites | best model |
|---|---|---|---|---|---|---|
| EOG092D4B4O | 413 | 80 | 188 | 365 | 48 | LG+I+G4 |
| EOG092D2ERC | 589 | 82 | 213 | 580 | 9 | LG+I+G4 |
| EOG092D3JQI | 394 | 82 | 202 | 331 | 63 | JTT+I+G4 |
| EOG092D0AE9 | 1390 | 82 | 697 | 1138 | 252 | JTT+F+I+G4 |
| EOG092D1M2S | 553 | 83 | 394 | 541 | 12 | LG+F+I+G4 |
| EOG092D1CM1 | 627 | 82 | 315 | 617 | 10 | JTT+F+I+G4 |
| EOG092D4HJN | 114 | 80 | 69 | 99 | 15 | LG+I+G4 |
| EOG092D0FXM | 640 | 69 | 247 | 611 | 29 | LG+F+I+G4 |
| EOG092D430G | 336 | 80 | 88 | 307 | 29 | JTT+F+I+G4 |
| EOG092D1KRH | 665 | 83 | 83 | 493 | 172 | JTT+I+G4 |
| EOG092D0S6R | 788 | 80 | 415 | 698 | 90 | LG+I+G4 |
| EOG092D3G7O | 559 | 76 | 309 | 488 | 71 | LG+F+I+G4 |
| EOG092D4J1H | 138 | 80 | 46 | 78 | 60 | LG+I+G4 |
| EOG092D04KS | 1711 | 79 | 682 | 1528 | 183 | JTT+F+I+G4 |
| EOG092D4S9W | 133 | 82 | 90 | 116 | 17 | LG+I+G4 |
| EOG092D4PXN | 120 | 82 | 74 | 98 | 22 | LG+I+G4 |
| EOG092D3DNZ | 336 | 81 | 163 | 255 | 81 | JTTDCMut+I+G4 |
| EOG092D4E28 | 165 | 73 | 98 | 132 | 33 | LG+I+G4 |
| EOG092D2YAG | 425 | 77 | 289 | 406 | 19 | LG+I+G4 |
| EOG092D31CV | 283 | 82 | 127 | 231 | 52 | LG+I+G4 |
| EOG092D4A4K | 143 | 78 | 78 | 105 | 38 | LG+G4 |
| EOG092D3F6H | 256 | 78 | 135 | 196 | 60 | LG+I+G4 |
| EOG092D18SO | 505 | 79 | 334 | 473 | 32 | LG+F+I+G4 |
| EOG092D0T8J | 667 | 82 | 342 | 610 | 57 | JTT+I+G4 |
| EOG092D3DMS | 855 | 81 | 234 | 802 | 53 | JTT+F+I+G4 |
| EOG092D3R6J | 230 | 81 | 102 | 152 | 78 | LG+I+G4 |
| EOG092D2MK3 | 593 | 78 | 388 | 560 | 33 | JTT+F+I+G4 |
| EOG092D33VX | 307 | 81 | 119 | 298 | 9 | LG+F+I+G4 |
| EOG092D3R78 | 311 | 79 | 108 | 302 | 9 | WAG+I+G4 |
| EOG092D3AVM | 608 | 80 | 129 | 536 | 72 | JTT+F+I+G4 |
| EOG092D2PJJ | 443 | 79 | 241 | 335 | 108 | LG+I+G4 |
| EOG092D3J6B | 245 | 81 | 67 | 141 | 104 | LG+I+G4 |
| EOG092D0320 | 1381 | 76 | 299 | 1330 | 51 | JTT+I+G4 |
| EOG092D3DN6 | 298 | 75 | 93 | 172 | 126 | LG+I+G4 |
| EOG092D0KG3 | 1293 | 71 | 434 | 1266 | 27 | JTT+F+I+G4 |
| EOG092D3M3Y | 445 | 82 | 50 | 438 | 7 | JTT+I+G4 |
| EOG092D1SLT | 422 | 56 | 286 | 374 | 48 | LG+I+G4 |
| EOG092D01WX | 1865 | 81 | 472 | 1793 | 72 | JTT+F+I+G4 |
| EOG092D2KS9 | 406 | 78 | 280 | 361 | 45 | LG+I+G4 |
| EOG092D1YHF | 467 | 83 | 201 | 364 | 103 | LG+I+G4 |
| EOG092D0H0I | 1103 | 73 | 572 | 1055 | 48 | JTT+F+I+G4 |
| EOG092D4IME | 223 | 81 | 122 | 196 | 27 | LG+I+G4 |
| EOG092D4MUW | 116 | 73 | 67 | 103 | 13 | LG+I+G4 |
| EOG092D25WL | 346 | 82 | 213 | 276 | 70 | LG+I+G4 |
| EOG092D4JQM | 300 | 62 | 60 | 287 | 13 | JTT+I+G4 |
| EOG092D4AOT | 276 | 80 | 207 | 261 | 15 | LG+F+I+G4 |
| EOG092D3CSS | 296 | 83 | 178 | 257 | 39 | LG+I+G4 |
| EOG092D0S8J | 759 | 66 | 410 | 705 | 54 | LG+F+I+G4 |
| EOG092D2Q89 | 419 | 79 | 94 | 409 | 10 | LG+I+G4 |
| EOG092D0WLN | 737 | 80 | 277 | 690 | 47 | LG+F+I+G4 |
| EOG092D00R3 | 2203 | 82 | 1010 | 1452 | 751 | LG+I+G4 |
| EOG092D3YU3 | 453 | 80 | 166 | 418 | 35 | JTT+I+G4 |
| EOG092D1QEF | 708 | 80 | 357 | 618 | 90 | LG+I+G4 |
| EOG092D3NPH | 233 | 81 | 139 | 190 | 43 | LG+I+G4 |
| EOG092D4J7I | 116 | 83 | 71 | 92 | 24 | JTT+I+G4 |
| EOG092D3K72 | 246 | 78 | 145 | 175 | 71 | LG+F+I+G4 |
| EOG092D28HS | 474 | 80 | 189 | 287 | 187 | WAG+I+G4 |

Table 7: Overview of alignments of genes used for phylogenomic reconstruction. Table generated with phylociraptor. *(continued)*

| gene | length | no. of sequences | no. of parsimony informative sites | no. of variable sites | no. of fixed sites | best model |
|---|---|---|---|---|---|---|
| EOG092D3VON | 242 | 80 | 107 | 194 | 48 | LG+I+G4 |
| EOG092D1A1J | 553 | 79 | 343 | 473 | 80 | LG+I+G4 |
| EOG092D44FK | 1236 | 80 | 127 | 1222 | 14 | JTTDCMut+F+I+G4 |
| EOG092D07ZP | 996 | 81 | 569 | 928 | 68 | LG+F+I+G4 |
| EOG092D26IC | 349 | 76 | 186 | 321 | 28 | LG+I+G4 |
| EOG092D3TNE | 214 | 75 | 153 | 197 | 17 | LG+I+G4 |
| EOG092D2OL1 | 361 | 82 | 225 | 301 | 60 | LG+I+G4 |
| EOG092D4HAV | 183 | 81 | 98 | 124 | 59 | LG+I+G4 |
| EOG092D4WYA | 128 | 72 | 42 | 122 | 6 | JTT+I+G4 |
| EOG092D3KII | 262 | 79 | 129 | 198 | 64 | LG+I+G4 |
| EOG092D23UZ | 628 | 82 | 154 | 462 | 166 | JTT+I+G4 |
| EOG092D0FGS | 940 | 80 | 497 | 766 | 174 | LG+F+I+G4 |
| EOG092D2WRH | 291 | 78 | 239 | 273 | 18 | LG+I+G4 |
| EOG092D2COI | 1184 | 81 | 785 | 1127 | 57 | LG+I+G4 |
| EOG092D2O8B | 1026 | 81 | 283 | 954 | 72 | JTT+F+I+G4 |
| EOG092D46HE | 279 | 78 | 127 | 259 | 20 | WAG+G4 |
| EOG092D4OAJ | 154 | 80 | 72 | 120 | 34 | WAG+I+G4 |
| EOG092D272T | 506 | 82 | 169 | 470 | 36 | LG+I+G4 |
| EOG092D49WK | 319 | 77 | 188 | 276 | 43 | LG+F+I+G4 |
| EOG092D11XU | 652 | 73 | 356 | 549 | 103 | LG+I+G4 |
| EOG092D3PF3 | 547 | 81 | 231 | 505 | 42 | LG+I+G4 |
| EOG092D344A | 316 | 83 | 171 | 267 | 49 | LG+I+G4 |
| EOG092D0BMY | 936 | 81 | 506 | 687 | 249 | LG+I+G4 |
| EOG092D0UJ2 | 593 | 77 | 280 | 576 | 17 | JTT+I+G4 |
| EOG092D29KI | 449 | 78 | 168 | 219 | 230 | LG+I+G4 |
| EOG092D3N3B | 279 | 77 | 138 | 229 | 50 | Dayhoff+I+G4 |
| EOG092D412G | 170 | 80 | 90 | 134 | 36 | LG+I+G4 |
| EOG092D47Y3 | 187 | 69 | 77 | 145 | 42 | LG+I+G4 |
| EOG092D31X5 | 392 | 60 | 239 | 315 | 77 | LG+I+G4 |
| EOG092D04HM | 1476 | 82 | 527 | 1408 | 68 | JTT+F+I+G4 |
| EOG092D0GXN | 909 | 83 | 407 | 742 | 167 | LG+F+I+G4 |
| EOG092D3F1G | 547 | 82 | 210 | 518 | 29 | JTT+I+G4 |
| EOG092D4D9K | 409 | 74 | 224 | 380 | 29 | LG+I+G4 |
| EOG092D49VZ | 165 | 74 | 74 | 94 | 71 | LG+I+G4 |
| EOG092D0KBR | 824 | 80 | 454 | 679 | 145 | LG+F+I+G4 |
| EOG092D3HUP | 243 | 80 | 125 | 209 | 34 | LG+I+G4 |
| EOG092D3K5F | 312 | 79 | 112 | 179 | 133 | LG+I+G4 |
| EOG092D3KV2 | 331 | 81 | 173 | 242 | 89 | LG+F+I+G4 |
| EOG092D0J9G | 913 | 80 | 334 | 540 | 373 | LG+I+G4 |
| EOG092D29QC | 548 | 78 | 271 | 535 | 13 | LG+F+I+G4 |
| EOG092D1YZO | 416 | 82 | 151 | 281 | 135 | JTT+I+G4 |
| EOG092D3LCX | 231 | 79 | 138 | 185 | 46 | LG+I+G4 |
| EOG092D1GVT | 557 | 81 | 159 | 259 | 298 | LG+F+I+G4 |
| EOG092D0072 | 4255 | 81 | 2225 | 3204 | 1051 | JTT+F+I+G4 |
| EOG092D4JRM | 188 | 78 | 72 | 174 | 14 | LG+F+I+G4 |
| EOG092D07D8 | 1708 | 81 | 237 | 1598 | 110 | JTT+I+G4 |
| EOG092D3P54 | 237 | 76 | 133 | 204 | 33 | JTT+F+I+G4 |
| EOG092D1MNK | 486 | 82 | 174 | 307 | 179 | LG+I+G4 |
| EOG092D3QV4 | 219 | 80 | 48 | 120 | 99 | LG+I+G4 |
| EOG092D2CAF | 355 | 82 | 131 | 274 | 81 | LG+I+G4 |
| EOG092D4ASP | 226 | 60 | 139 | 203 | 23 | LG+G4 |
| EOG092D3H8K | 297 | 82 | 134 | 199 | 98 | LG+I+G4 |
| EOG092D4I8P | 152 | 79 | 81 | 113 | 39 | LG+I+G4 |
| EOG092D2LX2 | 346 | 79 | 139 | 220 | 126 | LG+I+G4 |
| EOG092D4S3D | 169 | 60 | 123 | 153 | 16 | LG+G4 |
| EOG092D3SP4 | 339 | 76 | 203 | 268 | 71 | LG+F+I+G4 |
| EOG092D3E1M | 605 | 80 | 204 | 557 | 48 | JTTDCMut+I+G4 |

Table 7: Overview of alignments of genes used for phylogenomic reconstruction. Table generated with phylociraptor. *(continued)*

| gene | length | no. of sequences | no. of parsimony informative sites | no. of variable sites | no. of fixed sites | best model |
|---|---|---|---|---|---|---|
| EOG092D1S33 | 495 | 81 | 237 | 345 | 150 | LG+I+G4 |
| EOG092D0K9C | 857 | 78 | 284 | 803 | 54 | JTT+F+I+G4 |
| EOG092D1G2Z | 564 | 80 | 108 | 421 | 143 | LG+F+I+G4 |
| EOG092D4CIZ | 255 | 78 | 143 | 239 | 16 | LG+F+I+G4 |
| EOG092D164Q | 539 | 75 | 306 | 464 | 75 | LG+F+I+G4 |
| EOG092D0LVZ | 844 | 82 | 561 | 758 | 86 | LG+F+I+G4 |
| EOG092D33FU | 424 | 79 | 253 | 372 | 52 | LG+I+G4 |
| EOG092D0BYJ | 1048 | 82 | 486 | 670 | 378 | LG+F+I+G4 |
| EOG092D0T3V | 593 | 77 | 278 | 537 | 56 | LG+F+I+G4 |
| EOG092D2XS7 | 465 | 83 | 327 | 424 | 41 | LG+I+G4 |
| EOG092D0OXK | 838 | 80 | 459 | 636 | 202 | LG+I+G4 |
| EOG092D1CKF | 876 | 81 | 376 | 745 | 131 | LG+I+G4 |
| EOG092D1N0P | 744 | 81 | 484 | 705 | 39 | LG+F+I+G4 |
| EOG092D07JG | 1086 | 75 | 494 | 758 | 328 | LG+I+G4 |
| EOG092D4H8I | 133 | 64 | 80 | 116 | 17 | LG+G4 |
| EOG092D1YID | 478 | 83 | 193 | 350 | 128 | LG+I+G4 |
| EOG092D1ZEJ | 442 | 81 | 131 | 356 | 86 | LG+I+G4 |
| EOG092D056B | 1222 | 82 | 349 | 1193 | 29 | WAG+F+I+G4 |
| EOG092D0HGD | 1189 | 81 | 209 | 1163 | 26 | JTT+F+I+G4 |
| EOG092D4CW1 | 200 | 51 | 59 | 155 | 45 | JTT+I+G4 |
| EOG092D065A | 1269 | 72 | 705 | 1198 | 71 | LG+I+G4 |
| EOG092D2925 | 685 | 80 | 303 | 644 | 41 | JTT+F+I+G4 |
| EOG092D406X | 201 | 82 | 103 | 159 | 42 | JTT+I+G4 |
| EOG092D0QAH | 929 | 81 | 564 | 851 | 78 | LG+I+G4 |
| EOG092D1O7A | 495 | 82 | 149 | 279 | 216 | LG+I+G4 |
| EOG092D03RY | 1318 | 79 | 503 | 1302 | 16 | JTT+F+I+G4 |
| EOG092D0LJK | 853 | 79 | 369 | 624 | 229 | LG+F+I+G4 |
| EOG092D2BJH | 524 | 82 | 266 | 365 | 159 | LG+I+G4 |
| EOG092D3K5U | 305 | 78 | 171 | 294 | 11 | LG+I+G4 |
| EOG092D3MS3 | 213 | 81 | 157 | 202 | 11 | LG+I+G4 |
| EOG092D03PE | 1494 | 76 | 886 | 1441 | 53 | JTT+F+I+G4 |
| EOG092D4MP9 | 123 | 81 | 55 | 88 | 35 | LG+G4 |
| EOG092D0ILO | 900 | 81 | 326 | 878 | 22 | JTT+F+I+G4 |
| EOG092D3X2Q | 228 | 75 | 162 | 200 | 28 | JTT+I+G4 |
| EOG092D2G7T | 356 | 77 | 198 | 241 | 115 | LG+I+G4 |
| EOG092D4GG0 | 133 | 74 | 106 | 129 | 4 | LG+I+G4 |
| EOG092D32SI | 301 | 77 | 155 | 268 | 33 | LG+F+I+G4 |
| EOG092D4J3B | 105 | 73 | 67 | 84 | 21 | LG+I+G4 |
| EOG092D1H8I | 518 | 78 | 209 | 457 | 61 | LG+F+I+G4 |
| EOG092D1VKT | 534 | 81 | 264 | 449 | 85 | LG+I+G4 |
| EOG092D3PPI | 252 | 75 | 73 | 205 | 47 | VT+I+G4 |
| EOG092D4QSJ | 117 | 74 | 70 | 99 | 18 | LG+I+G4 |
| EOG092D1MQ6 | 571 | 82 | 186 | 256 | 315 | LG+I+G4 |
| EOG092D0P0O | 1013 | 75 | 305 | 1001 | 12 | JTT+F+I+G4 |
| EOG092D2Z5Z | 298 | 80 | 144 | 205 | 93 | LG+I+G4 |
| EOG092D2FDT | 404 | 74 | 260 | 390 | 14 | LG+F+I+G4 |
| EOG092D2QYY | 360 | 76 | 189 | 342 | 18 | LG+F+I+G4 |
| EOG092D23WW | 424 | 81 | 229 | 372 | 52 | LG+I+G4 |
| EOG092D2O1N | 999 | 80 | 280 | 976 | 23 | JTT+F+I+G4 |
| EOG092D3THJ | 272 | 77 | 141 | 238 | 34 | LG+I+G4 |
| EOG092D0R9A | 774 | 80 | 417 | 732 | 42 | JTT+F+I+G4 |
| EOG092D0ULK | 1003 | 83 | 373 | 846 | 157 | JTT+I+G4 |
| EOG092D0905 | 1276 | 80 | 480 | 1197 | 79 | JTT+F+I+G4 |
| EOG092D3PHH | 146 | 80 | 84 | 141 | 5 | LG+I+G4 |
| EOG092D40TG | 200 | 77 | 87 | 134 | 66 | LG+I+G4 |
| EOG092D080M | 1037 | 80 | 570 | 865 | 172 | LG+I+G4 |
| EOG092D02KN | 1828 | 81 | 576 | 1739 | 89 | JTT+I+G4 |

Table 7: Overview of alignments of genes used for phylogenomic reconstruction. Table generated with phylociraptor. *(continued)*

| gene | length | no. of sequences | no. of parsimony informative sites | no. of variable sites | no. of fixed sites | best model |
|---|---|---|---|---|---|---|
| EOG092D2AZW | 479 | 78 | 179 | 435 | 44 | JTTDCMut+I+G4 |
| EOG092D36RY | 314 | 76 | 146 | 293 | 21 | LG+G4 |
| EOG092D4241 | 322 | 79 | 124 | 302 | 20 | LG+I+G4 |
| EOG092D3NKD | 322 | 77 | 114 | 300 | 22 | JTTDCMut+F+I+G4 |
| EOG092D3X8S | 185 | 72 | 116 | 172 | 13 | WAG+I+G4 |
| EOG092D4X1O | 163 | 81 | 79 | 135 | 28 | JTT+F+I+G4 |
| EOG092D0UFO | 770 | 83 | 412 | 626 | 144 | WAG+I+G4 |
| EOG092D0W9K | 600 | 81 | 205 | 381 | 219 | LG+I+G4 |
| EOG092D0S5N | 609 | 81 | 302 | 605 | 4 | LG+I+G4 |
| EOG092D1AOO | 308 | 74 | 159 | 217 | 91 | LG+I+G4 |
| EOG092D2VWC | 382 | 81 | 129 | 377 | 5 | LG+F+I+G4 |
| EOG092D4145 | 343 | 81 | 139 | 297 | 46 | LG+F+I+G4 |
| EOG092D43XW | 200 | 82 | 116 | 187 | 13 | LG+I+G4 |
| EOG092D13PM | 709 | 82 | 364 | 600 | 109 | JTT+F+I+G4 |
| EOG092D45IW | 170 | 78 | 113 | 161 | 9 | LG+I+G4 |
| EOG092D3HVD | 293 | 81 | 182 | 244 | 49 | LG+I+G4 |
| EOG092D0Q00 | 1074 | 79 | 642 | 1010 | 64 | LG+I+G4 |
| EOG092D19EB | 1730 | 81 | 391 | 1620 | 110 | JTT+F+I+G4 |
| EOG092D3HBU | 511 | 76 | 272 | 506 | 5 | LG+F+I+G4 |
| EOG092D3WPA | 291 | 79 | 141 | 235 | 56 | JTT+I+G4 |
| EOG092D2LER | 407 | 78 | 152 | 329 | 78 | LG+F+I+G4 |
| EOG092D4PH2 | 197 | 60 | 129 | 172 | 25 | JTT+I+G4 |
| EOG092D0ANR | 1776 | 83 | 801 | 1662 | 114 | JTT+F+I+G4 |
| EOG092D1WA5 | 487 | 81 | 294 | 407 | 80 | LG+I+G4 |
| EOG092D28YM | 640 | 83 | 142 | 598 | 42 | JTT+F+I+G4 |
| EOG092D0ZVY | 644 | 77 | 254 | 518 | 126 | JTT+F+I+G4 |
| EOG092D0DNM | 1513 | 83 | 508 | 1214 | 299 | JTT+F+I+G4 |
| EOG092D1RSS | 312 | 83 | 154 | 285 | 27 | LG+I+G4 |
| EOG092D1AOA | 535 | 71 | 258 | 449 | 86 | LG+I+G4 |
| EOG092D32P7 | 263 | 78 | 124 | 209 | 54 | LG+I+G4 |
| EOG092D4M0K | 128 | 80 | 49 | 82 | 46 | Dayhoff+I+G4 |
| EOG092D05Q0 | 1205 | 83 | 657 | 1132 | 73 | JTT+I+G4 |
| EOG092D3XA6 | 351 | 82 | 173 | 317 | 34 | LG+F+I+G4 |
| EOG092D1884 | 836 | 82 | 178 | 700 | 136 | JTT+F+I+G4 |
| EOG092D3IJD | 399 | 83 | 237 | 361 | 38 | JTT+I+G4 |
| EOG092D05X9 | 1356 | 82 | 890 | 1198 | 158 | LG+I+G4 |
| EOG092D0NP5 | 1193 | 82 | 579 | 1162 | 31 | JTT+I+G4 |
| EOG092D048Q | 1877 | 80 | 479 | 1842 | 35 | JTT+F+I+G4 |
| EOG092D3Z6V | 511 | 77 | 267 | 418 | 93 | JTT+F+I+G4 |
| EOG092D0Q23 | 796 | 76 | 273 | 521 | 275 | LG+I+G4 |
| EOG092D3WSA | 297 | 83 | 105 | 241 | 56 | JTT+F+I+G4 |
| EOG092D40PM | 161 | 77 | 79 | 143 | 18 | LG+I+G4 |
| EOG092D2RFR | 365 | 76 | 160 | 215 | 150 | LG+I+G4 |
| EOG092D360D | 421 | 81 | 225 | 398 | 23 | LG+I+G4 |
| EOG092D4UA6 | 75 | 74 | 40 | 68 | 7 | LG+G4 |
| EOG092D2VYQ | 309 | 62 | 195 | 291 | 18 | LG+I+G4 |
| EOG092D3LGX | 242 | 41 | 170 | 226 | 16 | LG+I+G4 |
| EOG092D485T | 268 | 80 | 101 | 227 | 41 | JTTDCMut+F+I+G4 |
| EOG092D3NUF | 306 | 80 | 92 | 252 | 54 | Dayhoff+I+G4 |
| EOG092D4OIQ | 132 | 80 | 70 | 98 | 34 | LG+I+G4 |
| EOG092D4TT8 | 101 | 80 | 49 | 83 | 18 | LG+I+G4 |
| EOG092D01IY | 1782 | 81 | 919 | 1511 | 271 | JTT+I+G4 |
| EOG092D0FHW | 861 | 82 | 418 | 755 | 106 | LG+F+I+G4 |
| EOG092D0UNZ | 555 | 78 | 333 | 526 | 29 | LG+I+G4 |
| EOG092D4C5U | 386 | 82 | 111 | 361 | 25 | JTT+F+I+G4 |
| EOG092D16QM | 535 | 77 | 307 | 519 | 16 | LG+I+G4 |
| EOG092D0O61 | 1080 | 80 | 425 | 756 | 324 | JTT+F+I+G4 |

Table 7: Overview of alignments of genes used for phylogenomic reconstruction. Table generated with phylociraptor. *(continued)*

| gene | length | no. of sequences | no. of parsimony informative sites | no. of variable sites | no. of fixed sites | best model |
|---|---|---|---|---|---|---|
| EOG092D2W3S | 519 | 82 | 195 | 463 | 56 | JTT+I+G4 |
| EOG092D3LD3 | 364 | 83 | 167 | 328 | 36 | LG+F+I+G4 |
| EOG092D4DOH | 142 | 76 | 102 | 115 | 27 | LG+G4 |
| EOG092D465Q | 219 | 79 | 146 | 199 | 20 | LG+F+I+G4 |
| EOG092D0EUY | 954 | 82 | 638 | 894 | 60 | JTT+F+I+G4 |
| EOG092D02DP | 1584 | 82 | 918 | 1465 | 119 | JTT+F+I+G4 |
| EOG092D1V62 | 681 | 83 | 122 | 669 | 12 | JTT+I+G4 |
| EOG092D4POO | 205 | 82 | 70 | 193 | 12 | JTT+I+G4 |
| EOG092D47QN | 319 | 79 | 151 | 275 | 44 | JTTDCMut+I+G4 |
| EOG092D02YC | 1628 | 80 | 549 | 1482 | 146 | JTT+I+G4 |
| EOG092D1OV6 | 521 | 73 | 145 | 361 | 160 | JTT+I+G4 |
| EOG092D310Y | 376 | 81 | 214 | 339 | 37 | LG+I+G4 |
| EOG092D1US9 | 608 | 81 | 223 | 557 | 51 | LG+F+I+G4 |
| EOG092D2A51 | 576 | 81 | 265 | 563 | 13 | LG+I+G4 |
| EOG092D3Y3L | 216 | 82 | 58 | 97 | 119 | WAG+I+G4 |
| EOG092D37Y7 | 332 | 81 | 92 | 175 | 157 | JTT+I+G4 |
| EOG092D3RNW | 576 | 83 | 244 | 503 | 73 | JTT+I+G4 |
| EOG092D4M8A | 137 | 79 | 24 | 63 | 74 | LG+I+G4 |
| EOG092D4QXS | 116 | 79 | 53 | 100 | 16 | WAG+I+G4 |
| EOG092D3U3F | 235 | 81 | 137 | 189 | 46 | LG+I+G4 |
| EOG092D17TE | 546 | 82 | 250 | 346 | 200 | LG+I+G4 |
| EOG092D2QGC | 347 | 82 | 256 | 324 | 23 | LG+F+G4 |
| EOG092D0GSV | 818 | 79 | 481 | 731 | 87 | LG+I+G4 |
| EOG092D0MSY | 850 | 79 | 547 | 723 | 127 | LG+I+G4 |
| EOG092D1AQR | 791 | 82 | 297 | 679 | 112 | JTT+I+G4 |
| EOG092D4TKN | 130 | 80 | 73 | 110 | 20 | LG+I+G4 |
| EOG092D12VX | 775 | 82 | 302 | 742 | 33 | LG+F+I+G4 |
| EOG092D0NR0 | 878 | 80 | 336 | 640 | 238 | LG+F+I+G4 |
| EOG092D2DAM | 374 | 82 | 192 | 321 | 53 | LG+I+G4 |
| EOG092D1Q5G | 865 | 81 | 175 | 628 | 237 | JTT+F+I+G4 |
| EOG092D30CB | 327 | 67 | 170 | 316 | 11 | LG+I+G4 |
| EOG092D0YYX | 1133 | 82 | 434 | 968 | 165 | JTT+F+I+G4 |
| EOG092D4RXG | 98 | 81 | 68 | 86 | 12 | LG+G4 |
| EOG092D3W5S | 226 | 79 | 90 | 216 | 10 | LG+I+G4 |
| EOG092D3JLV | 251 | 82 | 110 | 171 | 80 | LG+I+G4 |
| EOG092D22H1 | 497 | 82 | 276 | 437 | 60 | LG+I+G4 |
| EOG092D2KKF | 1514 | 80 | 208 | 1434 | 80 | JTT+I+G4 |
| EOG092D0124 | 2350 | 82 | 1003 | 2274 | 76 | JTT+F+I+G4 |
| EOG092D4A7L | 257 | 67 | 128 | 249 | 8 | WAG+F+I+G4 |
| EOG092D1CLY | 634 | 82 | 212 | 485 | 149 | LG+I+G4 |
| EOG092D1ODU | 465 | 77 | 249 | 434 | 31 | LG+I+G4 |
| EOG092D2KNO | 378 | 81 | 149 | 238 | 140 | LG+I+G4 |
| EOG092D0V4E | 659 | 80 | 367 | 525 | 134 | LG+F+I+G4 |
| EOG092D2D7L | 402 | 79 | 154 | 247 | 155 | LG+I+G4 |
| EOG092D3ETT | 370 | 82 | 178 | 306 | 64 | LG+I+G4 |
| EOG092D3T9X | 948 | 81 | 329 | 909 | 39 | JTT+I+G4 |
| EOG092D2570 | 649 | 81 | 324 | 571 | 78 | JTT+I+G4 |
| EOG092D091W | 974 | 83 | 597 | 859 | 115 | LG+I+G4 |
| EOG092D3RDW | 655 | 83 | 74 | 574 | 81 | JTT+F+I+G4 |
| EOG092D4GXA | 152 | 79 | 42 | 71 | 81 | LG+G4 |
| EOG092D3WBP | 348 | 80 | 241 | 336 | 12 | LG+I+G4 |
| EOG092D1XYV | 454 | 82 | 180 | 355 | 99 | LG+I+G4 |
| EOG092D2MDL | 532 | 80 | 129 | 415 | 117 | JTT+F+I+G4 |
| EOG092D338V | 401 | 79 | 213 | 340 | 61 | JTT+I+G4 |
| EOG092D3YUS | 203 | 37 | 85 | 129 | 74 | LG+I+G4 |
| EOG092D0IC1 | 786 | 79 | 484 | 710 | 76 | JTT+I+G4 |
| EOG092D1EM9 | 473 | 81 | 244 | 332 | 141 | LG+I+G4 |

Table 7: Overview of alignments of genes used for phylogenomic reconstruction. Table generated with phylociraptor. *(continued)*

| gene | length | no. of sequences | no. of parsimony informative sites | no. of variable sites | no. of fixed sites | best model |
|---|---|---|---|---|---|---|
| EOG092D1I4F | 535 | 82 | 273 | 461 | 74 | LG+G4 |
| EOG092D4CM0 | 209 | 77 | 120 | 149 | 60 | Dayhoff+I+G4 |
| EOG092D31FN | 268 | 79 | 166 | 244 | 24 | LG+F+I+G4 |
| EOG092D0KAC | 1060 | 81 | 376 | 1052 | 8 | JTT+F+I+G4 |
| EOG092D30F8 | 282 | 83 | 107 | 163 | 119 | LG+I+G4 |
| EOG092D4BMV | 173 | 82 | 58 | 127 | 46 | JTT+I+G4 |
| EOG092D3SHS | 267 | 74 | 119 | 202 | 65 | LG+I+G4 |
| EOG092D0FN1 | 1107 | 81 | 400 | 1061 | 46 | JTT+F+I+G4 |
| EOG092D3EJG | 292 | 77 | 146 | 255 | 37 | LG+I+G4 |
| EOG092D0QOX | 968 | 80 | 270 | 959 | 9 | LG+F+I+G4 |
| EOG092D2633 | 445 | 81 | 299 | 368 | 77 | LG+I+G4 |
| EOG092D31Z3 | 394 | 79 | 160 | 376 | 18 | VT+I+G4 |
| EOG092D3CY0 | 387 | 65 | 172 | 351 | 36 | LG+F+I+G4 |
| EOG092D3LN2 | 271 | 78 | 198 | 254 | 17 | LG+I+G4 |
| EOG092D2XBR | 514 | 79 | 317 | 453 | 61 | LG+F+I+G4 |
| EOG092D2Q04 | 371 | 78 | 147 | 255 | 116 | LG+I+G4 |
| EOG092D1SCZ | 589 | 80 | 273 | 543 | 46 | JTTDCMut+I+G4 |
| EOG092D06QZ | 1206 | 82 | 598 | 1021 | 185 | LG+F+I+G4 |
| EOG092D2APH | 382 | 77 | 283 | 359 | 23 | LG+I+G4 |
| EOG092D1HB6 | 571 | 83 | 271 | 458 | 113 | LG+I+G4 |
| EOG092D1J6A | 811 | 82 | 146 | 686 | 125 | JTT+F+I+G4 |
| EOG092D0ZEU | 1165 | 82 | 472 | 1030 | 135 | JTT+F+I+G4 |
| EOG092D3039 | 936 | 82 | 358 | 826 | 110 | JTT+F+I+G4 |
| EOG092D2LGS | 411 | 83 | 154 | 393 | 18 | JTT+F+I+G4 |
| EOG092D29J0 | 721 | 81 | 323 | 620 | 101 | JTTDCMut+I+G4 |
| EOG092D2GP9 | 489 | 80 | 279 | 437 | 52 | LG+I+G4 |
| EOG092D2O0K | 481 | 80 | 174 | 464 | 17 | LG+F+I+G4 |
| EOG092D2OL5 | 404 | 78 | 238 | 384 | 20 | LG+I+G4 |
| EOG092D49PH | 357 | 80 | 113 | 343 | 14 | LG+I+G4 |
| EOG092D22TL | 428 | 82 | 189 | 279 | 149 | LG+I+G4 |
| EOG092D3MM0 | 542 | 82 | 197 | 474 | 68 | JTT+I+G4 |
| EOG092D45EU | 248 | 81 | 119 | 237 | 11 | JTTDCMut+I+G4 |
| EOG092D3CMX | 743 | 75 | 309 | 733 | 10 | JTT+F+I+G4 |
| EOG092D4824 | 233 | 32 | 112 | 185 | 48 | JTTDCMut+I+G4 |
| EOG092D0433 | 2462 | 82 | 984 | 2432 | 30 | JTT+F+I+G4 |
| EOG092D3VPE | 271 | 80 | 197 | 247 | 24 | JTT+I+G4 |
| EOG092D13K4 | 1270 | 83 | 692 | 1196 | 74 | JTT+F+I+G4 |
| EOG092D3Y0X | 347 | 82 | 250 | 318 | 29 | LG+F+I+G4 |
| EOG092D38IH | 307 | 79 | 138 | 205 | 102 | LG+I+G4 |
| EOG092D08HR | 1855 | 81 | 747 | 1674 | 181 | JTT+F+I+G4 |
| EOG092D3JAM | 248 | 77 | 122 | 201 | 47 | LG+F+I+G4 |
| EOG092D3DOE | 393 | 83 | 208 | 364 | 29 | LG+I+G4 |
| EOG092D3WW4 | 204 | 80 | 64 | 129 | 75 | LG+I+G4 |
| EOG092D28PE | 396 | 76 | 126 | 385 | 11 | LG+F+I+G4 |
| EOG092D2UJS | 446 | 79 | 267 | 407 | 39 | LG+F+I+G4 |
| EOG092D4M5U | 172 | 80 | 79 | 151 | 21 | LG+I+G4 |
| EOG092D23RQ | 361 | 81 | 202 | 295 | 66 | LG+I+G4 |
| EOG092D4KK3 | 163 | 83 | 99 | 149 | 14 | LG+F+I+G4 |
| EOG092D2KHQ | 409 | 82 | 203 | 378 | 31 | JTT+I+G4 |
| EOG092D3X7S | 358 | 81 | 125 | 341 | 17 | VT+I+G4 |
| EOG092D4LDU | 172 | 75 | 42 | 87 | 85 | LG+I+G4 |
| EOG092D339O | 425 | 81 | 163 | 396 | 29 | LG+F+I+G4 |
| EOG092D2G09 | 566 | 83 | 249 | 456 | 110 | JTTDCMut+I+G4 |
| EOG092D1A87 | 565 | 83 | 366 | 492 | 73 | LG+I+G4 |
| EOG092D2MM2 | 197 | 72 | 108 | 174 | 23 | LG+F+I+G4 |
| EOG092D3JGK | 294 | 80 | 193 | 261 | 33 | LG+F+I+G4 |
| EOG092D00HH | 2892 | 82 | 1089 | 2600 | 292 | LG+F+I+G4 |

Table 7: Overview of alignments of genes used for phylogenomic reconstruction. Table generated with phylociraptor. *(continued)*

| gene | length | no. of sequences | no. of parsimony informative sites | no. of variable sites | no. of fixed sites | best model |
|---|---|---|---|---|---|---|
| EOG092D1BU7 | 587 | 74 | 328 | 521 | 66 | LG+I+G4 |
| EOG092D0KTI | 965 | 82 | 480 | 891 | 74 | LG+I+G4 |
| EOG092D1EK2 | 462 | 82 | 238 | 395 | 67 | LG+I+G4 |
| EOG092D2GVZ | 636 | 78 | 260 | 607 | 29 | JTTDCMut+I+G4 |
| EOG092D33Q2 | 299 | 79 | 135 | 175 | 124 | LG+I+G4 |
| EOG092D09N2 | 982 | 82 | 361 | 629 | 353 | LG+I+G4 |
| EOG092D3ODN | 299 | 55 | 187 | 280 | 19 | JTTDCMut+I+G4 |
| EOG092D3A58 | 378 | 81 | 185 | 326 | 52 | LG+F+I+G4 |
| EOG092D0EMK | 917 | 82 | 408 | 859 | 58 | JTT+I+G4 |
| EOG092D2AME | 521 | 81 | 197 | 431 | 90 | LG+I+G4 |
| EOG092D3YIQ | 321 | 79 | 38 | 172 | 149 | JTT+F+I+G4 |
| EOG092D0PER | 1124 | 80 | 452 | 999 | 125 | JTTDCMut+F+I+G4 |
| EOG092D0DVR | 1017 | 79 | 577 | 845 | 172 | LG+I+G4 |
| EOG092D1N80 | 528 | 81 | 227 | 434 | 94 | LG+F+I+G4 |
| EOG092D3MIU | 260 | 79 | 151 | 186 | 74 | LG+I+G4 |
| EOG092D1I8J | 601 | 83 | 183 | 573 | 28 | JTT+I+G4 |
| EOG092D4CHT | 178 | 70 | 108 | 167 | 11 | LG+G4 |
| EOG092D3VJX | 247 | 77 | 165 | 218 | 29 | LG+I+G4 |
| EOG092D2HJ9 | 295 | 81 | 183 | 259 | 36 | LG+F+I+G4 |
| EOG092D0PZY | 656 | 76 | 438 | 536 | 120 | LG+I+G4 |
| EOG092D48Y7 | 130 | 71 | 90 | 126 | 4 | LG+G4 |
| EOG092D2ZKG | 214 | 74 | 145 | 203 | 11 | JTT+I+G4 |
| EOG092D4HHP | 135 | 42 | 97 | 129 | 6 | LG+F+G4 |
| EOG092D1XPE | 333 | 64 | 141 | 308 | 25 | LG+I+G4 |
| EOG092D3FYF | 277 | 79 | 136 | 248 | 29 | LG+I+G4 |
| EOG092D0J2Q | 941 | 80 | 272 | 878 | 63 | JTT+F+I+G4 |
| EOG092D0AVU | 898 | 82 | 369 | 821 | 77 | JTT+I+G4 |
| EOG092D4D32 | 252 | 72 | 120 | 250 | 2 | LG+I+G4 |
| EOG092D4HWC | 123 | 75 | 83 | 105 | 18 | LG+F+I+G4 |
| EOG092D27O3 | 464 | 82 | 300 | 428 | 36 | LG+I+G4 |
| EOG092D2LO5 | 351 | 78 | 258 | 320 | 31 | LG+I+G4 |
| EOG092D3N04 | 253 | 81 | 129 | 208 | 45 | JTT+I+G4 |
| EOG092D26WR | 404 | 74 | 287 | 388 | 16 | LG+F+I+G4 |
| EOG092D3R6Y | 230 | 75 | 153 | 210 | 20 | LG+I+G4 |
| EOG092D2SJQ | 585 | 81 | 29 | 486 | 99 | JTT+F+I+G4 |
| EOG092D1GZC | 525 | 83 | 281 | 391 | 134 | LG+I+G4 |
| EOG092D24W9 | 632 | 59 | 346 | 625 | 7 | LG+F+I+G4 |
| EOG092D005G | 4901 | 80 | 1238 | 4650 | 251 | JTT+F+I+G4 |
| EOG092D2VEY | 584 | 80 | 282 | 520 | 64 | LG+I+G4 |
| EOG092D4K3B | 170 | 83 | 60 | 83 | 87 | LG+G4 |
| EOG092D2PVH | 481 | 82 | 249 | 463 | 18 | LG+I+G4 |
| EOG092D33G3 | 430 | 83 | 263 | 391 | 39 | LG+I+G4 |
| EOG092D1OVL | 745 | 83 | 350 | 694 | 51 | JTT+I+G4 |
| EOG092D0PHK | 1139 | 75 | 578 | 885 | 254 | JTT+I+G4 |
| EOG092D3C2F | 276 | 78 | 159 | 212 | 64 | LG+I+G4 |
| EOG092D0CUX | 914 | 83 | 435 | 689 | 225 | LG+I+G4 |
| EOG092D4074 | 205 | 80 | 99 | 178 | 27 | LG+I+G4 |
| EOG092D1AFQ | 579 | 76 | 332 | 487 | 92 | JTT+I+G4 |
| EOG092D2WPT | 293 | 68 | 92 | 281 | 12 | LG+G4 |
| EOG092D0SK9 | 828 | 81 | 255 | 753 | 75 | JTT+I+G4 |
| EOG092D34D0 | 262 | 82 | 116 | 190 | 72 | LG+I+G4 |
| EOG092D12XY | 774 | 77 | 377 | 728 | 46 | LG+I+G4 |
| EOG092D0FQ1 | 784 | 78 | 460 | 665 | 119 | LG+I+G4 |
| EOG092D39KW | 725 | 78 | 360 | 688 | 37 | JTT+I+G4 |
| EOG092D1IEY | 499 | 80 | 122 | 193 | 306 | LG+I+G4 |
| EOG092D32X3 | 392 | 75 | 200 | 327 | 65 | LG+I+G4 |
| EOG092D3JKF | 249 | 60 | 134 | 200 | 49 | JTTDCMut+G4 |

Table 7: Overview of alignments of genes used for phylogenomic reconstruction. Table generated with phylociraptor. *(continued)*

| gene | length | no. of sequences | no. of parsimony informative sites | no. of variable sites | no. of fixed sites | best model |
|---|---|---|---|---|---|---|
| EOG092D0454 | 1995 | 81 | 489 | 1912 | 83 | LG+F+I+G4 |
| EOG092D3VHN | 208 | 81 | 135 | 176 | 32 | JTT+I+G4 |
| EOG092D3EJQ | 321 | 75 | 212 | 258 | 63 | LG+I+G4 |
| EOG092D2G5H | 327 | 80 | 140 | 202 | 125 | JTT+I+G4 |
| EOG092D0XC3 | 737 | 80 | 417 | 630 | 107 | JTT+I+G4 |
| EOG092D1BAU | 620 | 81 | 263 | 489 | 131 | JTT+F+I+G4 |
| EOG092D2N1K | 240 | 81 | 143 | 221 | 19 | LG+I+G4 |
| EOG092D068O | 1708 | 81 | 727 | 1424 | 284 | LG+F+I+G4 |
| EOG092D4S06 | 126 | 78 | 65 | 107 | 19 | LG+I+G4 |
| EOG092D1J1B | 456 | 43 | 251 | 443 | 13 | LG+I+G4 |
| EOG092D3WUX | 350 | 73 | 216 | 343 | 7 | LG+F+G4 |
| EOG092D3ZRA | 238 | 82 | 118 | 196 | 42 | WAG+I+G4 |
| EOG092D24YM | 390 | 76 | 279 | 376 | 14 | LG+F+I+G4 |
| EOG092D1W7D | 653 | 80 | 186 | 568 | 85 | JTT+I+G4 |
| EOG092D2I29 | 426 | 80 | 208 | 392 | 34 | JTT+I+G4 |
| EOG092D03LV | 1572 | 47 | 477 | 1538 | 34 | JTT+F+I+G4 |
| EOG092D4HNO | 135 | 78 | 76 | 110 | 25 | WAG+G4 |
| EOG092D1QZ9 | 475 | 80 | 153 | 233 | 242 | LG+I+G4 |
| EOG092D3LSA | 240 | 82 | 110 | 163 | 77 | LG+I+G4 |
| EOG092D4H5J | 296 | 82 | 62 | 281 | 15 | JTT+I+G4 |
| EOG092D2BNK | 601 | 83 | 194 | 488 | 113 | JTT+F+I+G4 |
| EOG092D0B5M | 906 | 82 | 600 | 738 | 168 | LG+I+G4 |
| EOG092D1JY2 | 614 | 79 | 374 | 534 | 80 | LG+F+I+G4 |
| EOG092D40N7 | 224 | 80 | 136 | 182 | 42 | LG+I+G4 |
| EOG092D1THK | 547 | 71 | 228 | 462 | 85 | LG+F+I+G4 |
| EOG092D2R1Y | 323 | 74 | 244 | 280 | 43 | LG+F+I+G4 |
| EOG092D1TFV | 574 | 81 | 373 | 513 | 61 | LG+I+G4 |
| EOG092D3H3Y | 287 | 73 | 155 | 260 | 27 | LG+I+G4 |
| EOG092D304F | 342 | 82 | 186 | 273 | 69 | LG+I+G4 |
| EOG092D0KCS | 1659 | 80 | 617 | 1467 | 192 | JTT+I+G4 |
| EOG092D3I9A | 466 | 81 | 353 | 419 | 47 | LG+F+I+G4 |
| EOG092D2QPK | 357 | 80 | 188 | 246 | 111 | LG+F+I+G4 |
| EOG092D3Z9F | 190 | 79 | 96 | 169 | 21 | LG+F+I+G4 |
| EOG092D1CJS | 558 | 78 | 191 | 295 | 263 | JTT+I+G4 |
| EOG092D2ASD | 441 | 80 | 190 | 275 | 166 | LG+I+G4 |
| EOG092D3NOC | 323 | 82 | 185 | 301 | 22 | LG+I+G4 |
| EOG092D0H7X | 1016 | 83 | 429 | 964 | 52 | LG+F+I+G4 |
| EOG092D0NJO | 823 | 77 | 309 | 740 | 83 | JTT+I+G4 |
| EOG092D0XLN | 768 | 80 | 453 | 641 | 127 | LG+F+I+G4 |
| EOG092D3ZKF | 215 | 81 | 122 | 173 | 42 | LG+I+G4 |
| EOG092D3O42 | 210 | 81 | 144 | 180 | 30 | JTT+I+G4 |
| EOG092D01MX | 1847 | 83 | 580 | 1608 | 239 | JTT+I+G4 |
| EOG092D26V2 | 471 | 81 | 175 | 458 | 13 | LG+I+G4 |
| EOG092D0WCK | 847 | 81 | 267 | 798 | 49 | JTT+I+G4 |
| EOG092D1DJB | 663 | 82 | 136 | 610 | 53 | JTT+F+I+G4 |
| EOG092D383R | 1301 | 82 | 310 | 1229 | 72 | JTT+I+G4 |
| EOG092D0YTO | 891 | 81 | 427 | 792 | 99 | JTT+F+I+G4 |
| EOG092D4MIB | 176 | 80 | 116 | 153 | 23 | LG+I+G4 |
| EOG092D1VPG | 433 | 80 | 137 | 235 | 198 | LG+I+G4 |
| EOG092D343G | 298 | 80 | 153 | 243 | 55 | LG+I+G4 |
| EOG092D0CQQ | 542 | 79 | 323 | 488 | 54 | LG+I+G4 |
| EOG092D4N4A | 137 | 81 | 28 | 75 | 62 | Dayhoff+I+G4 |
| EOG092D1O8L | 439 | 80 | 186 | 298 | 141 | LG+I+G4 |
| EOG092D3QEX | 268 | 77 | 158 | 262 | 6 | JTTDCMut+G4 |
| EOG092D36LY | 351 | 80 | 169 | 219 | 132 | LG+I+G4 |
| EOG092D0Y1F | 623 | 81 | 313 | 525 | 98 | JTTDCMut+I+G4 |
| EOG092D18RK | 901 | 82 | 292 | 818 | 83 | JTT+F+I+G4 |

Table 7: Overview of alignments of genes used for phylogenomic reconstruction. Table generated with phylociraptor. *(continued)*

| gene | length | no. of sequences | no. of parsimony informative sites | no. of variable sites | no. of fixed sites | best model |
|---|---|---|---|---|---|---|
| EOG092D38PJ | 370 | 79 | 199 | 336 | 34 | LG+I+G4 |
| EOG092D47ZH | 208 | 79 | 123 | 170 | 38 | LG+I+G4 |
| EOG092D3LLM | 381 | 80 | 122 | 344 | 37 | JTT+I+G4 |
| EOG092D04RH | 1303 | 78 | 434 | 1274 | 29 | LG+F+I+G4 |
| EOG092D29YU | 535 | 82 | 198 | 466 | 69 | LG+I+G4 |
| EOG092D0VF1 | 586 | 78 | 308 | 573 | 13 | LG+F+I+G4 |
| EOG092D1CXA | 643 | 73 | 366 | 575 | 68 | LG+I+G4 |
| EOG092D074K | 1152 | 83 | 553 | 1044 | 108 | JTT+F+I+G4 |
| EOG092D4RL0 | 120 | 81 | 70 | 101 | 19 | LG+I+G4 |
| EOG092D4597 | 168 | 78 | 120 | 158 | 10 | LG+F+I+G4 |
| EOG092D131D | 807 | 76 | 185 | 797 | 10 | JTT+F+I+G4 |
| EOG092D2UGI | 229 | 75 | 131 | 174 | 55 | LG+I+G4 |
| EOG092D16LD | 786 | 82 | 312 | 612 | 174 | LG+I+G4 |
| EOG092D1HN1 | 472 | 81 | 243 | 423 | 49 | JTT+I+G4 |
| EOG092D3MYV | 580 | 81 | 100 | 503 | 77 | JTT+F+I+G4 |
| EOG092D3BM8 | 253 | 73 | 180 | 239 | 14 | LG+I+G4 |
| EOG092D4A3U | 228 | 80 | 145 | 212 | 16 | LG+I+G4 |
| EOG092D383L | 498 | 75 | 255 | 485 | 13 | LG+I+G4 |
| EOG092D145A | 601 | 79 | 194 | 541 | 60 | JTT+F+I+G4 |
| EOG092D3J2L | 331 | 75 | 128 | 318 | 13 | WAG+I+G4 |
| EOG092D2U8Y | 395 | 80 | 211 | 348 | 47 | LG+I+G4 |
| EOG092D0O9O | 1054 | 81 | 297 | 980 | 74 | JTT+I+G4 |
| EOG092D2F9R | 449 | 81 | 256 | 409 | 40 | LG+I+G4 |
| EOG092D1VHG | 403 | 76 | 271 | 363 | 40 | LG+F+I+G4 |
| EOG092D2VBX | 378 | 80 | 166 | 311 | 67 | JTTDCMut+I+G4 |
| EOG092D22SO | 1022 | 78 | 231 | 1014 | 8 | JTT+F+I+G4 |
| EOG092D1BN5 | 640 | 78 | 288 | 577 | 63 | JTT+I+G4 |
| EOG092D425I | 230 | 74 | 151 | 201 | 29 | LG+I+G4 |
| EOG092D3X9I | 264 | 71 | 137 | 253 | 11 | LG+I+G4 |
| EOG092D2W25 | 498 | 76 | 222 | 456 | 42 | LG+F+I+G4 |
| EOG092D06DZ | 1216 | 78 | 648 | 1005 | 211 | JTT+I+G4 |
| EOG092D3PY2 | 312 | 82 | 210 | 284 | 28 | LG+F+G4 |
| EOG092D0BHJ | 797 | 82 | 317 | 627 | 170 | LG+I+G4 |
| EOG092D4Q0H | 107 | 81 | 59 | 74 | 33 | LG+I+G4 |
| EOG092D4E56 | 309 | 81 | 145 | 300 | 9 | LG+I+G4 |
| EOG092D0KU4 | 1077 | 83 | 503 | 1019 | 58 | LG+F+I+G4 |
| EOG092D4P6F | 177 | 81 | 120 | 169 | 8 | LG+F+I+G4 |
| EOG092D0VZC | 684 | 72 | 368 | 674 | 10 | JTT+I+G4 |
| EOG092D2Y9W | 351 | 73 | 228 | 344 | 7 | LG+I+G4 |
| EOG092D0XEW | 797 | 81 | 214 | 698 | 99 | JTT+F+I+G4 |
| EOG092D0PA5 | 839 | 78 | 430 | 630 | 209 | JTTDCMut+I+G4 |
| EOG092D1ABF | 647 | 82 | 338 | 596 | 51 | LG+I+G4 |
| EOG092D4E8N | 159 | 82 | 92 | 125 | 34 | JTT+G4 |
| EOG092D274B | 496 | 83 | 321 | 416 | 80 | JTT+I+G4 |
| EOG092D2UFS | 431 | 78 | 262 | 358 | 73 | JTTDCMut+I+G4 |
| EOG092D02JB | 1604 | 83 | 828 | 1297 | 307 | JTT+I+G4 |
| EOG092D10MG | 871 | 67 | 545 | 714 | 157 | LG+I+G4 |
| EOG092D2T3Z | 364 | 80 | 151 | 244 | 120 | LG+I+G4 |
| EOG092D2N6I | 309 | 82 | 177 | 256 | 53 | WAG+I+G4 |
| EOG092D1CGS | 547 | 75 | 257 | 473 | 74 | LG+I+G4 |
| EOG092D0A4I | 873 | 81 | 348 | 719 | 154 | JTTDCMut+I+G4 |
| EOG092D0ZAJ | 433 | 61 | 266 | 395 | 38 | LG+I+G4 |
| EOG092D37VX | 375 | 80 | 74 | 205 | 170 | DCMut+F+I+G4 |
| EOG092D1CVU | 1067 | 79 | 219 | 975 | 92 | JTT+F+I+G4 |
| EOG092D2OW6 | 346 | 72 | 200 | 271 | 75 | LG+I+G4 |
| EOG092D1MLK | 477 | 75 | 260 | 425 | 52 | LG+I+G4 |
| EOG092D2OSF | 351 | 72 | 217 | 309 | 42 | JTT+I+G4 |

Table 7: Overview of alignments of genes used for phylogenomic reconstruction. Table generated with phylociraptor. *(continued)*

| gene | length | no. of sequences | no. of parsimony informative sites | no. of variable sites | no. of fixed sites | best model |
|---|---|---|---|---|---|---|
| EOG092D02CG | 1414 | 81 | 822 | 1192 | 222 | JTT+F+I+G4 |
| EOG092D0PTK | 627 | 82 | 329 | 588 | 39 | JTT+I+G4 |
| EOG092D383C | 399 | 83 | 155 | 278 | 121 | JTT+F+I+G4 |
| EOG092D2VZF | 405 | 82 | 242 | 372 | 33 | LG+I+G4 |
| EOG092D15UU | 637 | 81 | 220 | 348 | 289 | LG+I+G4 |
| EOG092D3I4P | 303 | 80 | 157 | 265 | 38 | WAG+I+G4 |
| EOG092D3GR4 | 1518 | 81 | 394 | 1489 | 29 | JTT+F+I+G4 |
| EOG092D2AWS | 504 | 80 | 197 | 472 | 32 | LG+F+I+G4 |
| EOG092D27Y2 | 370 | 76 | 214 | 306 | 64 | JTT+I+G4 |
| EOG092D2SZZ | 415 | 68 | 268 | 396 | 19 | JTT+F+I+G4 |
| EOG092D0ACX | 1063 | 83 | 381 | 971 | 92 | LG+F+I+G4 |
| EOG092D4EIS | 276 | 78 | 65 | 244 | 32 | JTTDCMut+I+G4 |
| EOG092D3HE2 | 355 | 82 | 126 | 313 | 42 | JTTDCMut+I+G4 |
| EOG092D1RW7 | 677 | 80 | 354 | 638 | 39 | JTT+F+I+G4 |
| EOG092D4CXI | 310 | 77 | 155 | 300 | 10 | LG+F+I+G4 |
| EOG092D0B9X | 1022 | 82 | 423 | 652 | 370 | JTT+I+G4 |
| EOG092D49XW | 147 | 76 | 66 | 109 | 38 | JTTDCMut+I+G4 |
| EOG092D33N6 | 567 | 80 | 259 | 544 | 23 | JTT+I+G4 |
| EOG092D3QYI | 295 | 78 | 201 | 271 | 24 | LG+I+G4 |
| EOG092D3F2O | 291 | 78 | 175 | 235 | 56 | LG+I+G4 |
| EOG092D2X2E | 240 | 80 | 55 | 185 | 55 | LG+I+G4 |
| EOG092D0VTV | 617 | 79 | 294 | 473 | 144 | LG+I+G4 |
| EOG092D1VHP | 341 | 77 | 263 | 336 | 5 | LG+F+I+G4 |
| EOG092D3ZLJ | 522 | 82 | 210 | 469 | 53 | JTT+F+I+G4 |
| EOG092D3JAF | 285 | 72 | 131 | 268 | 17 | LG+F+I+G4 |
| EOG092D10OW | 646 | 80 | 383 | 626 | 20 | JTT+F+I+G4 |
| EOG092D3EQH | 536 | 81 | 205 | 495 | 41 | JTTDCMut+F+I+G4 |
| EOG092D0C9Q | 1150 | 78 | 435 | 981 | 169 | JTT+I+G4 |
| EOG092D0GP7 | 752 | 81 | 410 | 505 | 247 | LG+I+G4 |
| EOG092D19EQ | 535 | 78 | 165 | 229 | 306 | LG+I+G4 |
| EOG092D0XJC | 640 | 82 | 260 | 634 | 6 | JTT+F+I+G4 |
| EOG092D2MHF | 576 | 82 | 332 | 551 | 25 | JTT+I+G4 |
| EOG092D0KYR | 1499 | 80 | 732 | 1272 | 227 | LG+I+G4 |
| EOG092D29XW | 387 | 78 | 123 | 269 | 118 | LG+G4 |
| EOG092D3OQJ | 269 | 78 | 150 | 255 | 14 | JTT+I+G4 |
| EOG092D4C0N | 269 | 80 | 175 | 247 | 22 | JTT+I+G4 |
| EOG092D2P7K | 349 | 79 | 152 | 221 | 128 | LG+I+G4 |
| EOG092D4A4D | 364 | 31 | 120 | 352 | 12 | LG+F+I+G4 |
| EOG092D25DG | 438 | 80 | 114 | 215 | 223 | LG+I+G4 |
| EOG092D0NWF | 943 | 77 | 414 | 804 | 139 | LG+F+I+G4 |
| EOG092D3SKZ | 274 | 79 | 104 | 219 | 55 | LG+I+G4 |
| EOG092D0DLV | 840 | 80 | 272 | 830 | 10 | JTT+F+I+G4 |
| EOG092D2NES | 381 | 81 | 136 | 243 | 138 | LG+I+G4 |
| EOG092D45DK | 135 | 80 | 85 | 119 | 16 | LG+G4 |
| EOG092D4A5H | 168 | 82 | 84 | 104 | 64 | LG+I+G4 |
| EOG092D0HLF | 745 | 79 | 561 | 689 | 56 | LG+I+G4 |
| EOG092D3IIV | 486 | 77 | 177 | 464 | 22 | JTTDCMut+I+G4 |
| EOG092D1EU1 | 469 | 83 | 79 | 197 | 272 | JTT+I+G4 |
| EOG092D46UJ | 223 | 79 | 145 | 193 | 30 | LG+I+G4 |
| EOG092D4IK8 | 213 | 81 | 39 | 141 | 72 | LG+G4 |
| EOG092D0JYM | 937 | 83 | 295 | 925 | 12 | LG+F+I+G4 |
| EOG092D4PJO | 97 | 81 | 34 | 60 | 37 | LG+I+G4 |
| EOG092D33CF | 457 | 70 | 212 | 405 | 52 | JTT+F+I+G4 |
| EOG092D3KPJ | 274 | 83 | 69 | 180 | 94 | WAG+I+G4 |
| EOG092D2J85 | 506 | 81 | 187 | 355 | 151 | LG+I+G4 |
| EOG092D4NLC | 151 | 80 | 68 | 130 | 21 | LG+I+G4 |
| EOG092D0I6N | 904 | 81 | 409 | 705 | 199 | JTT+F+I+G4 |

Table 7: Overview of alignments of genes used for phylogenomic reconstruction. Table generated with phylociraptor. *(continued)*

| gene | length | no. of sequences | no. of parsimony informative sites | no. of variable sites | no. of fixed sites | best model |
|---|---|---|---|---|---|---|
| EOG092D0UJA | 681 | 80 | 318 | 506 | 175 | LG+F+I+G4 |
| EOG092D1Z45 | 653 | 81 | 313 | 595 | 58 | JTT+I+G4 |
| EOG092D2CBO | 209 | 71 | 146 | 191 | 18 | WAG+I+G4 |
| EOG092D0LTX | 1116 | 79 | 485 | 1046 | 70 | JTT+F+I+G4 |
| EOG092D4SWY | 79 | 76 | 48 | 63 | 16 | Dayhoff+I+G4 |
| EOG092D23QW | 451 | 80 | 131 | 189 | 262 | LG+I+G4 |
| EOG092D0CKP | 913 | 79 | 558 | 790 | 123 | LG+I+G4 |
| EOG092D0LTY | 1185 | 77 | 556 | 1087 | 98 | JTT+F+I+G4 |
| EOG092D0TDX | 930 | 61 | 348 | 905 | 25 | JTT+I+G4 |
| EOG092D1TP5 | 1004 | 80 | 416 | 853 | 151 | JTTDCMut+F+I+G4 |
| EOG092D0877 | 1314 | 81 | 497 | 1285 | 29 | JTT+F+I+G4 |
| EOG092D0HY6 | 676 | 64 | 412 | 654 | 22 | LG+I+G4 |
| EOG092D1CX4 | 753 | 83 | 360 | 721 | 32 | JTT+I+G4 |
| EOG092D26CK | 1531 | 72 | 146 | 1524 | 7 | JTT+F+I+G4 |
| EOG092D3D9K | 402 | 80 | 120 | 356 | 46 | Dayhoff+I+G4 |
| EOG092D4497 | 226 | 82 | 103 | 171 | 55 | LG+G4 |
| EOG092D0OG6 | 809 | 75 | 480 | 752 | 57 | LG+F+I+G4 |
| EOG092D3TRJ | 188 | 79 | 98 | 153 | 35 | LG+I+G4 |
| EOG092D4HQ1 | 126 | 73 | 92 | 114 | 12 | LG+I+G4 |
| EOG092D2D6B | 475 | 80 | 256 | 402 | 73 | LG+I+G4 |
| EOG092D2YQC | 317 | 80 | 226 | 280 | 37 | LG+I+G4 |
| EOG092D44UA | 238 | 80 | 117 | 221 | 17 | LG+I+G4 |
| EOG092D2K9N | 1191 | 82 | 270 | 1112 | 79 | JTT+F+I+G4 |
| EOG092D2EKA | 420 | 79 | 69 | 146 | 274 | JTTDCMut+I+G4 |
| EOG092D1HJN | 985 | 83 | 183 | 939 | 46 | JTT+F+I+G4 |
| EOG092D08DO | 901 | 81 | 507 | 788 | 113 | LG+I+G4 |
| EOG092D0Y58 | 912 | 81 | 417 | 790 | 122 | JTT+I+G4 |
| EOG092D2L06 | 321 | 79 | 166 | 302 | 19 | JTTDCMut+I+G4 |
| EOG092D3OYK | 331 | 81 | 160 | 304 | 27 | LG+F+I+G4 |
| EOG092D136Q | 834 | 80 | 401 | 731 | 103 | LG+I+G4 |
| EOG092D2IC5 | 335 | 77 | 177 | 238 | 97 | LG+I+G4 |
| EOG092D4FLZ | 168 | 80 | 108 | 141 | 27 | LG+I+G4 |
| EOG092D0HUI | 1002 | 80 | 650 | 812 | 190 | LG+I+G4 |
| EOG092D3ASG | 317 | 80 | 134 | 257 | 60 | LG+I+G4 |
| EOG092D26R7 | 396 | 77 | 150 | 318 | 78 | JTT+I+G4 |
| EOG092D0OJS | 859 | 82 | 360 | 809 | 50 | JTT+I+G4 |
| EOG092D27S3 | 393 | 82 | 149 | 216 | 177 | LG+I+G4 |
| EOG092D10XC | 600 | 70 | 252 | 518 | 82 | LG+I+G4 |
| EOG092D3ZQD | 381 | 82 | 228 | 324 | 57 | LG+I+G4 |
| EOG092D4CWI | 158 | 77 | 46 | 101 | 57 | LG+I+G4 |
| EOG092D4D5H | 233 | 80 | 84 | 154 | 79 | LG+G4 |
| EOG092D2DDI | 499 | 77 | 305 | 400 | 99 | LG+F+I+G4 |
| EOG092D0BYP | 1219 | 78 | 531 | 1000 | 219 | JTT+F+I+G4 |
| EOG092D2SPT | 538 | 80 | 264 | 475 | 63 | JTTDCMut+I+G4 |
| EOG092D1TTT | 446 | 83 | 148 | 242 | 204 | JTT+I+G4 |
| EOG092D1H11 | 318 | 81 | 130 | 249 | 69 | LG+F+I+G4 |
| EOG092D12Y0 | 1041 | 82 | 498 | 850 | 191 | LG+F+I+G4 |
| EOG092D0FJ0 | 967 | 80 | 397 | 876 | 91 | JTTDCMut+I+G4 |
| EOG092D19VF | 571 | 82 | 170 | 305 | 266 | LG+I+G4 |
| EOG092D4D41 | 189 | 78 | 94 | 168 | 21 | LG+I+G4 |
| EOG092D2NAB | 565 | 81 | 284 | 508 | 57 | LG+I+G4 |
| EOG092D4A0Q | 149 | 81 | 92 | 120 | 29 | LG+I+G4 |
| EOG092D34V3 | 615 | 72 | 353 | 604 | 11 | LG+F+I+G4 |
| EOG092D1UVP | 514 | 81 | 176 | 293 | 221 | LG+I+G4 |
| EOG092D2CO6 | 626 | 80 | 157 | 620 | 6 | JTTDCMut+I+G4 |
| EOG092D4AE2 | 184 | 80 | 97 | 179 | 5 | LG+I+G4 |
| EOG092D2TE4 | 421 | 75 | 176 | 394 | 27 | JTTDCMut+F+I+G4 |

Table 7: Overview of alignments of genes used for phylogenomic reconstruction. Table generated with phylociraptor. *(continued)*

| gene | length | no. of sequences | no. of parsimony informative sites | no. of variable sites | no. of fixed sites | best model |
|---|---|---|---|---|---|---|
| EOG092D2BH7 | 460 | 78 | 250 | 426 | 34 | JTT+F+I+G4 |
| EOG092D4CIY | 202 | 72 | 69 | 189 | 13 | JTT+I+G4 |
| EOG092D483D | 173 | 80 | 120 | 157 | 16 | LG+G4 |
| EOG092D1NX0 | 558 | 80 | 313 | 460 | 98 | LG+F+I+G4 |
| EOG092D1CL2 | 604 | 82 | 267 | 527 | 77 | JTT+F+I+G4 |
| EOG092D3KT6 | 484 | 80 | 49 | 363 | 121 | LG+F+I+G4 |
| EOG092D2C3H | 419 | 72 | 276 | 382 | 37 | LG+I+G4 |
| EOG092D1B2T | 736 | 78 | 289 | 732 | 4 | JTT+F+I+G4 |
| EOG092D1SC1 | 529 | 79 | 294 | 402 | 127 | LG+I+G4 |
| EOG092D0FF8 | 960 | 81 | 381 | 950 | 10 | JTT+I+G4 |
| EOG092D181M | 672 | 83 | 211 | 643 | 29 | JTT+I+G4 |
| EOG092D4IDA | 192 | 83 | 132 | 176 | 16 | LG+I+G4 |
| EOG092D24UI | 588 | 67 | 349 | 472 | 116 | LG+I+G4 |
| EOG092D3D5H | 388 | 78 | 264 | 355 | 33 | LG+F+I+G4 |
| EOG092D3UZB | 422 | 82 | 228 | 382 | 40 | WAG+I+G4 |
| EOG092D0JYR | 842 | 81 | 204 | 773 | 69 | JTT+F+I+G4 |
| EOG092D03RC | 1377 | 83 | 747 | 1180 | 197 | LG+F+I+G4 |
| EOG092D4AAG | 272 | 80 | 60 | 260 | 12 | JTTDCMut+F+I+G4 |
| EOG092D0MFC | 805 | 64 | 476 | 700 | 105 | LG+I+G4 |
| EOG092D01QP | 1902 | 82 | 757 | 1807 | 95 | JTT+F+I+G4 |
| EOG092D45SG | 244 | 81 | 142 | 223 | 21 | LG+I+G4 |
| EOG092D3MZ3 | 294 | 82 | 160 | 233 | 61 | LG+I+G4 |
| EOG092D3GX9 | 289 | 76 | 167 | 283 | 6 | LG+I+G4 |
| EOG092D0PK7 | 501 | 78 | 385 | 474 | 27 | LG+I+G4 |
| EOG092D2B4A | 487 | 80 | 236 | 363 | 124 | JTTDCMut+I+G4 |
| EOG092D1ZGV | 546 | 77 | 274 | 517 | 29 | LG+I+G4 |
| EOG092D41FH | 339 | 81 | 199 | 325 | 14 | LG+I+G4 |
| EOG092D3VL7 | 155 | 80 | 102 | 137 | 18 | LG+I+G4 |
| EOG092D01ZK | 1450 | 81 | 450 | 1440 | 10 | JTT+F+I+G4 |
| EOG092D02MN | 1712 | 82 | 691 | 1680 | 32 | LG+F+I+G4 |
| EOG092D2X9K | 1006 | 82 | 92 | 957 | 49 | JTT+F+I+G4 |
| EOG092D3F1T | 227 | 78 | 149 | 210 | 17 | LG+I+G4 |
| EOG092D44X0 | 137 | 75 | 42 | 122 | 15 | LG+F+I+G4 |
| EOG092D2WL3 | 566 | 80 | 271 | 548 | 18 | JTTDCMut+I+G4 |
| EOG092D06C2 | 1197 | 75 | 852 | 1164 | 33 | JTT+F+I+G4 |
| EOG092D3SWM | 202 | 60 | 52 | 126 | 76 | LG+G4 |
| EOG092D08R4 | 1072 | 82 | 511 | 909 | 163 | LG+I+G4 |
| EOG092D3IQI | 448 | 69 | 224 | 432 | 16 | LG+I+G4 |
| EOG092D31TX | 351 | 80 | 154 | 288 | 63 | JTT+F+I+G4 |
| EOG092D4PES | 122 | 67 | 84 | 109 | 13 | LG+I+G4 |
| EOG092D4877 | 161 | 76 | 111 | 139 | 22 | LG+I+G4 |
| EOG092D06BS | 1368 | 78 | 1060 | 1295 | 73 | JTTDCMut+I+G4 |
| EOG092D4HIU | 214 | 82 | 57 | 204 | 10 | LG+F+G4 |
| EOG092D1DJJ | 654 | 77 | 330 | 540 | 114 | LG+I+G4 |
| EOG092D409L | 222 | 82 | 114 | 179 | 43 | LG+I+G4 |
| EOG092D0BFU | 1093 | 83 | 661 | 885 | 208 | LG+F+I+G4 |
| EOG092D2ACV | 512 | 79 | 165 | 312 | 200 | JTT+I+G4 |
| EOG092D1C4I | 500 | 70 | 244 | 398 | 102 | LG+I+G4 |
| EOG092D1HSA | 603 | 83 | 229 | 536 | 67 | LG+F+I+G4 |
| EOG092D1BH6 | 553 | 78 | 179 | 349 | 204 | LG+I+G4 |
| EOG092D0AI2 | 976 | 83 | 503 | 836 | 140 | JTT+I+G4 |
| EOG092D12N2 | 608 | 80 | 392 | 527 | 81 | LG+I+G4 |
| EOG092D1SQB | 748 | 82 | 424 | 654 | 94 | JTT+I+G4 |
| EOG092D0HBI | 912 | 80 | 464 | 643 | 269 | JTTDCMut+F+I+G4 |
| EOG092D0Q8X | 996 | 81 | 451 | 883 | 113 | LG+F+I+G4 |
| EOG092D03NE | 1885 | 71 | 675 | 1870 | 15 | JTT+F+I+G4 |
| EOG092D3VOQ | 283 | 80 | 177 | 235 | 48 | Dayhoff+I+G4 |

Table 7: Overview of alignments of genes used for phylogenomic reconstruction. Table generated with phylociraptor. *(continued)*

| gene | length | no. of sequences | no. of parsimony informative sites | no. of variable sites | no. of fixed sites | best model |
|---|---|---|---|---|---|---|
| EOG092D0564 | 1331 | 82 | 977 | 1235 | 96 | LG+I+G4 |
| EOG092D48PX | 188 | 78 | 124 | 157 | 31 | LG+I+G4 |
| EOG092D3HV9 | 332 | 79 | 176 | 294 | 38 | LG+I+G4 |
| EOG092D2SV5 | 351 | 80 | 136 | 339 | 12 | JTT+F+I+G4 |
| EOG092D1RSV | 597 | 78 | 249 | 533 | 64 | LG+I+G4 |
| EOG092D35RA | 517 | 80 | 262 | 437 | 80 | LG+F+I+G4 |
| EOG092D182U | 1198 | 67 | 337 | 1169 | 29 | JTT+I+G4 |
| EOG092D3RVC | 202 | 81 | 54 | 171 | 31 | JTTDCMut+F+I+G4 |
| EOG092D34F6 | 810 | 82 | 363 | 702 | 108 | JTT+I+G4 |
| EOG092D36TR | 988 | 81 | 644 | 951 | 37 | JTT+I+G4 |
| EOG092D0951 | 964 | 81 | 432 | 738 | 226 | LG+I+G4 |
| EOG092D2IG4 | 453 | 80 | 268 | 447 | 6 | LG+I+G4 |
| EOG092D2V2V | 451 | 72 | 257 | 398 | 53 | LG+F+I+G4 |
| EOG092D34RI | 286 | 82 | 154 | 240 | 46 | LG+I+G4 |
| EOG092D3ZTV | 267 | 82 | 165 | 248 | 19 | LG+I+G4 |
| EOG092D3TZJ | 413 | 83 | 129 | 344 | 69 | LG+F+I+G4 |
| EOG092D0QW3 | 1090 | 57 | 544 | 1004 | 86 | LG+F+I+G4 |
| EOG092D45Q9 | 165 | 21 | 76 | 118 | 47 | LG+G4 |
| EOG092D3SSR | 222 | 72 | 153 | 194 | 28 | LG+I+G4 |
| EOG092D0BA6 | 1354 | 82 | 462 | 1185 | 169 | JTT+I+G4 |
| EOG092D4U2S | 119 | 82 | 64 | 103 | 16 | LG+I+G4 |
| EOG092D3QDK | 251 | 79 | 119 | 211 | 40 | LG+I+G4 |
| EOG092D1HYF | 638 | 80 | 225 | 578 | 60 | LG+I+G4 |
| EOG092D3MYJ | 263 | 80 | 45 | 78 | 185 | DCMut+F+I+G4 |
| EOG092D4KGF | 298 | 77 | 182 | 287 | 11 | LG+I+G4 |
| EOG092D46GG | 159 | 79 | 77 | 130 | 29 | LG+I+G4 |
| EOG092D2OSE | 392 | 81 | 182 | 298 | 94 | LG+I+G4 |
| EOG092D0S7U | 745 | 82 | 177 | 699 | 46 | LG+I+G4 |
| EOG092D04FZ | 1786 | 72 | 613 | 1779 | 7 | JTT+F+I+G4 |
| EOG092D2J1E | 889 | 80 | 329 | 830 | 59 | JTT+I+G4 |
| EOG092D4JJW | 116 | 81 | 48 | 75 | 41 | LG+G4 |
| EOG092D4M39 | 166 | 70 | 49 | 143 | 23 | JTT+I+G4 |
| EOG092D1LUK | 660 | 81 | 273 | 453 | 207 | JTTDCMut+I+G4 |
| EOG092D3IT4 | 376 | 82 | 203 | 355 | 21 | JTTDCMut+I+G4 |
| EOG092D4LPB | 194 | 74 | 63 | 171 | 23 | JTTDCMut+I+G4 |
| EOG092D4K5O | 116 | 78 | 49 | 74 | 42 | LG+G4 |
| EOG092D0RY3 | 1097 | 81 | 511 | 1050 | 47 | JTT+F+I+G4 |
| EOG092D4199 | 184 | 76 | 108 | 153 | 31 | LG+G4 |
| EOG092D3X0O | 299 | 80 | 154 | 269 | 30 | LG+I+G4 |
| EOG092D0CTT | 1571 | 83 | 665 | 1523 | 48 | JTT+F+I+G4 |
| EOG092D0R2T | 728 | 80 | 428 | 636 | 92 | LG+I+G4 |
| EOG092D3OJ2 | 255 | 77 | 44 | 95 | 160 | JTTDCMut+I+G4 |
| EOG092D3NPG | 250 | 79 | 55 | 98 | 152 | LG+I+G4 |
| EOG092D0R62 | 870 | 83 | 256 | 813 | 57 | JTTDCMut+F+I+G4 |
| EOG092D05PP | 1173 | 82 | 473 | 1114 | 59 | JTT+I+G4 |
| EOG092D3RKB | 283 | 81 | 118 | 239 | 44 | LG+I+G4 |
| EOG092D07QT | 1127 | 82 | 318 | 671 | 456 | LG+I+G4 |
| EOG092D042R | 1298 | 80 | 857 | 1279 | 19 | JTT+F+I+G4 |
| EOG092D3VNW | 197 | 75 | 127 | 171 | 26 | LG+G4 |
| EOG092D4DJ2 | 283 | 82 | 98 | 275 | 8 | JTT+I+G4 |
| EOG092D16QX | 624 | 72 | 299 | 556 | 68 | LG+I+G4 |
| EOG092D1ZWB | 456 | 74 | 188 | 388 | 68 | LG+F+I+G4 |
| EOG092D1UOB | 562 | 81 | 279 | 449 | 113 | JTTDCMut+I+G4 |
| EOG092D1I2I | 399 | 81 | 207 | 389 | 10 | LG+I+G4 |
| EOG092D2E2U | 599 | 81 | 218 | 514 | 85 | JTT+F+I+G4 |
| EOG092D13SA | 654 | 83 | 276 | 509 | 145 | JTTDCMut+I+G4 |
| EOG092D47YD | 241 | 79 | 190 | 221 | 20 | JTTDCMut+I+G4 |

Table 7: Overview of alignments of genes used for phylogenomic reconstruction. Table generated with phylociraptor. *(continued)*

| gene | length | no. of sequences | no. of parsimony informative sites | no. of variable sites | no. of fixed sites | best model |
|---|---|---|---|---|---|---|
| EOG092D22TO | 483 | 81 | 229 | 461 | 22 | LG+F+I+G4 |
| EOG092D37RT | 171 | 81 | 112 | 143 | 28 | LG+I+G4 |
| EOG092D13G3 | 599 | 78 | 372 | 516 | 83 | LG+I+G4 |
| EOG092D3KBF | 247 | 75 | 125 | 183 | 64 | LG+I+G4 |
| EOG092D167J | 503 | 79 | 271 | 420 | 83 | JTT+I+G4 |
| EOG092D2W4H | 374 | 81 | 169 | 240 | 134 | LG+I+G4 |
| EOG092D4DWW | 251 | 55 | 110 | 246 | 5 | JTT+I+G4 |
| EOG092D0EC9 | 1473 | 81 | 650 | 1288 | 185 | LG+I+G4 |
| EOG092D4GNM | 191 | 74 | 105 | 187 | 4 | LG+I+G4 |
| EOG092D2SCW | 351 | 81 | 190 | 305 | 46 | LG+I+G4 |
| EOG092D3HQ3 | 280 | 81 | 108 | 244 | 36 | LG+F+I+G4 |
| EOG092D2HE0 | 492 | 79 | 203 | 415 | 77 | LG+I+G4 |
| EOG092D4HZE | 122 | 79 | 79 | 100 | 22 | LG+I+G4 |
| EOG092D3NQF | 420 | 62 | 190 | 404 | 16 | LG+F+I+G4 |
| EOG092D18W8 | 572 | 75 | 304 | 512 | 60 | LG+F+I+G4 |
| EOG092D1ML3 | 560 | 80 | 219 | 533 | 27 | LG+F+G4 |
| EOG092D4AOP | 216 | 80 | 130 | 195 | 21 | JTTDCMut+I+G4 |
| EOG092D3942 | 716 | 78 | 379 | 675 | 41 | JTT+F+I+G4 |
| EOG092D1V0I | 540 | 81 | 209 | 323 | 217 | LG+I+G4 |
| EOG092D2RBH | 347 | 82 | 222 | 298 | 49 | LG+I+G4 |
| EOG092D21TB | 520 | 80 | 292 | 449 | 71 | LG+I+G4 |
| EOG092D4GQ5 | 240 | 82 | 145 | 210 | 30 | LG+I+G4 |
| EOG092D42PJ | 220 | 81 | 139 | 177 | 43 | LG+I+G4 |
| EOG092D3LA0 | 295 | 82 | 185 | 280 | 15 | LG+I+G4 |
| EOG092D092M | 1070 | 82 | 491 | 726 | 344 | LG+I+G4 |
| EOG092D4679 | 176 | 81 | 119 | 165 | 11 | JTT+I+G4 |
| EOG092D20HF | 604 | 79 | 391 | 538 | 66 | JTTDCMut+I+G4 |
| EOG092D2Z0K | 613 | 80 | 246 | 571 | 42 | JTT+F+I+G4 |
| EOG092D0KIC | 997 | 81 | 539 | 766 | 231 | JTTDCMut+I+G4 |
| EOG092D3QQQ | 1606 | 83 | 269 | 1554 | 52 | JTT+F+I+G4 |
| EOG092D0GS4 | 1026 | 80 | 426 | 969 | 57 | JTT+F+I+G4 |
| EOG092D2502 | 428 | 77 | 308 | 383 | 45 | LG+I+G4 |
| EOG092D3GZM | 200 | 81 | 107 | 161 | 39 | LG+G4 |
| EOG092D0OYE | 998 | 82 | 457 | 909 | 89 | JTT+I+G4 |
| EOG092D0HR5 | 670 | 77 | 373 | 642 | 28 | JTT+F+I+G4 |
| EOG092D2A2K | 908 | 81 | 275 | 798 | 110 | LG+F+I+G4 |
| EOG092D28W9 | 969 | 79 | 477 | 914 | 55 | JTT+F+I+G4 |
| EOG092D1M7G | 760 | 81 | 394 | 677 | 83 | JTTDCMut+I+G4 |
| EOG092D06WH | 521 | 82 | 273 | 465 | 56 | LG+I+G4 |
| EOG092D326X | 589 | 82 | 116 | 583 | 6 | JTT+I+G4 |
| EOG092D3F3H | 720 | 80 | 148 | 702 | 18 | JTT+F+I+G4 |
| EOG092D3TIF | 310 | 81 | 139 | 231 | 79 | LG+F+I+G4 |
| EOG092D4JCC | 151 | 59 | 71 | 133 | 18 | LG+G4 |
| EOG092D1OF2 | 444 | 81 | 293 | 413 | 31 | LG+F+I+G4 |
| EOG092D14TO | 846 | 73 | 203 | 812 | 34 | JTT+F+I+G4 |
| EOG092D1LYU | 549 | 79 | 349 | 488 | 61 | LG+I+G4 |
| EOG092D3SXY | 297 | 82 | 150 | 283 | 14 | LG+F+I+G4 |
| EOG092D2Y6K | 317 | 78 | 172 | 225 | 92 | LG+I+G4 |
| EOG092D25D9 | 378 | 72 | 248 | 352 | 26 | LG+I+G4 |
| EOG092D3CIT | 216 | 74 | 133 | 193 | 23 | WAG+I+G4 |
| EOG092D0SSQ | 1627 | 81 | 261 | 1378 | 249 | JTTDCMut+I+G4 |
| EOG092D1BBT | 795 | 82 | 250 | 738 | 57 | JTT+I+G4 |
| EOG092D3IQU | 247 | 81 | 86 | 164 | 83 | LG+F+I+G4 |
| EOG092D0R2H | 819 | 82 | 328 | 770 | 49 | JTT+F+I+G4 |
| EOG092D1L93 | 494 | 80 | 259 | 407 | 87 | LG+I+G4 |
| EOG092D2BBJ | 531 | 81 | 245 | 461 | 70 | JTTDCMut+I+G4 |
| EOG092D3076 | 349 | 79 | 197 | 238 | 111 | LG+I+G4 |

Table 7: Overview of alignments of genes used for phylogenomic reconstruction. Table generated with phylociraptor. *(continued)*

| gene | length | no. of sequences | no. of parsimony informative sites | no. of variable sites | no. of fixed sites | best model |
|---|---|---|---|---|---|---|
| EOG092D26XB | 394 | 72 | 216 | 369 | 25 | JTTDCMut+I+G4 |
| EOG092D2RFS | 282 | 75 | 214 | 261 | 21 | LG+I+G4 |

#### Table 8: Gene family expansion analysis summary for different runs

Table 8: Table summarizing singinficantly expanded CAZyme families in different CAFE5 runs.

| run | rate | error_model | expanded_families |
|---|---|---|---|
| 1 | single_rate | | GH43:p=0.004 AA3:p=0 CE10:p=0 AA7:p=0 |
| 2 | single_rate | yes | GH43:p=0 AA9:p=0.014 AA3:p=0 CE10:p=0 AA7:p=0 |
| 3 | single_rate | | GH43:p=0.005 AA3:p=0 CE10:p=0 AA7:p=0 |
| 4 | single_rate | yes | GH43:p=0 AA9:p=0.016 AA3:p=0 CE10:p=0 AA7:p=0 |
| 5 | single_rate | | GH43:p=0.006 AA3:p=0 CE10:p=0 AA7:p=0 |
| 6 | single_rate | yes | GH43:p=0.001 AA9:p=0.018 AA3:p=0 CE10:p=0 AA7:p=0 |
| 7 | single_rate | | GH43:p=0.008 AA3:p=0 CE10:p=0 AA7:p=0 |
| 8 | single_rate | yes | GH43:p=0 AA9:p=0.013 AA3:p=0 CE10:p=0 AA7:p=0 |
| 9 | single_rate | | GH43:p=0.006 AA3:p=0 CE10:p=0 AA7:p=0 |
| 10 | single_rate | yes | GH43:p=0 AA9:p=0.023 AA3:p=0 CE10:p=0 AA7:p=0 |
| 11 | two_rates | | GH43:p=0.005 AA3:p=0 CE10:p=0 AA7:p=0 |
| 12 | two_rates | yes | GH18:p=0.039 GH43:p=0.001 AA9:p=0.021 AA3:p=0 CE10:p=0 AA7:p=0 |
| 13 | two_rates | | GH43:p=0.006 AA3:p=0 CE10:p=0 AA7:p=0 |
| 14 | two_rates | yes | GH18:p=0.041 GH43:p=0 AA9:p=0.019 AA3:p=0 CE10:p=0 AA7:p=0 |
| 15 | two_rates | | GH43:p=0.006 AA3:p=0 CE10:p=0 AA7:p=0 |
| 16 | two_rates | yes | GH43:p=0.001 AA9:p=0.024 AA3:p=0 CE10:p=0 AA7:p=0 |
| 17 | two_rates | | GH43:p=0.007 AA3:p=0 CE10:p=0 AA7:p=0 |
| 18 | two_rates | yes | GH18:p=0.045 GH43:p=0.001 AA9:p=0.027 AA3:p=0 CE10:p=0 AA7:p=0 |
| 19 | two_rates | | GH43:p=0.006 AA3:p=0 CE10:p=0 AA7:p=0 |
| 20 | two_rates | yes | GH18:p=0.047 GH43:p=0.001 AA9:p=0.021 AA3:p=0 CE10:p=0 AA7:p=0 |

#### Table 9, 10 and 11: Comparison of CAZyme number of LFS groups and other fungi

Table 9: This table shows mean total number of genes of different groups of genomes in different CAZyme classes and the difference in percent between LFS groups and other Ascomycete fungi included in this study.

| CAZyme | Lecanoromycetes (mean) | Lecanoromycetidae (mean) | Ostropomycetidae (mean) | Ostropomycetidae w/o OG clade (mean) | OG clade (mean) | other fungi (mean) | diff. Lecanoromycetes vs. other fungi (%) | diff. Lecanoromycetidae vs. other fungi (%) | diff. Ostropomycetidae vs. other fungi (%) | diff. Ostropomycetidae (w/o OG clade) vs. other fungi (%) | diff. OG clade vs. other fungi (%) | diff. Lecanoromycetidae vs. Ostropomycetidae (%) |
|---|---|---|---|---|---|---|---|---|---|---|---|---|
| AA | 40.739 | 41.045 | 40.458 | 34.421 | 63.400 | 63.297 | 43.366 | 42.651 | 44.025 | 59.101 | 0.162 | 1.441 |
| CBM | 4.457 | 4.318 | 4.583 | 3.789 | 7.600 | 9.568 | 72.890 | 75.608 | 70.444 | 86.518 | 22.922 | 5.957 |
| CE | 24.391 | 19.773 | 28.625 | 22.947 | 50.200 | 43.405 | 56.092 | 74.813 | 41.039 | 61.664 | 14.518 | 36.581 |
| GH | 100.326 | 84.318 | 115.000 | 98.737 | 176.800 | 169.162 | 51.087 | 66.943 | 38.121 | 52.576 | 4.415 | 30.787 |
| GT | 64.196 | 64.182 | 64.208 | 64.684 | 62.400 | 67.243 | 4.637 | 4.659 | 4.618 | 3.879 | 7.472 | 0.041 |
| PL | 0.457 | 0.091 | 0.792 | 0.105 | 3.400 | 6.784 | 174.779 | 194.711 | 158.198 | 193.888 | 66.454 | 158.798 |

Table 10: This table shows mean total number of genes of different groups of genomes in different CAZyme classes. The values of groups of LFS from Lecanoromycetes are compared to all other fungi and p-values are calculated with Wilcoxon rank-sum tests.

| CAZyme | Lecanoromycetes (mean) | Lecanoromycetidae (mean) | Ostropomycetidae (mean) | Ostropomycetidae w/o OG clade (mean) | OG clade (mean) | other fungi (mean) | p-value Lecanoromycetes vs. other fungi | p-value Lecanoromycetidae vs. other fungi | p-value Ostropomycetidae vs. other fungi | p-value Ostropomycetidae (w/o OG clade) vs. other fungi | p-value OG clade vs. other fungi | p-value Lecanoromycetidae vs. Ostropomycetidae |
|---|---|---|---|---|---|---|---|---|---|---|---|---|
| AA | 40.739 | 41.045 | 40.458 | 34.421 | 63.400 | 63.297 | 0.004 | 0.026 | 0.009 | 0.001 | 0.741 | 0.567 |
| CBM | 4.457 | 4.318 | 4.583 | 3.789 | 7.600 | 9.568 | 0.000 | 0.002 | 0.005 | 0.000 | 0.572 | 0.982 |
| CE | 24.391 | 19.773 | 28.625 | 22.947 | 50.200 | 43.405 | 0.000 | 0.000 | 0.011 | 0.001 | 0.351 | 0.024 |
| GH | 100.326 | 84.318 | 115.000 | 98.737 | 176.800 | 169.162 | 0.000 | 0.000 | 0.005 | 0.000 | 0.614 | 0.000 |
| GT | 64.196 | 64.182 | 64.208 | 64.684 | 62.400 | 67.243 | 0.180 | 0.265 | 0.270 | 0.435 | 0.259 | 0.956 |
| PL | 0.457 | 0.091 | 0.792 | 0.105 | 3.400 | 6.784 | 0.000 | 0.000 | 0.000 | 0.000 | 0.495 | 0.130 |

Table 11: This table show by how much percent CAZyme numbers are increased (positive values) or decreased (negative values) when comparing LFS groups with other fungi.

| CAZyme | Lecanoromycetes (mean) | Lecanoromycetidae (mean) | Ostropomycetidae (mean) | Ostropomycetidae w/o OG clade (mean) | OG clade (mean) | other fungi (mean) | incr. Lecanoromycetes vs. other fungi (%) | incr. Lecanoromycetidae vs. other fungi (%) | incr. Ostropomycetidae vs. other fungi (%) | incr. Ostropomycetidae (w/o OG clade) vs. other fungi (%) | incr. OG clade vs. other fungi (%) | incr. Lecanoromycetidae vs. Ostropomycetidae (%) |
|---|---|---|---|---|---|---|---|---|---|---|---|---|
| AA | 40.739 | 41.045 | 40.458 | 34.421 | 63.400 | 63.297 | -35.638 | -35.154 | -36.082 | -45.620 | 0.162 | -1.430 |
| CBM | 4.457 | 4.318 | 4.583 | 3.789 | 7.600 | 9.568 | -53.421 | -54.866 | -52.095 | -60.393 | -20.565 | 6.140 |
| CE | 24.391 | 19.773 | 28.625 | 22.947 | 50.200 | 43.405 | -43.806 | -54.446 | -34.052 | -47.132 | 15.654 | 44.770 |
| GH | 100.326 | 84.318 | 115.000 | 98.737 | 176.800 | 169.162 | -40.692 | -50.155 | -32.018 | -41.632 | 4.515 | 36.388 |
| GT | 64.196 | 64.182 | 64.208 | 64.684 | 62.400 | 67.243 | -4.532 | -4.553 | -4.513 | -3.806 | -7.203 | 0.041 |
| PL | 0.457 | 0.091 | 0.792 | 0.105 | 3.400 | 6.784 | -93.270 | -98.660 | -88.330 | -98.448 | -49.880 | 770.833 |

### Figure 1: Metrics of studied genomes

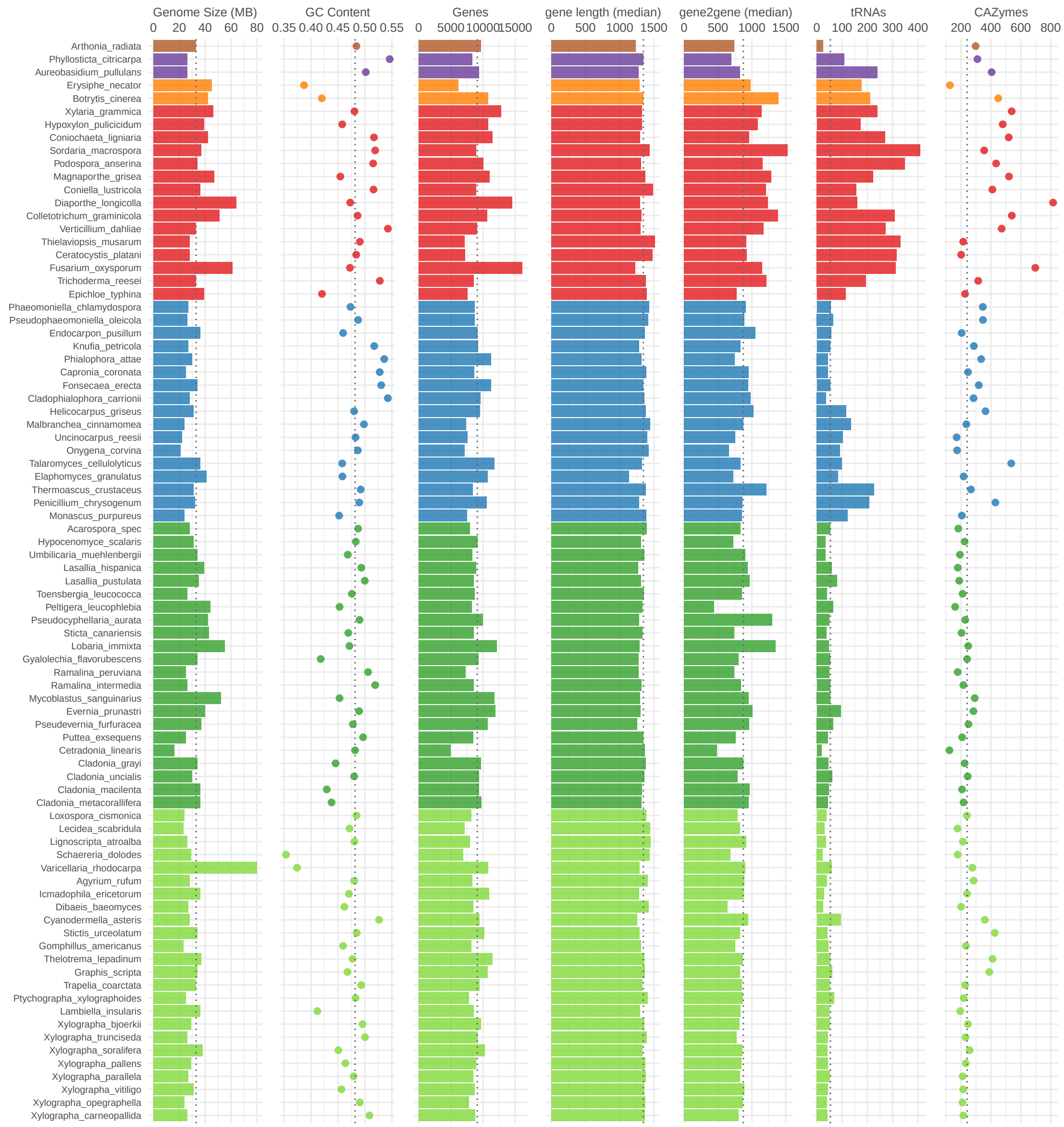

Figure 1: Different characteristics of the 83 fungal genomes studied here. Colors refer to different taxonomic groups. Brown - Arthoniomycetes, purple - Dothideomycetes, orange - Leotiomycetes, red - Sordariomycetes, blue - Eurotiomycetes, dark green - Lecanoromycetidae, light green - Ostropomycetidae

### Figure 2: BUSCO completeness of studied genomes

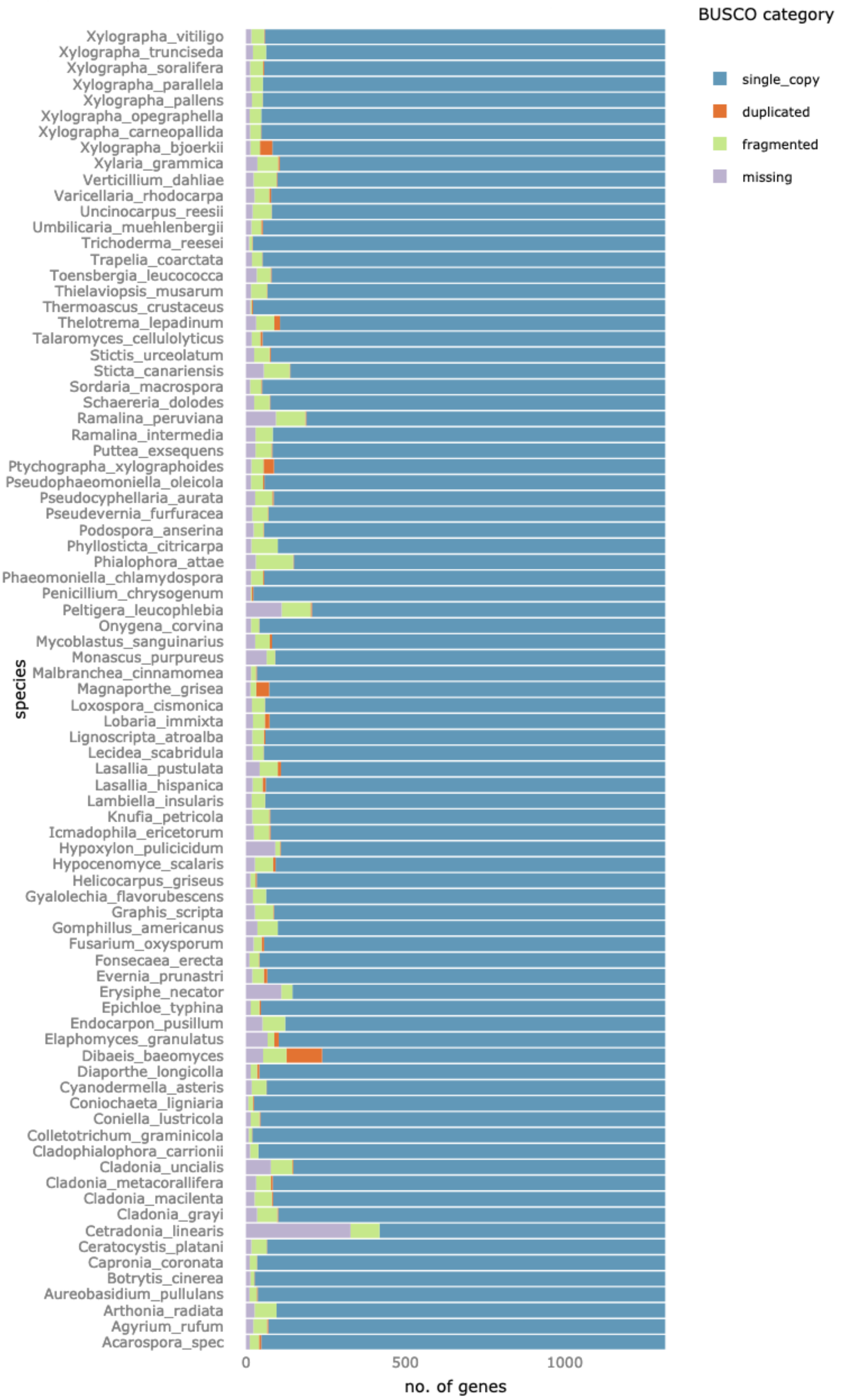

Figure 2: BUSCO completeness of core Ascomycota genes (ascomycota_odb9) in the 83 studied genomes. Figure generated with phylociraptor.

### Figure 3 and 4: Phylogenomic trees

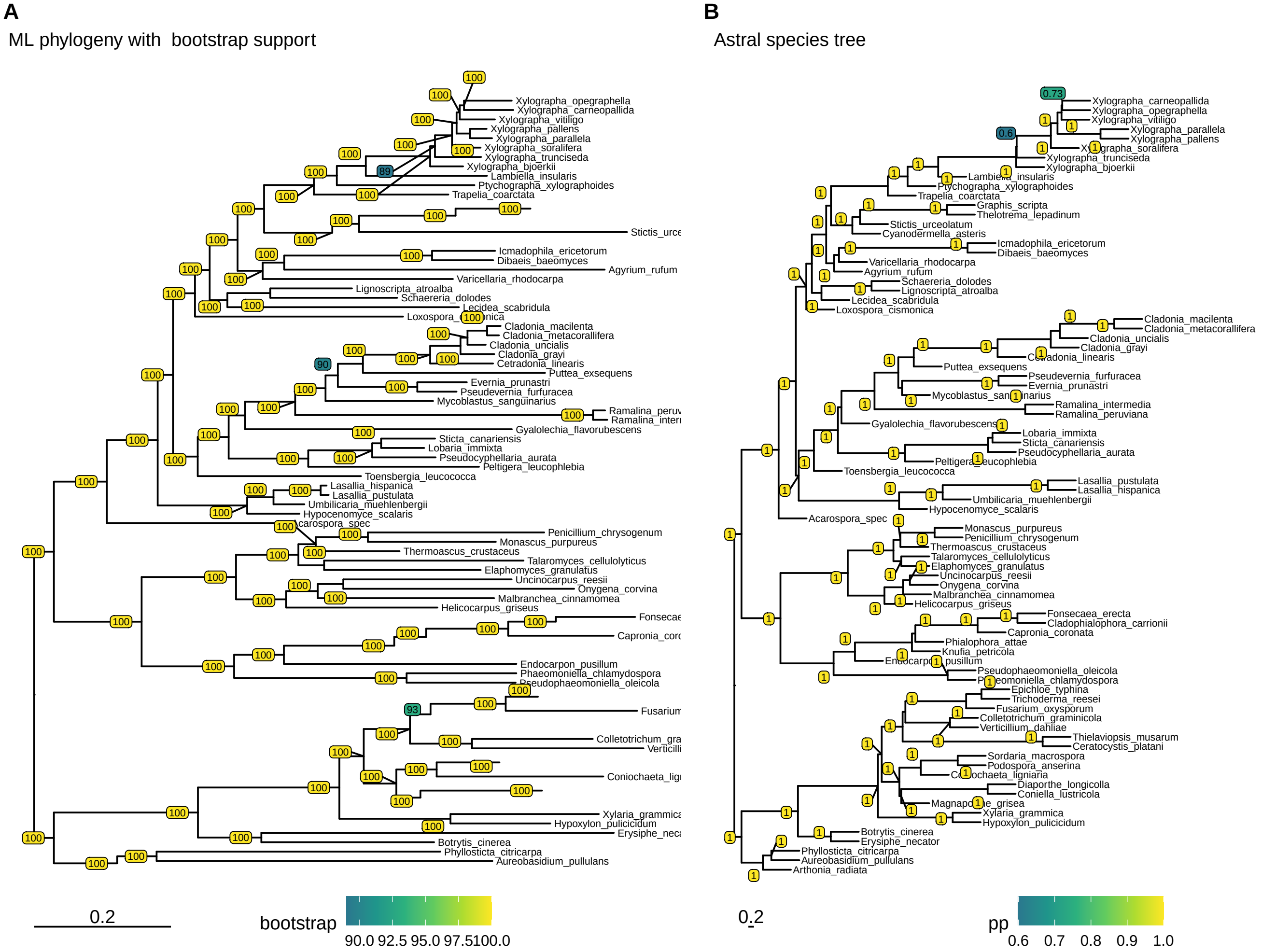

Figure 3: Phylogenomic trees of higher Ascomycetes based on 1310 alignments of single-copy genes. A) IQ-Tree maximum-likelihood tree of a concatenated supermatrix alignment with bootstrap node-support values. B) ASTRAL species tree calculated from individual maximum-likelihood gene-trees. Node support given as Posterior Probabilities (PP).

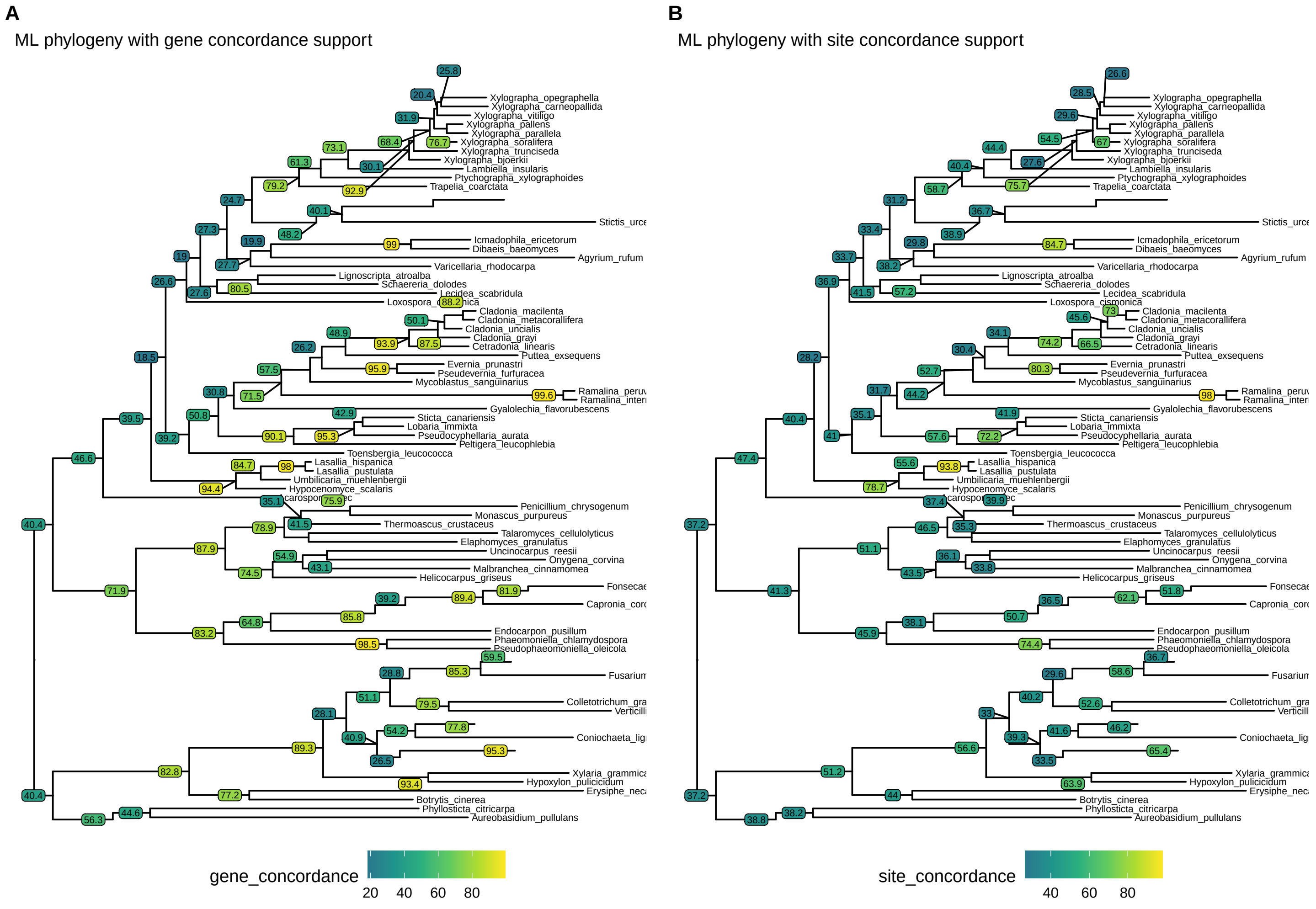

Figure 4: Maximum-Likelihood phylogenomic tree based in a concatenated alignment of 1310 singl-copy gene alignments. Node-support given as gene-concordance and site-concordance factors calculated with IQ-Tree.

#### Figure 5: Distribution of sugar-transporter orthologues

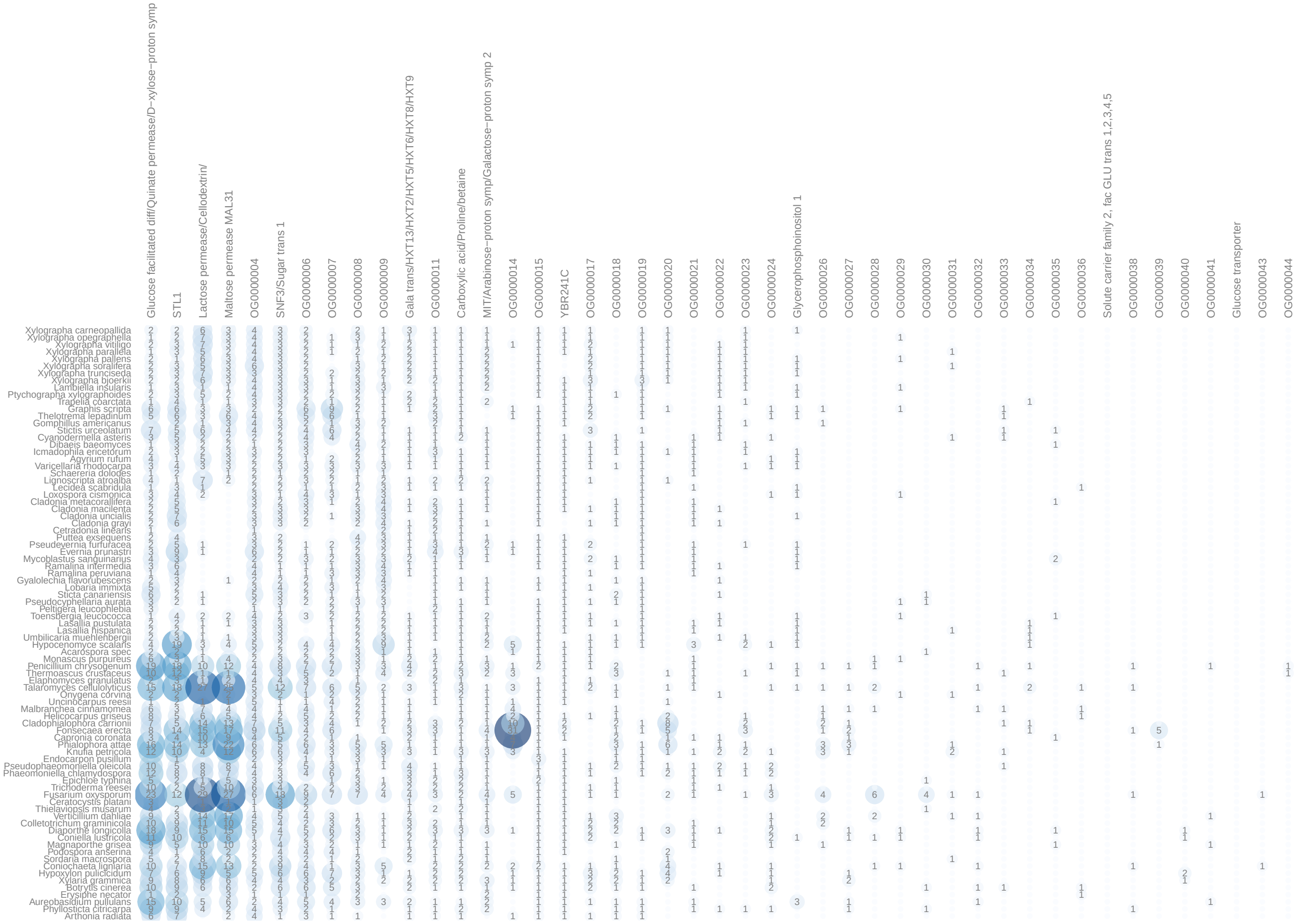

Figure 5: Overview of sugar transporters with PF00083 annotations in 83 studied genomes. Each column represents one orthogroup inferred by Orthofinder. Names of columns were assigned based on the presence of characterized transporter sequences in the respective orthogroup. Columns without any characterized transporter sequences still have names assigned by Orthofinder.

### Figure 6: Distribution of peroxidase orthologues

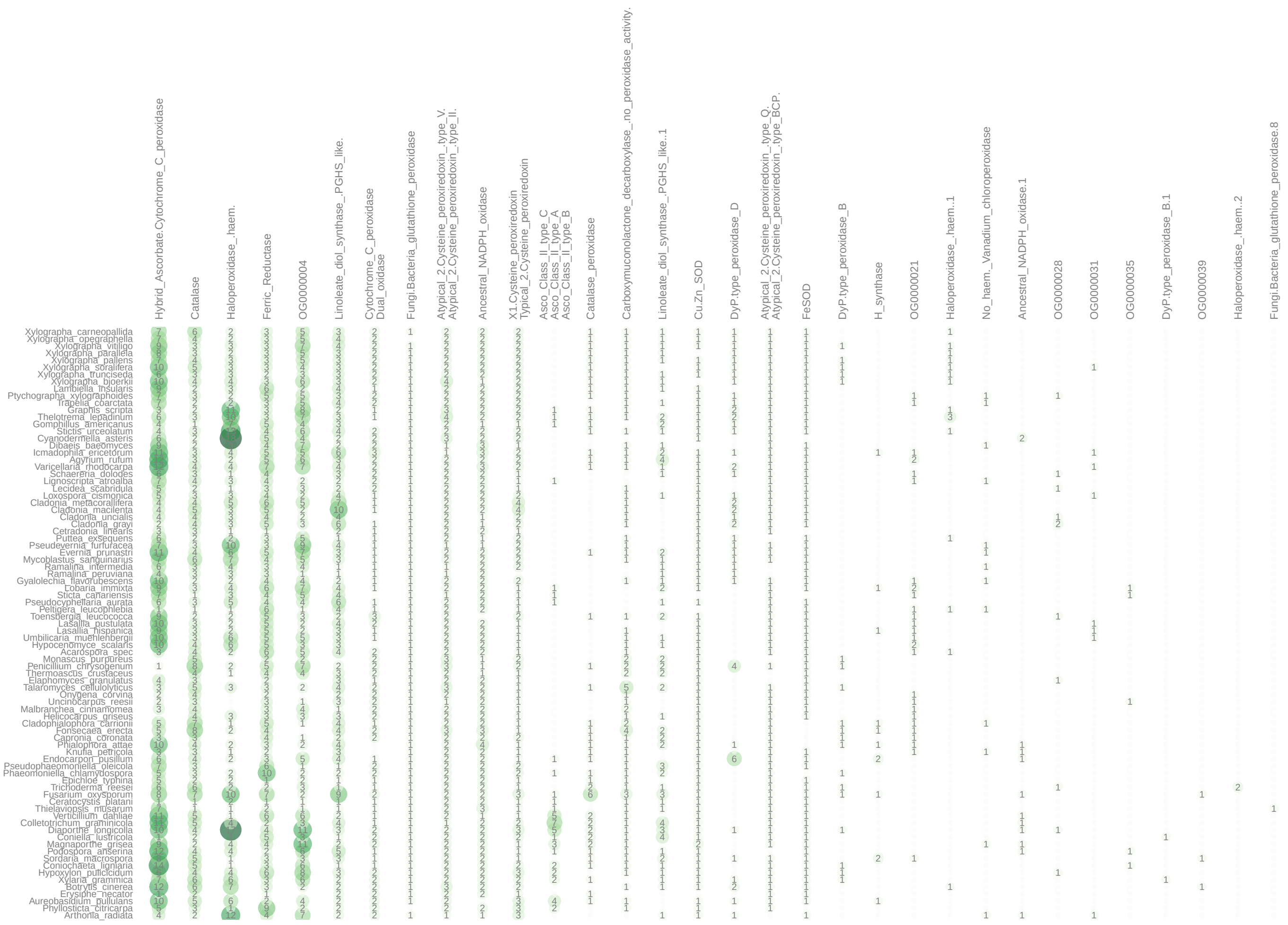

Figure 6: Overview of fungal peroxidases downloaded from RedOxibase in 83 genomes. Each column represents one orthogroup inferred by Orthofinder. Names of columns were assigned based on the presence of characterized peroxidase sequences in the respective orthogroup. Columns without any characterized peroxidase sequences still have names assigned by Orthofinder. Peroxidases which had no orthologes in any of the 83 studied genomes where excluded from the figure.

#### Figure 7: Ancestral state reconstruction results for (hemi-)cellulose degrading CAZymes

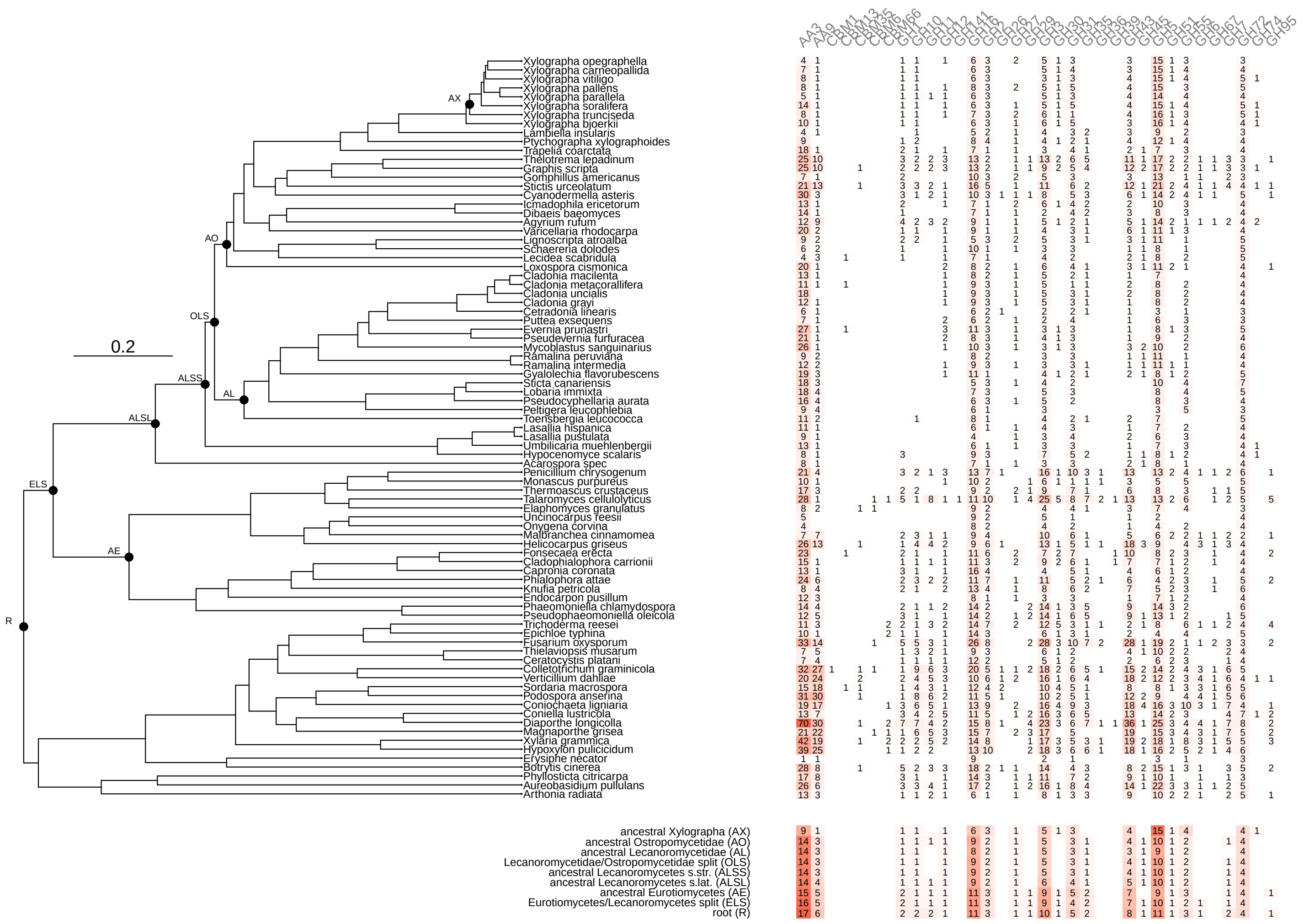

Figure 7: Heatmap of extant CAZyme numbers of families involved in cellulose and hemicellulose breakdown according to Table 6. Values below the tree are taken from ancestral state reconstructions of family sizes at different nodes.

#### Figure 8: Ancestral state reconstruction results for pectin degrading CAZymes

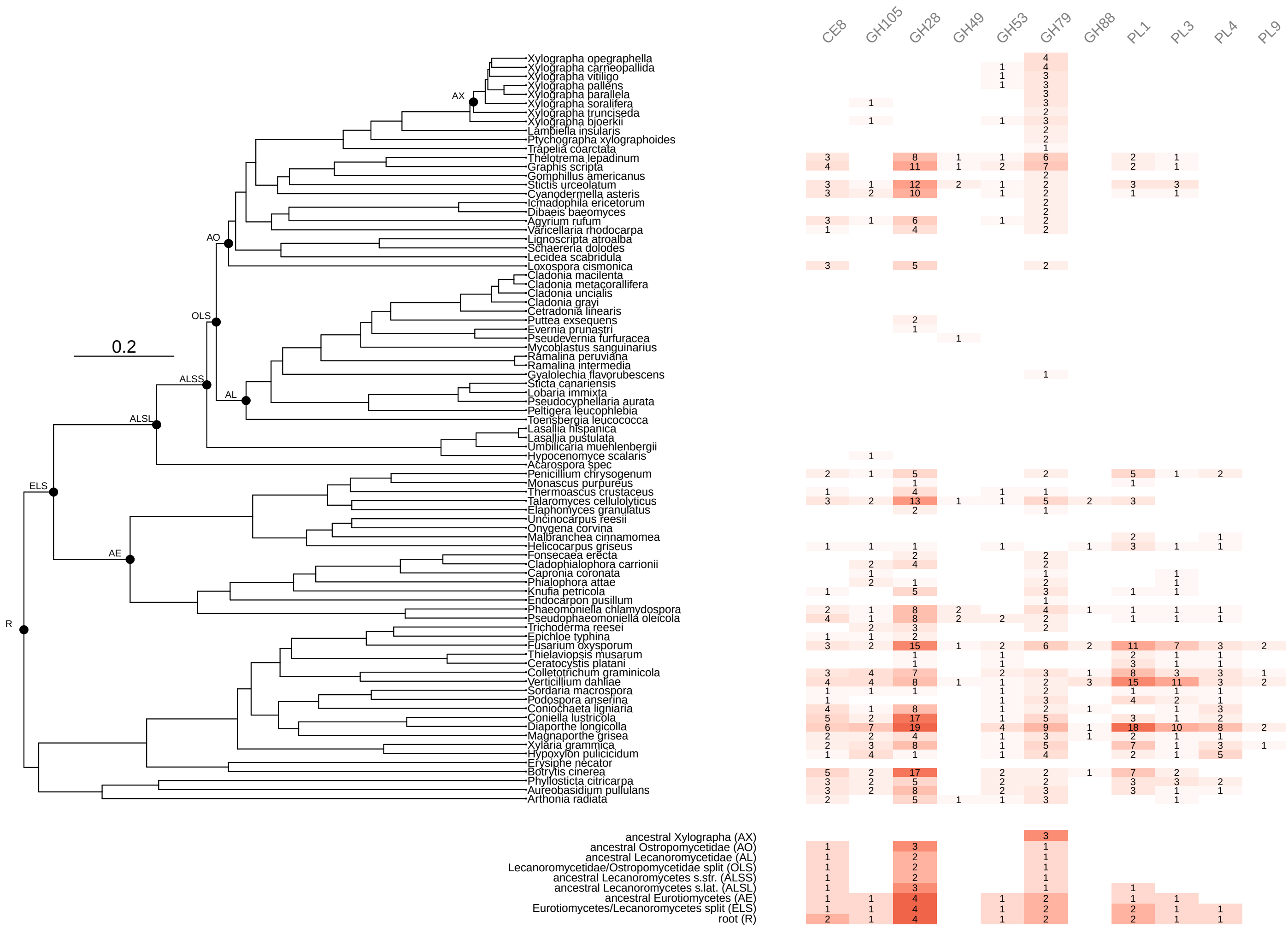

Figure 8: Heatmap of extant CAZyme numbers of families involved in pectin breakdown according to Table 6. Values below the tree are taken from ancestral state reconstructions of family sizes at different nodes.

#### Figure 9: Ancestral state reconstruction results for lignin degrading CAZymes

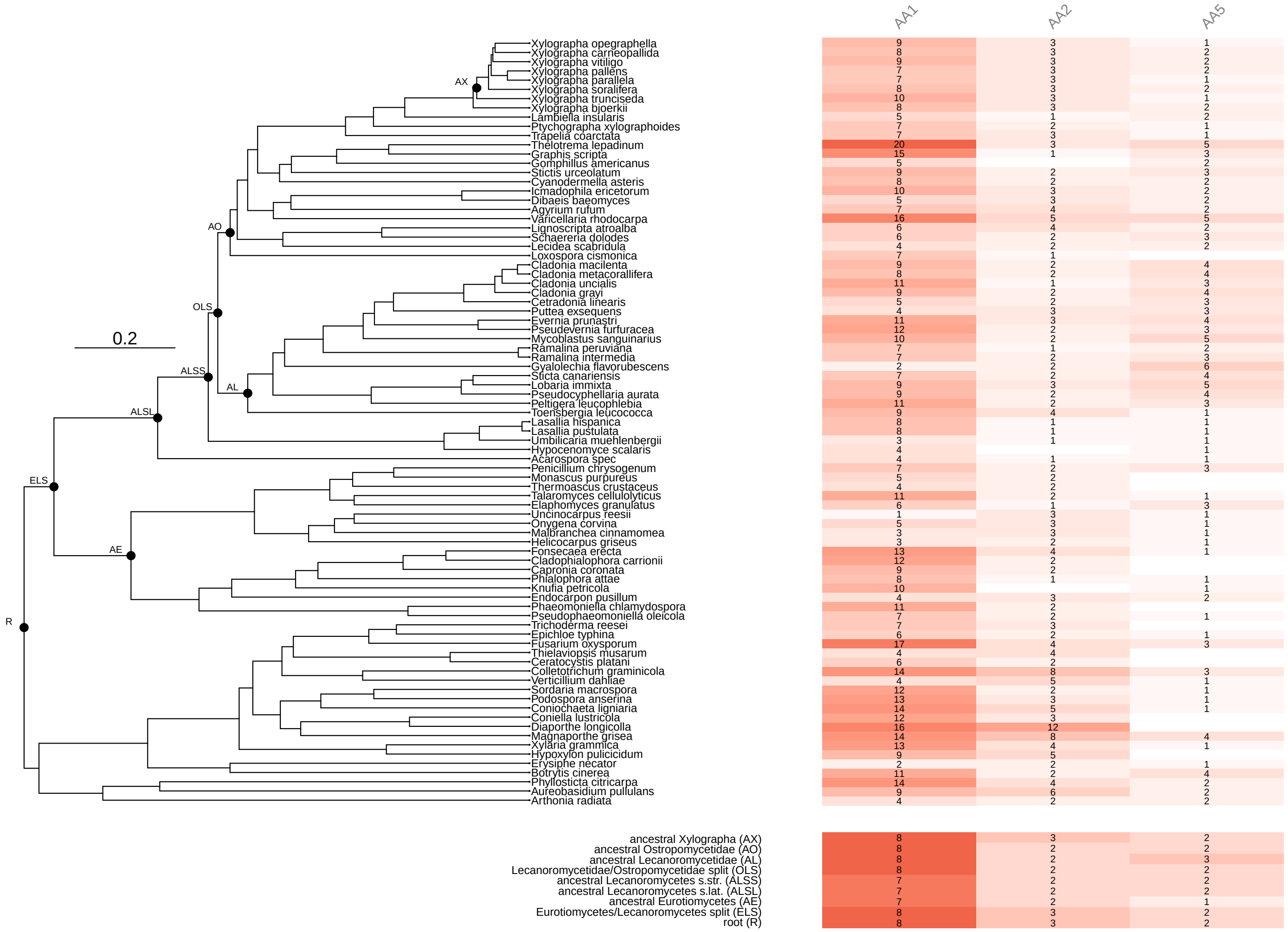

Figure 9: Heatmap of extant CAZyme numbers of families involved in lignin breakdown according to Table 6. Values below the tree are taken from ancestral state reconstructions of family sizes at different nodes.

#### Figure 10: Similarity of CAZyme sets based on PCA

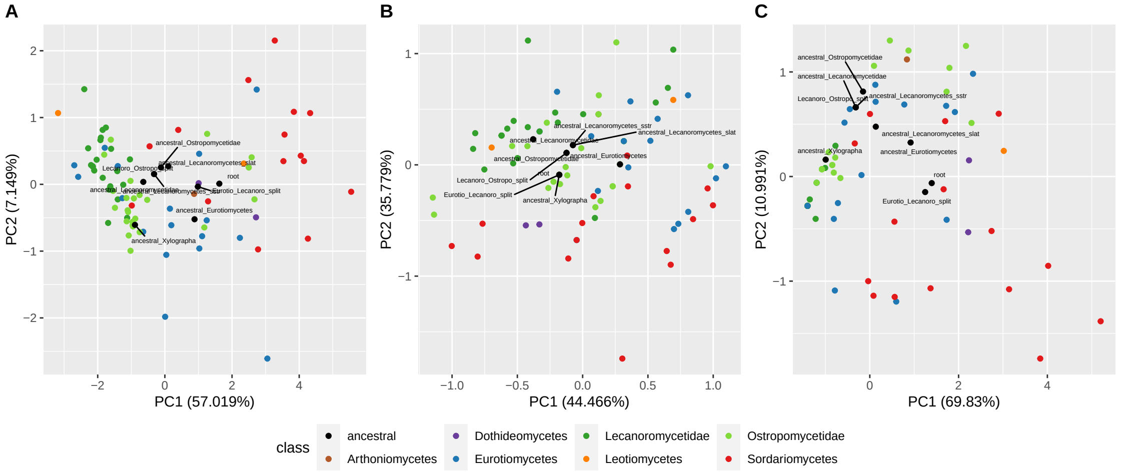

Figure 10: Overall similarity of CAZyme sets involved in (hemi)cellulose (A), lignin(B) and pectin(C) breakdown. Each dot represents the CAZyme composition of one genome. Dots are colored by broad-scale taxonomic assignment on class-level or subclass level (Lecanoromycetes). Labeled black dots refer to the composition reconstructed at ancestral nodes.

### Figure 11: Distribution and ancestral states of all CAZyme families

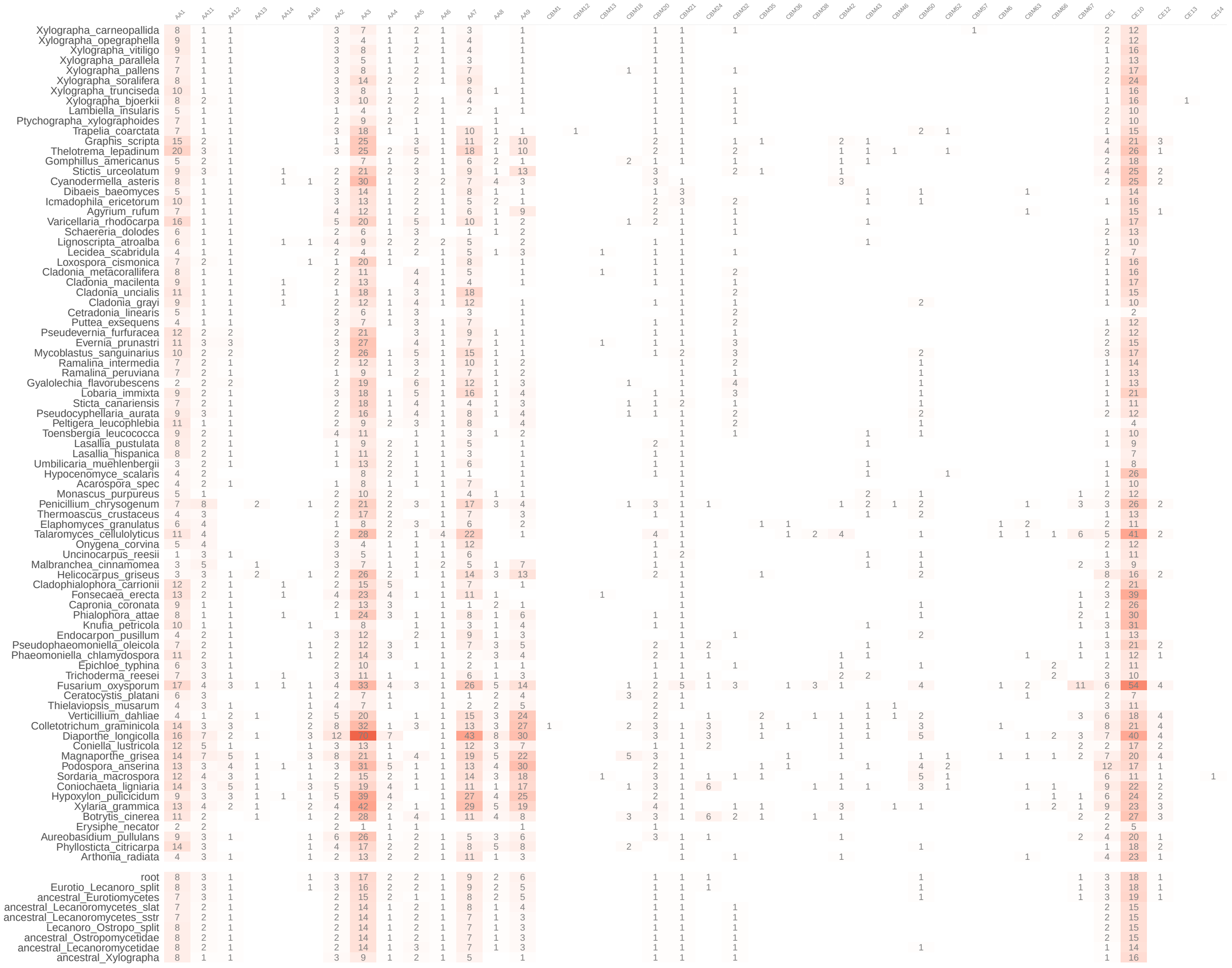

(a) part 1 of 5

Figure 11: Overall similarity of CAZyme sets involved in (hemi)cellulose (A), lignin(B) and pectin(C) breakdown. Each dot represents the CAZyme composition of one genome. Dots are colored by broad-scale taxonomic assignment on class-level or subclass level (Lecanoromycetes). Labeled black dots refer to the composition reconstructed at ancestral nodes.

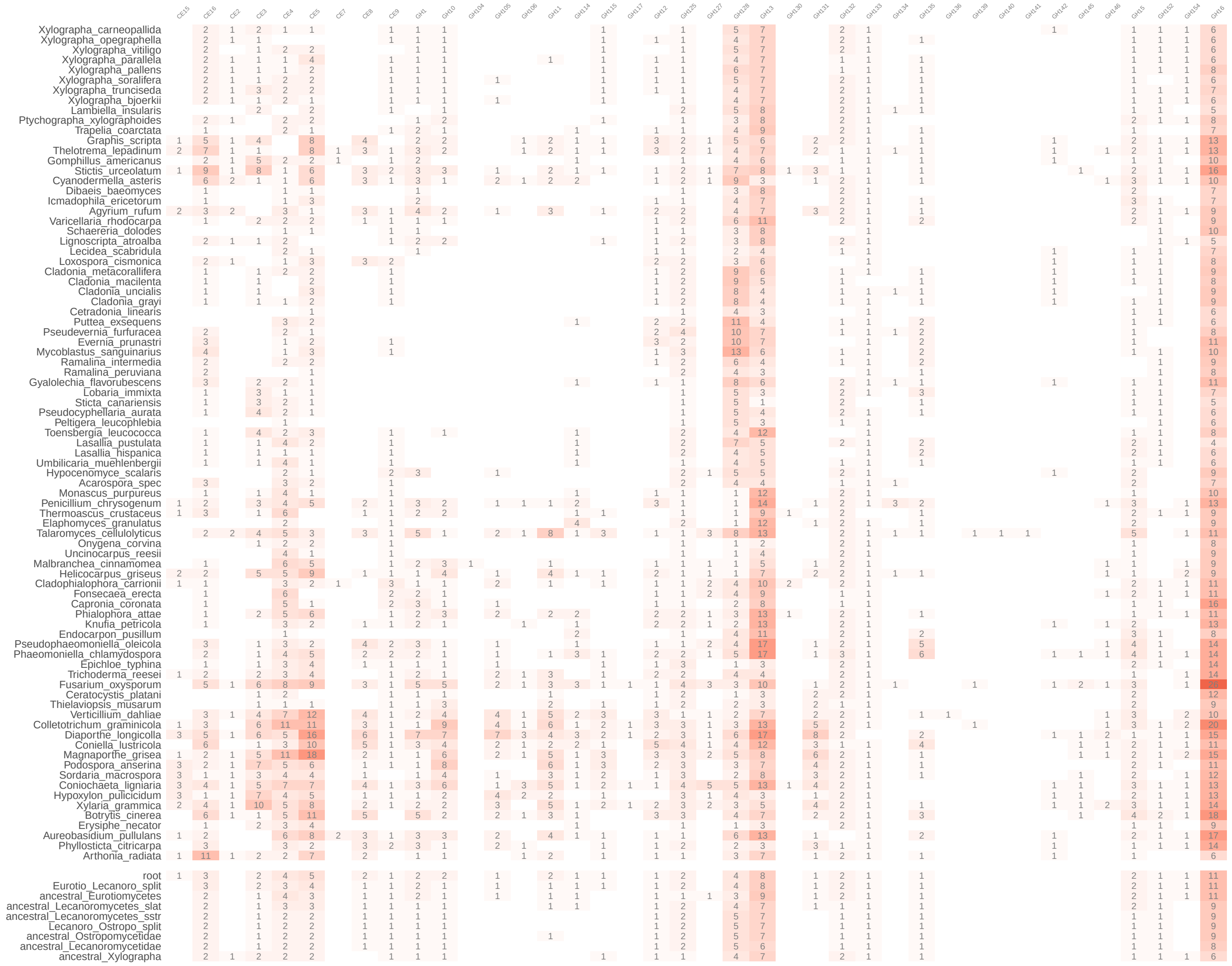

(b) part 2 of 5

Figure 11: Overall similarity of CAZyme sets involved in (hemi)cellulose (A), lignin(B) and pectin(C) breakdown. Each dot represents the CAZyme composition of one genome. Dots are colored by broad-scale taxonomic assignment on class-level or subclass level (Lecanoromycetes). Labeled black dots refer to the composition reconstructed at ancestral nodes.

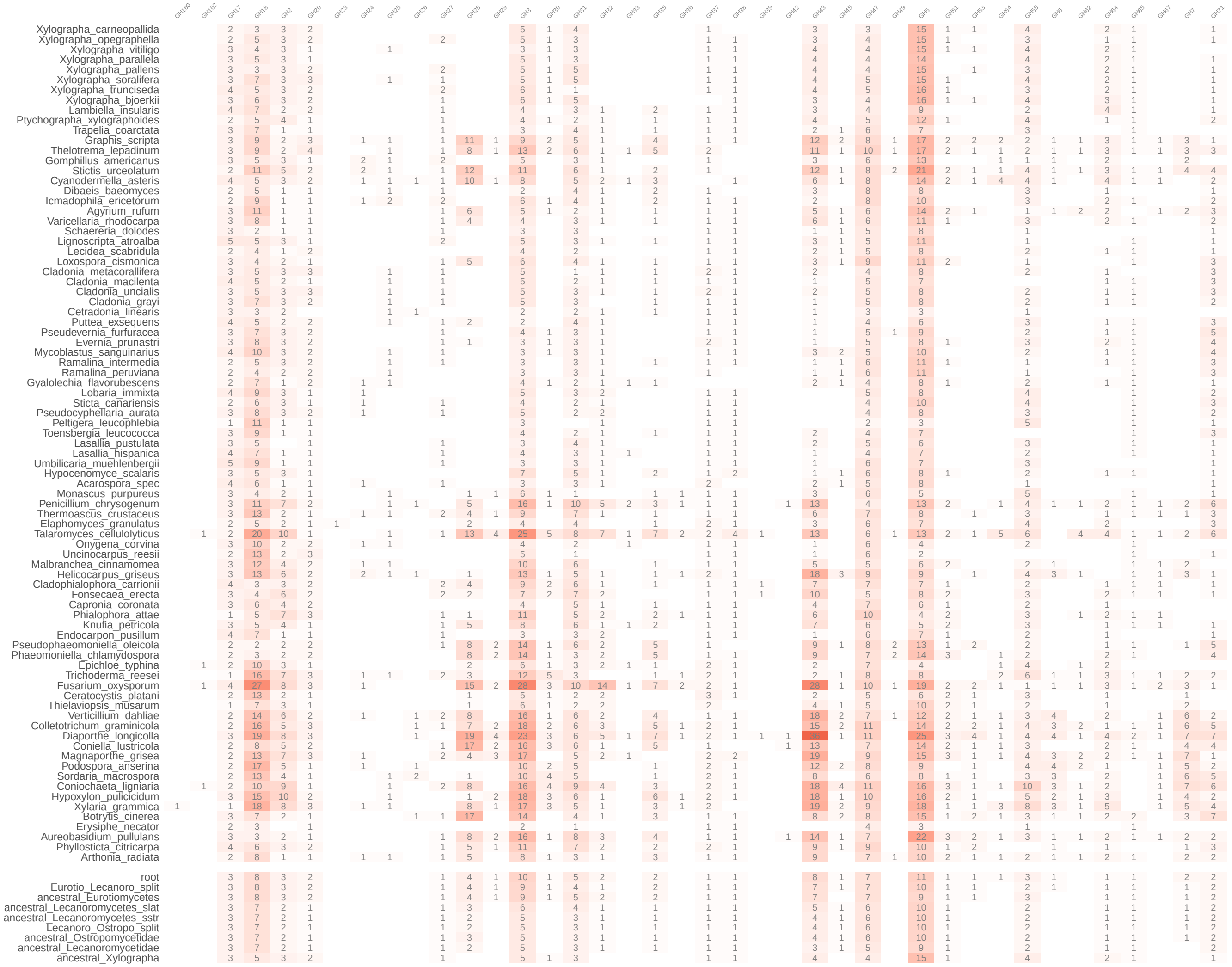

(c) part 3 of 5

Figure 11: Overall similarity of CAZyme sets involved in (hemi)cellulose (A), lignin(B) and pectin(C) breakdown. Each dot represents the CAZyme composition of one genome. Dots are colored by broad-scale taxonomic assignment on class-level or subclass level (Lecanoromycetes). Labeled black dots refer to the composition reconstructed at ancestral nodes.

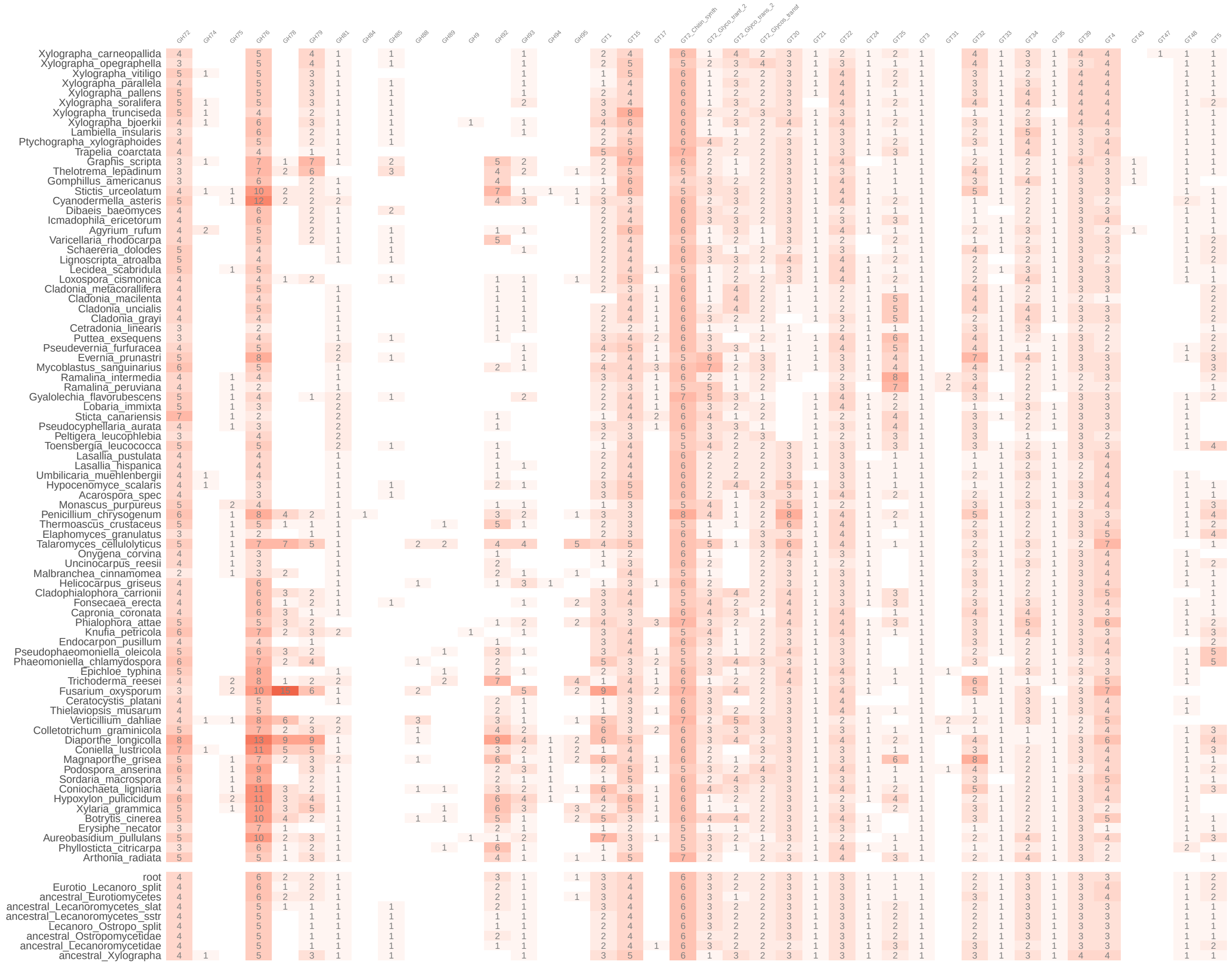

(d) part 4 of 5

Figure 11: Overall similarity of CAZyme sets involved in (hemi)cellulose (A), lignin(B) and pectin(C) breakdown. Each dot represents the CAZyme composition of one genome. Dots are colored by broad-scale taxonomic assignment on class-level or subclass level (Lecanoromycetes). Labeled black dots refer to the composition reconstructed at ancestral nodes.

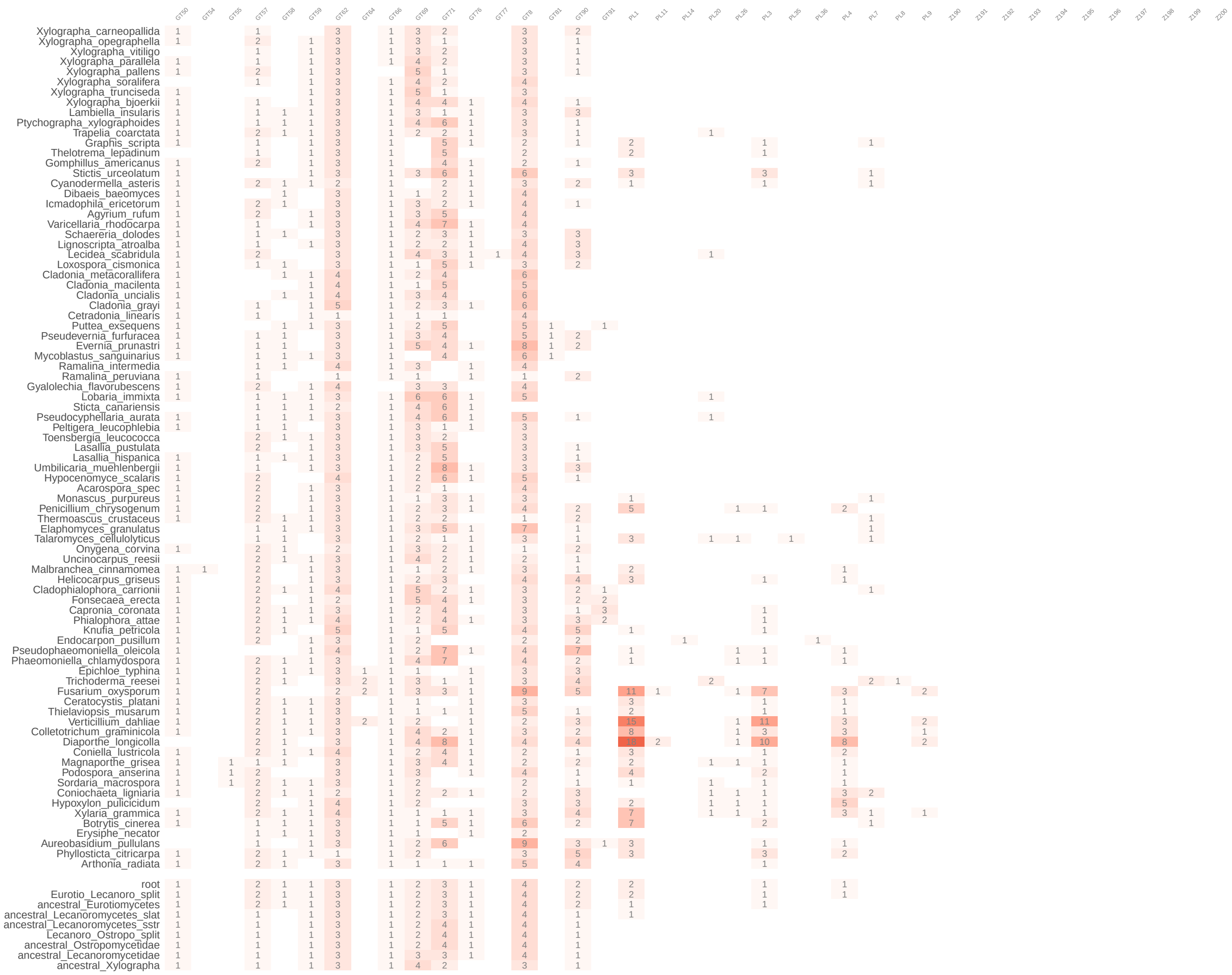

(e) part 5 of 5

Figure 11: Overall similarity of CAZyme sets involved in (hemi)cellulose (A), lignin(B) and pectin(C) breakdown. Each dot represents the CAZyme composition of one genome. Dots are colored by broad-scale taxonomic assignment on class-level or subclass level (Lecanoromycetes). Labeled black dots refer to the composition reconstructed at ancestral nodes.

### Figure 12: Overview of gene family expansion analyses results

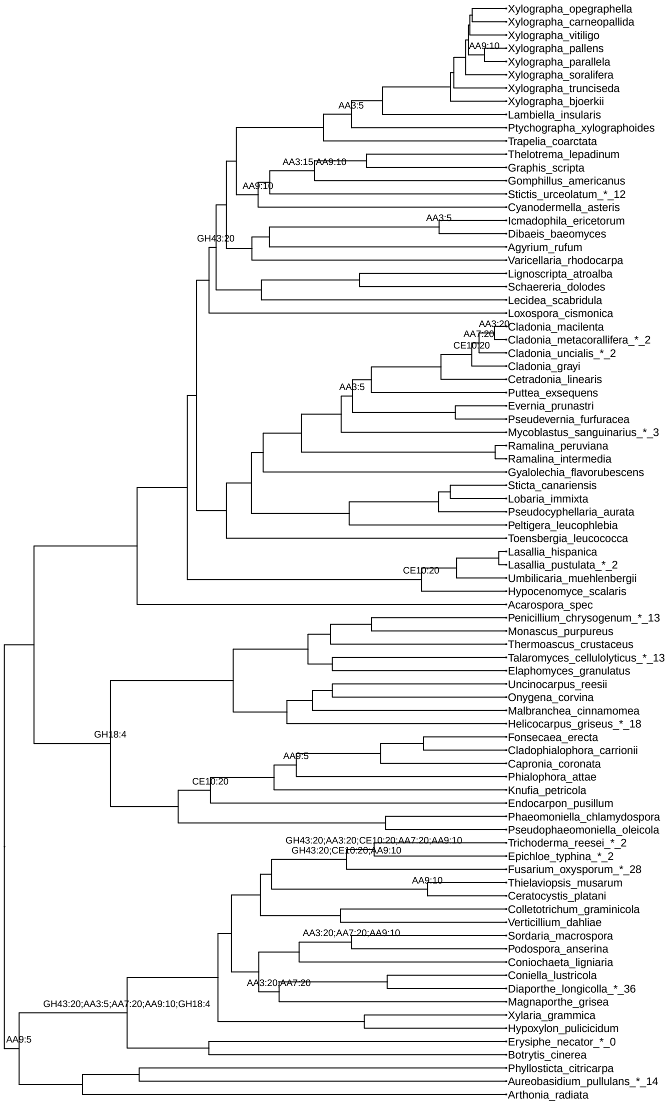

Figure 12: This tree gives a summary of our gene family expansion analyses using CAFE5. Indicated at nodes are signficantly expanded gene families and the number of independet runs it was found to be expanded, seperated by a colon. This data was used to create Figure 1.

#### Figure 13: Tree of putative invertases in GH32

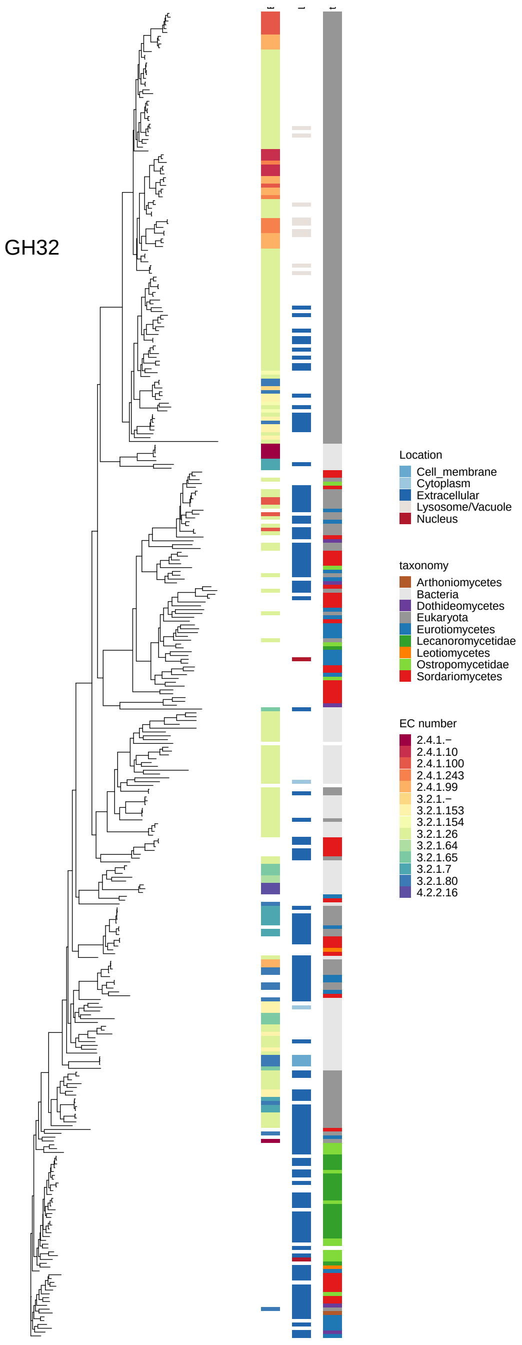

Figure 13: Maximum-likelihood gene tree of the Glycoside Hydrolase family 32 containing invertases (EC 3.1.2.26). It includes all experimentally characterized GH32 sequences from cazy.org as well as all sequences with GH32 annotations from 83 genomes. The three columns along tree tips provide additional information about the respective sequence. Left column: Enzyme Code (EC), middle column: Predicted subcellular location, Right column: Taxonomy.

#### Figures 14-64: Gene trees of CAZyme families generated with Saccharis

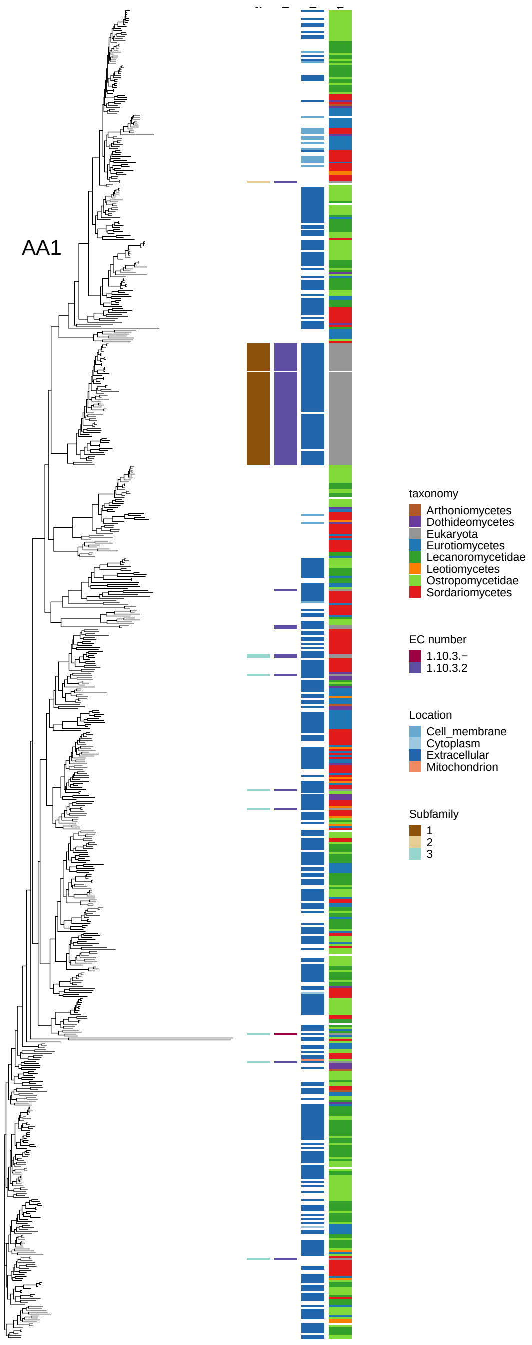

Figure 14: Maximum-likelihood gene tree of CAZyme families involved in (hemi-)cellulose, pectin and lignin breakdown. It includes all experimentally characterized sequences for the family downloaded from cazy.org as well as all sequences with annotations for this family in 83 genomes. The family name is given in the plot. The three columns along tree tips provide additional information about the respective sequence. Left column: Enzyme Code (EC), middle column: Predicted subcellular location, Right column: Taxonomy.

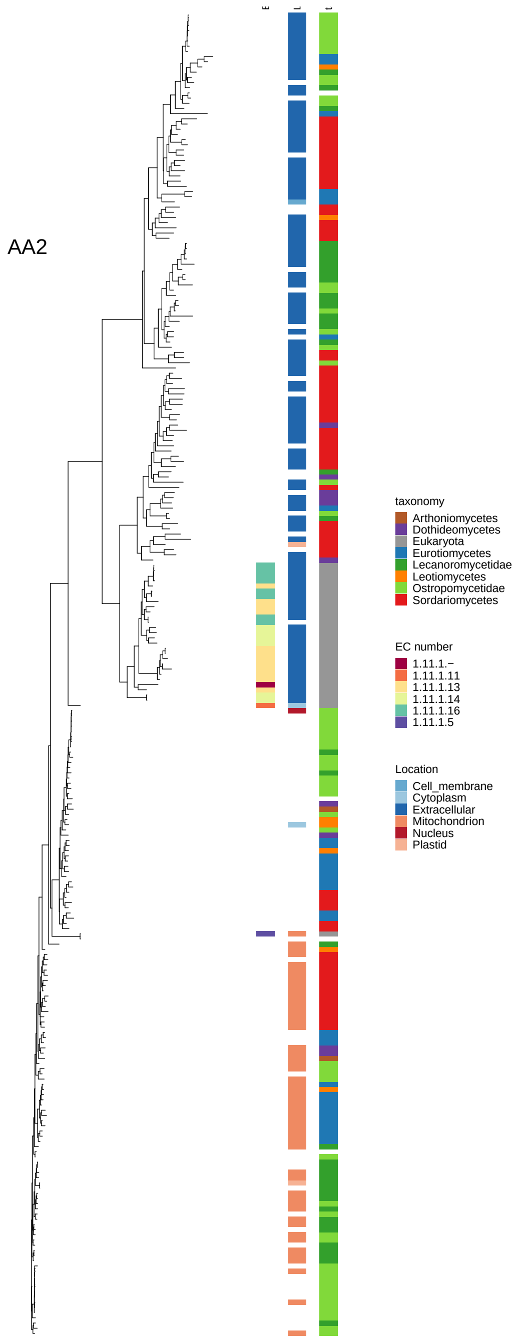

Figure 15: Maximum-likelihood gene tree of CAZyme families involved in (hemi-)cellulose, pectin and lignin breakdown. It includes all experimentally characterized sequences for the family downloaded from cazy.org as well as all sequences with annotations for this family in 83 genomes. The family name is given in the plot. The three columns along tree tips provide additional information about the respective sequence. Left column: Enzyme Code (EC), middle column: Predicted subcellular location, Right column: Taxonomy.

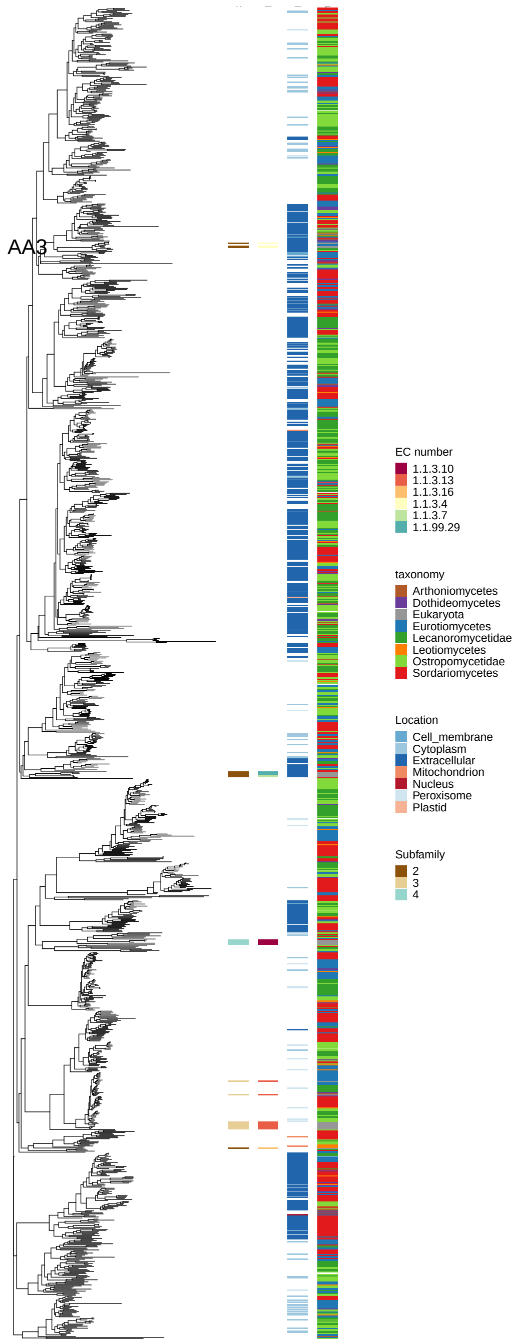

Figure 16: Maximum-likelihood gene tree of CAZyme families involved in (hemi-)cellulose, pectin and lignin breakdown. It includes all experimentally characterized sequences for the family downloaded from cazy.org as well as all sequences with annotations for this family in 83 genomes. The family name is given in the plot. The three columns along tree tips provide additional information about the respective sequence. Left column: Enzyme Code (EC), middle column: Predicted subcellular location, Right column: Taxonomy.

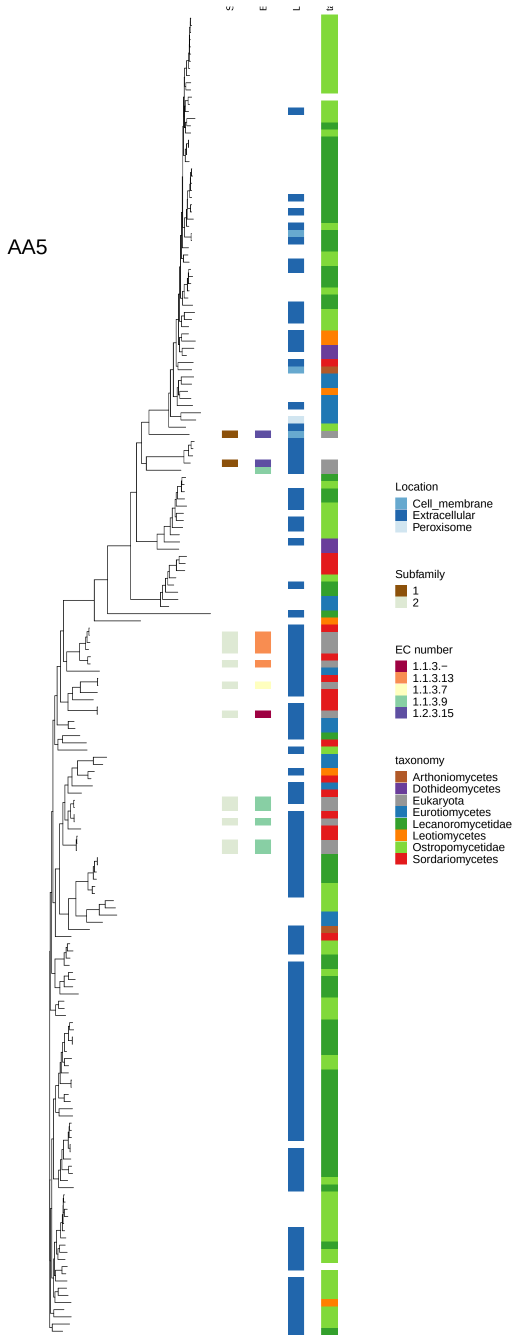

Figure 17: Maximum-likelihood gene tree of CAZyme families involved in (hemi-)cellulose, pectin and lignin breakdown. It includes all experimentally characterized sequences for the family downloaded from cazy.org as well as all sequences with annotations for this family in 83 genomes. The family name is given in the plot. The three columns along tree tips provide additional information about the respective sequence. Left column: Enzyme Code (EC), middle column: Predicted subcellular location, Right column: Taxonomy.

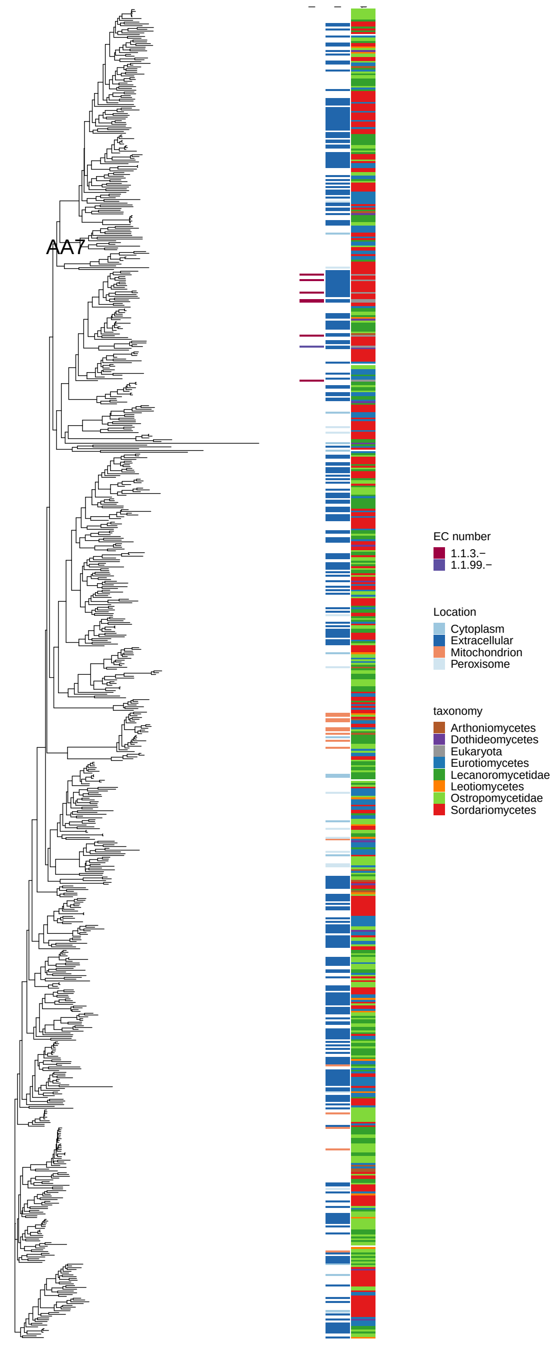

Figure 18: Maximum-likelihood gene tree of CAZyme families involved in (hemi-)cellulose, pectin and lignin breakdown. It includes all experimentally characterized sequences for the family downloaded from cazy.org as well as all sequences with annotations for this family in 83 genomes. The family name is given in the plot. The three columns along tree tips provide additional information about the respective sequence. Left column: Enzyme Code (EC), middle column: Predicted subcellular location, Right column: Taxonomy.

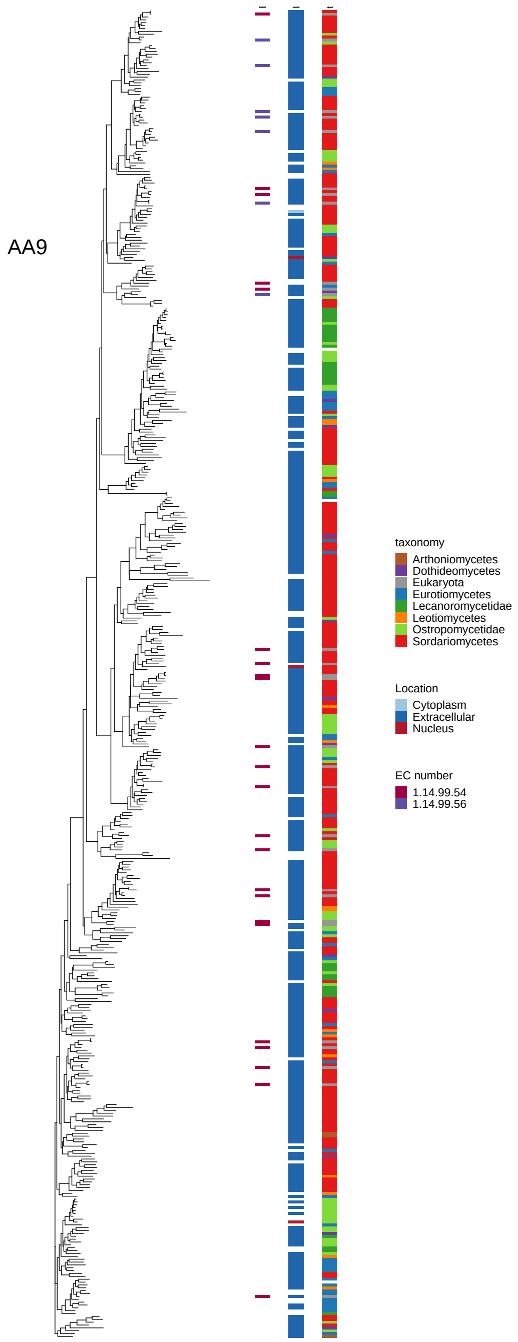

Figure 19: Maximum-likelihood gene tree of CAZyme families involved in (hemi-)cellulose, pectin and lignin breakdown. It includes all experimentally characterized sequences for the family downloaded from cazy.org as well as all sequences with annotations for this family in 83 genomes. The family name is given in the plot. The three columns along tree tips provide additional information about the respective sequence. Left column: Enzyme Code (EC), middle column: Predicted subcellular location, Right column: Taxonomy.

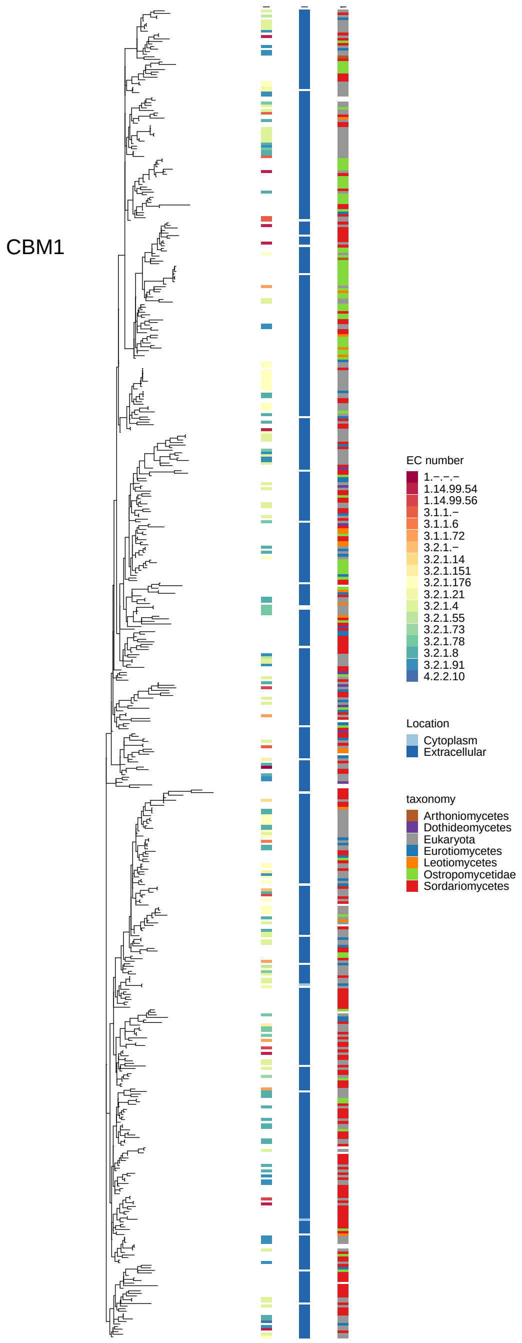

Figure 20: Maximum-likelihood gene tree of CAZyme families involved in (hemi-)cellulose, pectin and lignin breakdown. It includes all experimentally characterized sequences for the family downloaded from cazy.org as well as all sequences with annotations for this family in 83 genomes. The family name is given in the plot. The three columns along tree tips provide additional information about the respective sequence. Left column: Enzyme Code (EC), middle column: Predicted subcellular location, Right column: Taxonomy.

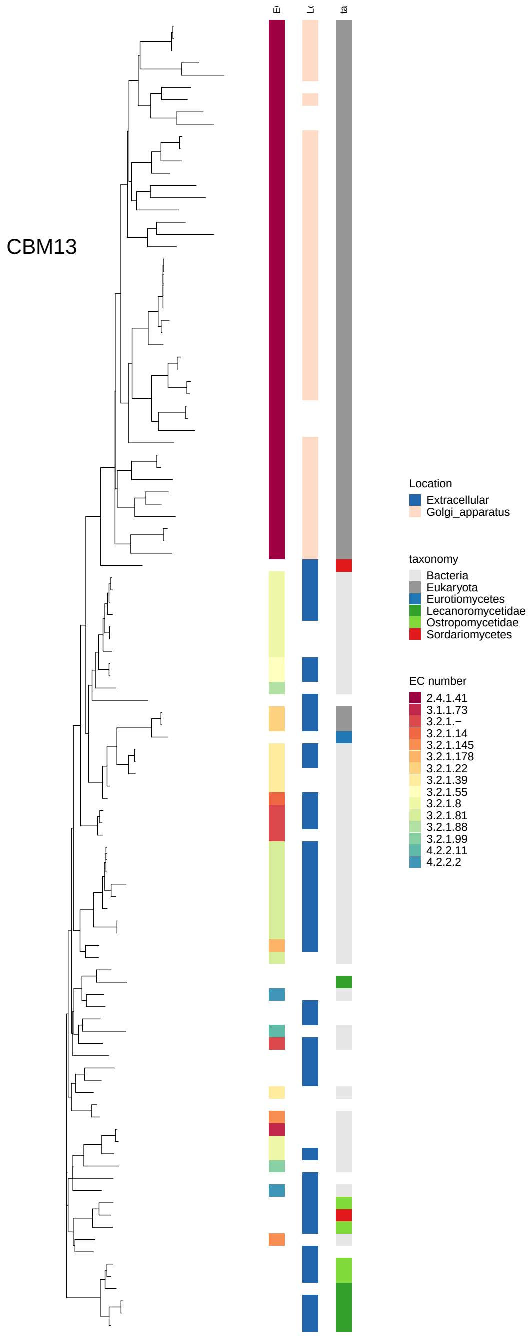

Figure 21: Maximum-likelihood gene tree of CAZyme families involved in (hemi-)cellulose, pectin and lignin breakdown. It includes all experimentally characterized sequences for the family downloaded from cazy.org as well as all sequences with annotations for this family in 83 genomes. The family name is given in the plot. The three columns along tree tips provide additional information about the respective sequence. Left column: Enzyme Code (EC), middle column: Predicted subcellular location, Right column: Taxonomy.

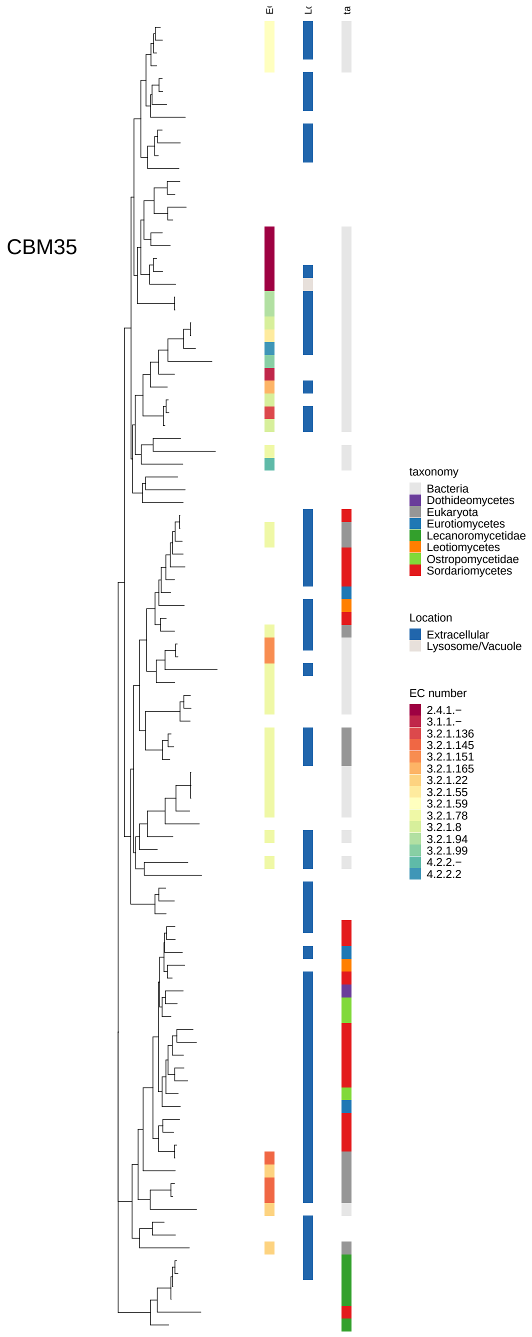

Figure 22: Maximum-likelihood gene tree of CAZyme families involved in (hemi-)cellulose, pectin and lignin breakdown. It includes all experimentally characterized sequences for the family downloaded from cazy.org as well as all sequences with annotations for this family in 83 genomes. The family name is given in the plot. The three columns along tree tips provide additional information about the respective sequence. Left column: Enzyme Code (EC), middle column: Predicted subcellular location, Right column: Taxonomy.

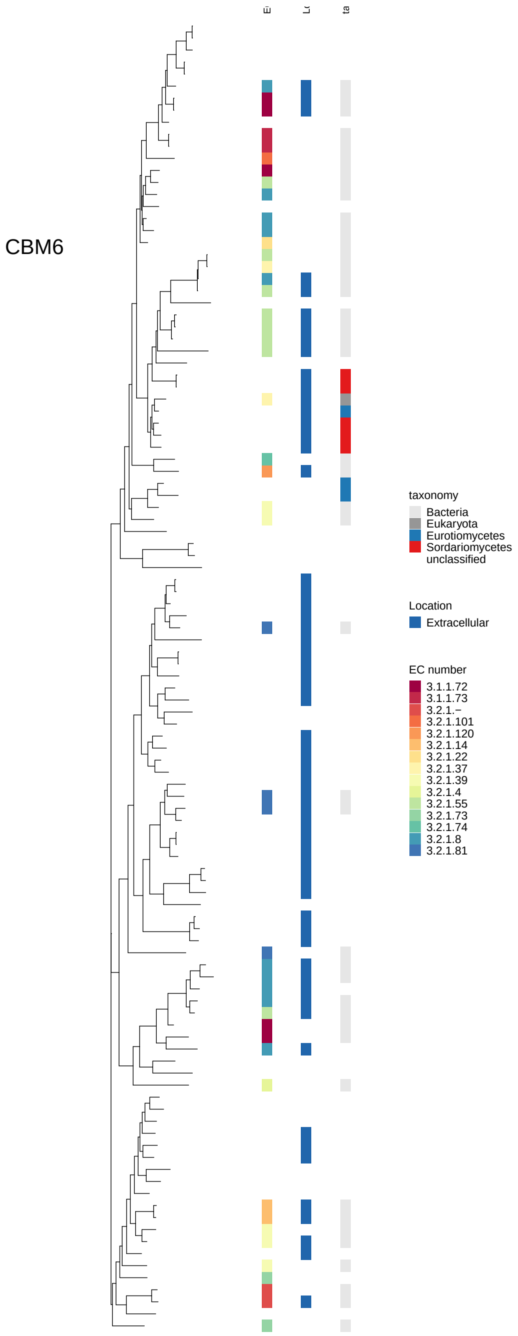

Figure 23: Maximum-likelihood gene tree of CAZyme families involved in (hemi-)cellulose, pectin and lignin breakdown. It includes all experimentally characterized sequences for the family downloaded from cazy.org as well as all sequences with annotations for this family in 83 genomes. The family name is given in the plot. The three columns along tree tips provide additional information about the respective sequence. Left column: Enzyme Code (EC), middle column: Predicted subcellular location, Right column: Taxonomy.

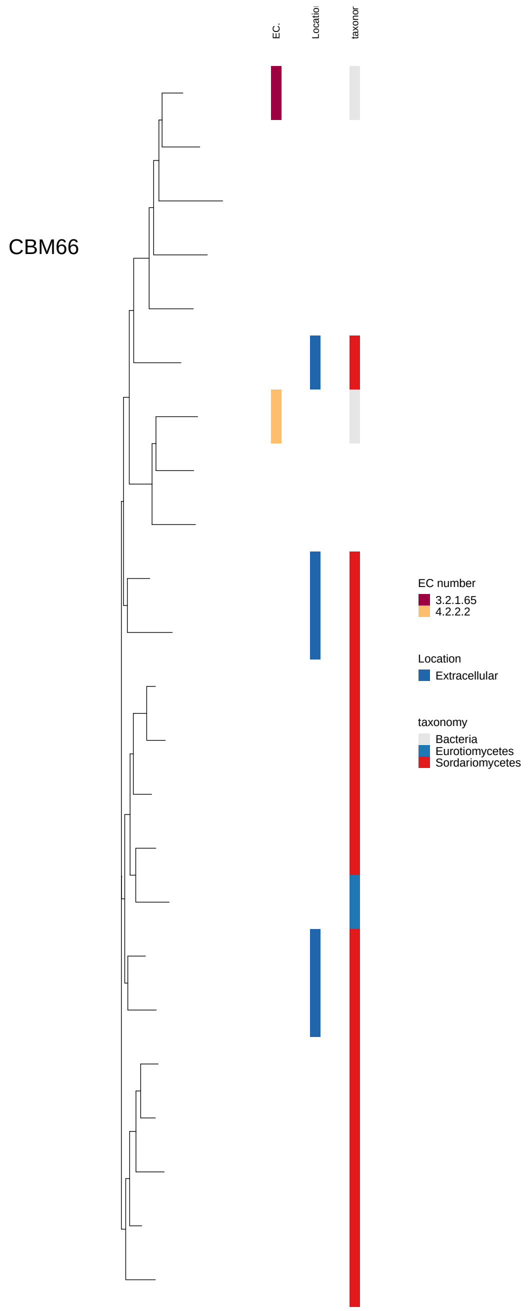

Figure 24: Maximum-likelihood gene tree of CAZyme families involved in (hemi-)cellulose, pectin and lignin breakdown. It includes all experimentally characterized sequences for the family downloaded from cazy.org as well as all sequences with annotations for this family in 83 genomes. The family name is given in the plot. The three columns along tree tips provide additional information about the respective sequence. Left column: Enzyme Code (EC), middle column: Predicted subcellular location, Right column: Taxonomy.

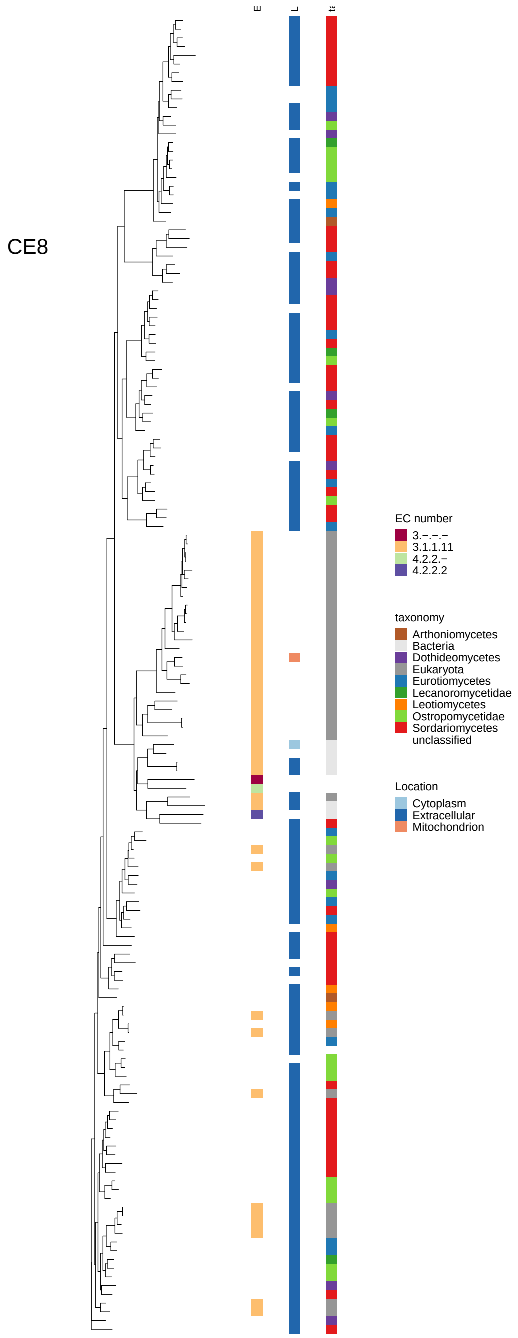

Figure 25: Maximum-likelihood gene tree of CAZyme families involved in (hemi-)cellulose, pectin and lignin breakdown. It includes all experimentally characterized sequences for the family downloaded from cazy.org as well as all sequences with annotations for this family in 83 genomes. The family name is given in the plot. The three columns along tree tips provide additional information about the respective sequence. Left column: Enzyme Code (EC), middle column: Predicted subcellular location, Right column: Taxonomy.

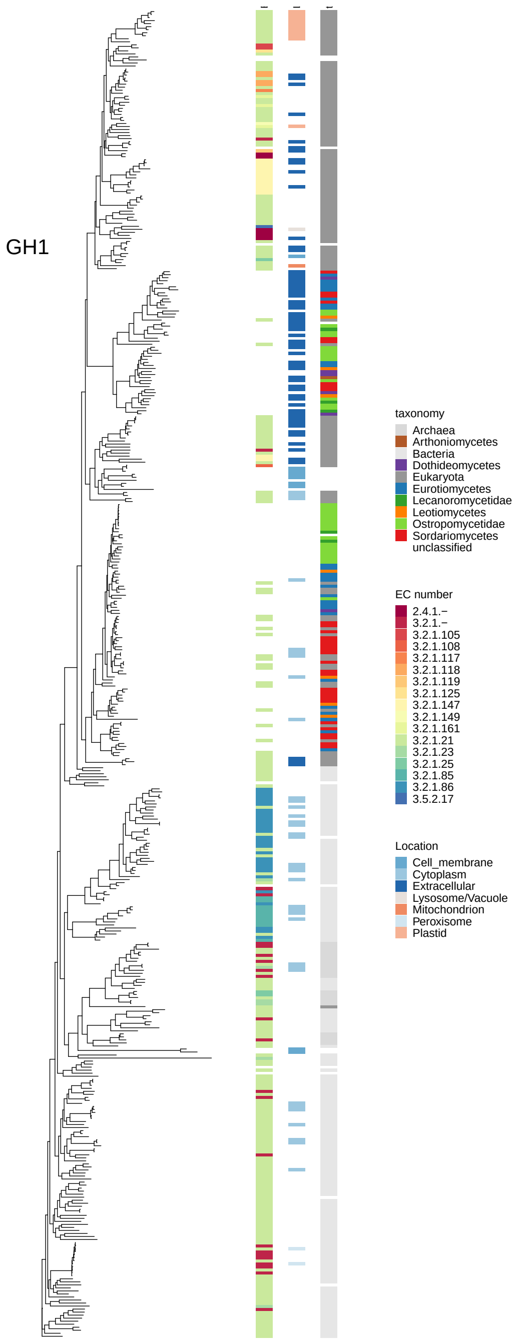

Figure 26: Maximum-likelihood gene tree of CAZyme families involved in (hemi-)cellulose, pectin and lignin breakdown. It includes all experimentally characterized sequences for the family downloaded from cazy.org as well as all sequences with annotations for this family in 83 genomes. The family name is given in the plot. The three columns along tree tips provide additional information about the respective sequence. Left column: Enzyme Code (EC), middle column: Predicted subcellular location, Right column: Taxonomy.

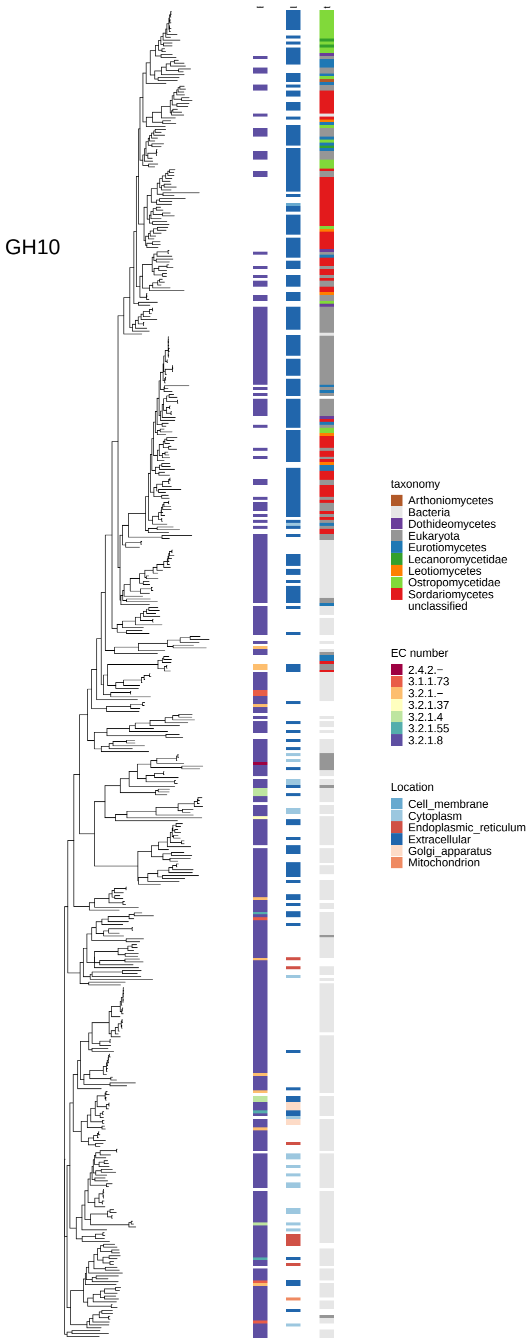

Figure 27: Maximum-likelihood gene tree of CAZyme families involved in (hemi-)cellulose, pectin and lignin breakdown. It includes all experimentally characterized sequences for the family downloaded from cazy.org as well as all sequences with annotations for this family in 83 genomes. The family name is given in the plot. The three columns along tree tips provide additional information about the respective sequence. Left column: Enzyme Code (EC), middle column: Predicted subcellular location, Right column: Taxonomy.

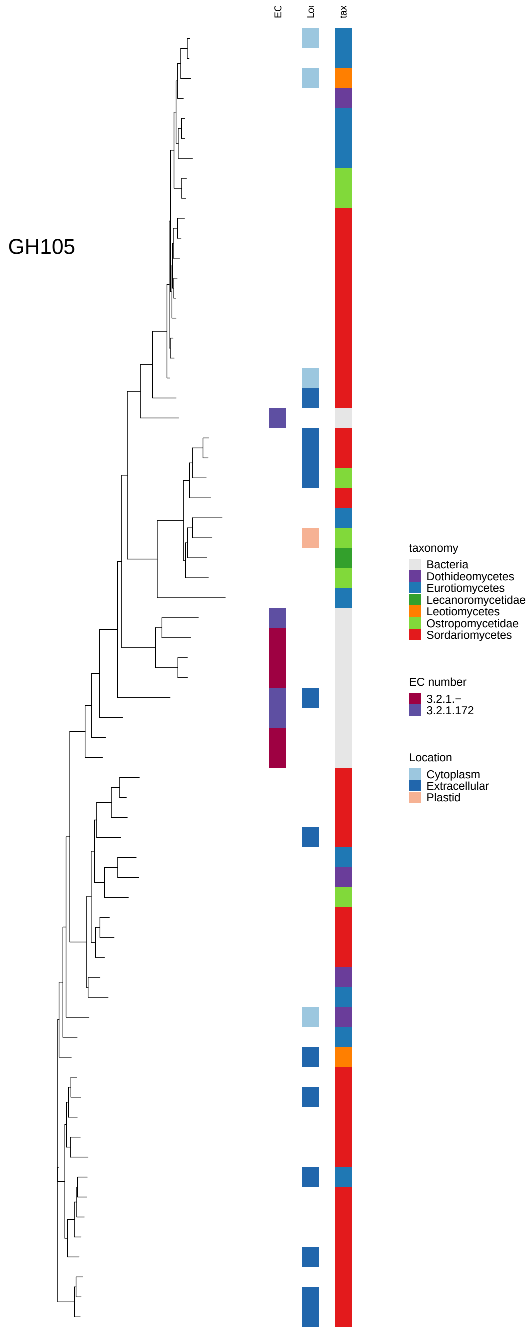

Figure 28: Maximum-likelihood gene tree of CAZyme families involved in (hemi-)cellulose, pectin and lignin breakdown. It includes all experimentally characterized sequences for the family downloaded from cazy.org as well as all sequences with annotations for this family in 83 genomes. The family name is given in the plot. The three columns along tree tips provide additional information about the respective sequence. Left column: Enzyme Code (EC), middle column: Predicted subcellular location, Right column: Taxonomy.

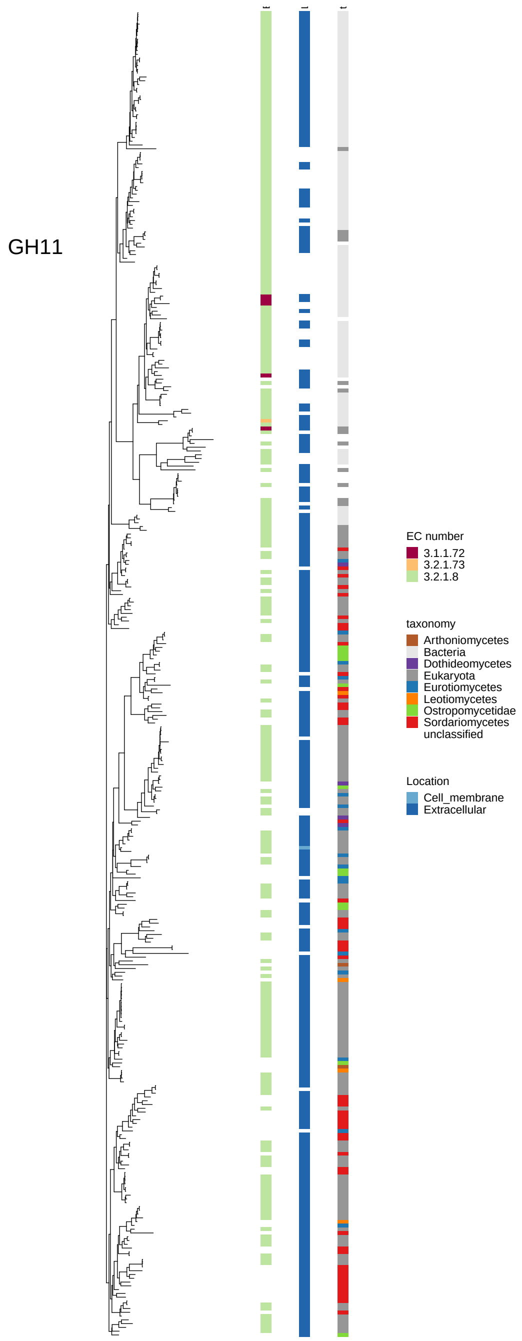

Figure 29: Maximum-likelihood gene tree of CAZyme families involved in (hemi-)cellulose, pectin and lignin breakdown. It includes all experimentally characterized sequences for the family downloaded from cazy.org as well as all sequences with annotations for this family in 83 genomes. The family name is given in the plot. The three columns along tree tips provide additional information about the respective sequence. Left column: Enzyme Code (EC), middle column: Predicted subcellular location, Right column: Taxonomy.

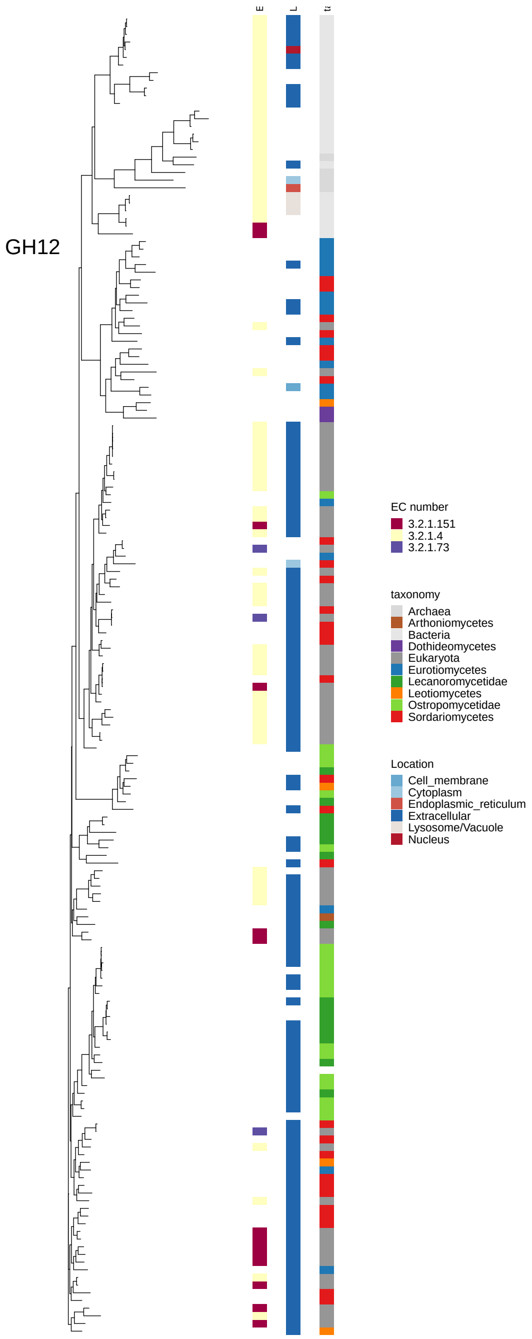

Figure 30: Maximum-likelihood gene tree of CAZyme families involved in (hemi-)cellulose, pectin and lignin breakdown. It includes all experimentally characterized sequences for the family downloaded from cazy.org as well as all sequences with annotations for this family in 83 genomes. The family name is given in the plot. The three columns along tree tips provide additional information about the respective sequence. Left column: Enzyme Code (EC), middle column: Predicted subcellular location, Right column: Taxonomy.

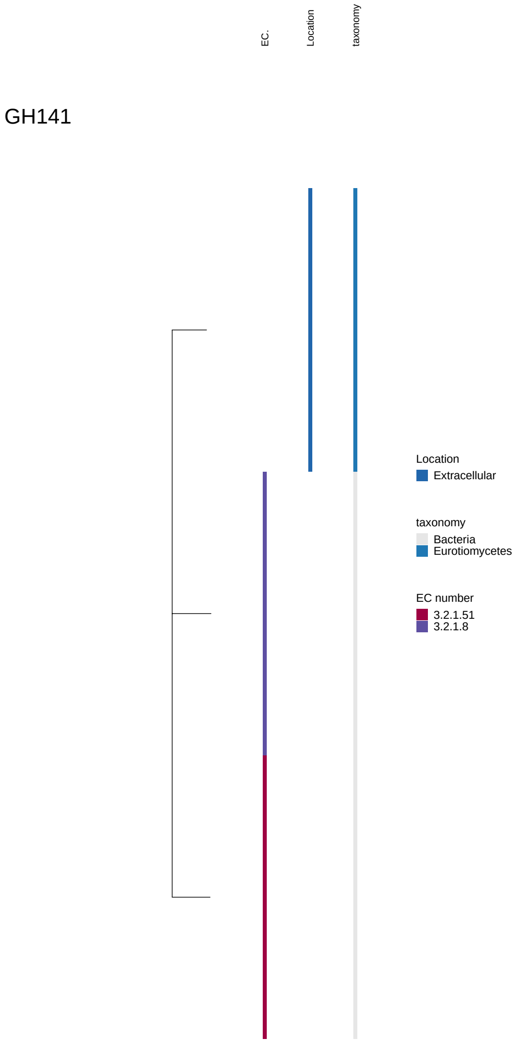

Figure 31: Maximum-likelihood gene tree of CAZyme families involved in (hemi-)cellulose, pectin and lignin breakdown. It includes all experimentally characterized sequences for the family downloaded from cazy.org as well as all sequences with annotations for this family in 83 genomes. The family name is given in the plot. The three columns along tree tips provide additional information about the respective sequence. Left column: Enzyme Code (EC), middle column: Predicted subcellular location, Right column: Taxonomy.

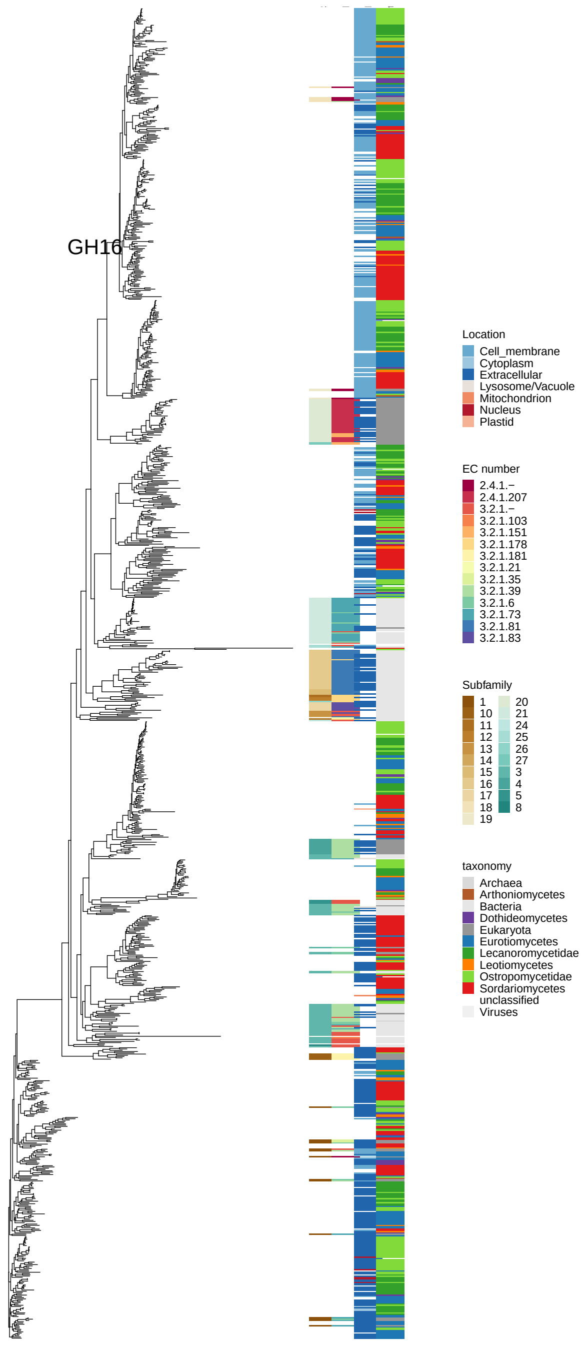

Figure 32: Maximum-likelihood gene tree of CAZyme families involved in (hemi-)cellulose, pectin and lignin breakdown. It includes all experimentally characterized sequences for the family downloaded from cazy.org as well as all sequences with annotations for this family in 83 genomes. The family name is given in the plot. The three columns along tree tips provide additional information about the respective sequence. Left column: Enzyme Code (EC), middle column: Predicted subcellular location, Right column: Taxonomy.

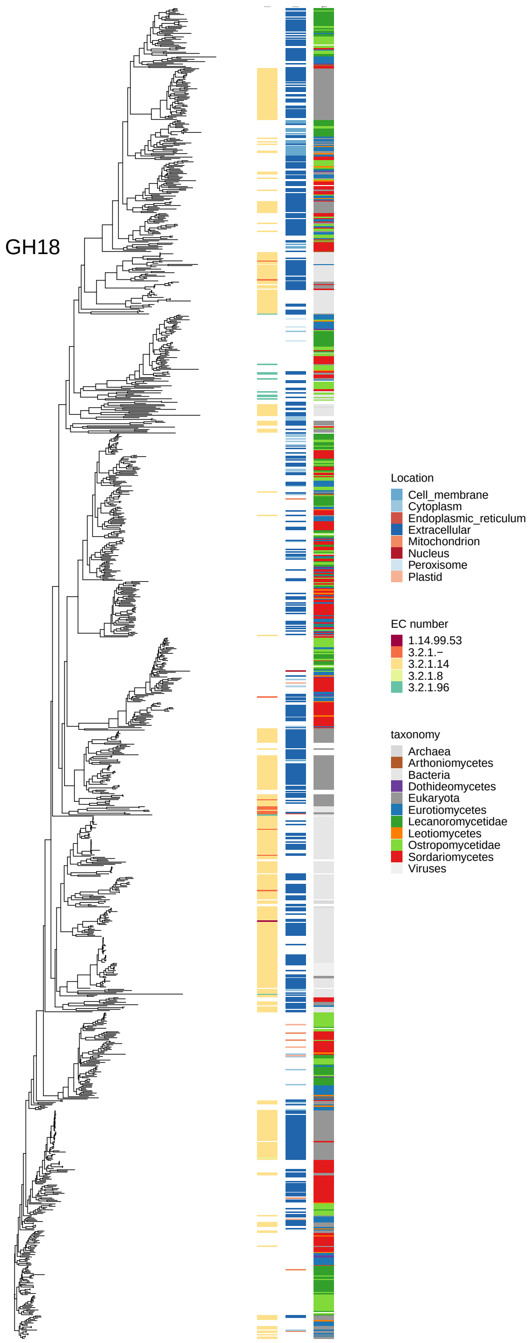

Figure 33: Maximum-likelihood gene tree of CAZyme families involved in (hemi-)cellulose, pectin and lignin breakdown. It includes all experimentally characterized sequences for the family downloaded from cazy.org as well as all sequences with annotations for this family in 83 genomes. The family name is given in the plot. The three columns along tree tips provide additional information about the respective sequence. Left column: Enzyme Code (EC), middle column: Predicted subcellular location, Right column: Taxonomy.

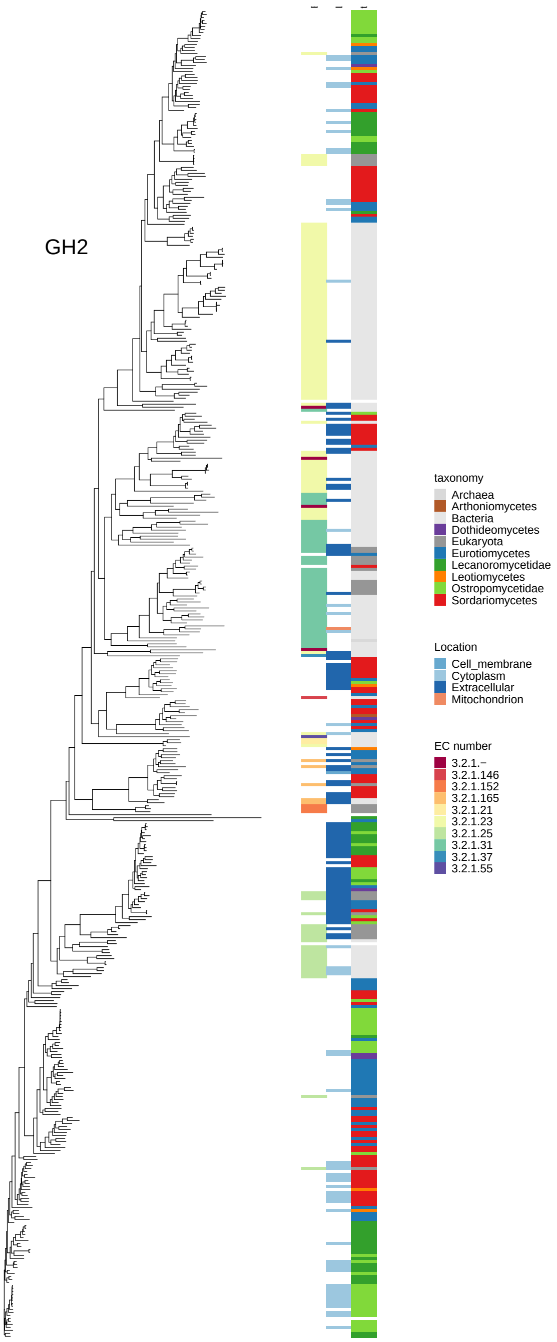

Figure 34: Maximum-likelihood gene tree of CAZyme families involved in (hemi-)cellulose, pectin and lignin breakdown. It includes all experimentally characterized sequences for the family downloaded from cazy.org as well as all sequences with annotations for this family in 83 genomes. The family name is given in the plot. The three columns along tree tips provide additional information about the respective sequence. Left column: Enzyme Code (EC), middle column: Predicted subcellular location, Right column: Taxonomy.

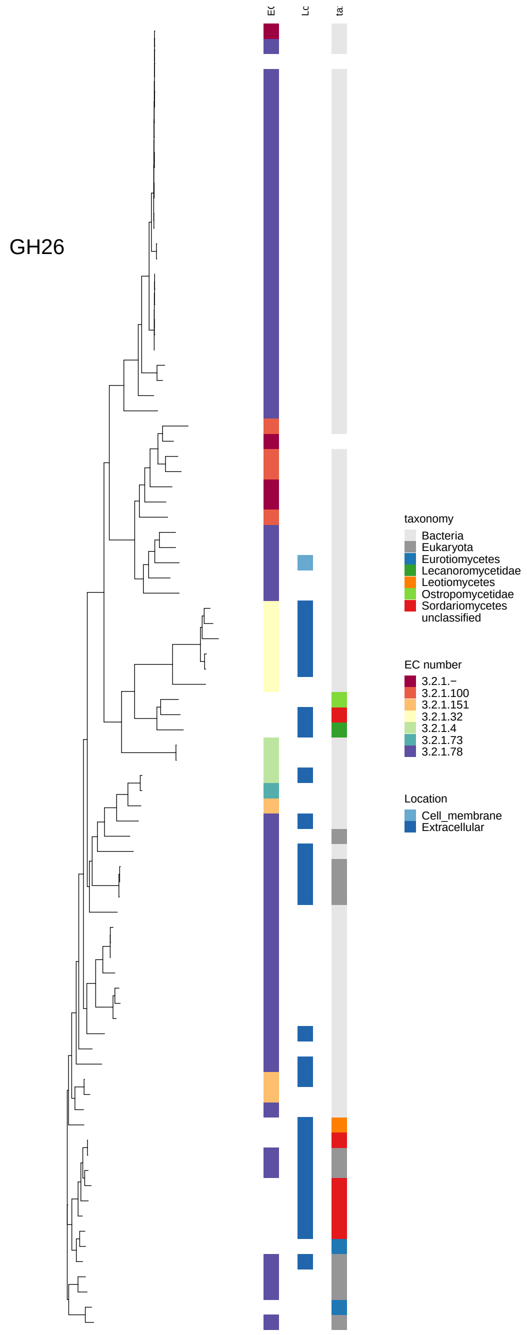

Figure 35: Maximum-likelihood gene tree of CAZyme families involved in (hemi-)cellulose, pectin and lignin breakdown. It includes all experimentally characterized sequences for the family downloaded from cazy.org as well as all sequences with annotations for this family in 83 genomes. The family name is given in the plot. The three columns along tree tips provide additional information about the respective sequence. Left column: Enzyme Code (EC), middle column: Predicted subcellular location, Right column: Taxonomy.

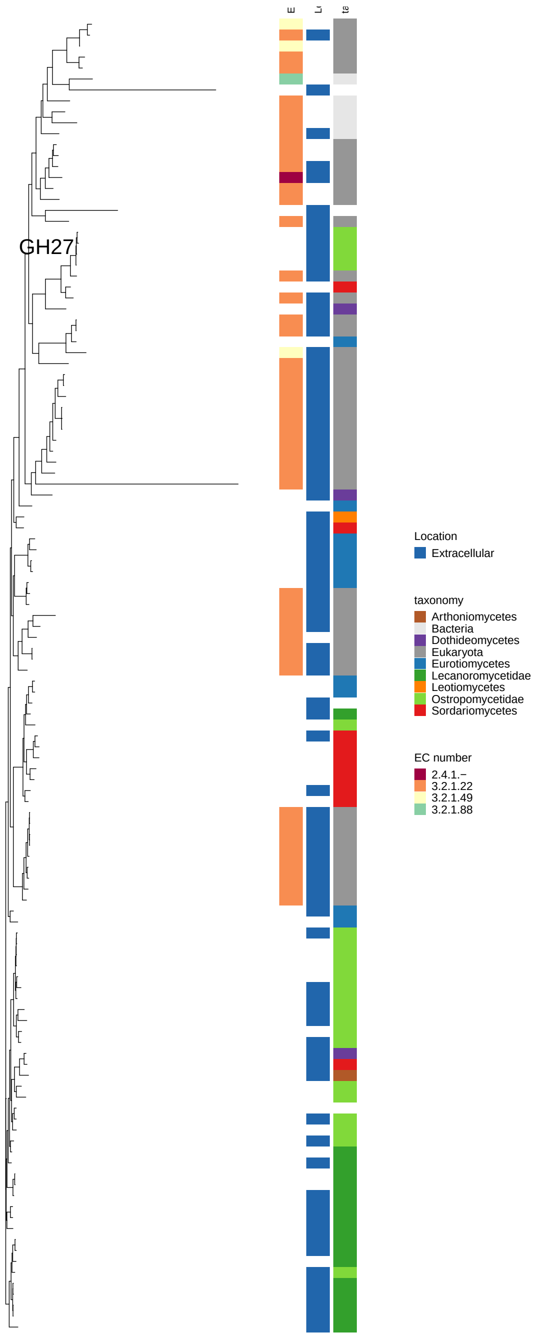

Figure 36: Maximum-likelihood gene tree of CAZyme families involved in (hemi-)cellulose, pectin and lignin breakdown. It includes all experimentally characterized sequences for the family downloaded from cazy.org as well as all sequences with annotations for this family in 83 genomes. The family name is given in the plot. The three columns along tree tips provide additional information about the respective sequence. Left column: Enzyme Code (EC), middle column: Predicted subcellular location, Right column: Taxonomy.

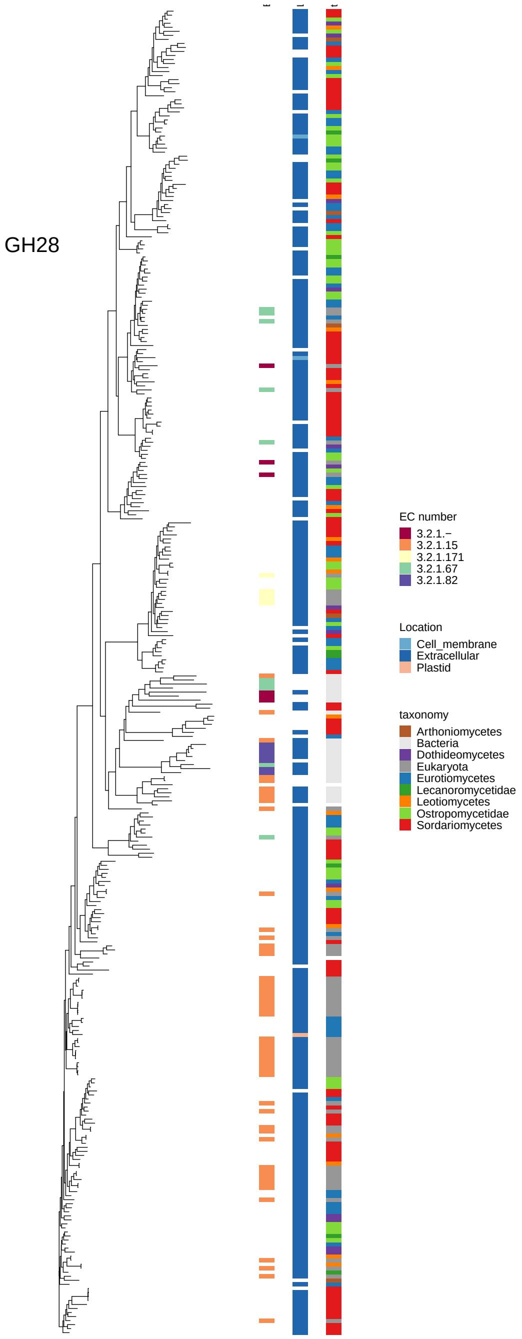

Figure 37: Maximum-likelihood gene tree of CAZyme families involved in (hemi-)cellulose, pectin and lignin breakdown. It includes all experimentally characterized sequences for the family downloaded from cazy.org as well as all sequences with annotations for this family in 83 genomes. The family name is given in the plot. The three columns along tree tips provide additional information about the respective sequence. Left column: Enzyme Code (EC), middle column: Predicted subcellular location, Right column: Taxonomy.

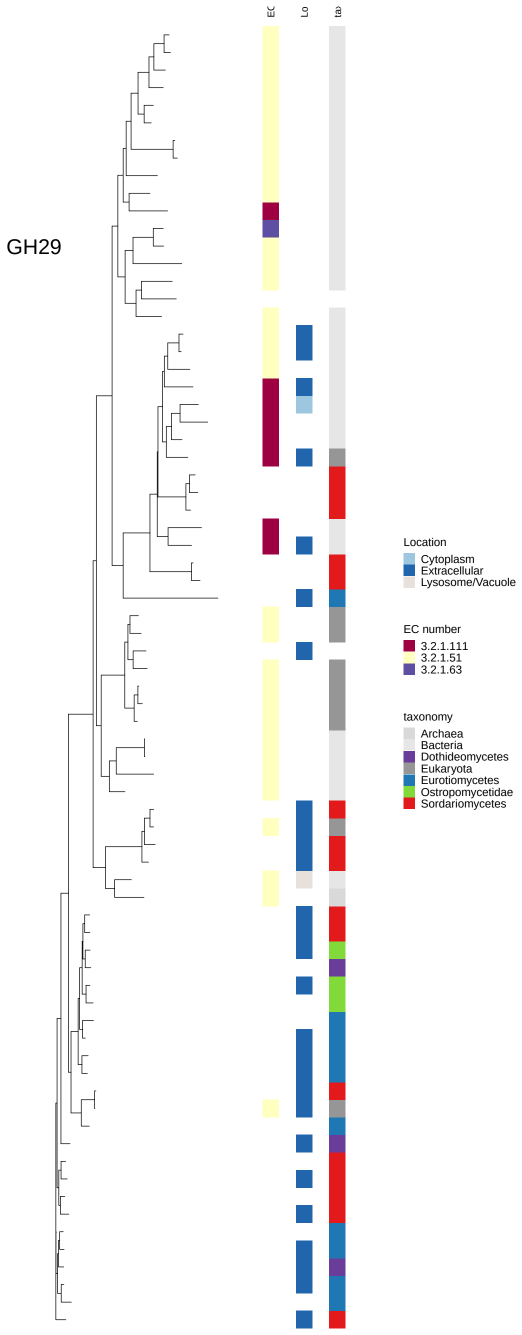

Figure 38: Maximum-likelihood gene tree of CAZyme families involved in (hemi-)cellulose, pectin and lignin breakdown. It includes all experimentally characterized sequences for the family downloaded from cazy.org as well as all sequences with annotations for this family in 83 genomes. The family name is given in the plot. The three columns along tree tips provide additional information about the respective sequence. Left column: Enzyme Code (EC), middle column: Predicted subcellular location, Right column: Taxonomy.

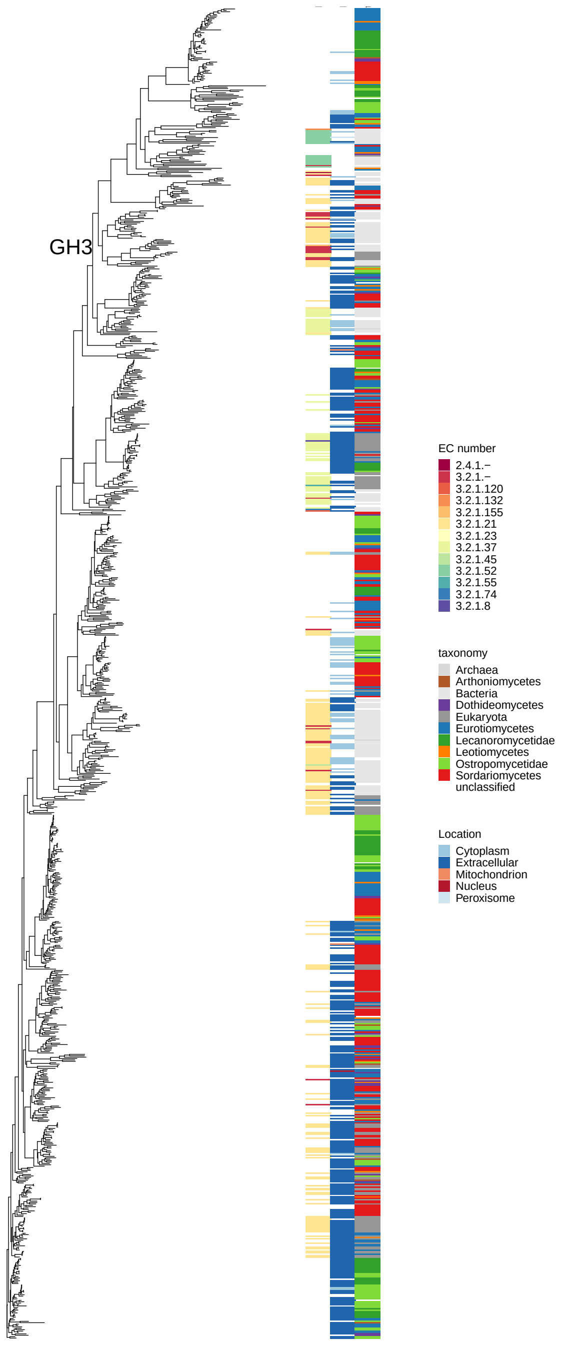

Figure 39: Maximum-likelihood gene tree of CAZyme families involved in (hemi-)cellulose, pectin and lignin breakdown. It includes all experimentally characterized sequences for the family downloaded from cazy.org as well as all sequences with annotations for this family in 83 genomes. The family name is given in the plot. The three columns along tree tips provide additional information about the respective sequence. Left column: Enzyme Code (EC), middle column: Predicted subcellular location, Right column: Taxonomy.

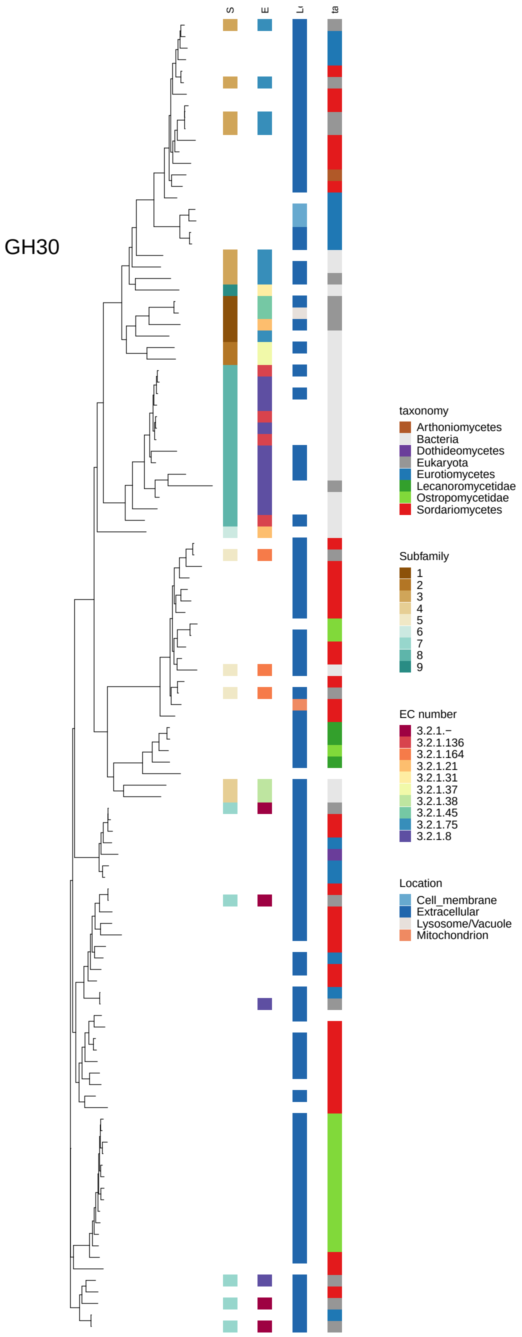

Figure 40: Maximum-likelihood gene tree of CAZyme families involved in (hemi-)cellulose, pectin and lignin breakdown. It includes all experimentally characterized sequences for the family downloaded from cazy.org as well as all sequences with annotations for this family in 83 genomes. The family name is given in the plot. The three columns along tree tips provide additional information about the respective sequence. Left column: Enzyme Code (EC), middle column: Predicted subcellular location, Right column: Taxonomy.

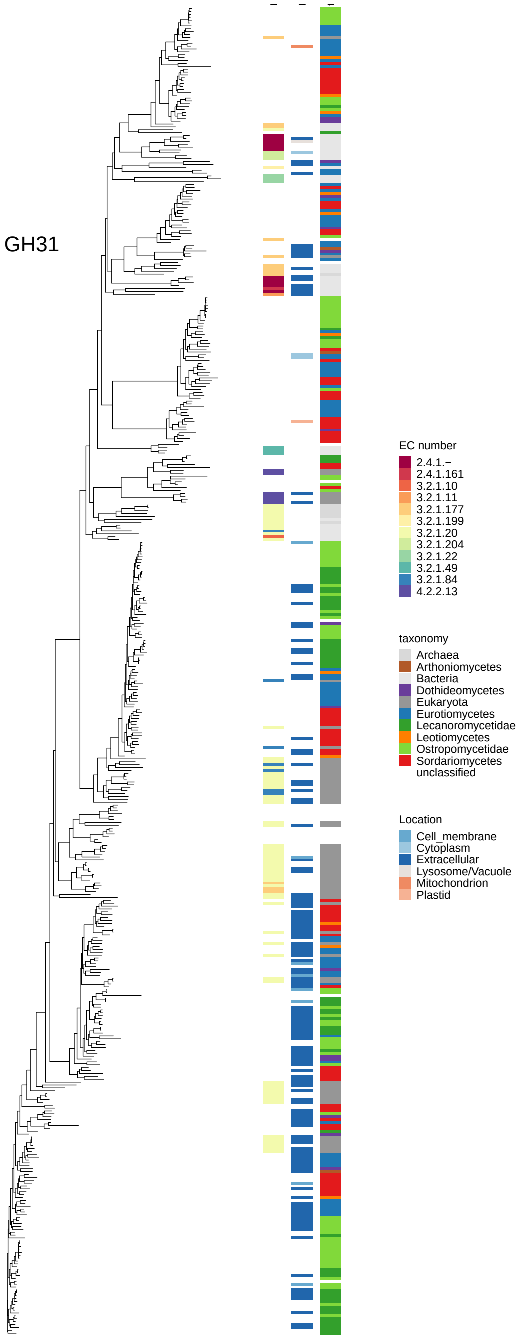

Figure 41: Maximum-likelihood gene tree of CAZyme families involved in (hemi-)cellulose, pectin and lignin breakdown. It includes all experimentally characterized sequences for the family downloaded from cazy.org as well as all sequences with annotations for this family in 83 genomes. The family name is given in the plot. The three columns along tree tips provide additional information about the respective sequence. Left column: Enzyme Code (EC), middle column: Predicted subcellular location, Right column: Taxonomy.

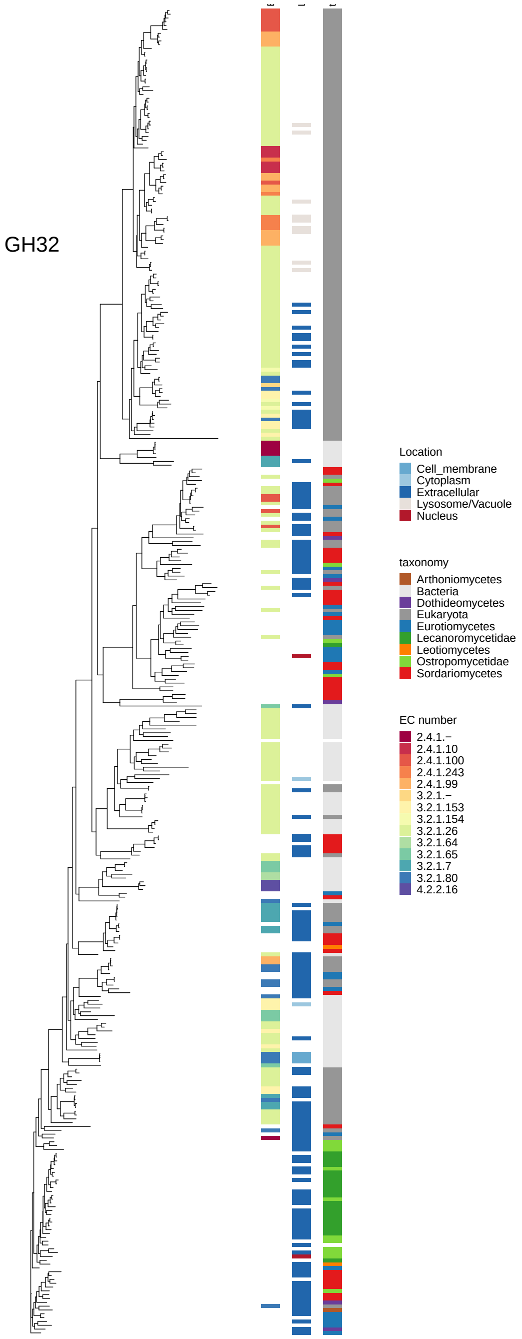

Figure 42: Maximum-likelihood gene tree of CAZyme families involved in (hemi-)cellulose, pectin and lignin breakdown. It includes all experimentally characterized sequences for the family downloaded from cazy.org as well as all sequences with annotations for this family in 83 genomes. The family name is given in the plot. The three columns along tree tips provide additional information about the respective sequence. Left column: Enzyme Code (EC), middle column: Predicted subcellular location, Right column: Taxonomy.

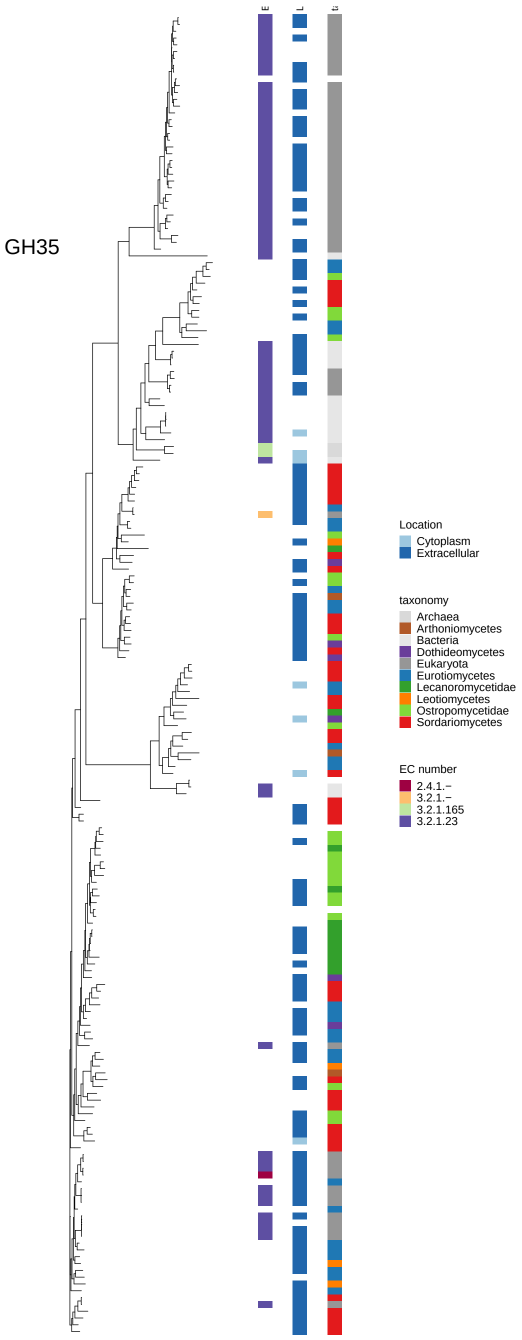

Figure 43: Maximum-likelihood gene tree of CAZyme families involved in (hemi-)cellulose, pectin and lignin breakdown. It includes all experimentally characterized sequences for the family downloaded from cazy.org as well as all sequences with annotations for this family in 83 genomes. The family name is given in the plot. The three columns along tree tips provide additional information about the respective sequence. Left column: Enzyme Code (EC), middle column: Predicted subcellular location, Right column: Taxonomy.

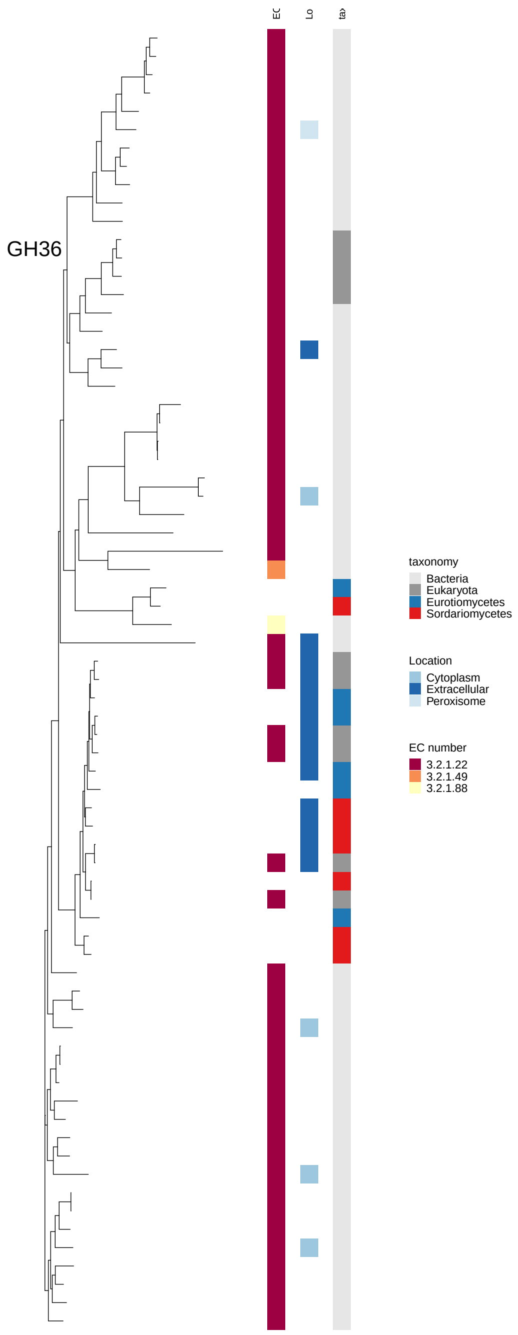

Figure 44: Maximum-likelihood gene tree of CAZyme families involved in (hemi-)cellulose, pectin and lignin breakdown. It includes all experimentally characterized sequences for the family downloaded from cazy.org as well as all sequences with annotations for this family in 83 genomes. The family name is given in the plot. The three columns along tree tips provide additional information about the respective sequence. Left column: Enzyme Code (EC), middle column: Predicted subcellular location, Right column: Taxonomy.

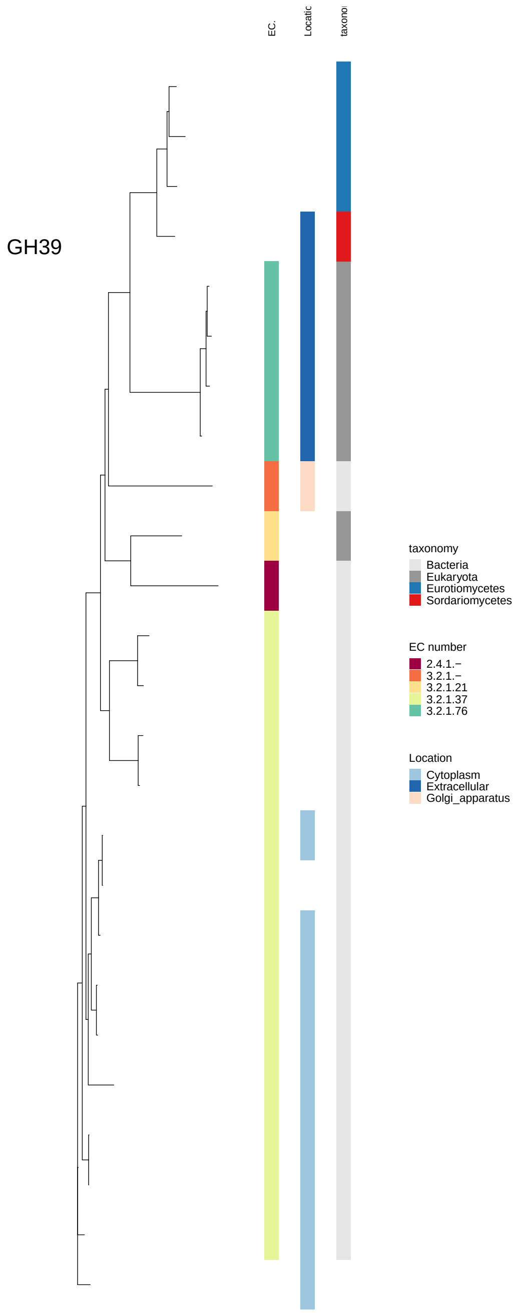

Figure 45: Maximum-likelihood gene tree of CAZyme families involved in (hemi-)cellulose, pectin and lignin breakdown. It includes all experimentally characterized sequences for the family downloaded from cazy.org as well as all sequences with annotations for this family in 83 genomes. The family name is given in the plot. The three columns along tree tips provide additional information about the respective sequence. Left column: Enzyme Code (EC), middle column: Predicted subcellular location, Right column: Taxonomy.

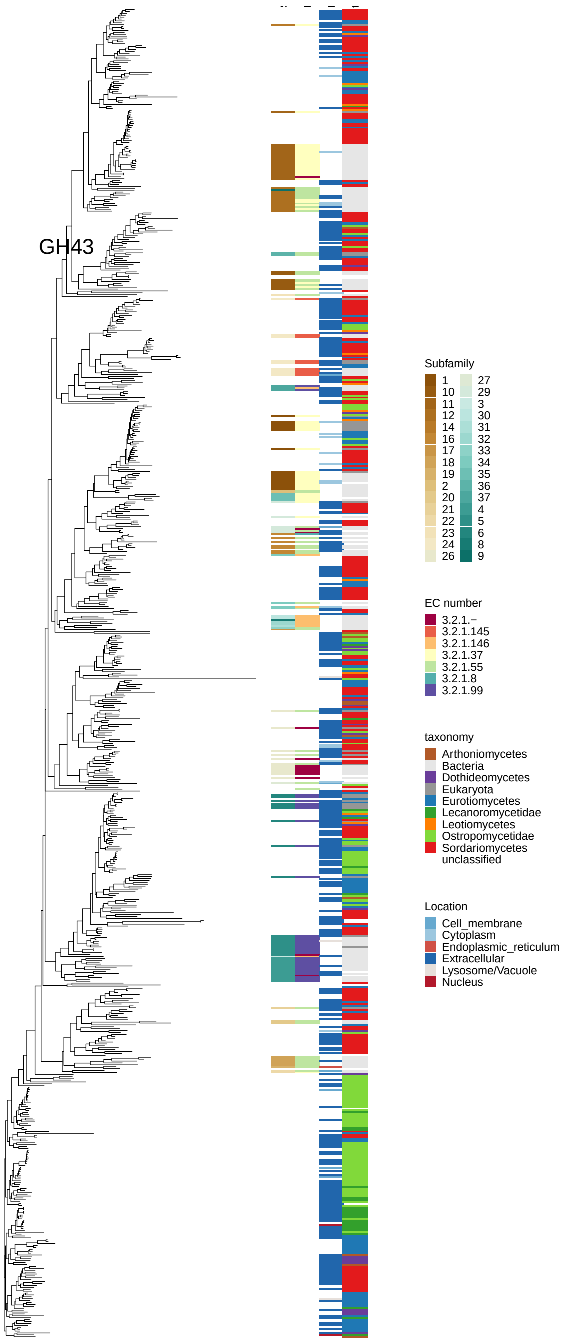

Figure 46: Maximum-likelihood gene tree of CAZyme families involved in (hemi-)cellulose, pectin and lignin breakdown. It includes all experimentally characterized sequences for the family downloaded from cazy.org as well as all sequences with annotations for this family in 83 genomes. The family name is given in the plot. The three columns along tree tips provide additional information about the respective sequence. Left column: Enzyme Code (EC), middle column: Predicted subcellular location, Right column: Taxonomy.

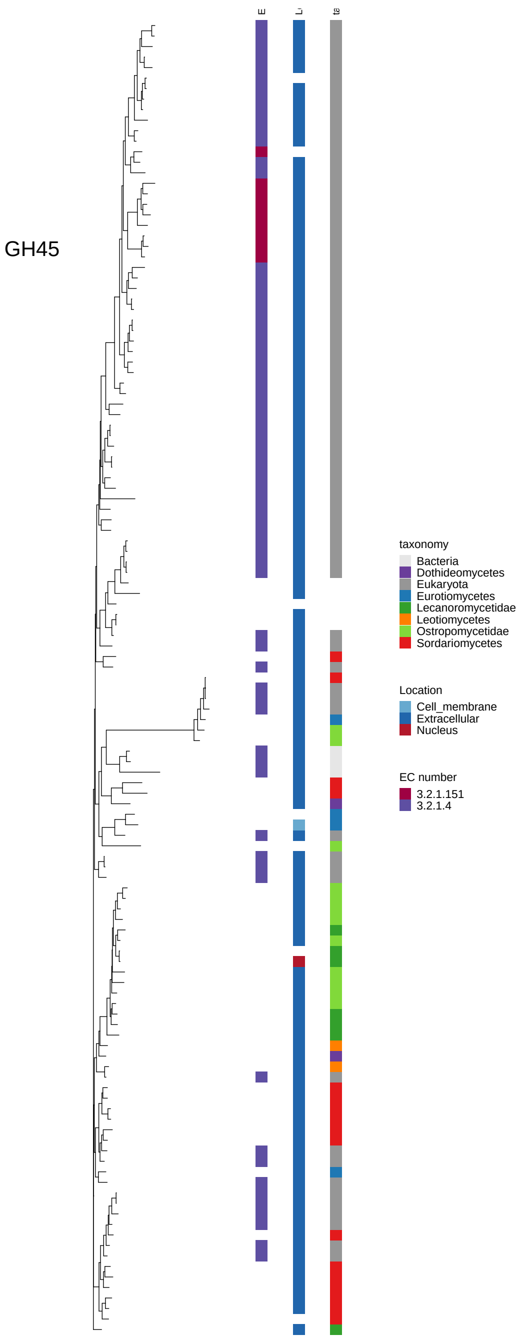

Figure 47: Maximum-likelihood gene tree of CAZyme families involved in (hemi-)cellulose, pectin and lignin breakdown. It includes all experimentally characterized sequences for the family downloaded from cazy.org as well as all sequences with annotations for this family in 83 genomes. The family name is given in the plot. The three columns along tree tips provide additional information about the respective sequence. Left column: Enzyme Code (EC), middle column: Predicted subcellular location, Right column: Taxonomy.

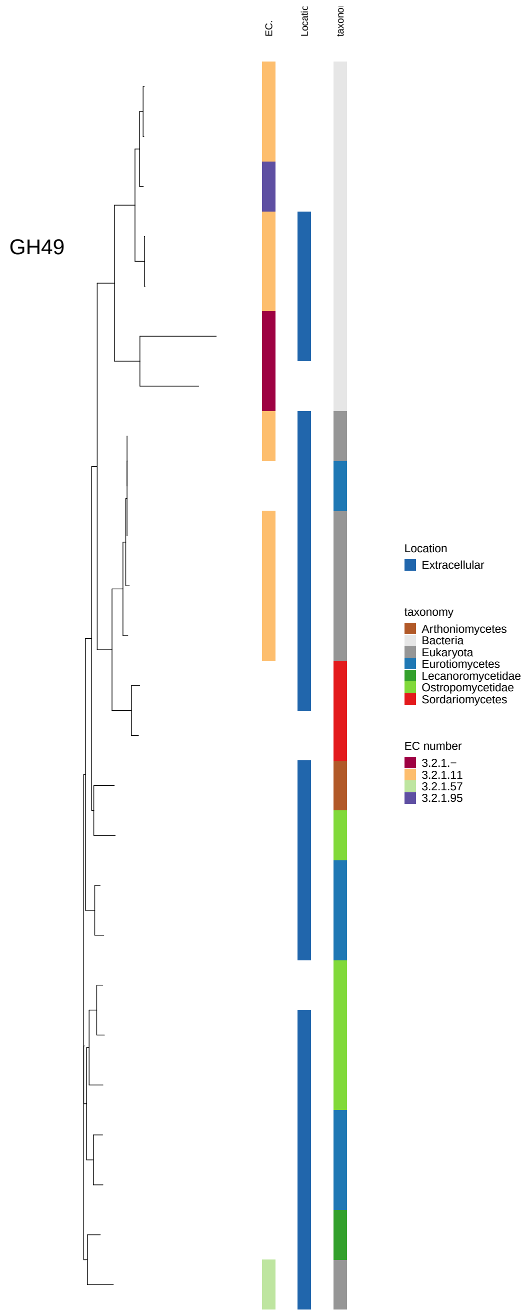

Figure 48: Maximum-likelihood gene tree of CAZyme families involved in (hemi-)cellulose, pectin and lignin breakdown. It includes all experimentally characterized sequences for the family downloaded from cazy.org as well as all sequences with annotations for this family in 83 genomes. The family name is given in the plot. The three columns along tree tips provide additional information about the respective sequence. Left column: Enzyme Code (EC), middle column: Predicted subcellular location, Right column: Taxonomy.

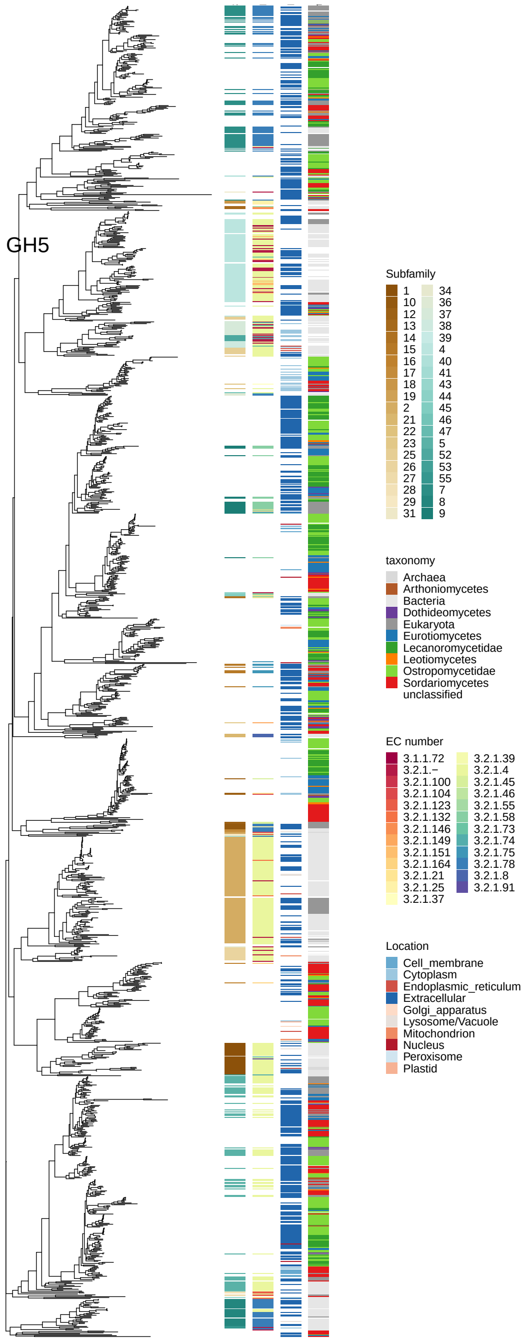

Figure 49: Maximum-likelihood gene tree of CAZyme families involved in (hemi-)cellulose, pectin and lignin breakdown. It includes all experimentally characterized sequences for the family downloaded from cazy.org as well as all sequences with annotations for this family in 83 genomes. The family name is given in the plot. The three columns along tree tips provide additional information about the respective sequence. Left column: Enzyme Code (EC), middle column: Predicted subcellular location, Right column: Taxonomy.

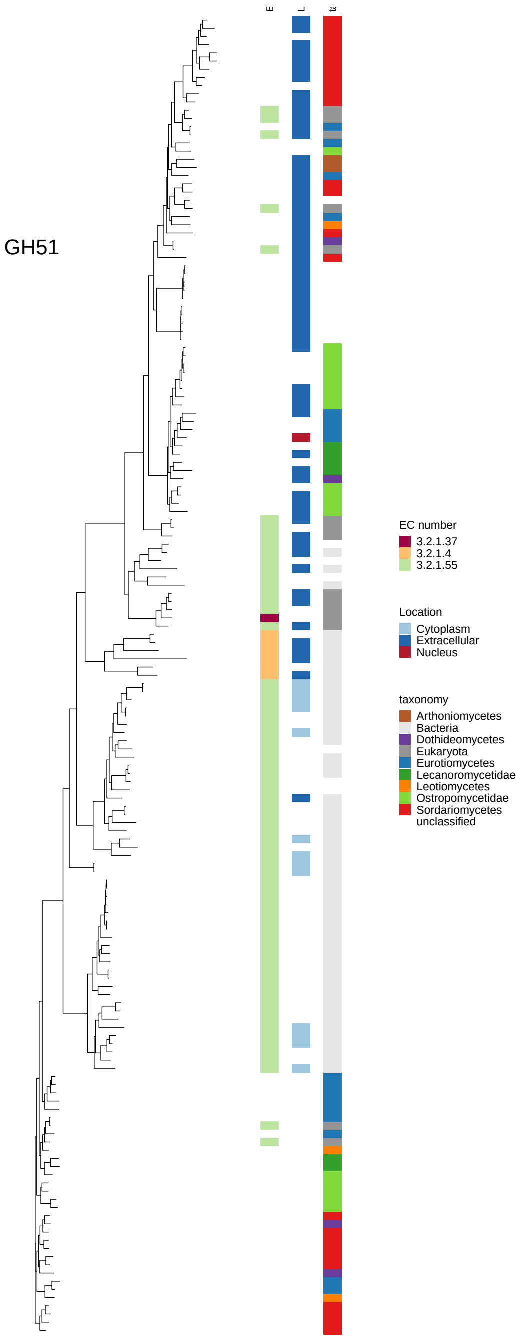

Figure 50: Maximum-likelihood gene tree of CAZyme families involved in (hemi-)cellulose, pectin and lignin breakdown. It includes all experimentally characterized sequences for the family downloaded from cazy.org as well as all sequences with annotations for this family in 83 genomes. The family name is given in the plot. The three columns along tree tips provide additional information about the respective sequence. Left column: Enzyme Code (EC), middle column: Predicted subcellular location, Right column: Taxonomy.

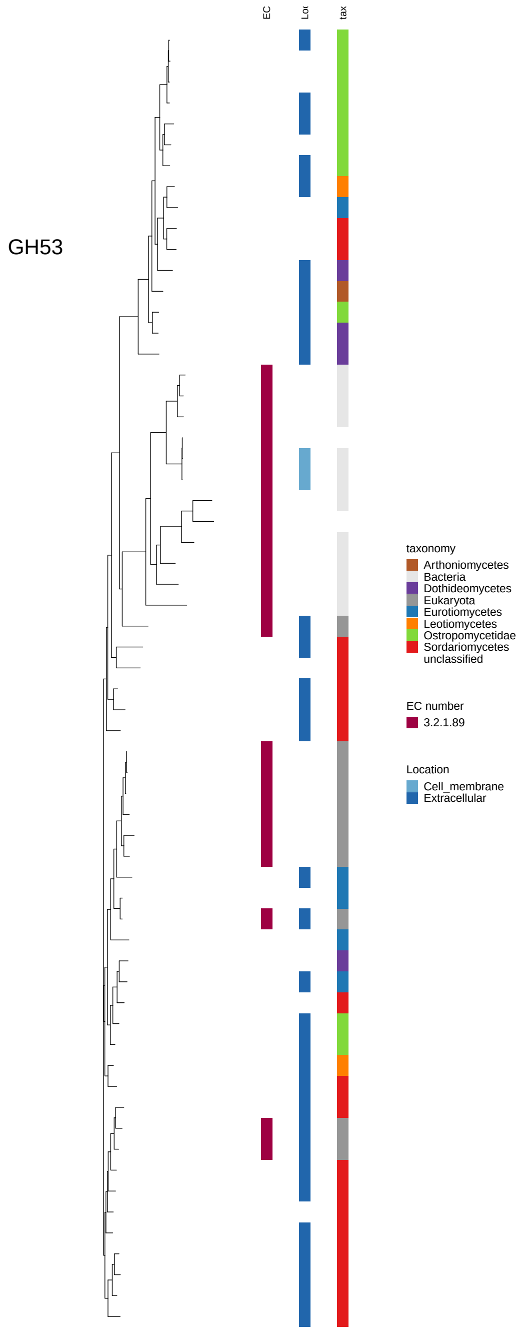

Figure 51: Maximum-likelihood gene tree of CAZyme families involved in (hemi-)cellulose, pectin and lignin breakdown. It includes all experimentally characterized sequences for the family downloaded from cazy.org as well as all sequences with annotations for this family in 83 genomes. The family name is given in the plot. The three columns along tree tips provide additional information about the respective sequence. Left column: Enzyme Code (EC), middle column: Predicted subcellular location, Right column: Taxonomy.

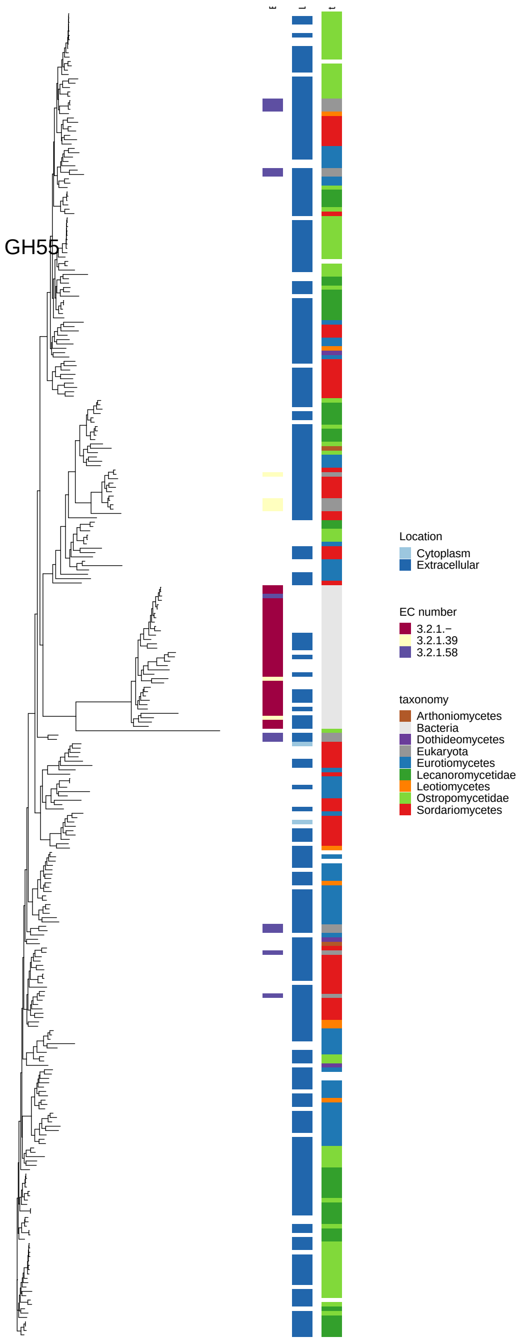

Figure 52: Maximum-likelihood gene tree of CAZyme families involved in (hemi-)cellulose, pectin and lignin breakdown. It includes all experimentally characterized sequences for the family downloaded from cazy.org as well as all sequences with annotations for this family in 83 genomes. The family name is given in the plot. The three columns along tree tips provide additional information about the respective sequence. Left column: Enzyme Code (EC), middle column: Predicted subcellular location, Right column: Taxonomy.

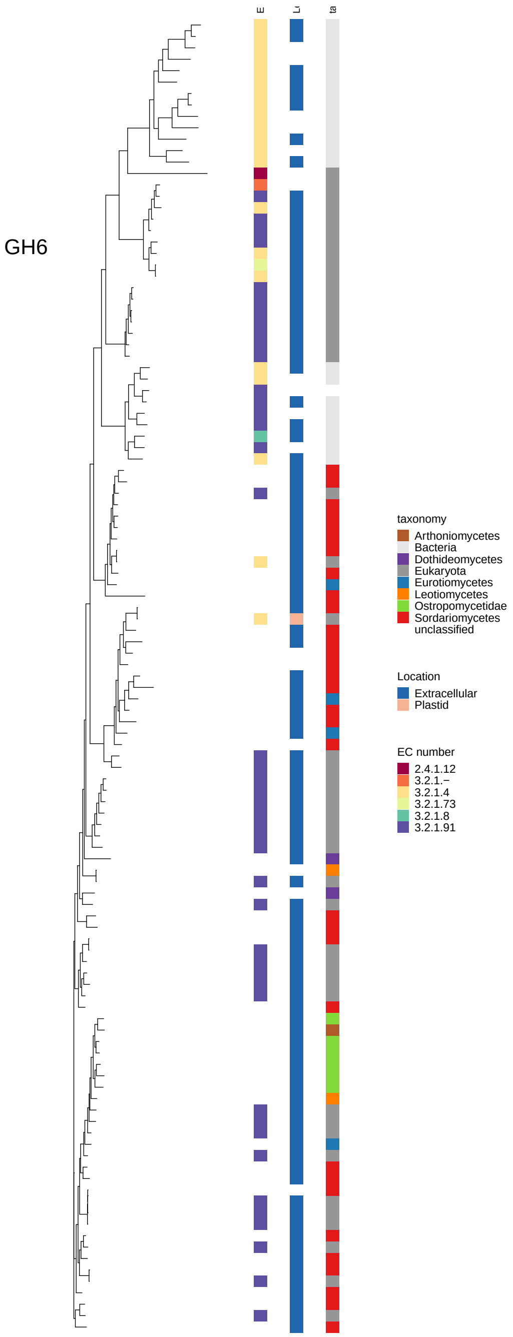

Figure 53: Maximum-likelihood gene tree of CAZyme families involved in (hemi-)cellulose, pectin and lignin breakdown. It includes all experimentally characterized sequences for the family downloaded from cazy.org as well as all sequences with annotations for this family in 83 genomes. The family name is given in the plot. The three columns along tree tips provide additional information about the respective sequence. Left column: Enzyme Code (EC), middle column: Predicted subcellular location, Right column: Taxonomy.

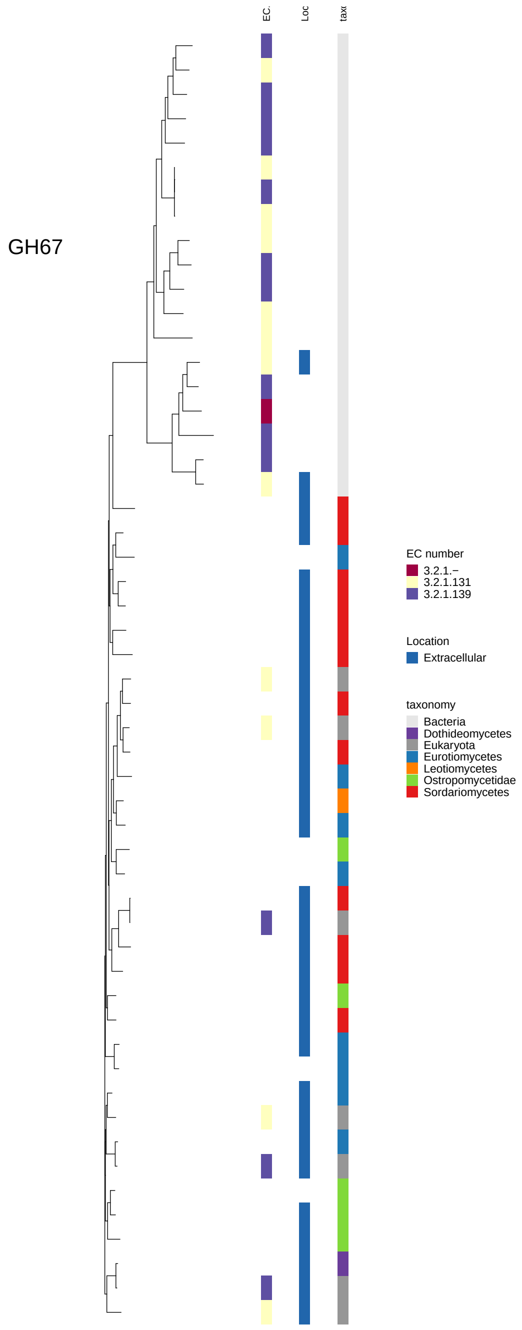

Figure 54: Maximum-likelihood gene tree of CAZyme families involved in (hemi-)cellulose, pectin and lignin breakdown. It includes all experimentally characterized sequences for the family downloaded from cazy.org as well as all sequences with annotations for this family in 83 genomes. The family name is given in the plot. The three columns along tree tips provide additional information about the respective sequence. Left column: Enzyme Code (EC), middle column: Predicted subcellular location, Right column: Taxonomy.

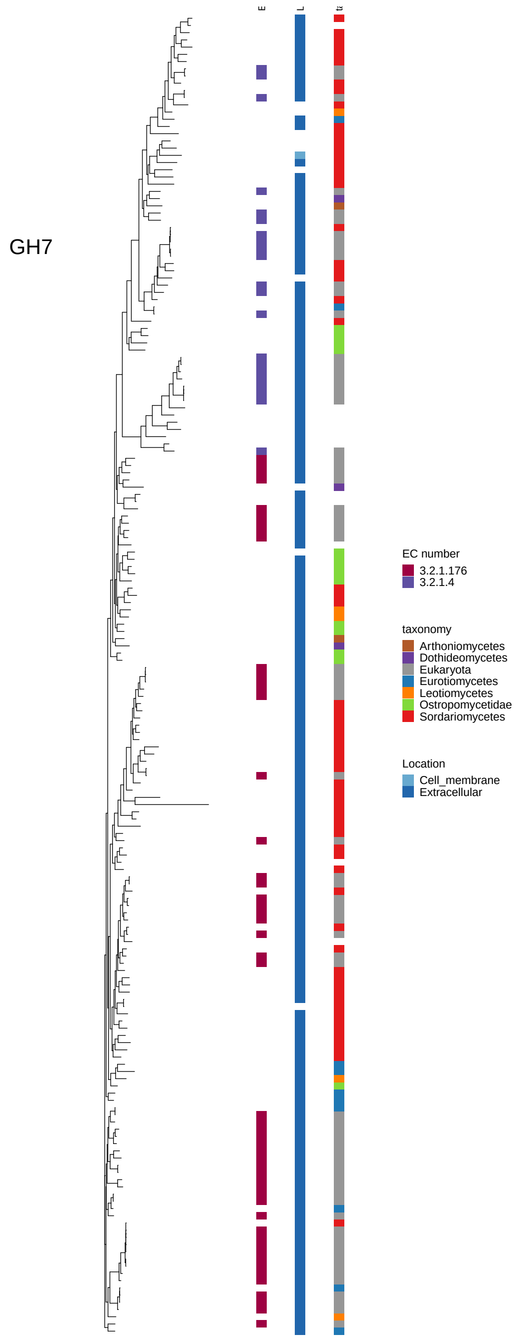

Figure 55: Maximum-likelihood gene tree of CAZyme families involved in (hemi-)cellulose, pectin and lignin breakdown. It includes all experimentally characterized sequences for the family downloaded from cazy.org as well as all sequences with annotations for this family in 83 genomes. The family name is given in the plot. The three columns along tree tips provide additional information about the respective sequence. Left column: Enzyme Code (EC), middle column: Predicted subcellular location, Right column: Taxonomy.

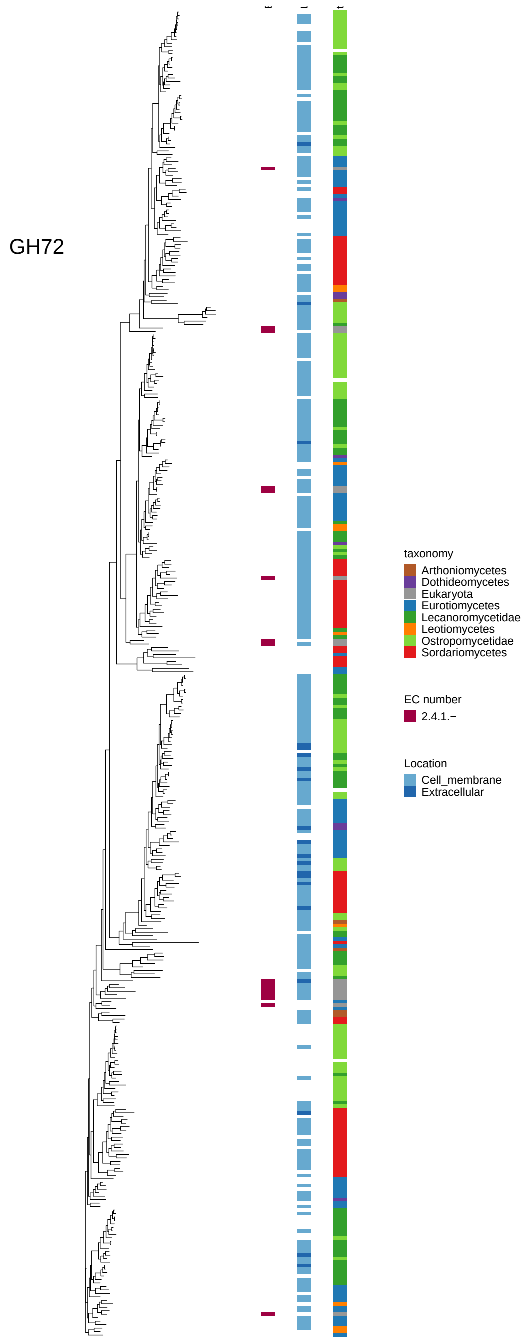

Figure 56: Maximum-likelihood gene tree of CAZyme families involved in (hemi-)cellulose, pectin and lignin breakdown. It includes all experimentally characterized sequences for the family downloaded from cazy.org as well as all sequences with annotations for this family in 83 genomes. The family name is given in the plot. The three columns along tree tips provide additional information about the respective sequence. Left column: Enzyme Code (EC), middle column: Predicted subcellular location, Right column: Taxonomy.

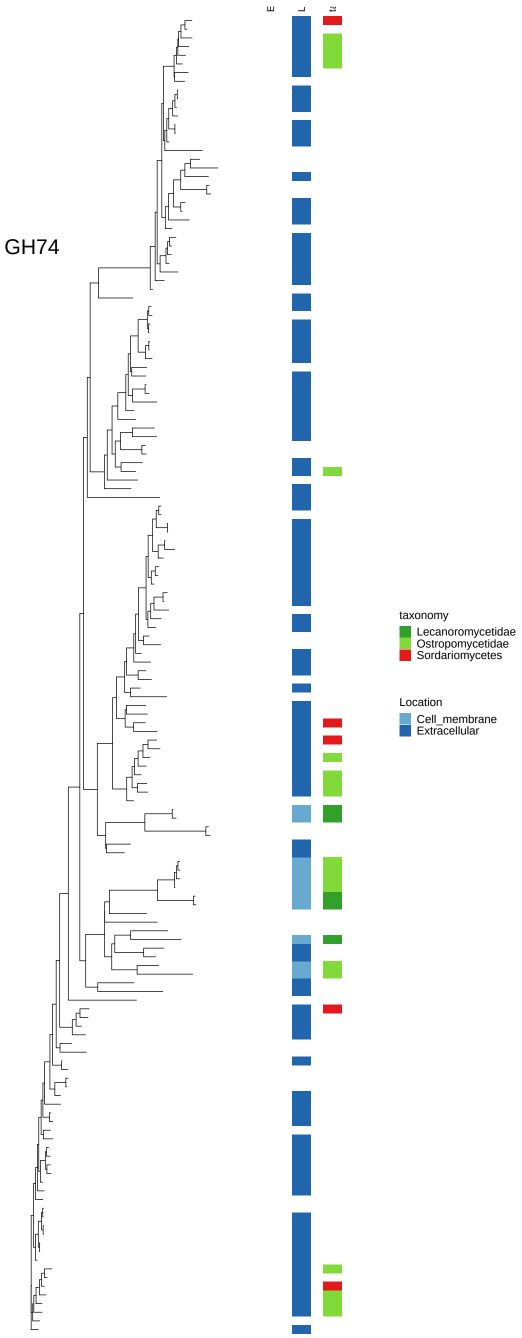

Figure 57: Maximum-likelihood gene tree of CAZyme families involved in (hemi-)cellulose, pectin and lignin breakdown. It includes all experimentally characterized sequences for the family downloaded from cazy.org as well as all sequences with annotations for this family in 83 genomes. The family name is given in the plot. The three columns along tree tips provide additional information about the respective sequence. Left column: Enzyme Code (EC), middle column: Predicted subcellular location, Right column: Taxonomy.

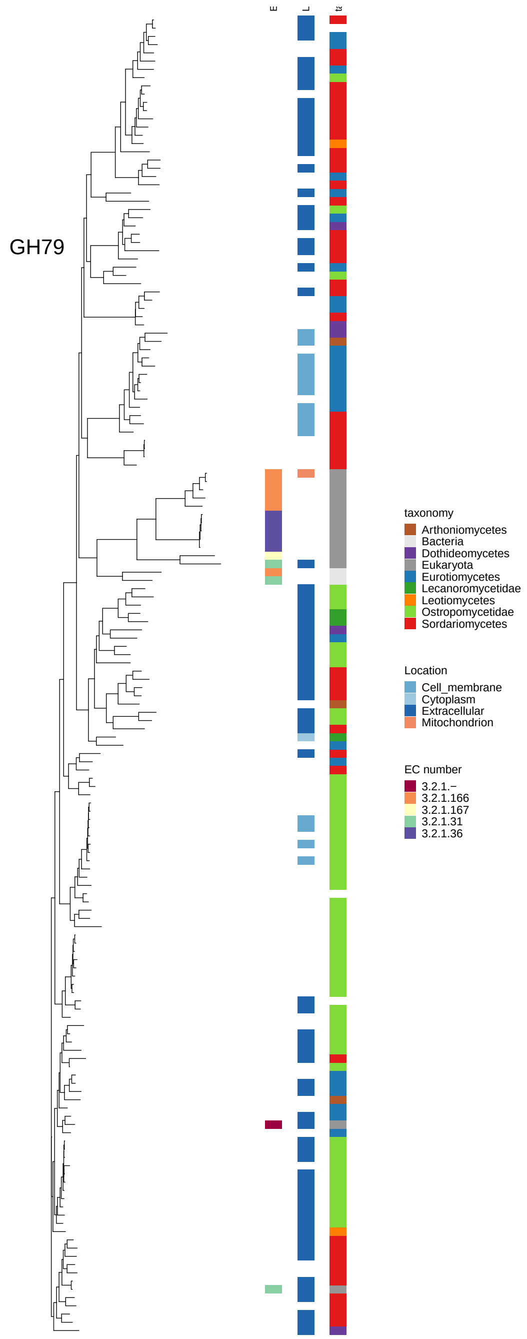

Figure 58: Maximum-likelihood gene tree of CAZyme families involved in (hemi-)cellulose, pectin and lignin breakdown. It includes all experimentally characterized sequences for the family downloaded from cazy.org as well as all sequences with annotations for this family in 83 genomes. The family name is given in the plot. The three columns along tree tips provide additional information about the respective sequence. Left column: Enzyme Code (EC), middle column: Predicted subcellular location, Right column: Taxonomy.

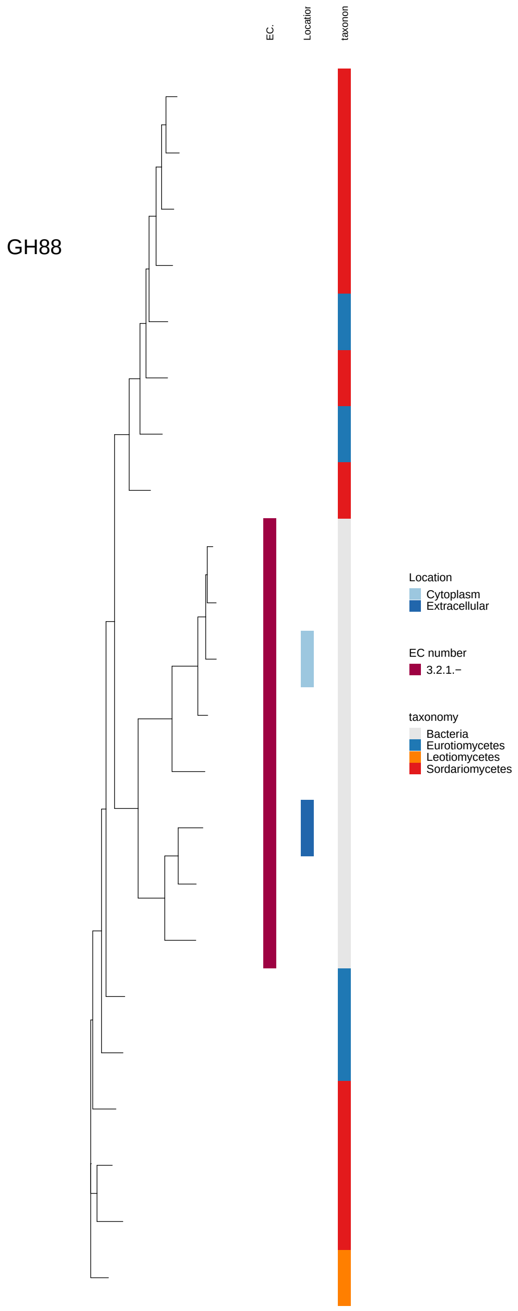

Figure 59: Maximum-likelihood gene tree of CAZyme families involved in (hemi-)cellulose, pectin and lignin breakdown. It includes all experimentally characterized sequences for the family downloaded from cazy.org as well as all sequences with annotations for this family in 83 genomes. The family name is given in the plot. The three columns along tree tips provide additional information about the respective sequence. Left column: Enzyme Code (EC), middle column: Predicted subcellular location, Right column: Taxonomy.

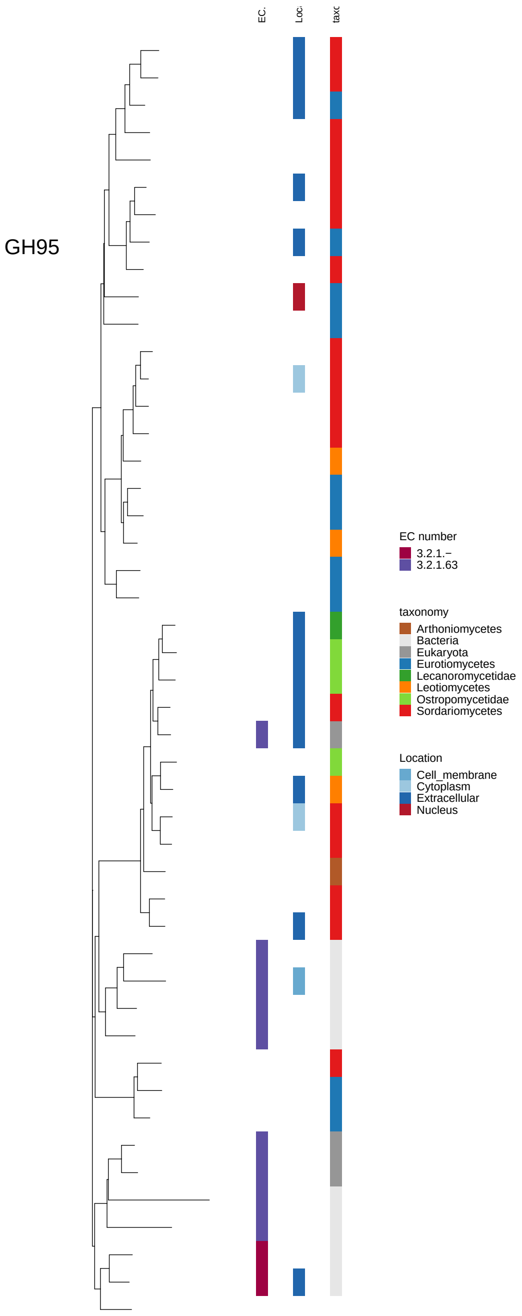

Figure 60: Maximum-likelihood gene tree of CAZyme families involved in (hemi-)cellulose, pectin and lignin breakdown. It includes all experimentally characterized sequences for the family downloaded from cazy.org as well as all sequences with annotations for this family in 83 genomes. The family name is given in the plot. The three columns along tree tips provide additional information about the respective sequence. Left column: Enzyme Code (EC), middle column: Predicted subcellular location, Right column: Taxonomy.

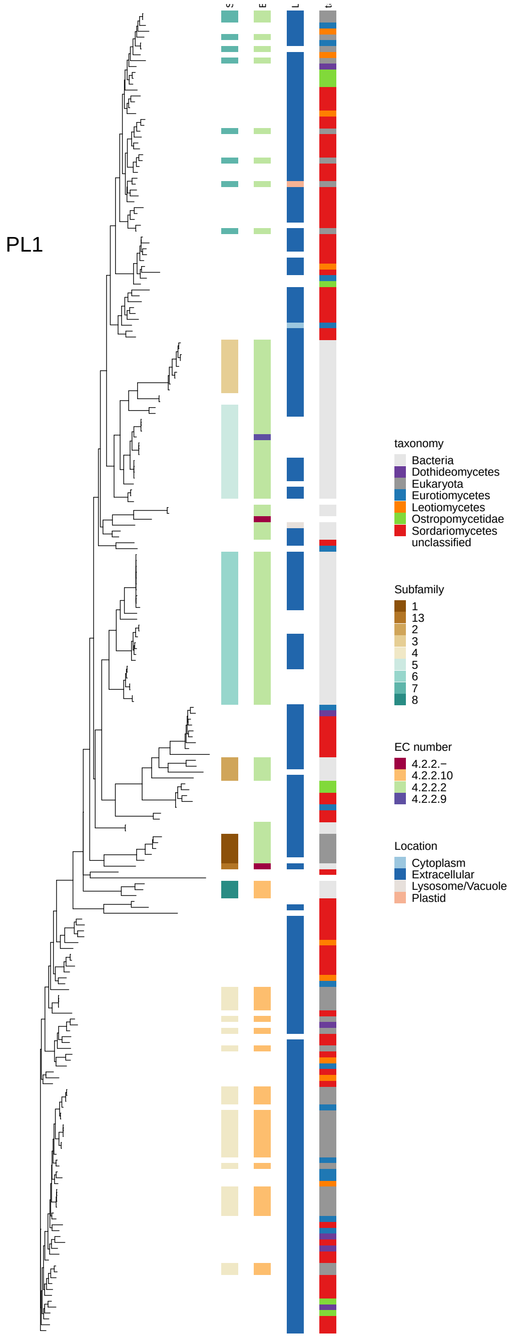

Figure 61: Maximum-likelihood gene tree of CAZyme families involved in (hemi-)cellulose, pectin and lignin breakdown. It includes all experimentally characterized sequences for the family downloaded from cazy.org as well as all sequences with annotations for this family in 83 genomes. The family name is given in the plot. The three columns along tree tips provide additional information about the respective sequence. Left column: Enzyme Code (EC), middle column: Predicted subcellular location, Right column: Taxonomy.

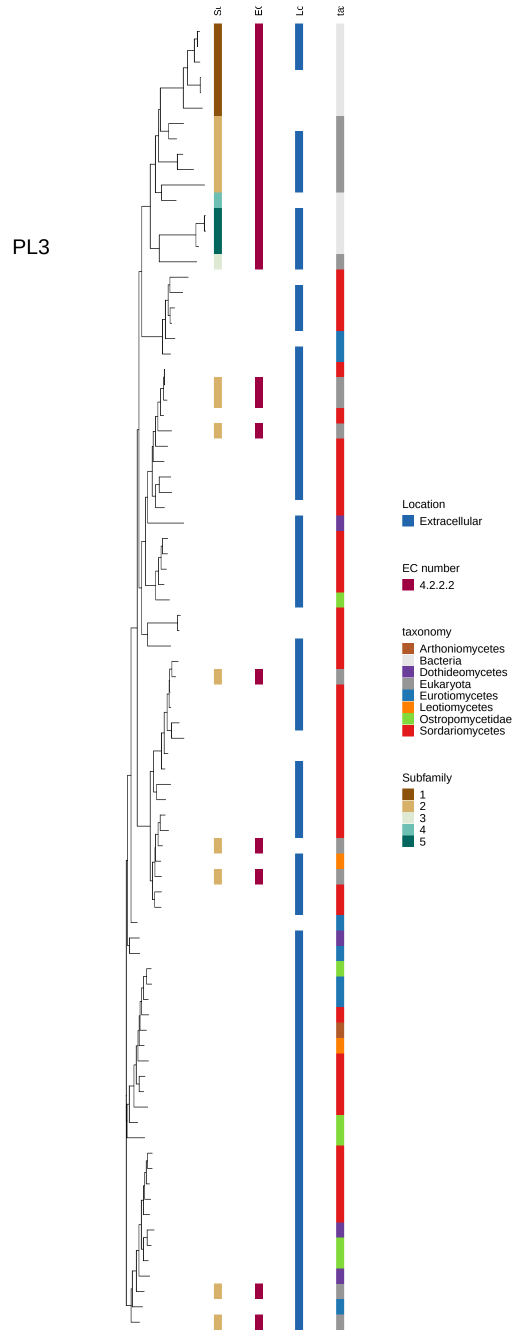

Figure 62: Maximum-likelihood gene tree of CAZyme families involved in (hemi-)cellulose, pectin and lignin breakdown. It includes all experimentally characterized sequences for the family downloaded from cazy.org as well as all sequences with annotations for this family in 83 genomes. The family name is given in the plot. The three columns along tree tips provide additional information about the respective sequence. Left column: Enzyme Code (EC), middle column: Predicted subcellular location, Right column: Taxonomy.

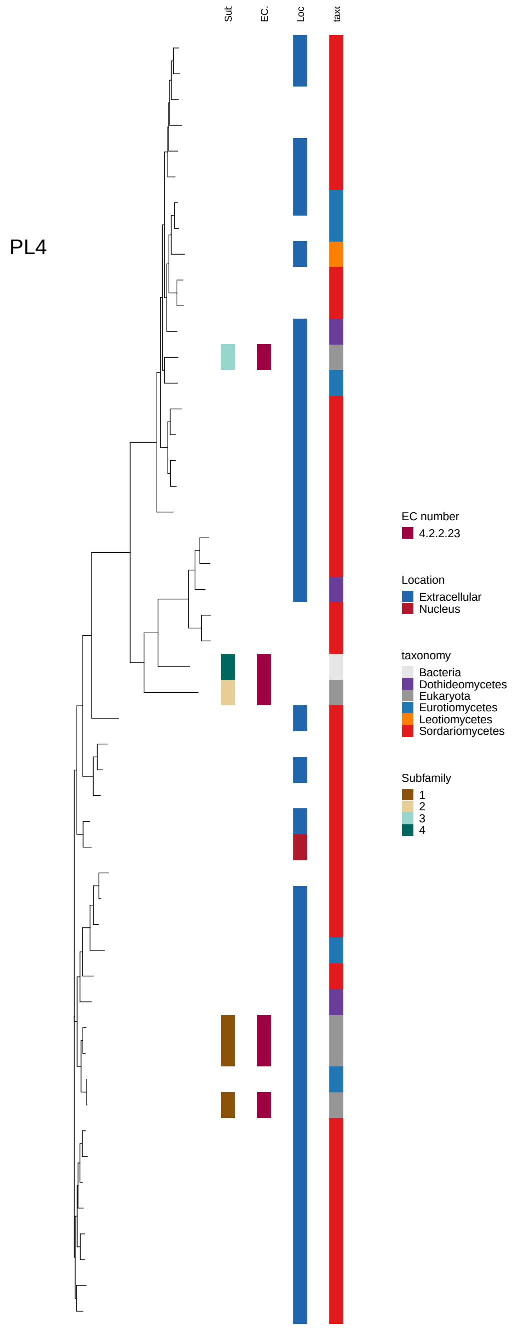

Figure 63: Maximum-likelihood gene tree of CAZyme families involved in (hemi-)cellulose, pectin and lignin breakdown. It includes all experimentally characterized sequences for the family downloaded from cazy.org as well as all sequences with annotations for this family in 83 genomes. The family name is given in the plot. The three columns along tree tips provide additional information about the respective sequence. Left column: Enzyme Code (EC), middle column: Predicted subcellular location, Right column: Taxonomy.

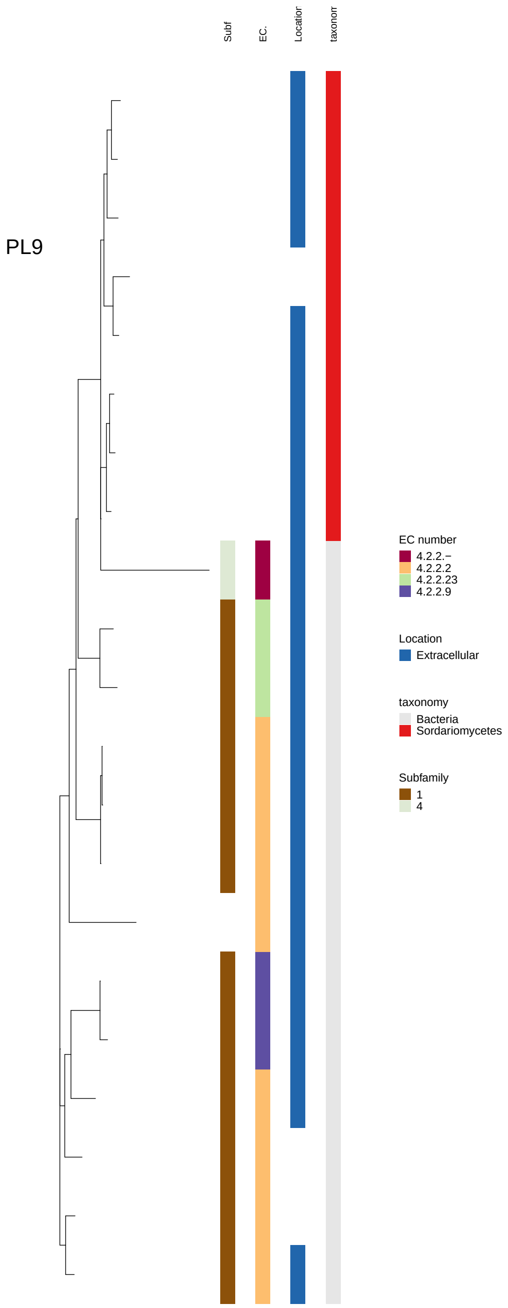

Figure 64: Maximum-likelihood gene tree of CAZyme families involved in (hemi-)cellulose, pectin and lignin breakdown. It includes all experimentally characterized sequences for the family downloaded from cazy.org as well as all sequences with annotations for this family in 83 genomes. The family name is given in the plot. The three columns along tree tips provide additional information about the respective sequence. Left column: Enzyme Code (EC), middle column: Predicted subcellular location, Right column: Taxonomy.

Chikhi, Rayan, and Guillaume Rizk. 2012. "Space-Efficient and Exact de Bruijn Graph Representation Based on a Bloom Filter." In *WABI.*

Edgar, Robert C. 2004. "MUSCLE: multiple sequence alignment with high accuracy and high throughput." *Nucleic Acids Research.*

Emms, David M, and Steven Kelly. 2019. "OrthoFinder: phylogenetic orthology inference for comparative genomics."

Floudas, D, M Binder, R Riley, Kerrie Barry, Robert A Blanchette, B Henrissat, Angel T Martıénez, et al. 2012. "The Paleozoic origin of enzymatic lignin decomposition reconstructed from 31 fungal genomes." *Science.*

"GitHub - reslp/binner: script for running multiple metagenome filtering programs using Docker containers." 2020.

Gurevich, A, V Saveliev, N Vyahhi, and G Tesler. 2013. "QUAST: quality assessment tool for genome assemblies." *Bioinformatics.*

Henrissat, B, M Claeyssens, P Tomme, L Lemesle, and J P Mornon. 1989. "Cellulase Families Revealed by Hydrophobic Cluster-Analysis." *Gene.*

Hoang, D T, O Chernomor Molecular biology, and 2017. n.d. "UFBoot2: improving the ultrafast bootstrap approximation." *Molecular Biology and Evolution.*

Huang, X, N Duan, H Xu, T N Xie, Y R Xue, and C H Liu. 2018. "CTAB-PEG DNA Extraction from Fungi with High Contents of Polysaccharides." *Molecular Biology.*

Huerta-Cepas, J, K Forslund, Luis Pedro Coelho, Damian Szklarczyk, Lars Juhl Jensen, Christian von Mering, and Peer Bork. 2017. "Fast Genome-Wide Functional Annotation through Orthology Assignment by eggNOG-Mapper." *Molecular Biology and Evolution.*

Jackman, Shaun D, Benjamin P Vandervalk, Hamid Mohamadi, Justin Chu, Sarah Yeo, S Austin Hammond, Golnaz Jahesh, et al. 2017. "ABySS 2.0: resource-efficient assembly of large genomes using a Bloom filter." *Genome Research.*

Jones, Darryl R, Dallas Thomas, Nicholas Alger, Ata Ghavidel, G Douglas Inglis, and D Wade Abbott. 2018. "SACCHARIS: an automated pipeline to streamline discovery of carbohydrate active enzyme activities within polyspecific families and de novo sequence datasets." *Biotechnology for Biofuels.*

Jones, P, D Binns, H-Y Chang, M Fraser, W Li, C McAnulla, Hamish McWilliam, et al. 2014. "InterProScan 5: genome-scale protein function classification." *Bioinformatics.*

Kajitani, Rei, Kouta Toshimoto, Hideki Noguchi, Atsushi Toyoda, Yoshitoshi Ogura, Miki Okuno, Mitsuru Yabana, et al. 2014. "Efficient de novo assembly of highly heterozygous genomes from whole-genome shotgun short reads." *Genome Research.*

Katoh, K, and D M Standley. 2013. "MAFFT multiple sequence alignment software version 7: improvements in performance and usability." *Molecular Biology and Evolution.*

Kohler, A, Alan Kuo, Laszlo G Nagy, Emmanuelle Morin, Kerrie W Barry, Francois Buscot, Björn Canbäck, et al. 2015. "Convergent losses of decay mechanisms and rapid turnover of symbiosis genes in mycorrhizal mutualists." *Nature Genetics.*

Korf, I. 2004. "Gene finding in novel genomes." *BMC Bioinformatics.*

Laetsch, Dominik R, and Mark L Blaxter. 2017. "BlobTools: Interrogation of genome assemblies." *F1000Research.*

Lowe, T M, and S R Eddy. 1997. "tRNAscan-SE: a program for improved detection of transfer RNA genes in genomic sequence." *Nucleic Acids Research.*

Majoros, W H, M Pertea, and S L Salzberg. 2004. "TigrScan and GlimmerHMM: two open source ab initio eukaryotic gene-finders." *Bioinformatics.*

Mathé, Catherine, Nizar Fawal, Christophe Roux, and Christophe Dunand. 2019. "In silico definition of new ligninolytic peroxidase sub-classes in fungi and putative relation to fungal life style." *Scientific Reports.*

Mendes, Fábio K, Dan Vanderpool, Ben Fulton, and M W Hahn. 2020. "CAFE 5 models variation in evolutionary rates among gene families." *Bioinformatics.*

Mewis, Keith, Nicolas Lenfant, V Lombard, and B Henrissat. 2016. "Dividing the Large Glycoside Hydrolase Family 43 into Subfamilies: a Motivation for Detailed Enzyme Characterization." *Applied and Environmental Microbiology.*

Minh, Bui Quang, M W Hahn, and R Lanfear. 2020. "New Methods to Calculate Concordance Factors for Phylogenomic Datasets." *Molecular Biology and Evolution.*

Minh, Bui Quang, Heiko A Schmidt, Olga Chernomor, Dominik Schrempf, Michael D Woodhams, Arndt von Haeseler, and R Lanfear. 2020. "IQ-TREE 2: New Models and Efficient Methods for Phylogenetic Inference in the Genomic Era." *Molecular Biology and Evolution.*

Mirarab, S, R Reaz, Md S Bayzid, T Zimmermann, M S Swenson, and T Warnow. 2014. "ASTRAL: genome-scale coalescent-based species tree estimation." *Bioinformatics.*

Mistry, Jaina, R D Finn, S R Eddy, Alex Bateman, and Marco Punta. 2013. "Challenges in homology search: HMMER3 and convergent evolution of coiled-coil regions." *Nucleic Acids Research.*

Miyauchi, Shingo, Enikő Kiss, Alan Kuo, Elodie Drula, A Kohler, Marisol Sánchez-Garcıéa, Emmanuelle Morin, et al. 2020. "Large-scale genome sequencing of mycorrhizal fungi provides insights into the early evolution of symbiotic traits." *Nature Communications.*

Molina, M del Carmen, and A Crespo. 2000. "Comparison of development of axenic cultures of five species of lichen-forming fungi." *Mycological Research.*

Palmer, Jon, and Jason Stajich. 2021. "nextgenusfs/funannotate: funannotate v1.8.7. Available online: https://github.com/nextgenusfs/funannotate."

Pogoda, Cloe S, Kyle G Keepers, J C Lendemer, Nolan C Kane, and Erin A Tripp. 2018. "Reductions in Complexity of Mitochondrial Genomes in Lichen-Forming Fungi Shed Light on Genome Architecture of Obligate Symbioses." *Molecular Ecology.*

Price, Morgan N, Paramvir S Dehal, and Adam P Arkin. 2010. "FastTree 2-Approximately Maximum-Likelihood Trees for Large Alignments." *PLoS ONE.*

Resl, P, and C Hahn. 2021. "phylociraptor: rapid phylogenomic tree calculator. Available online: https://github.com/reslp/phylociraptor."

Revell, L J. 2009. "Size-correction and principal components for interspecific comparative studies." *Evolution.*

———. 2013. "A comment on the use of stochastic character maps to estimate evolutionary rate variation in a continuously valued trait." *Systematic Biology.*

Salichos, L, and A Rokas. 2013. "Inferring ancient divergences requires genes with strong phylogenetic signals." *Nature.*

Sanderson, M J. 2003. "r8s: inferring absolute rates of molecular evolution and divergence times in the absence of a molecular clock." *Bioinformatics.*

Sievers, Fabian, Andreas Wilm, David Dineen, Toby J Gibson, Kevin Karplus, W Li, Rodrigo Lopez, et al. 2011. "Fast, scalable generation of high-quality protein multiple sequence alignments using Clustal Omega." *Molecular Systems Biology.*

Smit, AFA, and R Hubley. n.d. "RepeatModeler Open-1.0. Available online: http://www.repeatmasker.org."

Smit, AFA, R Hubley, and P Green. 2013. "*RepeatMasker Open-4.0*. Available online: http://www.repeatmasker.org."

Stamatakis, A. 2014. "RAxML version 8: A tool for Phylogenetic Analysis and Post-Analysis of Large Phylogenies." *Bioinformatics.*

Stanke, M, O Schöffmann, B Morgenstern, and S Waack. 2006. "Gene prediction in eukaryotes with a generalized hidden Markov model that uses hints from external sources." *BMC Bioinformatics.*

Ter-Hovhannisyan, V, A Lomsadze, Y O Chernoff, and M Borodovsky. 2008. "Gene prediction in novel fungal genomes using an ab initio algorithm with unsupervised training." *Genome Research.*

Uritskiy, Gherman V, Jocelyne DiRuggiero, and James Taylor. 2018. "MetaWRAPa flexible pipeline for genome-resolved metagenomic data analysis."

Waterhouse, R M, Mathieu Seppey, F A Simão, Mosè Manni, P Ioannidis, Guennadi Klioutchnikov, E V Kriventseva, and E M Zdobnov. 2017. "BUSCO Applications from Quality Assessments to Gene Prediction and Phylogenomics." *Molecular Biology and Evolution.*

Wickham, H. 2016. *ggplot2: Elegant Graphics for Data Analysis.* Springer-Verlag New York.

Zerbino, D R, and E Birney. 2008. "Velvet: Algorithms for de novo short read assembly using de Bruijn graphs." *Genome Research.*
